# Evolution of thermal tolerance in marine diatoms: Metabolic strategies under heat stress

**DOI:** 10.1101/2024.11.27.625584

**Authors:** María Aranguren-Gassis, Angel P. Diz, María Huete-Ortega, Andrew Allen, Elena Litchman

## Abstract

In the last decade, numerous laboratory experiments have demonstrated that when marine phytoplankton are exposed to thermal stress, they can evolve high temperature tolerance in a short time (weeks to months). This evolutionary potential may ensure the persistence of marine phytoplankton species under current and future global warming. However, the effect of such adaptation on the phytoplankton interaction with the environment and other organisms depends on how cellular metabolism shifts during the evolutionary process. In order to elucidate which cellular strategies allow the emergence of thermo-tolerant populations, we analyzed the proteomics response of a marine diatom (*Chaetoceros simplex*) to both thermal acclimation and evolutionary adaptation. We found that high temperature-tolerant populations exhibit a conservative cellular strategy when acclimated to high, above-optimal temperature, where recycling and reallocation is favored at the expense of new structures’ biosynthesis. While this strategy gives the populations that evolved high temperature tolerance an advantage under thermal stress, the shift to resource reallocation may explain the absence of high-temperature adaptation when cells are exposed to low nitrate availability.

## INTRODUCTION

Temperature is key selective agent for marine species, including diatoms (Koester *et al,* 2013), and its importance is increasing under the ongoing climate change. Ocean is warming at a non-precedented rate (IPCC, 2021), and temperature increase is already affecting marine species distribution and extinction rates (Barton *et al.,* 2016; Lenoir *et al,* 2020, Pinsky *et al,* 2019 and 2020). This effect could be less dramatic for unicellular organisms, such as phytoplankton, which may be able to readily adapt to novel conditions due to their short generation times, large population numbers, and typically high genetic diversity (Rynearson & Armbrust 2000,and 2005). Thermal adaptation in unicellular organisms shifts their cellular strategies allowing species to persist in new environments (Schaum *et al,* 2018a). Those metabolic changes can modify the adapted cells’ interaction with the environment and other organisms (Aranguren-Gassis *et al,* 2019; Padfield *et al,* 2015; Schaum *et al,* 2018b; Yvon-Durocher *et al,* 2015) but are not well characterized. Because phytoplankton form the base of most food webs in the ocean and drive biogeochemical cycles (Falkowski *et al,* 2008; Yvon-Durocher *et al,* 2017), we must unravel the cellular mechanisms that drive phytoplankton adaptation to predict its impact on marine ecosystems under the global change.

The short-term response (acclimation) of phytoplankton to elevated temperatures has been widely studied, but long-term adaptation may be different (Boyd *et al.,* 2018; Chan *et al.,* 2021 and Collins *et al.,* 2020 for reviews), and plasticity itself can evolve (Collins *et al.,* 2020). From the current literature, some generalities on phytoplankton adaptation to thermal stress emerge, for example, adaptive changes involve such traits as cell growth rate, nutrient and polyunsaturated fatty acid content, photo-physiological performance or extracellular level (O’Donnell *et al.,* 2019, 2021, Chan *et al.,* 2021 and references therein). However, information on the cellular process and the reorganization of metabolic pathways that drive such physiological changes is scarce. Such information is crucial to envisage long-term responses in new environments where temperature and other environmental drivers can interact. By characterizing cellular mechanisms behind thermal adaptation, we can infer potential trade-offs and synergies with responses to other drivers that improve our understanding of adaptive outcomes and constraints.

To better understand the cellular mechanisms driving phytoplankton adaptation to high temperatures, we analyzed proteomic response of high temperature-resistant populations to an above-optimal temperature. We obtained those populations during an evolution experiment with a marine diatom (Aranguren-Gassis *et al,* 2019). The results revealed specific cellular strategies under stressful thermal conditions that allowed the populations to persist. We discuss how shifts in metabolic pathways improved cells’ survival under high, previously lethal temperatures, and why the interaction with other environmental drivers, such as low nitrate availability, suppressed high temperature tolerance.

## MATERIALS AND METHODS

After evolving under above-optimal temperatures (31°C) for ∼200 generations, marine diatom populations became tolerant to extremely high temperatures (34°C) (Aranguren-Gassis *et al.,* 2019). Here we analyzed the proteome response of those lines that evolved extremely high temperature tolerance when they grew at control (25°C) and high temperature (34°C) to identify cellular processes that allowed such adaptation (figure 1). We compared their responses to the responses of ancestral populations and the populations that were maintained at benign temperature (25°C) when they grew at 25°C and high temperature (31°C) (figure 1).

**Figure 1:**
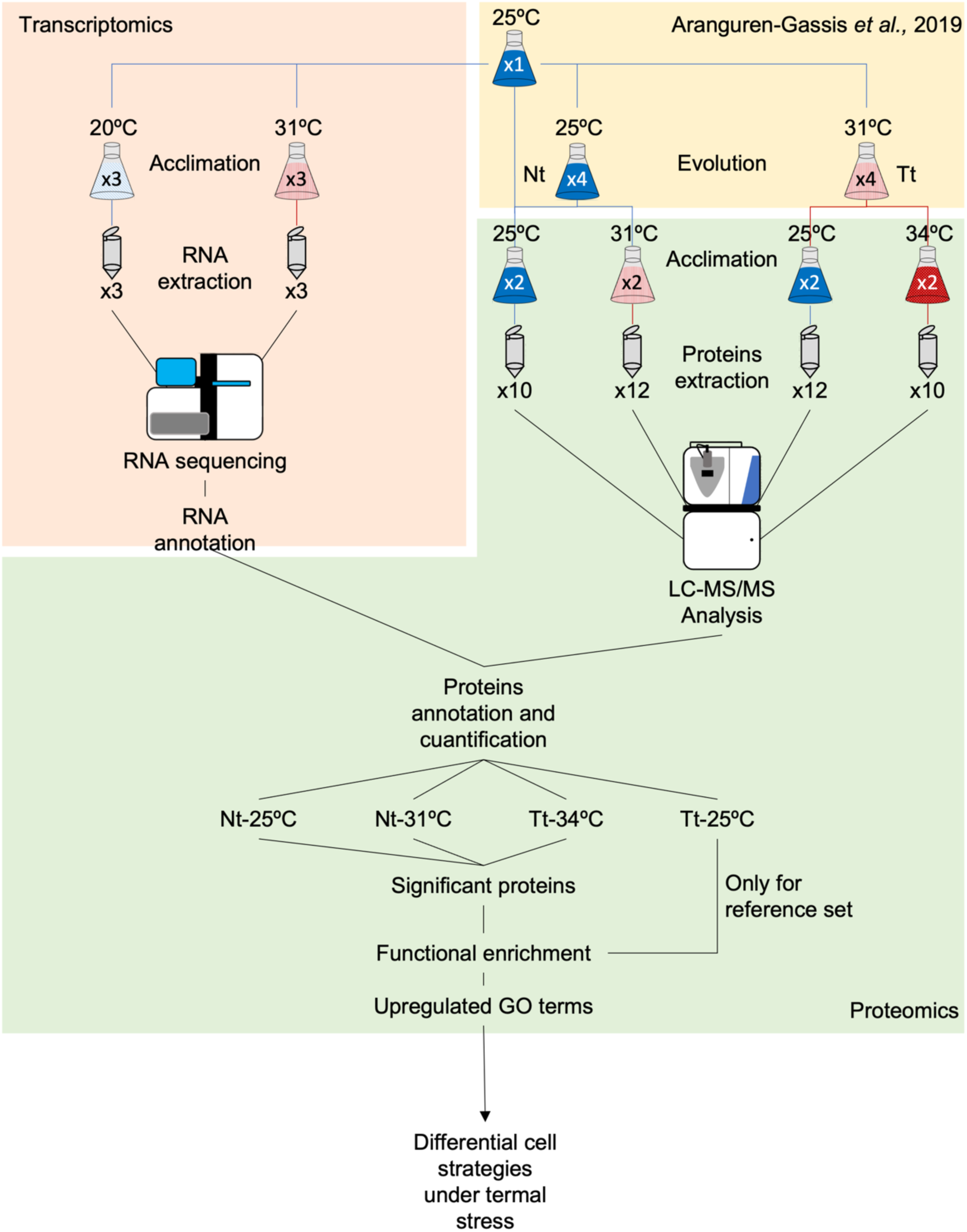
Methods workflow. The pink square on the upper left describes the transcriptomics analysis, the yellow square in the upper right describes the evolution experiment described in Aranguren-Gassis *et al*, 2019, the green area includes the proteomic analysis, and the white part on the bottom resumes the results. *Nt* means Non tolerant populations, *Tt* means thermo-tolerant populations. Numbers inside flasks with a *x* indicates the number of replicates.

### Culture conditions

The details of the evolution experiment are described in Aranguren-Gassis *et al*. (2019). Briefly, four populations of *Chaetoceros simplex* Ostenfeld (strain CCMP 200 from the National Center for Marine Algae and Microbiota, NCMA) were cultured during ∼200 generations under above-optimum temperature (31°C), and one population was maintained at 25°C as a control. Additional replicates of the ancestral population were cryopreserved and defrosted as additional controls. All of them were maintained in regular nutrient-enriched artificial L1 seawater medium from NCMA, modified from Guillard and Hargraves (1993). Several of those populations adapted to warm conditions developed the ability to grow at high, previously lethal temperatures (34°C).

#### 1- Proteomics

Two evolved populations were randomly selected from the ones that developed high-temperature (34°C) tolerance (thermo-tolerant, Tt populations, see Box 1), and other two populations were randomly selected from the control replicates (nontolerant, Nt populations, see Box 1), one maintained at 25°C and one from the cryopreserved ancestral populations. Those populations were grown at the ancestral optimum temperature (25°C) and at the above-optimal temperatures (31°C for the Nt and 34°C for the Tt, see table 1). 34°C was selected as a stressful over-optimal temperature for the Tt populations at which the Nt populations were not able to grow (Aranguren-Gassis *et al,* 2019). Populations were maintained in the same L1 medium specified before, in 500 ml glass flasks, at 100 μmol quanta m^-2^ s^-1^ cool white fluorescent light on the 14/10 hour day/night cycle. We gently swirled and randomly repositioned flasks daily. Every three days, 5-10 ml from each population were transferred to fresh medium. We monitored populations by measuring *in vivo* chlorophyll-*a* fluorescence daily (excitation wavelength: 436 nm, emission wavelength: 680 nm) using a SpectraMax M5 microplate reader (Molecular Devices, Sunnyvale, CA).

**Table 1:**
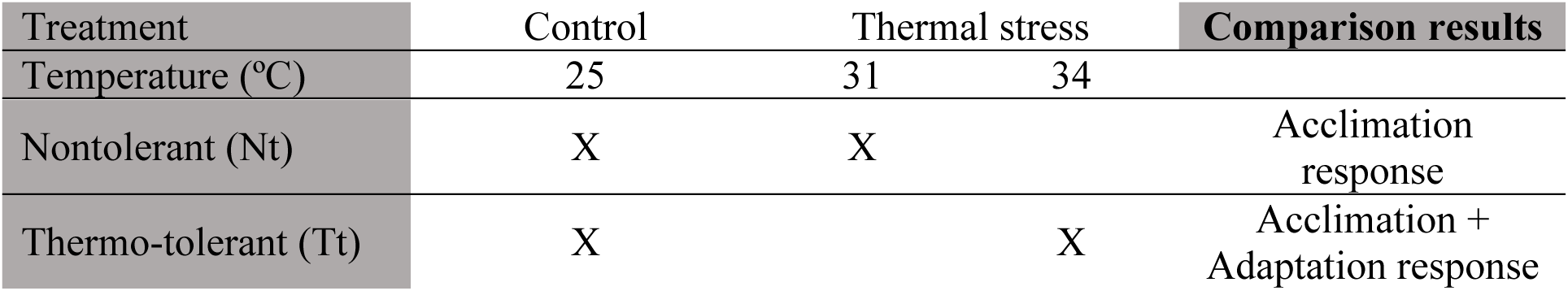
Temperature growth treatments for proteomic analyses. The final column (Comparison results) shows how the comparison results of the checked treatments were interpreted.

After at least 2 days of acclimation, and while cultures were growing exponentially, 100 ml of each culture were filtered onto 0.2 μm pore size polycarbonate membrane filters with gentle vacuum pressure. The filters were immediately frozen in liquid nitrogen and preserved at −80°C until protein extraction.

We obtained 10-12 samples for each treatment:

- Two biological samples (those coming from different experimental strains) with 5 replicates (different filtration dates) each for control populations at 25°C (Nt25) and evolved populations at 34°C (Tt34).
- Four biological samples with 3 replicates each for control at 31°C (Nt31).
- Two biological samples with 5 replicates and two additional biological samples with only one replicate for the evolved populations at 25°C (Tt25).

#### 2- Transcriptomics

For transcriptomics, three replicates of the ancestral population were grown in similar conditions than described before, at 20 and 31°C. Filtration was made as described above for proteomics. The results from these transcriptomic analyses were used only for proteins annotation in this work.

### RNA extraction and analysis

Total RNA was extracted from filters using Macherey-Nagel NucleoMag RNA kit (Macherey-Nagel, Düren, Germany). Cleared lysate was loaded into a 96 deep-well plate and put on an epMotion 5075 TMX liquid handler (Eppendorf, Hamburg, Germany) to complete the RNA extraction following the Macherey-Nagel standard protocol. RNA quality was analyzed on a 2200 TapeStation System with Agilent RNA ScreenTape System (Agilent Technologies, Santa Clara, CA, USA) and quantified using Quant-iT RiboGreen RNA Assay Kit (ThermoFisher, Waltham, MA, USA). PolyA enrichment mRNA transcriptome was constructed using 400ng-1ug of total RNA as input into the Illumina TruSeq RNA Sample Preparation Kit (Illumina, San Diego, CA, USA), following the manufacture’s Low-Throughput protocol. Library quality was analyzed on a 2200 TapeStation System with Agilent High Sensitivity D1000 ScreenTape System (Agilent Technologies). Resulting libraries were subjected to paired-end sequencing via Illumina HiSeq4000.

### Proteins extraction

The equipment was washed before extraction, first with methanol, then with ethanol 70%, and finally with 100% ethanol. All reactants were prepared with HPLC grade water, and the tubes used for processing samples were low binding types. Frozen filters were washed with 0.5 M Tetraethylammonium bicarbonate (TEAB) buffer solution with 0.1% sodium dodecyl sulfate (SDS) detergent. The filtered biomass was ground in a precooled mortar with liquid nitrogen and 1ml 1% protease inhibitor solution (Thermo Fisher protease and phosphatase inhibition cocktail without EDTA). The ground samples were sonicated for 5 minutes with ice, and then 100 μl glass beads (Sigma acid washed glass beads 425-600 µm) were added to each sample. They were bead beaten 5 times, alternating 2 minutes beating and 2 minutes resting in ice. After beating, samples were centrifugated at 4°C and 18000g for 30 minutes. The supernatant was then transferred to new tubes and stored at −80°C.

### Proteomic analyses

#### 1- Quantification

Bicinchoninic acid (BCA) microplate assay (Thermo Scientific Pierce BCA Protein Assay Kit) was used for protein quantification with an albumin (Sigma Aldrich bovine serum albumin) calibration standard. For each sample, 15 μg protein was diluted in acetone to obtain 1 μg/μl protein concentration and stored at −20°C for 12 hours. Samples were then centrifugated (4°C 10000 g 10 minutes), and the pellets were resuspended in 15 μl of TEAB 20 mM solution.

#### 2- In-gel digestion of proteins

Protein samples containing 10 µg of total protein were loaded onto a standard Laemmli-type polyacrylamide gel and allowed to stack and enter the resolving gel, but not allowed to separate. Comassie blue stain was used. Then, gel pieces were excised and subjected to in-gel digestion. Briefly, gel pieces were washed sequentially with ammonium bicarbonate 25 mM and 50% ACN/ammonium bicarbonate 25 mM, in an ultrasonic bath; proteins were reduced by treatment with 10mM DTT for 1 hour, and alkylated with 55mM IAA for 30 min. Protein digestion was accomplished with 40 ng of trypsin at 37 °C overnight and tryptic peptides were extracted from the gel matrix in two steps with 0,5% TFA and 100% ACN.

#### 3- LC-MS/MS analysis

The tryptic peptides were dried in a speed-vacuum concentrator at 45°C (Concentrator plus, Eppendorf, Hamburg, Germany), reconstituted in LC/MS-grade water containing 0.1% (v/v) formic acid and analysed by electrospray ionization-tandem mass spectrometry on a hybrid high-resolution LTQ-Orbitrap Elite mass spectrometer coupled to a Proxeon Easy-nLC 1000 UHPLC system (ThermoFisher Scientific).

The peptides were delivered onto a reverse phase column (PepMap® RSLC C18, 2µm, 100 Å, 75µm x 50cm, Thermo Fisher Scientific) and were eluted with an ACN gradient of 5-30% containing 0.1% formic acid for 240 min. The elutes were directly delivered into the mass spectrometer, which was set in positive ion mode in a data-dependent manner. A full MS scan was performed from 380−1600 m/z with resolution at 120,000. The tandem mass spectrometer (MS/MS) CID (Collision-Induced Dissociation was performed with top 15 at 38% of the normalized collision energy (NCE) with a dynamic exclusion time at 30 s, a minimum signal threshold at 1000, a resolution at 30,000, and an isolation width at 1.50 Da.

### Proteins annotation

#### 1- Transcriptomics

Transcripts were assembled using the RNAseq Annotation Pipeline (RAP). The raw paired-end Illumina fastq files were trimmed of any existing primers and the low quality bases (primer-trim & qtrim) and filtered for rRNA sequences. The preprocessed (filtered) sequences were initially used for assembling the contigs at library level. The resulting contigs were then merged by groups. Open reading frames (ORFs) were identified for the merged assembled contigs. Read counts for each ORF were tabulated by mapping the original reads to the assembled contigs of each group. Functional annotation of each ORF was generated using BLAST, HMMER, TMHMM, transporter, organelle & KOG searches of the curated reference sequences in phyloDB database.

#### 2- Protein identification and quantification

PEAKS Studio v.8.0 software (Bioinformatics Solutions Inc., Waterloo, Canada) was used for matching MS/MS spectra against the database (open reading frames from RNAseq) obtained from the transcriptomic analyses described above. Prior to identification, a data refinement was made with default predefined parameters.

PEAKS DB tool was used for the proteins identification against *Ch. simplex* transcriptomics results. 10 ppm and 0.02 Da were set for precursor and fragment ion tolerance respectively; up to one missed trypsin cleavage site allowed. Oxidation of methionine and N-terminal acetylation were set as variable modifications, while carbamidomethylation of cysteine residues was set as fixed modifications. False Discovery Rate (FDR) was set at 0.1% at PSM level.

For Label Free quantification, peptide intensities for each sample were normalized based on the total ion chromatogram (TIC). Only identifications of unique peptides were used for protein quantification. Parameter settings were: Mass Error Tolerance: 10.0 ppm; Retention Time Shift Tolerance: 5.0 min; Dependent on PID: 46; FDR Threshold: 1%. From the identified proteins, only those present in at least 50% of the replicates for each treatment were considered for the subsequent analyses.

### Statistical data analysis

All analyses were performed with R 4.1.3 version (R Core Team 2022).

#### 1- Missing value imputation

Missing values were replaced by the minimum intensity value of that same sample (Liu & Dongre, 2021).

#### 2- Comparison and significant proteins

*S*ignificantly expressed proteins were identified for five comparisons between treatments. In this work we focus in three of those comparisons, providing information about response to heat:

- Nt25 vs. Nt31 (Nt25-Nt31) to identify proteins differentially expressed at over-optimum temperature in acclimated non thermo-tolerant populations.
- Nt25 vs. Tt34 (Nt25-Tt34) to identify proteins differentially expressed at over-optimum temperature in acclimated thermo-tolerant populations.
- Nt31 vs. Tt34 (Nt31-Tt34) to identify proteins differentially expressed at over-optimum temperatures that are different for acclimated non thermo-tolerant and thermo-tolerant populations.

A Student’s t-test was performed for each protein at each comparison in order to evaluate whether the observed differences in the average expression can be explained just by chance (p-value < 0.05). Following Diz *et al*. (2011) recommendation, several methods were explored to correct for multiple comparisons:

- Sequential goodness of fit (SGoF) method. Two variants of this method were tested:

▪ Binomial SGoF method (Carvajal-Rodriguez *et al,* 2009), with the function Binomial.SGoF from the package sgof (Castro-Conde & de Uña-Álvarez 2020) version 2.3.3.
▪ Conservative SGoF mehod (de Uña-Álvarez 2011): recommended for large number of comparisons, with the function SGoF from the package sgof (Castro-Conde & de Uña-Álvarez 2020) version 2.3.3.
- Benjamini & Hochberg method (Benjamini & Hochberg 1995): with the function BH from the package sgof (Castro-Conde & de Uña-Álvarez 2022) version 2.3.3.
- Sequential Fisher method (Diz *et al,* 2011): data were first sorted by the p-value from lowest to highest, then Fisher correction was applied using fisher function from the package poolr (Cinar & Viechtbauer 2022) version 1.1.1. The result was assigned as the corrected p-value for the one protein on the top of the list. Fisher was applied again for all but the first protein. The new result was assigned to the second protein, and so on.
- Sequential Bonferroni method (Holm 1979): with the function *holm* from the package stats (R Core Team 2022).

For each protein at each comparison the q-value was calculated, using the function q-value from the package q-value (Storey *et al.,* 2021) from the Bioconductor version 3.14. This value indicates how many of the significant proteins are actually false positives.

#### 3- Functional enrichment analysis

Functional annotation for all the identified proteins and a gene ontology (GO) enrichment analysis based on Fisher exact test (FDR = 0.05) were performed using Omicsbox software (version 1.4.12). To identify the enriched terms for each comparison, the set of differentially expressed proteins was used as test set whereas all quantified proteins in the experiment were considered the reference set. The reduced option was used for the analysis, showing more specific GO terms. Each enriched GO term was manually classified in a functional category depending on its role in cell metabolism. The categories used are: carbon (C) metabolism, cell organization, DNA-RNA, energy use and storage, fatty acid metabolism, nitrogen (N) metabolism, photosynthesis, protein metabolism (including protein biosynthesis, protein catabolism and protein regulation), respiration, stress response and transport. Those GO terms with general functions that cannot be uniquely assigned to a specific category are classified in the other category.

#### 4- Upregulated GO terms for each treatment and three comparisons

For each enriched GO term at each comparison, the number of proteins upregulated in each treatment was checked. A GO term is considered upregulated for a particular treatment in a comparison, when at least 75% of the significant proteins that correspond to such GO term are upregulated in that treatment.

## RESULTS

Mass spectrometry data analysis identified and quantified 7089 peptides corresponding to 1544 annotated proteins. After removing those proteins that were not detected in at least 50% of the replicates for each treatment, there were 1509 proteins for further analysis.

For each comparison, the number of proteins showing a statistically significant difference (p<0.05 test t-student) between treatments was: 611 for Nt25-Nt31, 440 for Nt25-Tt34, and 247 for Nt31-Tt34. From all multiple comparison corrections tested, Binomial SGoF was the one that maximizes the number of significant proteins with an acceptable q-value (q<0.2, at least 80% are true positives) (supplementary table 1) resulting in the following number of proteins showing statistically significant differences between treatments: 521 for Nt25-Nt31; 350 for Nt25-Tt34; and 157 for Nt31-Tt34. Results after applying this correction method were used for subsequent analyses.

The functional enrichment analysis identifies those biological processes, molecular functions, or cellular components that are enriched (over-represented) compared to the reference set in each comparison. The number of enriched GO terms identified is 113 for Nt25-Nt31, 70 for Nt25-Tt34, and 22 for Nt31-Tt34 (supplementary table 2), all overrepresented compared to the reference set. Those GO terms with % sequences higher than 2 for the test set are shown in figures 2, 3, and 4:

**Figure 2:**
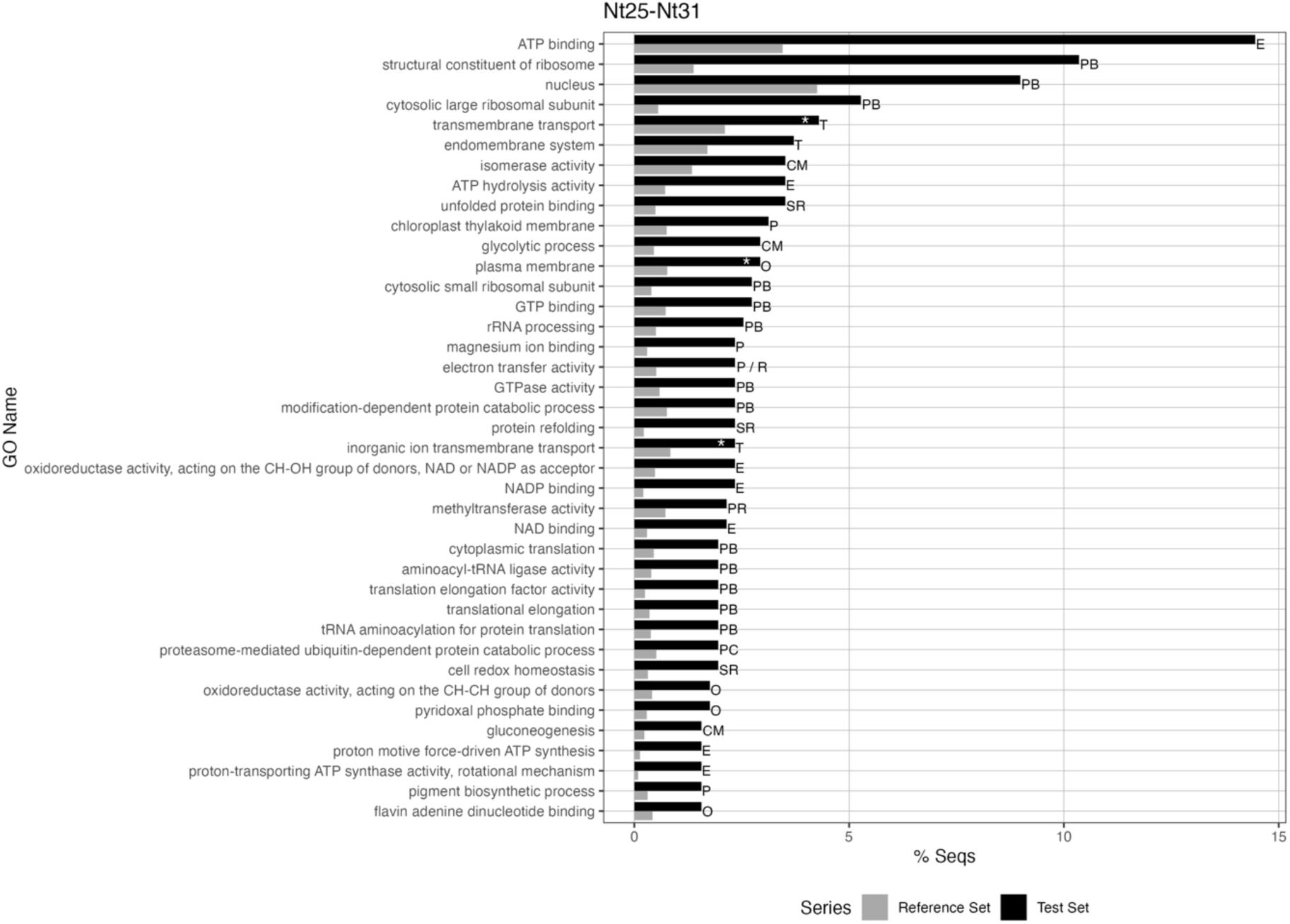
Bar chart of enriched GO terms for the Nt25-Nt31 comparison. *% Seqs* is the percentage of sequences for each GO term, and *GO Name* is the name of the corresponding GO term. This chart only shows those terms with *% Seq* >2. Grey bars represent the reference set (all quantified proteins in the experiment), and black bars are the test set (differentially expressed proteins for this comparison). This is the reduced version of the chart with only the most specific terms. White asterisks indicate terms that are not clearly upregulated for one treatment (none with more than 75% upregulated proteins). Letters on the right are abbreviations for functional categories: CM (carbon metabolism), E (energy related cellular processes), P (photosynthesis), PB (protein biosynthesis), PC (protein catabolism), R (respiration), SR (stress response), T (transport), O (others). Several abbreviations in the same term indicate that it can be related to different categories.

### - Nt25-Nt31

The most enriched GO terms (sequences >5% for the test set) are the molecular functions ATP binding and structural constituent of ribosome; the cellular components nucleus and cytosolic large ribosomal subunit; and the biological process transmembrane transport (figure 2). There are 4 enriched GO terms (with % sequences < 2) that are upregulated for the Nt25 treatment, all of them from the N metabolism functional category (supplementary table 2). Furthermore, there are 3 GO terms not clearly upregulated for Nt31 nor Nt25 (less than 75% of significant proteins for any of them, see methods); all three are related to the membrane and the ion transport through it. From those three terms, proteins related to nitrate transport are upregulated in the Nt25 treatment.

The most represented functional category in the pool of enriched GO terms is protein biosynthesis, which, together with protein regulation, and catabolism, represent 35% of all enriched GO terms (Protein metabolism category in figure 5), all upregulated in Nt31. Details of the functional categories represented by enriched GO terms are described below, in order of term abundance (figure 5):

- Protein metabolism: Most enriched terms in this category are related to transcription and translation. Six enriched GO terms in this category are related to ribosome constitution (cytosolic large ribosomal subunit, cytosolic small ribosomal subunit, ribosomal large subunit assembly, ribosomal small subunit assembly, structural constituent of ribosome, and large ribosomal subunit rRNA binding). Another six enriched GO terms are related to protein catabolism (proteasomal ubiquitin-independent protein catabolic process, proteasome core complex, proteasome-mediated ubiquitin-dependent protein catabolic process, amino acid catabolic process, threonine biosynthetic process, and threonine-type endopeptidase activity).
- C metabolism: glycolysis, gluconeogenesis, and the pentose phosphate pathway are enriched and significantly upregulated in Nt31 populations compared to control (Nt25).
- Stress response: we observed in Nt31 the upregulation and enrichment of GO terms related to ROS scavenging, chaperone proteins activity, and DNA repair (supplementary table 2).
- Energy: Several terms related to the generation, transport, storage, and use of ATP and NAD/NADP are enriched and upregulated in Nt31 (supplementary table 2).
- Photosynthesis: the upregulated and enriched GO terms (supplementary table 2) are related to both photosynthetic pigments and C fixation processes (Calvin cycle and electron transport chain).
- Respiration: the enriched GO terms in this category are related to the tricarboxylic acid cycle.
- Fatty acid metabolism: the enriched GO terms in this category are related to fatty acid biosynthesis.
- Transport: The GO terms from this category enriched and upregulated in Nt31 indicate the increase of cell investment in protein transport and the endomembrane system, which includes the endoplasmic reticulum, Golgi bodies, vesicles, cell membrane, and nuclear envelope.
- DNA-RNA: upregulation and enrichment of the biosynthesis of pyrimidine-containing compounds and purine ribonucleoside monophosphate.
- N metabolism: On the one side, two GO terms are enriched and upregulated in Nt31, the glutamate-ammonia ligase activity (catalyzing the reaction of glutamate with ammonia to form glutamine) and the agmatinase activity (catalyzing the generation of urea and putrescine from agmatine). On the other side, four enriched GO terms related to N metabolism were upregulated in Nt25: nitrate transmembrane transport, nitrate transmembrane transporter activity, nitrite transmembrane transporter activity, and nitrite transport.

### - Nt25-Tt34

The most enriched GO terms (sequences >5% for the test set) are the molecular functions ATP binding and chlorophyll binding; the cellular components cytosol and chloroplast thylakoid membrane; and the biological process cellular component organization, all them upregulated in Tt34 (figure 3). There are 8 enriched GO terms (with % sequences < 2) that are upregulated for the Nt25 treatment, all of them from the N metabolism functional category (supplementary table 2).

**Figure 3:**
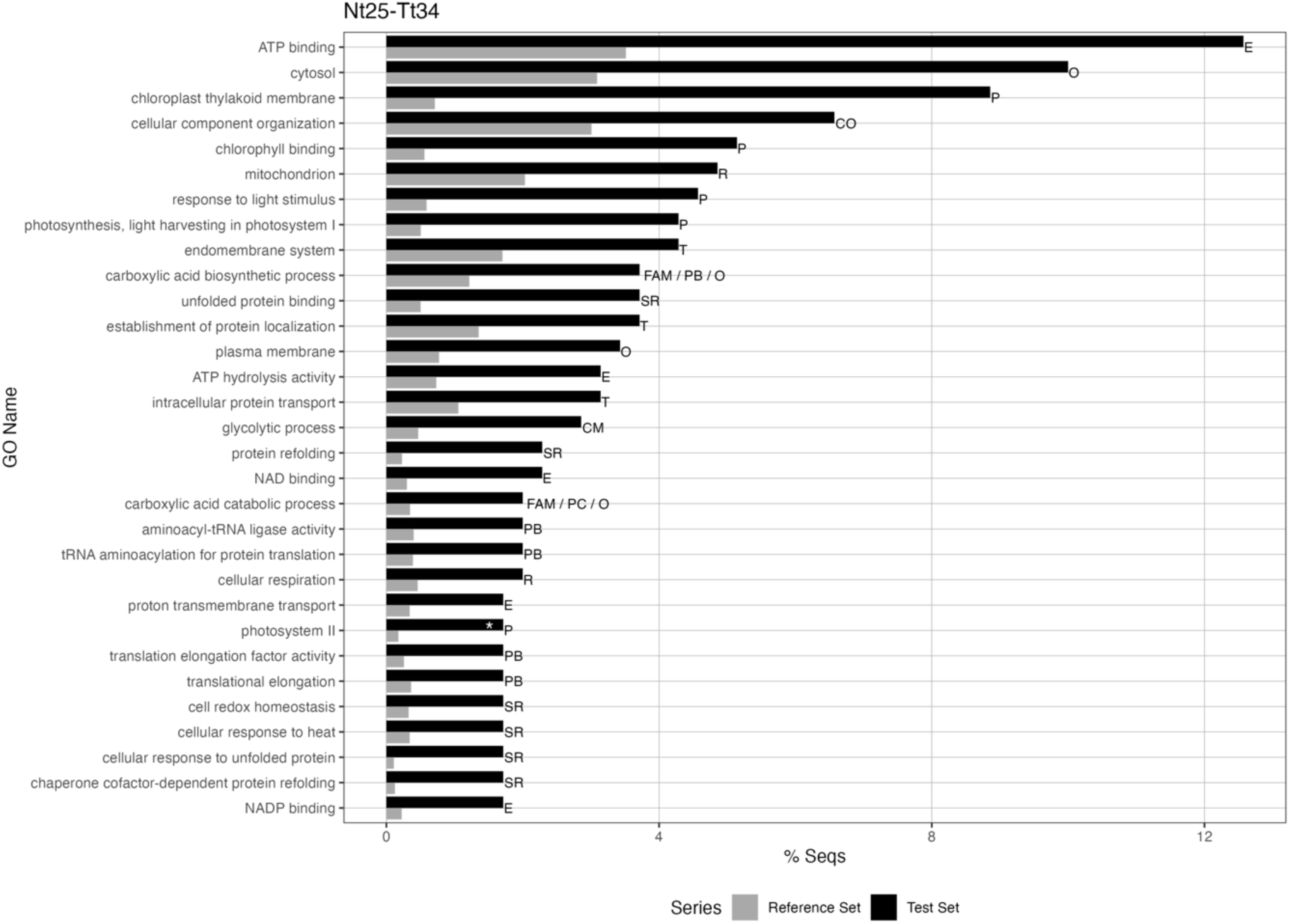
Bar chart of enriched GO terms for the Nt25-Tt34 comparison. *% Seqs* is the percentage of sequences for each GO term, and *GO Name* is the name of the corresponding GO term. This chart only shows those terms with *% Seq* >2. Grey bars represent the reference set (all quantified proteins in the experiment), and black bars are the test set (differentially expressed proteins for this comparison). This is the reduced version of the chart with only the most specific terms. White asterisks indicate terms that are not clearly upregulated for one treatment (none with more than 75% upregulated proteins). Letters on the right are abbreviations for functional categories: CM (carbon metabolism), CO (cell organization), E (cell energy related processes), FAM (fatty acid metabolism), P (photosynthesis), PB (protein biosynthesis), PC (protein catabolism), R (respiration), SR (stress response), T (transport), O (others). Several abbreviations in the same term indicate that it can be related to different categories.

The most represented functional categories are stress response and energy use and storage (figure 5). Details of the functional categories represented by enriched GO terms are described below, in order of terms abundance (figure 5):

- Stress response: The GO terms enriched and upregulated from this category in the Tt34 populations are mostly related to the chaperone proteins activity or heat shock proteins.
- Energy: GO terms related to cellular energy use and storage are enriched and upregulated in Tt34.
- Protein metabolism: GO terms enriched and upregulated in Tt34 are related to translation (aminoacyl-tRNA ligase activity, translation elongation factor activity, translational elongation, tRNA aminoacylation for protein translation) and protein catabolism (aminopeptidase activity, metalloexopeptidase activity).
- Photosynthesis: The enriched and upregulated GO terms from this category in Tt34 populations are related to light harvesting and C fixation.
- Transport: The enriched GO terms are related to transmembrane transport and the activity of the endoplasmic reticulum.
- C metabolism: The four enriched GO terms upregulated for Tt34 in this category (gluconeogenesis, glycolytic process, transketolase activity, and glucose 6-phosphate metabolic process) are related to carbohydrates metabolism.
- Fatty acid metabolism: Two enriched GO terms upregulated in Tt34 are specific for the fatty acid metabolism category: acyl-CoA dehydrogenase activity and fatty acid beta-oxidation using acyl-CoA dehydrogenase
- Respiration: Two GO terms were enriched in Tt34 related to this category: cellular respiration and mitochondrion.
- Cell organization: two terms from this category were enriched in Tt34: cellular component organization and protein insertion into the membrane.
- DNA-RNA: the GO term purine-nucleoside phosphorylase activity is the only one from this category enriched and upregulated for Tt34 compared to Nt25.
- N metabolism: On the one side, there is only one GO term enriched and upregulated for Tt34: agmatinase activity. On the other side, eight GO terms in this category are enriched and upregulated in Nt25 (supplementary table 2).

### - Nt31-Tt34

The most enriched GO terms (sequences >5% for the test set) are the molecular functions structural constituent of ribosome, oxidoreductase activity, RNA binding, and ATP binding; and the cellular components chloroplast, cytosolic large ribosomal subunit, and thylakoid. From those terms, all but thylakoid are upregulated in Nt31 populations (figure 4). No GO term is clearly upregulated for Tt34 populations, and the only 2 GO terms not clearly upregulated for Nt31 are related to the chloroplast (figure 4).

**Figure 4:**
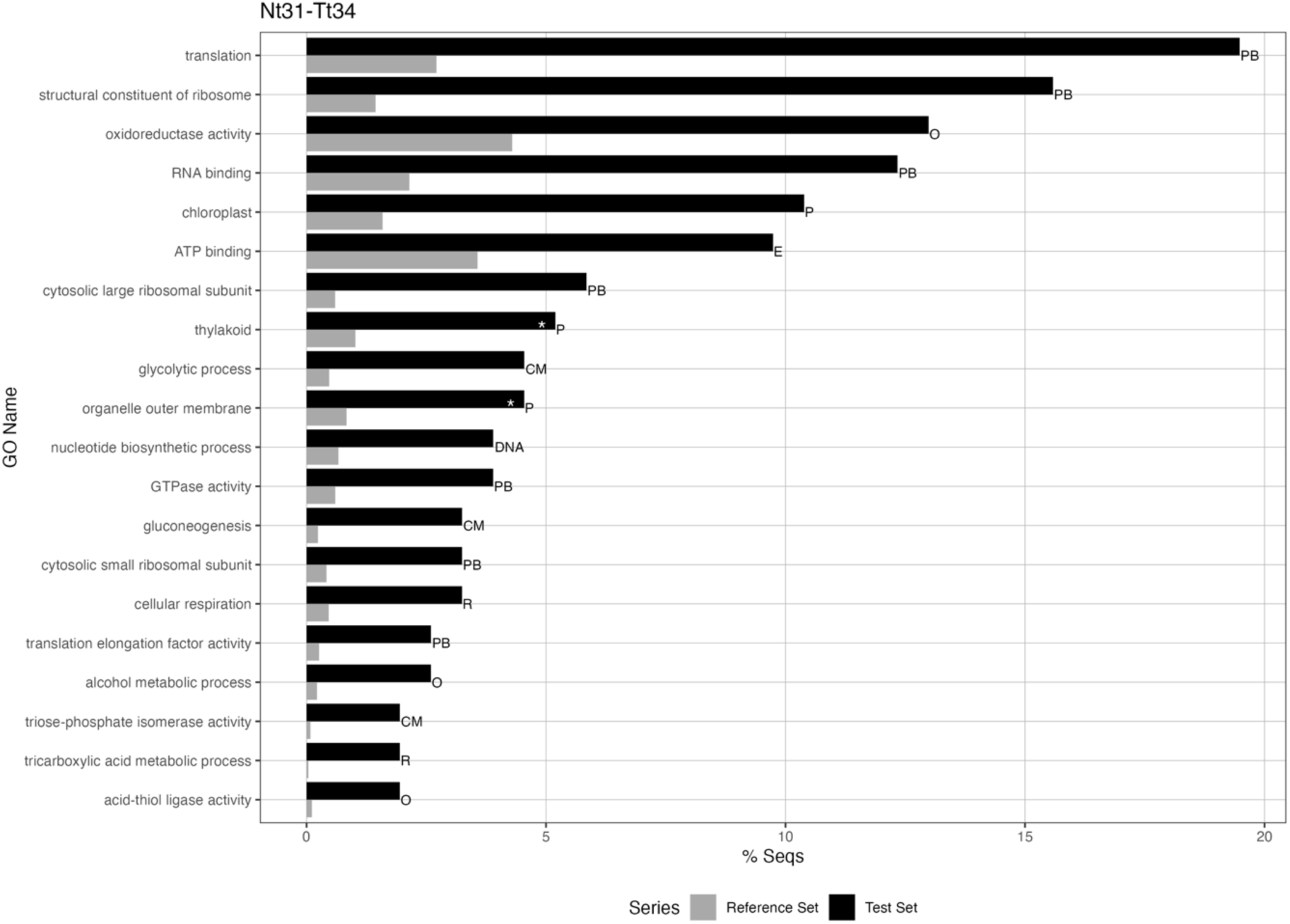
Enriched GO terms bar chart for the comparison Nt31-Tt34. *% Seqs* is the percentage of sequences for each GO term, and *GO Name* is the name of the corresponding GO term. This chart only shows those terms with *% Seq* >2. Grey bars represent the reference set (all quantified proteins in the experiment), and black bars are the test set (differentially expressed proteins for this comparison). This is the reduced version of the chart with only the most specific terms. White asterisks indicate terms that are not clearly upregulated for one treatment (none with more than 75% upregulated proteins). Letters on the right are abbreviations for functional categories: CM (carbon metabolism), DNA (DNA and RNA related processes), E (cell energy related processes), P (photosynthesis), PB (protein biosynthesis), R (respiration), O (others). Several abbreviations in the same term indicate that it can be related to different categories.

The most represented functional category is protein biosynthesis. Details of the functional categories represented by enriched GO terms (all of them enriched in Nt31 populations), are described below in order of terms abundance (figure 5):

- Protein metabolism: the GO terms in this functional category are related to protein biosynthesis and ribosomes, upregulated in Nt31.
- C metabolism and respiration: The GO terms enriched in these categories are related to C fixation, glycolysis, gluconeogenesis, and tricarboxylic acid metabolism (cellular respiration).
- Energy: one enriched GO term in this category (ATP binding) was upregulated in Nt31.
- Photosynthesis: There were three enriched GO terms related to this category, and only one upregulated in Nt31 is chloroplast.
- DNA-RNA: One GO term enriched and upregulated in Nt31: nucleotide biosynthetic process.

**Figure 5:**
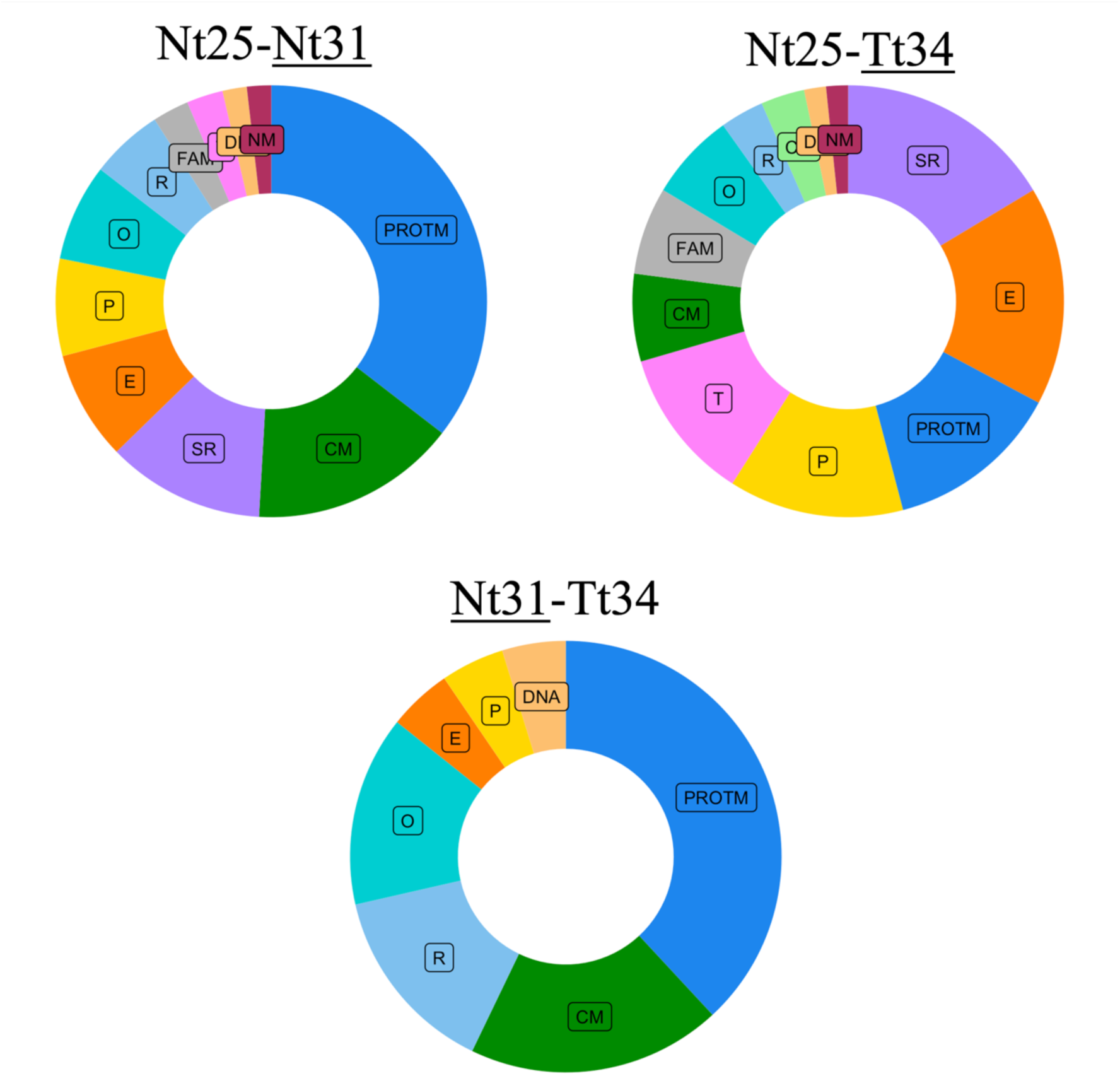
Proportion (%) of GO terms representing each functional category for the three population thermal response comparisons, considering 113, 70, and 22 (for the Nt25-Nt31, Nt25-Tt34, and Nt31-Tt34 comparisons, respectively) total GO terms enriched and upregulated in the underlined population. Labels are abbreviations for the functional categories as follow: PROTM (protein metabolism, including proteins biosynthesis, catabolism, and organization), CM (carbon metabolism), SR (stress response), E (energy), P (photosynthesis), R (respiration), CO (cell organization), DNA (DNA-RNA), FAM (fatty acid metabolism), NM (nitrogen metabolism), T (transport), and O (others). Those terms assigned to more than one category (supplementary table 2) are counted in all of them.

## DISCUSSION

Research has shown that the duration of exposure to stressful environments affects how organisms respond (Benner *et al,* 2013; Chakravarti *et al,* 2020; Collins *et al,* 2020; Padfield *et al,* 2015). In this discussion, we will reference only the literature that specifically describes the exposure time and where the time scale is comparable to our study.

### - Acclimated non-tolerant populations’ response to thermal stress (Nt25-Nt31)

The treatment Nt25 is used in this work as an absolute control because it reveals the response of non-thermo-tolerant populations at their optimal temperature for growth. The enriched GO terms in the comparison Nt25-Nt31 reveal the acclimation response to thermal stress of non-tolerant populations.

In this study, we observed in Nt31 the upregulation of chaperones, ROS protection mechanisms, and repair mechanisms, a trend that has been widely reported as a cellular response to temperature increase (Chakravarti et al, 2020; Liang et al, 2019; Tomanek et al, 2011; Williams et al, 2011; Xu et al, 2014; Zhang et al, 2022), confirming that the cells are suffering from heat stress. The enrichment and upregulation of C metabolism processes as glycolysis, gluconeogenesis and the pentose phosphate pathway, combined with the enrichment of terms related to energy use and storage, suggests that acclimation process in Nt31 is highly energy-demanding. This observation agrees with previous work describing the upregulation of energy generation to cope with thermal stress (Bonora *et al,* 2012; Tomanek *et al,* 2011; Xu *et al,* 2014). The cell needs to increase the ATP generation to manage the increase in investment in metabolic processes. This results also agree with the upregulation of the tricarboxylic acid cycle, a pathway that releases stored energy in the cells and suggest the increase of respiration, as it has been previously described during thermal acclimation (Barton *et al,* 2020; Toseland, 2013). Furthermore, the response of the C fixation and photosynthetic apparatus to heat stress is not universal in marine algae. Some studies have reported a decrease in photosynthesis and/or chlorophyll as a response to heat (Fan *et al,* 2018; Liang *et al,* 2019; Rhee & Gotham, 1981; Xu *et al,* 2014), while others report an increase (Boyd, 2016; Raven & Geider, 1988; Thrane, 2017) or a more complex response depending on the variable studied and temperature tested (Baker *et al,* 2016; Zhang *et al,* 2022); Results from the present work suggest the increase of cellular investment in both photosynthetic pigments and C fixation in non-tolerant populations when acclimated to above-optimal temperatures.

Additionally, the enriched terms in the protein metabolism category suggest that cellular investment in protein biosynthesis increased during the acclimation to high temperature in Nt populations. This trend was previously described for marine algae (Xu *et al,* 2014). Moreover, the enrichment of terms related to ribosome constitution and protein catabolism suggests the concurrent increase of the ribosome concentration as a response to heat (Fan *et al,* 2018) and the lysis of damaged proteins to protect the cell from its toxicity (Chakravarti *et al,* 2020). Likewise, the terms upregulated in the DNA-RNA category suggest the increase of cell investment in nucleic acid biosynthesis, which could be related to the increase in ribosomes.

In relation to fatty acids metabolism, the increase or the composition shift of cellular fatty acids as a response to thermal stress has been previously reported for algae (Chakravarti *et al,* 2020; Xing *et al,* 2018), including diatoms (Feijão *et al,* 2020; O’Donnell *et al.,* 2019, Rousch *et al,* 2004). However, the observed trends are contradictory, with some papers reporting an increase in the total amount or synthesis of fatty acids (Rousch *et al,* 2004; Xing *et al,* 2018), and others reporting an increase in fatty acid abundance (Feijão *et al,* 2020) or catabolism (Chakravarti *et al,* 2020) under thermal stress. Whatever change is observed, it is usually hypothesized to be related to the cellular membrane composition. In our study, the upregulation and enrichment of fatty acid biosynthesis indicate increased cell investment in lipids in the Nt31 population. While we cannot determine the function of those new fatty acids with the available data, it might be linked to the observed enrichment of terms related to transport membranes that may be related to cell growth and duplication.

More surprising is the response we observed in the N metabolism. The enriched terms in such category upregulated in Nt31, suggest an increase in cellular investment in ammonia metabolism (probably coming from protein catabolism or photorespiration) and the reallocation of N for the synthesis of proteins (Allen *et al,* 2011; Guerra *et al,* 2013; Parker *et al,* 2008). However, those terms upregulated in Nt25 indicate the upregulation of cell investment in nitrate uptake and metabolism in Nt25 or, from another perspective, the downregulation of nitrate assimilation in Nt31. It has been described before that nitrate uptake decreases with temperature increase in diatoms (Lomas and Glibert, 1999 and references therein; Parker & Armbrust, 2005). Those studies hypothesize that diatoms upregulate nitrate metabolism as a diffusive pathway for the excess of energy generated at lower temperatures when light and dark photosynthesis reactions are decoupled. So at higher temperatures, less diffusion is needed, and N metabolism is not upregulated. However, when above-optimal temperatures are considered in marine diatoms, nitrate reductase activity (NR, the enzyme catalyzing the first reaction in nitrate assimilation) decreases at higher temperatures (Gleich *et al,* 2020; Kristiansen, 1983), according to the lower optimum temperature for NR activity (Rhee & Gotham, 1981). However, at the same time, the genetic expression for NR is higher (Gleich *et al,* 2020), probably compensating for the lack of NR efficiency. Conversely, we found that at the above-optimum temperatures (both for NR activity and population growth), NR abundance does not significantly change for the non-tolerant populations (NR is not in the list of significant proteins, supplementary table 1). So, contrary to the findings of Gleich *et al,* (2020), the expression level of NR does not compensate for the NR efficacy reduction due to temperature. In addition, nitrate or nitrite transporters are downregulated. In summary, cell investment in nitrate uptake and assimilation decreased in Nt31 populations, and cells compensated for the decline of N from this source, increasing the reallocation of N from ammonia, probably to synthesize new components. The activation of ammonia recycling is a general response to a decrease in N availability in diatoms (Hockin *et al,* 2012; Remmers *et al,* 2018; Scarsini *et al,* 2022).

Summing up, in Nt31, we observed a classical short-term (acclimation) thermal stress response with increased energy demand to sustain the upregulation of protein synthesis, C fixation, and cellular protection mechanisms. But we also found the downregulation of nitrate assimilation under stressful temperature conditions, probably due to lesser efficacy of the NR enzyme at above-optimal temperature, and the upregulation of ammonia metabolism, probably as a strategy to provide N to the highly demanding biosynthesis of proteins and photosynthetic pigments.

### - Acclimated thermo-tolerant populations’ response to thermal stress (Nt25-Tt34) and its comparison with nontolerant populations

The enriched GO terms when comparing the Nt25 and Tt34 responses reveal the acclimated thermal response of the thermo-tolerant populations, resulting from both acclimation and adaptation of the Tt populations (Aranguren-Gassis *et al,* 2019; Collins *et al,* 2020).

As described before for the Nt31 populations, enrichment in energy and respiration categories suggest the increase of cellular energy demand when Tt34 populations are under thermal stress, and similarly to our observations in Nt31 response, thermal acclimation involves increased protein biosynthesis in Tt34 populations. However, the simultaneous enrichment of protein catabolism suggests the recycling of damaged proteins to obtain amino acids. The acclimated response of Tt34 to the above-optimum temperature did not show the enrichment of ribosome-related GO terms described for Nt31. Eight ribosomal proteins were significantly upregulated for Tt34 compared to Nt25 (supplementary table 1); however, there were 50 ribosomal proteins upregulated in Nt31. From those, only six were also significant for Tt34 (the mean p-value for the others is 0.85± 0.05 standard error). So, contrary to what we observed in the non-tolerant population, the increase of ribosomes is not a primary cellular response to heat stress for thermo-tolerant populations.

Enriched terms related to stress response suggest that Tt34 populations activates chaperone proteins to ensure the correct protein’s function during the heat stress, a commonly observed cellular response to heat stress (Shi *et al,* 2017; Tomanek *et al,* 2011; Williams *et al,* 2011). On the contrary, several terms related to ROS protection that were enriched and upregulated for Nt31 were not enriched in Tt34 compared to Nt25 (supplementary table 2). However, some proteins related to those GO terms are significantly upregulated in Tt34 (supplementary table 1). So, there was an increase in ROS protection for Tt34 during acclimation to stressful high temperatures, but such an effect is more pronounced in Nt31 than in Tt34.

We observed similar trends for other functions, such as C metabolism or photosynthesis, suggesting an increase in cellular investment in these processes, though less than in Nt31 populations. For example, there were only four C metabolism enriched GO terms upregulated for Tt34, compared to the 17 in Nt31 for this same category and the enriched and upregulated GO terms from the photosynthesis category in Tt34 populations suggest the increase of cellular investment in light harvesting and C fixation, but not as much for photosynthetic pigments synthesis as in Nt31.

About fatty acids metabolism, the described enriched terms imply an increase of cellular investment in fatty acid catabolism and desaturation, contributing to cellular respiration (Ghisla & Thorp, 2004). Previous observations in high temperature-adapted diatoms showed that thermo-resistant species had higher fatty acid content, and the synthesis of fatty acids increased more when exposed to thermal stress than in not thermo-tolerant species (Rousch *et al,* 2004). Our results did not suggest this same trend; on the contrary, it suggests the increase of fatty acid biosynthesis in the non-tolerant populations under thermal stress but an increase of catabolism in the thermo-tolerant populations.

There were also responses in the cellular transport and cell division GO terms that we didn’t detect for the Nt31. Transport GO terms are related to transmembrane transport and the activity of the endoplasmic reticulum, suggesting the increase of intracellular reallocation of compounds. And the two terms enriched in the cell organization category describe the constitution of cellular structures like membranes, microtubules, or cytoplasm. The upregulation and enrichment of these terms can be related to different processes, like membrane organization for cell division or the cytoskeleton reorganization process previously reported for other organisms during thermal stress (Chakravarti *et al,* 2020; Xu *et al,* 2014). In addition, the purine-nucleoside phosphorylase activity enrichment suggests increased cellular investment in nucleic acid biosynthesis, possibly related to cell division and population growth.

Concerning N metabolism, on the one side, there is only one GO term enriched and upregulated for Tt34: agmatinase activity. This term suggests, similarly to Nt31, the increase of cellular investment in ammonia metabolism. On the other side, eight GO terms in this category are enriched and upregulated in Nt25 (supplementary table 2), suggesting a strong effect of thermal stress on the downregulation of cellular investment in nitrate metabolism in the thermo-tolerant populations. In this case, thermal stress affects not only the transporters but also the nitrate reductase (significantly more abundant in Tt34, supplementary table 1), so nitrate assimilation seems more inhibited for thermal-adapted populations.

In summary, in the same way as non-tolerant populations, the acclimation of the thermo-tolerant populations to thermal stress implies the upregulation of core metabolic pathways such as proteins metabolism, photosynthesis, and respiration, the increase of cellular protection mechanisms, and the increase of the energy metabolism to supply the increased cells demand. However, several differences suggest divergent cellular strategies to cope with thermal stress. Non-tolerant populations invest more in ribosomes, photosynthetic pigments, carbon fixation, and fatty acid biosynthesis, prioritizing growth and cell division. Thermo-tolerant populations, however, invest less in photosynthetic pigments and C fixation, probably reducing ROS generation, which allows reducing expense of ROS protection. At the same time, instead of increasing ribosomes, the increase of proteins and fatty acid catabolism and transport indicate higher recycling activity and C and N reallocation in Tt34 populations. Surprisingly, both populations downregulated nitrate metabolism, but the inhibition seems stronger in thermo-tolerant ones. However, lower investment in pigments, ribosomes, and protein biosynthesis (al rich in N) in the thermo-tolerant populations would reduce the cellular N demand allowing the cells to cope with lower N assimilation.

### - Thermo-tolerant populations’ adaptive response to thermal stress (Nt31-Tt34)

The enriched GO terms in the comparison between Nt31 and Tt34 reveal the processes that differentiate the acclimated response of the non-tolerant and thermo-tolerant populations to temperature stressful conditions, that is, the result of the evolutionary adaptation to thermal stress. The disparities could be the consequence of different temperatures affecting enzyme dynamics. If that were the case, we would expect a similar response from both populations at 34°C. However, Nt doesn’t survive at such temperature, but Tt does, so the direct effect of temperature increase is insufficient to explain our observations. Another possible explanation for the differences found is that 34°C is more stressful for Tt populations than 31°C for Nt populations. However, both temperatures are at a similar thermal distance (6°C for Nt and 5°C for Tt) from the corresponding optimum temperatures (25°C for Nt and 29.2°C for Tt, Aranguren-Gassis *et al*, 2019). So, in any case, stress levels may be higher for Nt31 than Tt34, but growth rates are comparable, being slightly higher for Tt31 (Nt31=1.1 ± 0.01; Tt34=0.9 ± 0.01 growth rate ± standard error; from fig 1.c in Aranguren-Gassis *et al,* 2019). Therefore, we consider that the differences observed between Nt31 and Tt34 come from the inherent ability of Tt populations to tolerate high temperatures.

The GO terms enriched in the C metabolism, respiration and energy categories are upregulated in Nt31, suggesting higher upregulation of cellular investment in those pathways in Nt, probably to cope with the higher demand of energy and C skeletons from the active metabolism in Nt31. The energy cost for the response to thermal stress in the Nt populations may be higher than for Tt populations. Also, as was expected from the results described above, the GO terms related to protein biosynthesis and ribosomes, were upregulated in Nt31. This difference confirms that the response to high temperatures in Nt31 implies higher cellular investment in protein synthesis and ribosomes than in Tt34. The upregulation of photosynthesis related terms agrees with the increase in photosynthetic pigment biosynthesis described in Nt31, and there is one GO term enriched and upregulated in Nt31 for the DNA-RNA category, suggesting higher investment in cell division by non-tolerant populations.

This comparison confirms the main differences between the two cellular strategies described above: (1) non-thermo-tolerant populations invest more in growing, upregulating the proteins and photosynthetic structures biosynthesis, and increasing C fixation and respiration to satisfy higher energy demand. Upregulation of chaperones and ROS protection is needed to maintain the enhanced cell’s metabolism under thermal stress. (2) The thermo-tolerant populations invest more in recycling and protein repair or protection than new structures. They increase C fixation, but not so much to deal with excessive ROS or to need so much energy. It suggests a more conservative strategy that prioritizes survival over rapid growth.

The remaining question is, why this strategy allows Tt populations to survive at higher temperatures? It has been demonstrated that warming increases the minimum N requirement (N*), especially at above-optimal temperatures (Lewington-Pearce *et al.,* 2019; Thomas *et al.,* 2017), but at the same time, according to our results and previous observations (Lomas and Glibert, 1999 and references therein; Parker & Armbrust, 2005) it reduces N uptake. In such a context, the N-conservation strategy in the thermo-tolerant populations may allow them to maintain cellular functioning at higher temperatures, promoting N recycling and reducing highly N-consuming processes.

### - Adaptation to high temperatures under low nitrate availability

One of the most exciting results of our evolution experiment (Aranguren-Gassis *et al,* 2019) was that thermo-tolerant populations did not persist when populations were evolving under low nitrate availability. We hypothesized that N requirements for thermo-tolerant populations were higher due to investment in protection. Consequently, under low nitrate availability, growth rate would be more limited for thermo-tolerant than non-tolerant populations, so the latter would dominate the community after some time. Both the model proposed and the lower C:N ratio in the thermo-tolerant populations supported this explanation.

However, our proteomic analysis provides an alternative explanation. Contrary to what we proposed previously, proteomics suggests that non-tolerant populations invest more in protection, and this strategy requires more energy than in thermo-tolerant populations. Regardless of the different cellular strategies uncovered with proteomics, at 31°C, both types of populations had comparable growth rates (1.1±0.1 for both types of populations, from fig 1.c in Aranguren-Gassis *et al,* 2019). In addition, if temperature limited nitrate assimilation, as proteomics suggests, we can suppose that N uptake was similar for both population types at 31°C, regardless of N availability. In that case, the non-tolerant and thermo-tolerant populations may coexist during the evolution experiment under high N availability conditions. At 31°C, closer to its optimum temperature for growth (Aranguren-Gassis *et al.,* 2019), and with the advantage of better using the available N, the thermo-tolerant genotype gradually became more frequent.

Still, this doesn’t explain why thermo-tolerant populations did not persist under low N availability (Aranguren-Gassis *et al.,* 2019). According to our proteomic analysis, the strategy of the non-tolerant populations was to invest in protection and growth. In contrast, thermo-tolerant populations activated more recycling pathways and downregulated metabolism, reducing oxidative stress. The resulting reallocation of components agrees with the lower C:N ratios we measured in the thermo-tolerant compared to non-tolerant populations (Aranguren-Gassis *et al,* 2019) because, even though N uptake should be similar (NR equally active), the non-tolerant populations invested more in C fixation, potentially increasing the C:N ratio.

The results indicate that thermo-tolerant populations may be better competitors at low N availability conditions. However, we need to also consider the cellular response to N deprivation, which would interact with the thermal response at 31°C and low N availability.

Some general acclimation responses of diatoms to low N availability at the cellular level can be derived from the literature: downregulation of photosynthesis (Hockin *et al,* 2012; Scarsini *et al,* 2022), through both carbon fixation (Bender *et al,* 2014; Guerra *et al,* 2013; Remmers *et al,* 2018) and photosynthetic pigments (Guerra *et al,* 2013; Remmers *et al,* 2018; Scarsini *et al,* 2022), an increase of N recycling from intracellular compounds (Hockin *et al,* 2012; Remmers *et al,* 2018; Scarsini *et al,* 2022; with some differences on the pathways depending on the species or studies considered), downregulation of ribosomes (Bender *et al,* 2014; Guerra *et al,* 2013; Hockin *et al,* 2012; Scarsini *et al,* 2022) and protein synthesis (Hockin *et al,* 2012) and the upregulation of protein catabolism (Bender *et al,* 2014; Hockin *et al,* 2012; Scarsini *et al,* 2022). This acclimation response to low N resembles the acclimated thermal response of the thermo-tolerant populations and is the opposite to the thermal response of the non-tolerant populations in our study. Hence, N limitation may reinforce the cellular strategy observed for the thermo-tolerant populations but may moderate the thermal response for the non-tolerant ones. As a result, in non-tolerant populations, the available N would be invested in supporting as much photosynthesis and protein biosynthesis as possible, i.e., growth, while in thermo-tolerant populations, growth would be more limited, even if they can better tolerate N deprivation (because of the lesser cellular N demand). In the long term, two things can favor the prevalence of the non-tolerant populations: on one hand, these cellular strategies can result in a slightly higher growth rate of the non-tolerant populations, which would allow them to out-compete the thermo-tolerant ones; on the other hand, if growth is not promoted and cell division is more limited in thermo-tolerant populations, the potential to adapt to the new environment (stressful temperature and N limitation) is lower than for the non-tolerant ones, so we can expect the thermo-tolerant population to go extinct in the long term. However, all the information available on the diatoms’ response to low N availability comes from short-term observations, that is acclimation. We do not know of any studies analyzing the diatoms’ adaptative strategies to deal with N limitation, and unfortunately, we did not examine the populations from our evolution experiment. Further experiments are needed to understand diatoms’ adaptive cellular response to simultaneous thermal stress and N limitation.

### - Ecological implications

The patterns observed are consistent with physiological measurements (growth rates and C:N ratios, Aranguren-Gassis *et al,* 2019) and involve core metabolic pathways that probably drive the primary cellular response.

From our observations, temperature increase due to climate change may allow the coexistence of populations with different metabolic strategies, especially in natural environments where nutrient availability fluctuates. This is unsurprising, as high intraspecific diversity has been described for marine diatom species (Rynearson & Armbrust, 2000 and 2005). However, according to our results, only some strains will cope well with future marine heat waves, events of extremely high temperature that are expected to increase in frequency, duration, and maximum temperatures reached (Frolicher *et al,* 2018). Those extreme thermal conditions will favor the most thermo-tolerant strains, whose cellular strategies may be similar to the one described for the thermo-tolerant populations in this study. The increase in the abundance of cells with a more conservative strategy (like the one in our thermo-tolerant populations) may restrict the biomass produced by the diatom blooms. Even under high N availability, thermo-tolerant populations inhibit protein biosynthesis and C fixation in response to high temperature. In addition, the C:N ratio of such populations would be lower; thus, less biomass with different nutritional value would be available for upper trophic levels (John & Davidson, 2001).

We realize that generalizations from the current results may not be possible. Limitations come from several aspects: First, we have studied one species from warm waters, and the thermal response may be species-specific (Aranguren-Gassis & Litchman, 2020). Second, as in many proteomics analyses of non-model organisms, only a portion of existing proteins were identified, and among them some have unknown function, resulting in a significant loss of information, and a complete map of the metabolic pathways does not exist. However, we can still derive conclusions that could motivate future research. Even though a generalization from our laboratory experiments may not be straightforward, our findings reveal successful cellular strategies that warrant further consideration and analysis. This will facilitate better predictions of phytoplankton responses to warming and other stressors in the near future.

In conclusion, evolutionary adaptation may ensure the persistence of marine diatoms species, but the resulting cellular strategies can alter the ecological function of those species. Moreover, the interaction of diverse environmental selective factors would influence the adaptation outcome and may limit the potential to adapt to extreme temperatures.

## AKNOWLEDGMENTS

We thank P. Wilburn for helping with genomic sampling and initial collaborator contacts; J. L. Herrera-Cortijo for helping with R coding; P. Álvarez from the CACTI’s proteomic lab for her help and patience, and E. Marañón for providing laboratory equipment. Funding was provided by a Xunta de Galicia postdoctoral fellowship in 2013 and 2016 and the University of Vigo research talent retention program 2019 to M.A.G.; National Science Foundation grant (OCE-1638958) to E.L. Xunta de Galicia (“Grupos de Referencia Competitiva” ED431C 2020/05) and Fondos Feder (ERDF, European Commission) funded A.P.D. This is Kellogg Biological Station contribution number 2353

## DATA ACCESSIBILITY AND BENEFIT-SHARING

Data accessibility: Original data, analyses scripts and results, will be available upon peer-reviewed publication in Github. Transcriptomic data had been deposited in the NCBI portal. The mass spectrometry proteomics data have been deposited to the ProteomeXchange Consortium via the PRIDE partner repository. Data bases will be publicly available upon peer-reviewed publication.

Benefit-sharing: Benefits from this research accrue from the sharing of our data and results on public databases as described above.

Competing interests: The authors declare that they have no competing interests.

## AUTHORS CONTRIBUTION

MAG, EL, and MHO conceived the idea and designed the experiments. MAG performed the experiments. AA performed the transcriptomics analysis. MAG, APD, and MHO analyzed the proteomics. MAG wrote the manuscript and all authors revised the manuscript and gave final approval.

## BOX 1.

**Terminology used throughout the text**

### Non-tolerant populations (Nt)

one population comes from cultures maintained at 25

°C and nitrate replete conditions, and the other was cryopreserved at the beginning of the evolution experiment, thawed just before the assay and maintained in the N-replete medium at 25 °C (called control and ancestral populations respectively in Aranguren-Gassis *et al,* 2019). None of them were able to grow at 34°C.

### Thermo-tolerant populations (Tt)

cultures that evolved experimentally over ∼200 generations at 31 °C and nitrate replete conditions and could grow at 34°C (called tolerant strains in Aranguren-Gassis *et al,* 2019).

### Nt25

non-tolerant (control) populations grown at 25°C and nitrate replete conditions for the proteomics analysis. We consider this the absolute control (optimal temperature for growth).

### Nt31

non-tolerant populations grown at 31°C (above-optimum temperature) and nitrate replete conditions for the proteomics analysis.

### Tt34

thermo-tolerant populations grown at 34°C (above-optimum temperature) and nitrate replete conditions for the proteomics analysis.

## REFERENCES

1. Allen, A. E., Dupont, C. L., Oborník, M., Horák, A., Nunes-Nesi, A., McCrow, J. P., Zheng, H., Johnson, D. A., Hu, H., Fernie, A. R. & Bowler, C. (2011). Evolution and metabolic significance of the urea cycle in photosynthetic diatoms. Nature, 473, 203–208.

2. Aranguren-Gassis, M., Kremer, C. T., Klausmeier, C. A. & Litchman, E. (2019). Nitrogen limitation inhibits marine diatom adaptation to high temperatures. Ecology Letters, 22, 1860–1869.

3. Aranguren-Gassis, M. & Litchman, E. (2020). Thermal performance of marine diatoms under contrasting nitrate availability. Journal of Plankton Research, 42 (6), 680–688.

4. Baker, K. G., Robinson, C. M., Radford, D. T., McInnes, A. S., Evenhuis, C., & Doblin, M. A. (2016). Thermal performance curves of functional traits aid understanding of thermally induced changes in diatom-mediated biogeochemical fluxes. Frontier in Marine Sciences, 3, 44.

5. Barton, A. D., Irwin, A. J., Finkel, Z. V. & Stock, C. A. (2016). Anthropogenic climate change drives shift and shuffle in North Atlantic phytoplankton communities. Proceedings of the National Academy of Sciences USA, 113, 2964–2969.

6. Barton, S., Jenkins, J., Buckling, A., Schaum, E., Smirnoff, N., Raven, J. A., & Yvon-Durocher, G. (2020). Evolutionary temperature compensation of carbon fixation in marine phytoplankton. Ecology Letters, 23, 722–733.

7. Bender, S. J., Durkin, C. A., Berthiaume, C. T., Morales, R. L. & Armbrust, E. V. (2014). Transcriptional responses of three model diatoms to nitrate limitation of growth. Frontiers in Marine Scicinces, 1, 1–15.

8. Benjamini, Y. & Hochberg, Y. (1995). Controlling the false discovery rate: A practical and power-full approach to multiple testing. Journal of the Royal Statistical Society Series B (Methodological), 57, 289–300.

9. Benner, I. Diner, R. E., Lefebvre, S. C., Li1, D., Komada, T., Carpenter, E. J., & Stillman, J. H. (2013). *Emiliania huxleyi* increases calcification but not expression of calcification-related genes in long-term exposure to elevated temperature and pCO2. Philosophical Transactions of the Royal Society B: Biological Sciences, 368, 20130049–20130049.

10. Bonora, M., Patergnani, S., Rimessi, A., De Marchi, E., Suski, J. M., Bononi, A., Giorgi, C., Marchi, S., Missiroli, S., Poletti, F., Wieckowski, M. R., & Pinton, P. (2012). ATP synthesis and storage. Purinergic Signalling, 8, 343–357.

11. Boyd, P. W., Dillingham, P. W., McGraw, C. M., Armstrong, E. A., Cornwall, C. E., Feng, Y., Hurd, C. L., Gault-Ringold, M., Roleda, M. Y., Timmins-Schi, E. & Nunn, B. L. (2016). Physiological responses of a Southern Ocean diatom to complex future ocean conditions. Nature Climate change, 6, 207–213.

12. Carvajal Rodríguez, A., de Uña Álvarez, J., & Rolán-Álvarez, E. (2009). A new multitest correction (SGoF) that increases its statistical power when increasing the number of tests. BMC Bioinformatics, 10, 209.

13. Chakravarti, L. J., Buerger, P., Levin, R. A. & van Oppen, M. J. H. (2020).Gene regulation underpinning increased thermal tolerance in a laboratory-evolved coral photosymbiont. Molecular Ecology, 29, 1684–1703.

14. Chan, W. Y., Oakeshott, J. G., Buerger, P., Edwards, O. R. & van Oppen, M. J. H. (2021). Adaptive responses of free-living and symbiotic microalgae to simulated future ocean conditions. Global Change Biology, 27:1737–1754.

15. Cinar, O., & Viechtbauer, W. (2022). The poolr package for combining independent and dependent p values. Journal of Statistical Software, 101(1), 1–42.

16. Collins, S., Boyd, P. W. & Doblin, M. A. (2020). Evolution, microbes, and changing Ocean conditions. Annual Review of Marine Sciences, 12, 13.1–13.28.

17. de Uña Álvarez, J. (2011). On the statistical properties of SGoF multitesting method. Statistical Applications in Genetics and Molecular Biology, 10(1), 18.

18. Diz, A. P., Carvajal-Rodriguez, A. & Skibinski, D. O. F. (2011). Multiple hypothesis testing in proteomics: A strategy for experimental work. Molecular Cellular Proteomics, 10, M110.004374.

19. Falkowski, P. G., Fenchel, T. & Delong, E. F. (2008). The microbial engines that drive earth’s biogeochemical cycles. Science 320, 1034–1039.

20. Fan, M., Sun, X., Liao, Z., Wang, J., Li, Y. & Xu, N. (2018). Comparative proteomic analysis of Ulva prolifera response to high temperature stress. Proteome Science, 16:17, 1–22.

21. Feijão, E., Franzitta, M., Cabrita, M. T., Caçador, I., Duarte, B., Gameiro, C. & Matos, A. R. (2020). Marine heat waves alter gene expression of key enzymes of membrane and storage lipids metabolism in *Phaeodactylum tricornutum*. Plant Physiology and Biochemistry, 156, 357–368.

22. Frölicher, T. L., Fischer, E. M. & Gruber, N. (2018). Marine heatwaves under global warming. Nature, 560, 360–376.

23. Ghisla, S. & Thorpe, C. (2004). Acyl-CoA dehydrogenases, a mechanistic overview. European Journal of Biochemistry, 271, 494–508.

24. Gleich, S. J., Plough, L. V. & Glibert, P. M. (2020). Photosynthetic efficiency and nutrient physiology of the diatom *Thalassiosira pseudonana* at three growth temperatures. Marine Biology, 167:124, 1–13.

25. Guerra, L. T., Levitan, O., Frada, M. J., Sun, J. S., Falkowski, P-G. & Dismukes, G. C. (2013). Regulatory branch points affecting protein and lipid biosynthesis in the diatom *Phaeodactylum tricornutum*. Biomass and Bioenergy, 59, 306–315.

26. Guillard, R. R. L. & Hargraves, P. E. (1993). *Stichochrysis immobilis* is a diatom, not a chrysophyte. Phycologia, 32, 234–236.

27. Hockin, N. L., Mock, T., Mulholland, F., Kopriva, S. & Malin, G. (2012). The response of diatom central carbon metabolism to nitrogen starvation is different from that of green algae and higher plants. Plant physiology, 158, 299–312.

28. Holm, S. A. (1979). Simple sequentially rejective multiple test procedure. Scandinavian Journal of Statistics, 6, 65–70.

29. Castro-Conde, I. & de Uña-Alvarez, J. (2020). sgof: Multiple hypothesis testing. R package version 2.3.2. https://CRAN.R-project.org/package=sgof

30. IPCC, (2021). Summary for Policymakers. In: Climate Change 2021: The Physical Science Basis. Contribution of Working Group I to the Sixth Assessment Report of the Intergovernmental Panel on Climate Change [Masson-Delmotte, V., P. Zhai, A. Pirani, S.L. Connors, C. Péan, S. Berger, N. Caud, Y. Chen, L. Goldfarb, M.I. Gomis, M. Huang, K. Leitzell, E. Lonnoy, J.B.R. Matthews, T.K. Maycock, T. Waterfield, O. Yelekçi, R. Yu, and B. Zhou (eds.)].

31. John, E. H. & Davidson, K. (2001). Prey selectivity and the influence of prey carbon: nitrogen ratio on microflagellate grazing. Journal of Experimental Marine Biology and Ecology, 260, 93–111.

32. Storey, J. D., Bass, A. J., Dabney, A. & Robinson, D. (2021). qvalue: Q-value estimation for false discovery rate control. R package version 2.26.0. http://github.com/jdstorey/qvalue

33. Koester, J. A., Swanson, W. J. & Armbrust, E. V. (2013). Positive selection within a diatom species acts on putative protein interactions and transcriptional regulation. Molecular Biology and Evolution, 30, 422–434.

34. Kristiansen, S. (1983). The temperature optimum of the nitrate reductase assay for marine phytoplankton. Limnology & Oceanography, 28, 776–780.

35. Lenoir, J., Bertrand, R., Comte, L., Bourgeaud, L., Hattab, T., Murienne, J., & Grenouillet, G. (2020). Species better track climate warming in the oceans than on land. Nature Ecology & Evolution, 4, 1044–1059.

36. Lewington-Pearce, L. Narwani, A., Thomas, M. K., Kremer, C. T., Vogler, H., & Kratina, P. (2019). Temperature-dependence of minimum resource requirements alters competitive hierarchies in phytoplankton. Oikos, 12, 369–12.

37. Liang, Y., Koester, J. A., Liefer, J. D., Irwin, A. J. & Finkel, Z. V. (2019). Molecular mechanisms of temperature acclimation and adaptation in marine diatoms. The ISME Journal, 13, 2415–2425.

38. Liu, M. & Dongre, A. (2021). Proper imputation of missing values in proteomics datasets for differential expression analysis. Briefings in bioinformatics, 22, 1–17.

39. Lomas, M. W. & Glibert, P. M. (1999).Temperature regulation of nitrate uptake: A novel hypothesis about nitrate uptake and reduction in cool-water diatoms. Limnology & Oceanography, 44, 556–572.

40. O’Donnell, D., Du, Z. & Litchman, E. (2019). Experimental evolution of phytoplankton fatty acid thermal reaction norms. Evolutionary Applications, 12, 1201– 1211.

41. O’Donnell, D., Beery, S. M. & Litchman, E. (2021). Temperature-dependent evolution of cell morphology and carbon and nutrient content in a marine diatom. Limnology & Oceanography, 66, 4334–4346.

42. Padfield, D., Yvon-Durocher, G., Buckling, A., Jennings, S. & Yvon-Durocher, G. (2015). Rapid evolution of metabolic traits explains thermal adaptation in phytoplankton. Ecology Letters, 19, 133–142.

43. Parker, M. S. & Armbrust, E. V. (2005). Synergestic effects of light, temperature, and nitrogen source on transcription of genes for carbon and nitrogen metabolism in the centric diatom *Thalassiosira pseudonana* (Bacyllariophyceae). Journal of Phycology, 41, 1142–1153.

44. Parker, M. S., Mock, T. & Armbrust, E. V. (2022). Genomic insights into marine microalgae. Annual Review of Genetics, 42, 619–645.

45. Pinsky, M. L., Eikeset, A. M., McCauley, D. J., Payne, J. L. & Sunday, J. M. (2019). Greater vulnerability to warming of marine versus terrestrial ectotherms. Nature, 569, 108–111.

46. Pinsky, M. L., Selden, R. L. & Kitchel, Z. J. (2020). Climate-driven shifts in marine species ranges: Scaling from organisms to communities. Annual Reviews of Marine Science, 12, 153–179.

47. Raven, J. A. & Geider, R. J. (1988). Temperature and algal growth. New Phytologist, 110, 441–461.

48. Remmers, I. M., D’Adamo, S., Martens, D. E., de Vos, R. C. H., Mumm, R., America, A. H. P., Cordewener, J. H. G., Bakker, L. V., Peters, S. A., Wijffels, R. H., & Lamers, P. P. (2018). Orchestration of transcriptome, proteome and metabolome in the diatom *Phaeodactylum tricornutum* during nitrogen limitation. Algal Research, 35, 33– 49.

49. Rhee, G. Y. & Gotham, I. J. (1981). The effect of environmental factors on phytoplankton growth: Temperature and the interactions of temperature with nutrient limitation. Limnology & Oceanography, 26, 635–648.

50. Rousch, J. M., Bingham, S. E. & Sommerfeld, M. R. (2004). Protein expression during heat stress in thermo-intolerant and thermo-tolerant diatoms. Journal of Experimental Marine Biology and Ecology, 306, 231–243.

51. Rynearson, T. A. & Armbrust, E. V. (2000). DNA fingerprinting reveals extensive genetic diversity in a field population of the centric diatom. Limnology & Oceanography, 45, 1329–1340.

52. Rynearson, T. A. & Armbrust, E. V. (2005). Maintenance of clonal diversity during a spring bloom of the centric diatom *Ditylum brightwellii*. Molecular Ecology, 14, 1631–1640.

53. Scarsini, M. Thiriet-Rupert, S., Veidl, B., Mondeguer, F., Hu, H., Marchand, J. & Schoefs, B. (2022). The transition toward nitrogen deprivation in diatoms requires chloroplast stand-by and deep metabolic reshuffling. Frontiers in Plant Sciences, 12, 760516.

54. Schaum, C. E., Buckling, A., Smirnoff, N., Studholme, D. J. & Yvon-Durocher, G. (2018)a. Environmental fluctuations accelerate molecular evolution of thermal tolerance in a marine diatom. Nature Communications, 9, 1719.

55. Schaum, C. E., Student Research Team, ffrench-Constant, R., Lowe, C., Olafsson, J. S., Padfield, D. & Yvon-Durocher, G. (2018)b.Temperature-driven selection on metabolic traits increases the strength of an algal-grazer interaction in naturally warmed streams. Global Change Biology, 24, 1793–1803.

56. Shi, J., Chen, Y., Xu, Y., Ji, D., Chen, C. & Xie, C. (2017). Differential proteomic analysis by iTRAQ reveals the mechanism of *Pyropia haitanensis* responding to high temperature stress. Scientific Reports, 7, 44734.

57. Thomas, M. K., Aranguren-Gassis, M., Kremer, C. T., Gould, M. R., Anderson, K., Klausmeier, C. A. & Litchman, E. (2017). Temperature-nutrient interactions exacerbate sensitivity to warming in phytoplankton. Global Change Biology, 23, 3269–3280.

58. Thrane, J. E., Hessen, D. O. & Andersen, T. (2017). Plasticity in algal stoichiometry: Experimental evidence of a temperature-induced shift in optimal supply N:P ratio. Limnology & Oceanography, 122, 1121–1129.

59. Tomanek, L. (2011). Environmental proteomics: Changes in the proteome of marine organisms in response to environmental stress, pollutants, infection, symbiosis, and development. Annual Review of Marine Sciences, 3, 373–399.

60. Toseland, A., Daines, S., Clark, J., Kirkham, A., Strauss, J., Uhlig, C., Lenton, T. M., Valentin, K., Pearson, G., Moulton, V., & Mock, T. (2013). The impact of temperature on marine phytoplankton resource allocation and metabolism. Nature Climate change, 3, 979–984.

61. Williams, T. J., Lauro, F. M., Ertan, H., Burg, D. W., Poljak, A., Raftery, M. J. & Cavicchioli, R. (2011). Defining the response of a microorganism to temperatures that span its complete growth temperature range (−2°C to 28°C) using multiplex quantitative proteomics. Environmental Microbiology, 13, 2186–2203.

62. Xing, G., Yuan, H. L., Yang, J. S., Li, J. Y., Gao, Q. X., Li, W. L. & Wang, E. T. (2018). Integrated analyses of transcriptome, proteome and fatty acid profilings of the oleaginous microalga *Auxenochlorella protothecoides* UTEX 2341 reveal differential reprogramming of fatty acid metabolism in response to low and high temperatures. Algal Research, 33, 16–27.

63. Xu, Y., Chen, C., Ji, D., Hang, N. & Xie, C. (2014). Proteomic profile analysis of *Pyropia haitanensis* in response to high-temperature stress. Journal of Applied Phycology, 26, 607–618.

64. Yvon-Durocher, G., Dossena, M., Trimmer, M., Woodward, G. & Allen, A. P. (2015). Temperature and the biogeography of algal stoichiometry. Global Ecology and Biogeography, 24, 562–570.

65. Yvon-Durocher, G., Schaum, C. E. & Trimmer, M. (2017). The temperature dependence of phytoplankton stoichiometry: Investigating the roles of species sorting and local adaptation. Frontiers in Microbiology, 8, E2182–14.

66. Zhang, N., Mattoon, E. M., McHargue, W., Venn, B., Zimmer, D., Pecani, K., Jeong, J., Anderson, C. M., Chen, C., Berry, J. C., Xia, M., Tzeng, S. C., Becker, E., Pazouki, L., Evans, B., Cross, F., Cheng, J., Czymmek, K. J., Schroda, M., Mühlhaus T. & Zhang, R. (2022). Systems-wide analysis revealed shared and unique responses to moderate and acute high temperatures in the green alga *Chlamydomonas reinhardtii*. Communications Biology, 5, 460.

