## Supplemental tables for "Evolution of thermal tolerance in marine diatoms: Metabolic strategies under heat stress"

**SUPLEMENT**

Supplementary table 1: Compilation of the p values corrected for multiple comparisons with five different methods and the q-value for each protein. The red color indicates $p < 0.05$.The proteinAccession is the identification code for each protein; Description is the descriptive identification of each protein; ttest_pvals are the original p values without correction; Bisnomial_SGoF is the Binomial SGoF method (Carvajal-Rodriguez *et al.,* 2009); Conservative_SGoF is the conservative SGoF method (de Uña-Álvarez 2011); BH is the Benjamini & Hochberg method (Benjamini and Hochberg 1995); SFisher is the sequential Fisher method (Diz *et al.,* 2011); and Bonferroni is the sequential Bonferroni method (Holm 1979); q_val is the q value. The original data can be downloaded from the repository specified in the main text of the paper (files Multiple_comparison_p_C25vsC31_Descr_Smallest.csv; Multiple_comparison_p_C25vsE34_Descr_Smallest.csv, and Multiple_comparison_p_C31vsE34_Descr_Smallest.csv)

Nt25 – Nt31

| proteinAccession | ttest_pvals | Binomial_SGoF | Conservative_SGoF | BH | SFisher | Bonferroni | q_val | Description |
|---|---|---|---|---|---|---|---|---|
| contig_6708_5 | 0 | 0 | 0 | 0 | 0 | 0 | 0 | hemagglutinin/amebocyte aggregation factor |
| contig_6069_2 | 0 | 0 | 0 | 0 | 0 | 0 | 0 | GLNA glutamine synthase |
| contig_16604_4 | 0 | 0 | 0 | 0 | 0 | 1E-04 | 0 | myo-inositol 2-dehydrogenase putative |
| contig_22629_4 | 0 | 0 | 0 | 0 | 0 | 1E-04 | 0 | acetyl-coa carboxylase |
| contig_10415_5 | 0 | 0 | 0 | 1E-04 | 0 | 4E-04 | 0 | DNA-binding protein 42 kDa |
| contig_14326_6 | 0 | 0 | 0 | 2E-04 | 0 | 0.0023 | 1E-04 | peroxiredoxin |
| contig_37313_1 | 0 | 0 | 0 | 2E-04 | 0 | 0.0023 | 1E-04 | predicted protein |
| contig_39738_3 | 0 | 0 | 0 | 2E-04 | 0 | 0.0024 | 1E-04 | Asparaginyl-tRNA synthetase |
| contig_23318_6 | 0 | 0 | 0 | 2E-04 | 0 | 0.0026 | 1E-04 | putative LOV domain-containing protein |
| contig_5982_4 | 0 | 0 | 0 | 2E-04 | 0 | 0.003 | 1E-04 | abnormal spindle-like microcephaly-associated protein homolog |
| contig_27950_6 | 0 | 0 | 0 | 2E-04 | 0 | 0.0031 | 1E-04 | aspartyl-tRNA synthetase |
| contig_30765_6 | 0 | 0 | 0 | 2E-04 | 0 | 0.0031 | 1E-04 | dynactin subunit 1 |
| contig_17134_1 | 0 | 0 | 0 | 2E-04 | 0 | 0.0032 | 1E-04 | 60S ribosomal protein L22 |
| contig_19340_6 | 0 | 0 | 0 | 2E-04 | 0 | 0.0035 | 1E-04 | acetyl-coa carboxylase |
| contig_26787_4 | 0 | 0 | 0 | 2E-04 | 0 | 0.0037 | 1E-04 | DUF4336 domain-containing protein |
| contig_35026_4 | 0 | 0 | 0 | 2E-04 | 0 | 0.0038 | 1E-04 | glyceraldehyde 3-phosphate dehydrogenase |

| | | | | | | | | |
|---|---|---|---|---|---|---|---|---|
| contig_14556_3 | 0 | 0 | 0 | 2E-04 | 0 | 0.004 | 1E-04 | anion-transporting ATPase |
| contig_38406_6 | 0 | 0 | 0 | 2E-04 | 0 | 0.0042 | 1E-04 | predicted protein |
| contig_40308_5 | 0 | 0 | 0 | 2E-04 | 0 | 0.0044 | 1E-04 | 40S ribosomal protein S6 |
| contig_13084_2 | 0 | 0 | 0 | 2E-04 | 0 | 0.005 | 1E-04 | 60S ribosomal protein L14 putative |
| contig_34173_4 | 0 | 0 | 0 | 2E-04 | 0 | 0.005 | 1E-04 | succinate dehydrogenase flavoprotein subunit |
| contig_5881_6 | 0 | 0 | 0 | 2E-04 | 0 | 0.0053 | 1E-04 | elongation factor Tu (chloroplast) |
| contig_22636_2 | 0 | 0 | 0 | 3E-04 | 0 | 0.0061 | 1E-04 | 60s acidic ribosomal protein p0 |
| contig_30318_6 | 0 | 0 | 0 | 3E-04 | 0 | 0.0067 | 1E-04 | pyruvate dehydrogenase complex dihydrolipoamide acetyltransferase |
| contig_28919_4 | 0 | 0 | 0 | 3E-04 | 0 | 0.0078 | 1E-04 | adenosylhomocysteinase |
| contig_5654_2 | 0 | 0 | 0 | 3E-04 | 0 | 0.0084 | 1E-04 | hypothetical protein THAOC_17223 partial |
| contig_28181_3 | 0 | 0 | 0 | 4E-04 | 0 | 0.0093 | 1E-04 | dihydroxy-acid dehydratase |
| contig_28501_6 | 0 | 0 | 0 | 4E-04 | 0 | 0.0111 | 2E-04 | predicted protein |
| contig_22011_5 | 0 | 0 | 0 | 4E-04 | 0 | 0.0117 | 2E-04 | argininosuccinate synthase |
| contig_17693_5 | 0 | 0 | 0 | 4E-04 | 0 | 0.0121 | 2E-04 | elongation factor-like |
| contig_33214_4 | 0 | 0 | 0 | 4E-04 | 0 | 0.0122 | 2E-04 | phosphoadenosine-phosphosulphate reductase |
| contig_17394_3 | 0 | 0 | 0 | 4E-04 | 0 | 0.013 | 2E-04 | proliferating cell nuclear antigen |
| contig_12284_5 | 0 | 0 | 0 | 4E-04 | 0 | 0.0136 | 2E-04 | ribosomal protein S9 |
| contig_28236_1 | 0 | 0 | 0 | 5E-04 | 0 | 0.0184 | 2E-04 | tRNA splicing endonuclease putative |
| contig_21931_4 | 0 | 0 | 0 | 5E-04 | 0 | 0.0191 | 2E-04 | chaperonin GroEL (chloroplast) |
| contig_28103_5 | 0 | 0 | 1E-04 | 5E-04 | 0 | 0.0191 | 2E-04 | magnesium chelatase subunit H |
| contig_40859_3 | 0 | 0 | 1E-04 | 6E-04 | 0 | 0.022 | 3E-04 | predicted protein |
| contig_39519_2 | 0 | 0 | 1E-04 | 6E-04 | 0 | 0.0225 | 3E-04 | predicted protein |
| contig_27675_6 | 0 | 0 | 1E-04 | 6E-04 | 0 | 0.0226 | 3E-04 | inosine-5'-monophosphate dehydrogenase |
| contig_28781_6 | 0 | 0 | 1E-04 | 6E-04 | 0 | 0.0237 | 3E-04 | heat shock protein 90 |
| contig_7982_5 | 0 | 0 | 1E-04 | 6E-04 | 0 | 0.0243 | 3E-04 | photosystem II protein V (chloroplast) |
| contig_26987_1 | 0 | 0 | 1E-04 | 6E-04 | 0 | 0.0246 | 3E-04 | predicted protein |
| contig_3037_3 | 0 | 0 | 1E-04 | 6E-04 | 0 | 0.0259 | 3E-04 | ribosomal protein S4 (chloroplast) |

| | | | | | | | | |
|---|---|---|---|---|---|---|---|---|
| contig_40308_6 | 0 | 0 | 1E-04 | 6E-04 | 0 | 0.026 | 3E-04 | 40S ribosomal protein S6 |
| contig_24967_5 | 0 | 0 | 1E-04 | 6E-04 | 0 | 0.0272 | 3E-04 | PGR5-like protein 1A chloroplastic |
| contig_9976_6 | 0 | 0 | 1E-04 | 6E-04 | 0 | 0.0285 | 3E-04 | poly(U)-binding-splicing factor PUF60 |
| contig_29625_1 | 0 | 0 | 1E-04 | 6E-04 | 0 | 0.0287 | 3E-04 | rRNA 2'-O-methyltransferase fibrillarin |
| contig_23664_6 | 0 | 0 | 1E-04 | 6E-04 | 0 | 0.0287 | 3E-04 | enoyl-acp reductase |
| contig_28903_4 | 0 | 0 | 1E-04 | 6E-04 | 0 | 0.0305 | 3E-04 | predicted protein |
| contig_37802_5 | 0 | 0 | 1E-04 | 6E-04 | 0 | 0.0305 | 3E-04 | predicted protein |
| contig_28927_1 | 0 | 0 | 1E-04 | 7E-04 | 0 | 0.0337 | 3E-04 | 60S ribosomal protein |
| contig_8347_5 | 0 | 0 | 1E-04 | 8E-04 | 0 | 0.0393 | 3E-04 | elongation factor Ts |
| contig_33347_4 | 0 | 0 | 1E-04 | 8E-04 | 0 | 0.04 | 3E-04 | V-type ATPase A subunit |
| contig_10223_6 | 0 | 0 | 1E-04 | 8E-04 | 0 | 0.0401 | 3E-04 | predicted protein |
| contig_22651_1 | 0 | 0 | 1E-04 | 8E-04 | 0 | 0.0436 | 3E-04 | glutamate synthase subunit alpha |
| contig_16887_1 | 0 | 0 | 1E-04 | 8E-04 | 0 | 0.0444 | 4E-04 | predicted protein |
| contig_27190_6 | 0 | 0 | 1E-04 | 0.001 | 0 | 0.0525 | 4E-04 | 20S proteasome subunit alpha 2 |
| contig_25695_5 | 0 | 0 | 1E-04 | 0.001 | 0 | 0.0563 | 4E-04 | 40S ribosomal protein S7 |
| contig_22474_6 | 0 | 0 | 1E-04 | 0.001 | 0 | 0.0592 | 4E-04 | succinyl-CoA ligase beta-chain mitochondrial precursor |
| contig_6911_6 | 0 | 0 | 1E-04 | 0.0011 | 0 | 0.0613 | 5E-04 | pyrophosphate-dependent phosphofructose kinase |
| contig_17954_4 | 0 | 0 | 1E-04 | 0.0011 | 0 | 0.0654 | 5E-04 | predicted protein |
| contig_22726_1 | 0 | 0 | 1E-04 | 0.0011 | 0 | 0.0662 | 5E-04 | Valine-tRNA ligase |
| contig_34750_5 | 0 | 0 | 1E-04 | 0.0011 | 0 | 0.0691 | 5E-04 | thioredoxin-disulfide reductase |
| contig_33684_3 | 0 | 1E-04 | 1E-04 | 0.0012 | 0 | 0.0712 | 5E-04 | nadh dehydrogenase |
| contig_32718_6 | 0 | 1E-04 | 1E-04 | 0.0012 | 0 | 0.0719 | 5E-04 | aspartate-semialdehyde dehydrogenase |
| contig_14441_4 | 1E-04 | 1E-04 | 1E-04 | 0.0012 | 0 | 0.0761 | 5E-04 | predicted protein |
| contig_8232_1 | 1E-04 | 1E-04 | 1E-04 | 0.0012 | 0 | 0.0783 | 5E-04 | alanine--glyoxylate aminotransferase family protein |
| contig_15521_3 | 1E-04 | 1E-04 | 1E-04 | 0.0012 | 0 | 0.0795 | 5E-04 | predicted protein |
| contig_16696_4 | 1E-04 | 1E-04 | 1E-04 | 0.0012 | 0 | 0.0807 | 5E-04 | pyruvate kinase |
| contig_10561_5 | 1E-04 | 1E-04 | 1E-04 | 0.0012 | 0 | 0.0816 | 5E-04 | 6-phosphogluconate dehydrogenase |

| | | | | | | | | |
|---|---|---|---|---|---|---|---|---|
| contig_39499_3 | 1E-04 | 1E-04 | 1E-04 | 0.0013 | 0 | 0.0851 | 5E-04 | adenylosuccinate synthetase |
| contig_11114_6 | 1E-04 | 1E-04 | 1E-04 | 0.0013 | 0 | 0.0878 | 5E-04 | predicted protein |
| contig_24360_5 | 1E-04 | 1E-04 | 1E-04 | 0.0013 | 0 | 0.0881 | 5E-04 | splicing factor 3B subunit 3-like |
| contig_36968_1 | 1E-04 | 1E-04 | 1E-04 | 0.0013 | 0 | 0.09 | 5E-04 | 60S ribosomal protein L38 |
| contig_21030_6 | 1E-04 | 1E-04 | 2E-04 | 0.0013 | 0 | 0.0909 | 5E-04 | 60S ribosomal protein L39 |
| contig_14853_4 | 1E-04 | 1E-04 | 2E-04 | 0.0013 | 0 | 0.0916 | 5E-04 | peptidyl-prolyl cis-trans isomerase FKBP19 chloroplastic |
| contig_6050_1 | 1E-04 | 1E-04 | 2E-04 | 0.0014 | 0 | 0.1002 | 6E-04 | heat shock protein 70 (chloroplast) |
| contig_17251_3 | 1E-04 | 1E-04 | 2E-04 | 0.0015 | 0 | 0.1087 | 6E-04 | CBS domain-containing protein CBSCBSPB5-like |
| contig_33630_6 | 1E-04 | 1E-04 | 2E-04 | 0.0015 | 0 | 0.1094 | 6E-04 | actin |
| contig_17853_5 | 1E-04 | 1E-04 | 2E-04 | 0.0015 | 0 | 0.1101 | 6E-04 | predicted protein |
| contig_10303_5 | 1E-04 | 1E-04 | 2E-04 | 0.0015 | 0 | 0.1151 | 6E-04 | 40S ribosomal protein S3-3 |
| contig_25619_1 | 1E-04 | 1E-04 | 2E-04 | 0.0015 | 0 | 0.1158 | 6E-04 | ornithine--oxo-acid transaminase |
| contig_28450_6 | 1E-04 | 1E-04 | 2E-04 | 0.0015 | 0 | 0.1223 | 7E-04 | transketolase |
| contig_30640_5 | 1E-04 | 1E-04 | 2E-04 | 0.0015 | 0 | 0.1226 | 7E-04 | predicted protein |
| contig_37567_5 | 1E-04 | 1E-04 | 2E-04 | 0.0016 | 0 | 0.126 | 7E-04 | ATP synthase gamma subunit |
| contig_9376_2 | 1E-04 | 1E-04 | 2E-04 | 0.0016 | 0 | 0.1287 | 7E-04 | predicted protein |
| contig_18580_6 | 1E-04 | 1E-04 | 2E-04 | 0.0017 | 0 | 0.136 | 7E-04 | predicted protein |
| contig_29981_4 | 1E-04 | 1E-04 | 2E-04 | 0.0017 | 0 | 0.1395 | 7E-04 | 20S proteasome subunit alpha 4 |
| contig_26583_1 | 1E-04 | 1E-04 | 2E-04 | 0.0017 | 0 | 0.1408 | 7E-04 | predicted protein |
| contig_29073_1 | 1E-04 | 1E-04 | 2E-04 | 0.0017 | 0 | 0.1411 | 7E-04 | 26S proteasome non-ATPase regulatory subunit 6 |
| contig_8751_4 | 1E-04 | 1E-04 | 2E-04 | 0.0017 | 0 | 0.1429 | 7E-04 | predicted protein |
| contig_40094_2 | 1E-04 | 1E-04 | 2E-04 | 0.0017 | 0 | 0.1481 | 7E-04 | 40S ribosomal protein S14 |
| contig_16274_4 | 1E-04 | 1E-04 | 2E-04 | 0.0017 | 0 | 0.1501 | 7E-04 | predicted protein |
| contig_40261_6 | 1E-04 | 1E-04 | 2E-04 | 0.0017 | 0 | 0.1508 | 7E-04 | predicted protein |
| contig_22708_3 | 1E-04 | 1E-04 | 2E-04 | 0.0017 | 0 | 0.1513 | 7E-04 | Laminin G domain protein |
| contig_22960_6 | 1E-04 | 1E-04 | 2E-04 | 0.0017 | 0 | 0.152 | 7E-04 | urease alpha subunit |
| contig_27467_4 | 1E-04 | 1E-04 | 2E-04 | 0.0017 | 0 | 0.1562 | 7E-04 | histidinol-phosphate aminotransferase |

| | | | | | | | | |
|---|---|---|---|---|---|---|---|---|
| contig_28789_2 | 1E-04 | 1E-04 | 2E-04 | 0.0017 | 0 | 0.1565 | 7E-04 | predicted protein |
| contig_32573_3 | 1E-04 | 1E-04 | 2E-04 | 0.0017 | 0 | 0.1607 | 7E-04 | ATP synthase CF1 alpha subunit (chloroplast) |
| contig_38550_1 | 1E-04 | 1E-04 | 2E-04 | 0.0018 | 0 | 0.1643 | 7E-04 | polyadenylate-binding protein 1-B |
| contig_30735_2 | 1E-04 | 1E-04 | 3E-04 | 0.0018 | 0 | 0.1688 | 8E-04 | myo-inositol 2-dehydrogenase putative |
| contig_16094_5 | 1E-04 | 1E-04 | 3E-04 | 0.0018 | 0 | 0.1688 | 8E-04 | 40S ribosomal protein S23 |
| contig_9076_4 | 1E-04 | 1E-04 | 3E-04 | 0.0018 | 0 | 0.1726 | 8E-04 | 60S ribosomal protein L24 putative |
| contig_4611_6 | 1E-04 | 1E-04 | 3E-04 | 0.0018 | 0 | 0.1744 | 8E-04 | long chain acyl-coa synthetase |
| contig_32146_2 | 1E-04 | 1E-04 | 3E-04 | 0.0018 | 0 | 0.1801 | 8E-04 | silicon transporter |
| contig_32774_1 | 1E-04 | 1E-04 | 3E-04 | 0.0018 | 0 | 0.1809 | 8E-04 | 3-deoxy-7-phosphoheptulonate synthase |
| contig_24799_5 | 1E-04 | 1E-04 | 3E-04 | 0.0018 | 0 | 0.1811 | 8E-04 | predicted protein |
| contig_26510_5 | 1E-04 | 2E-04 | 3E-04 | 0.0019 | 0 | 0.1875 | 8E-04 | LL-diaminopimelate aminotransferase |
| contig_28347_6 | 1E-04 | 2E-04 | 3E-04 | 0.0019 | 0 | 0.194 | 8E-04 | DEAD-box ATP-dependent RNA helicase 8 |
| contig_23256_4 | 1E-04 | 2E-04 | 3E-04 | 0.002 | 0 | 0.2046 | 9E-04 | fumarate hydratase mitochondrial |
| contig_10672_3 | 2E-04 | 2E-04 | 3E-04 | 0.0021 | 0 | 0.2152 | 9E-04 | ATP synthase subunit |
| contig_6927_4 | 2E-04 | 2E-04 | 3E-04 | 0.0021 | 0 | 0.2153 | 9E-04 | predicted protein |
| contig_7380_4 | 2E-04 | 2E-04 | 3E-04 | 0.0021 | 0 | 0.2165 | 9E-04 | predicted protein |
| contig_21155_4 | 2E-04 | 2E-04 | 4E-04 | 0.0021 | 0 | 0.2172 | 9E-04 | heat shock 70 kDa protein |
| contig_28463_3 | 2E-04 | 2E-04 | 4E-04 | 0.0022 | 0 | 0.2384 | 0.001 | aspartate kinase |
| contig_25286_5 | 2E-04 | 2E-04 | 4E-04 | 0.0023 | 0 | 0.2433 | 0.001 | ap-2 complex subunit mu |
| contig_22686_4 | 2E-04 | 2E-04 | 4E-04 | 0.0023 | 0 | 0.2498 | 0.001 | predicted protein |
| contig_30865_4 | 2E-04 | 2E-04 | 4E-04 | 0.0023 | 0 | 0.2503 | 0.001 | 2-oxo acid dehydrogenase subunit E2 |
| contig_30060_4 | 2E-04 | 2E-04 | 4E-04 | 0.0023 | 0 | 0.2504 | 0.001 | nitrite transporter NAR1 |
| contig_26348_5 | 2E-04 | 2E-04 | 4E-04 | 0.0023 | 0 | 0.2582 | 0.001 | heat shock protein 70 |
| contig_19197_6 | 2E-04 | 2E-04 | 4E-04 | 0.0024 | 0 | 0.2637 | 0.001 | casein kinase I isoform X1 |
| contig_39276_6 | 2E-04 | 2E-04 | 4E-04 | 0.0024 | 0 | 0.2759 | 0.001 | orotidine 5'-phosphate decarboxylase |
| contig_16024_6 | 2E-04 | 2E-04 | 4E-04 | 0.0024 | 0 | 0.2771 | 0.001 | Chaperonin GroL |
| contig_25924_5 | 2E-04 | 2E-04 | 4E-04 | 0.0024 | 0 | 0.2789 | 0.001 | ATP-dependent Clp protease proteolytic subunit |

| | | | | | | | | |
|---|---|---|---|---|---|---|---|---|
| contig_34452_4 | 2E-04 | 2E-04 | 5E-04 | 0.0025 | 0 | 0.2815 | 0.001 | glyceraldehyde-3-phosphate dehydrogenase precursor |
| contig_17447_3 | 2E-04 | 2E-04 | 5E-04 | 0.0025 | 0 | 0.2891 | 0.0011 | tetrapyrrole-binding protein |
| contig_27062_5 | 2E-04 | 2E-04 | 5E-04 | 0.0025 | 0 | 0.29 | 0.0011 | heat shock protein 70 |
| contig_30141_2 | 2E-04 | 2E-04 | 5E-04 | 0.0025 | 0 | 0.2911 | 0.0011 | Lipoamide acyltransferase component of branched-chain alpha-keto acid dehydrogenase complex mitochondrial |
| contig_38361_6 | 2E-04 | 2E-04 | 5E-04 | 0.0025 | 0 | 0.2964 | 0.0011 | 60S ribosomal protein L7a |
| contig_24914_6 | 2E-04 | 2E-04 | 5E-04 | 0.0025 | 0 | 0.298 | 0.0011 | bifunctional dTDP-4-dehydrorhamnose 3 5-epimerase/dTDP-4-dehydrorhamnose reductase-like |
| contig_6709_6 | 2E-04 | 2E-04 | 5E-04 | 0.0025 | 0 | 0.3005 | 0.0011 | predicted protein |
| contig_14379_2 | 2E-04 | 2E-04 | 5E-04 | 0.0025 | 0 | 0.3005 | 0.0011 | probable aspartyl aminopeptidase |
| contig_29723_5 | 2E-04 | 2E-04 | 5E-04 | 0.0026 | 0 | 0.3148 | 0.0011 | predicted protein |
| contig_34140_1 | 2E-04 | 2E-04 | 5E-04 | 0.0026 | 0 | 0.3153 | 0.0011 | predicted protein |
| contig_10857_4 | 2E-04 | 2E-04 | 5E-04 | 0.0026 | 0 | 0.3255 | 0.0011 | predicted protein |
| contig_20080_5 | 2E-04 | 3E-04 | 5E-04 | 0.0027 | 0 | 0.3314 | 0.0011 | AP-1 complex subunit beta-1-like |
| contig_8784_1 | 2E-04 | 3E-04 | 5E-04 | 0.0027 | 0 | 0.3328 | 0.0011 | 40S ribosomal protein S10 |
| contig_41222_3 | 2E-04 | 3E-04 | 5E-04 | 0.0027 | 0 | 0.3359 | 0.0011 | predicted protein |
| contig_26303_6 | 2E-04 | 3E-04 | 5E-04 | 0.0027 | 0 | 0.3369 | 0.0011 | DEAD-domain-containing protein |
| contig_25490_5 | 3E-04 | 3E-04 | 6E-04 | 0.0028 | 0 | 0.3587 | 0.0012 | ribonucleoside-diphosphate reductase large subunit |
| contig_28628_1 | 3E-04 | 3E-04 | 6E-04 | 0.0029 | 0 | 0.3737 | 0.0012 | phosphoribosylaminoimidazolecarboxamide formyltransferase/IMP cyclohydrolase |
| contig_13540_6 | 3E-04 | 3E-04 | 6E-04 | 0.0029 | 0 | 0.3742 | 0.0012 | RNA polymerase beta subunit (chloroplast) |
| contig_27595_5 | 3E-04 | 3E-04 | 6E-04 | 0.003 | 0 | 0.3843 | 0.0013 | transitional endoplasmic reticulum ATPase |
| contig_34979_6 | 3E-04 | 3E-04 | 6E-04 | 0.003 | 0 | 0.3934 | 0.0013 | ubiquitin-like protein |
| contig_30076_1 | 3E-04 | 3E-04 | 6E-04 | 0.0032 | 0 | 0.4193 | 0.0014 | elongation factor 2 |
| contig_17285_6 | 3E-04 | 3E-04 | 7E-04 | 0.0032 | 0 | 0.4218 | 0.0014 | 40S ribosomal protein S27-4 putative |
| contig_26782_3 | 3E-04 | 3E-04 | 7E-04 | 0.0032 | 0 | 0.426 | 0.0014 | predicted protein |
| contig_17829_4 | 3E-04 | 3E-04 | 7E-04 | 0.0032 | 0 | 0.4327 | 0.0014 | predicted protein |
| contig_13715_6 | 3E-04 | 3E-04 | 7E-04 | 0.0034 | 0 | 0.4605 | 0.0015 | dnaJ homolog subfamily B member 6 isoform X2 |

| | | | | | | | | |
|---|---|---|---|---|---|---|---|---|
| contig_39493_ 5 | 3E-04 | 4E-04 | 7E-04 | 0.00 35 | 0 | 0.4742 | 0.00 15 | 60s ribosomal protein |
| contig_24139_ 1 | 3E-04 | 4E-04 | 7E-04 | 0.00 35 | 0 | 0.476 | 0.00 15 | chaperonin GroEL |
| contig_7234_4 | 3E-04 | 4E-04 | 8E-04 | 0.00 35 | 0 | 0.4765 | 0.00 15 | ras-related gtp-binding protein |
| contig_30834_ 3 | 3E-04 | 4E-04 | 8E-04 | 0.00 35 | 0 | 0.4771 | 0.00 15 | phosphoglycerate kinase |
| contig_18571_ 6 | 4E-04 | 4E-04 | 8E-04 | 0.00 35 | 0 | 0.4804 | 0.00 15 | phosphoribulokinase |
| contig_5427_6 | 4E-04 | 4E-04 | 8E-04 | 0.00 37 | 0 | 0.5218 | 0.00 16 | tyrosyl-tRNA synthetase |
| contig_28307_ 4 | 4E-04 | 4E-04 | 8E-04 | 0.00 37 | 0 | 0.5244 | 0.00 16 | atp-dependent rna helicase uap56 |
| contig_24606_ 4 | 4E-04 | 4E-04 | 8E-04 | 0.00 38 | 0 | 0.534 | 0.00 16 | beta-ketoacyl-ACP synthase I |
| contig_35366_ 6 | 4E-04 | 4E-04 | 8E-04 | 0.00 38 | 0 | 0.5366 | 0.00 16 | chaperonin GroEL (chloroplast) |
| contig_14697_ 5 | 4E-04 | 5E-04 | 9E-04 | 0.00 4 | 0 | 0.5718 | 0.00 17 | fructose 1 6-bisphosphatase |
| contig_21768_ 6 | 4E-04 | 5E-04 | 9E-04 | 0.00 4 | 0 | 0.5812 | 0.00 17 | 40S ribosomal protein |
| contig_29386_ 5 | 4E-04 | 5E-04 | 9E-04 | 0.00 4 | 0 | 0.5825 | 0.00 17 | predicted protein |
| contig_17741_ 5 | 4E-04 | 5E-04 | 9E-04 | 0.00 42 | 0 | 0.6015 | 0.00 18 | endonuclease 4 |
| contig_25862_ 1 | 4E-04 | 5E-04 | 9E-04 | 0.00 42 | 0 | 0.6102 | 0.00 18 | predicted protein |
| contig_25894_ 5 | 5E-04 | 5E-04 | 9E-04 | 0.00 42 | 0 | 0.6183 | 0.00 18 | predicted protein |
| contig_16700_ 6 | 5E-04 | 5E-04 | 9E-04 | 0.00 42 | 0 | 0.621 | 0.00 18 | serine decarboxylase |
| contig_30397_ 1 | 5E-04 | 5E-04 | 9E-04 | 0.00 42 | 0 | 0.6231 | 0.00 18 | FAD-dependent monooxygenase |
| contig_21909_ 6 | 5E-04 | 5E-04 | 0.001 | 0.00 43 | 0 | 0.6404 | 0.00 18 | 26S proteasome beta type 7 subunit |
| contig_32502_ 6 | 5E-04 | 5E-04 | 0.001 | 0.00 43 | 0 | 0.6484 | 0.00 18 | predicted protein |
| contig_22539_ 3 | 5E-04 | 5E-04 | 0.001 | 0.00 45 | 0 | 0.6723 | 0.00 19 | sulfate adenylyltransferase |
| contig_41330_ 2 | 5E-04 | 5E-04 | 0.001 | 0.00 45 | 0 | 0.6727 | 0.00 19 | 60S ribosomal protein L23a |
| contig_28453_ 2 | 5E-04 | 5E-04 | 0.001 | 0.00 45 | 0 | 0.6778 | 0.00 19 | methionine-dependent methyltransferase |
| contig_5987_6 | 5E-04 | 5E-04 | 0.0011 | 0.00 45 | 0 | 0.68 | 0.00 19 | predicted protein |
| contig_30723_ 5 | 5E-04 | 5E-04 | 0.0011 | 0.00 45 | 0 | 0.6866 | 0.00 19 | 20S proteasome subunit beta 1 |
| contig_30106_ 6 | 5E-04 | 5E-04 | 0.0011 | 0.00 45 | 0 | 0.6867 | 0.00 19 | predicted protein |
| contig_8777_3 | 5E-04 | 5E-04 | 0.0011 | 0.00 45 | 0 | 0.7045 | 0.00 19 | elongation factor G |
| contig_7984_6 | 5E-04 | 6E-04 | 0.0011 | 0.00 46 | 0 | 0.7092 | 0.00 19 | ribosomal protein L1 (chloroplast) |

| | | | | | | | | |
|---|---|---|---|---|---|---|---|---|
| contig_37452_ 3 | 5E-04 | 6E-04 | 0.0012 | 0.00 46 | 0 | 0.7172 | 0.00 19 | ADP-ribosylation factor |
| contig_25987_ 5 | 5E-04 | 6E-04 | 0.0012 | 0.00 46 | 0 | 0.7242 | 0.00 2 | polyadenylate-binding protein 1-B |
| contig_33607_ 5 | 5E-04 | 6E-04 | 0.0012 | 0.00 46 | 0 | 0.7271 | 0.00 2 | S-adenosyl-L-methionine-dependent methyltransferase |
| contig_27592_ 6 | 5E-04 | 6E-04 | 0.0013 | 0.00 46 | 0 | 0.7347 | 0.00 2 | zeta-carotene desaturase |
| contig_30881_ 4 | 6E-04 | 6E-04 | 0.0013 | 0.00 48 | 0 | 0.7668 | 0.00 2 | pyruvate carboxylase |
| contig_24939_ 3 | 6E-04 | 6E-04 | 0.0013 | 0.00 49 | 0 | 0.7849 | 0.00 21 | beta-ketoacyl-[acyl-carrier-protein] synthase II |
| contig_15800_ 4 | 6E-04 | 7E-04 | 0.0013 | 0.00 5 | 0 | 0.8055 | 0.00 21 | asparagine synthase |
| contig_36382_ 5 | 6E-04 | 7E-04 | 0.0014 | 0.00 51 | 0 | 0.82 | 0.00 22 | Ubiquitin-40S ribosomal protein S27a |
| contig_20441_ 2 | 6E-04 | 7E-04 | 0.0014 | 0.00 52 | 0 | 0.8438 | 0.00 22 | putative glyoxalase I |
| contig_27259_ 5 | 6E-04 | 7E-04 | 0.0014 | 0.00 53 | 0 | 0.8655 | 0.00 22 | ATP synthase F1 gamma subunit |
| contig_6795_4 | 6E-04 | 7E-04 | 0.0014 | 0.00 53 | 0 | 0.8655 | 0.00 22 | WSC domain protein putative |
| contig_10170_ 1 | 7E-04 | 8E-04 | 0.0014 | 0.00 54 | 0 | 0.8958 | 0.00 23 | predicted protein |
| contig_14595_ 3 | 7E-04 | 8E-04 | 0.0014 | 0.00 55 | 0 | 0.9103 | 0.00 23 | predicted protein |
| contig_38598_ 5 | 7E-04 | 8E-04 | 0.0015 | 0.00 55 | 0 | 0.9198 | 0.00 24 | Adenylyl cyclase-associated protein |
| contig_12195_ 1 | 7E-04 | 8E-04 | 0.0016 | 0.00 56 | 0 | 0.9282 | 0.00 24 | thioredoxin-like protein 4A isoform X1 |
| contig_39862_ 1 | 7E-04 | 8E-04 | 0.0016 | 0.00 57 | 0 | 0.9494 | 0.00 24 | pyruvate kinase |
| contig_28396_ 1 | 7E-04 | 8E-04 | 0.0017 | 0.00 58 | 0 | 0.9788 | 0.00 25 | chaperone DnaK |
| contig_789_5 | 8E-04 | 8E-04 | 0.0017 | 0.00 59 | 0 | 0.9999 | 0.00 25 | ribulose-1 5-bisphosphate carboxylase/oxygenase large subunit (chloroplast) |
| contig_15223_ 4 | 8E-04 | 9E-04 | 0.0017 | 0.00 59 | 0 | 1 | 0.00 25 | predicted protein |
| contig_22594_ 4 | 8E-04 | 9E-04 | 0.0017 | 0.00 59 | 0 | 1 | 0.00 25 | magnesium chelatase subunit H |
| contig_13616_ 6 | 8E-04 | 9E-04 | 0.0017 | 0.00 6 | 0 | 1 | 0.00 25 | hsp70-like protein |
| contig_10358_ 5 | 8E-04 | 9E-04 | 0.0017 | 0.00 6 | 0 | 1 | 0.00 26 | 60S ribosomal protein L12 |
| contig_27196_ 4 | 8E-04 | 9E-04 | 0.0017 | 0.00 63 | 0 | 1 | 0.00 27 | predicted protein |
| contig_35648_ 5 | 8E-04 | 9E-04 | 0.0018 | 0.00 64 | 0 | 1 | 0.00 27 | ubiquitin / ribosomal protein CEP52 fusion protein putative |
| contig_8636_5 | 9E-04 | 9E-04 | 0.0018 | 0.00 65 | 0 | 1 | 0.00 28 | isocitrate dehydrogenase |
| contig_27301_ 5 | 9E-04 | 9E-04 | 0.0019 | 0.00 65 | 0 | 1 | 0.00 28 | enolase |

| | | | | | | | | |
|---|---|---|---|---|---|---|---|---|
| contig_23563_5 | 9E-04 | 0.001 | 0.0019 | 0.0065 | 0 | 1 | 0.0028 | FK506-binding protein 2 isoform X2 |
| contig_29943_6 | 9E-04 | 0.001 | 0.0019 | 0.0065 | 0 | 1 | 0.0028 | Got1-domain-containing protein |
| contig_13566_4 | 9E-04 | 0.001 | 0.0019 | 0.0065 | 0 | 1 | 0.0028 | 60S ribosomal protein L35a-4 |
| contig_29531_5 | 9E-04 | 0.001 | 0.0019 | 0.0065 | 0 | 1 | 0.0028 | oxidoreductase putative |
| contig_33265_1 | 9E-04 | 0.001 | 0.002 | 0.0066 | 0 | 1 | 0.0028 | phenylalanine-tRNA ligase beta subunit |
| contig_22589_6 | 9E-04 | 0.0011 | 0.002 | 0.0066 | 0 | 1 | 0.0028 | Peptidase_M16-domain-containing protein |
| contig_17958_6 | 9E-04 | 0.0011 | 0.002 | 0.0066 | 0 | 1 | 0.0028 | luminal-binding protein |
| contig_38780_2 | 0.001 | 0.0011 | 0.002 | 0.0071 | 0 | 1 | 0.003 | carbamoyl-phosphate synthase large subunit |
| contig_25542_6 | 0.001 | 0.0011 | 0.002 | 0.0072 | 0 | 1 | 0.003 | ---NA--- |
| contig_9330_5 | 0.001 | 0.0011 | 0.002 | 0.0073 | 0 | 1 | 0.0031 | short-chain alcohol dehydrogenase with NAD or NADP as acceptor |
| contig_29419_6 | 0.001 | 0.0011 | 0.002 | 0.0073 | 0 | 1 | 0.0031 | leucyl-tRNA synthetase |
| contig_26248_5 | 0.001 | 0.0012 | 0.0021 | 0.0074 | 0 | 1 | 0.0032 | predicted protein |
| contig_27719_4 | 0.0011 | 0.0013 | 0.0021 | 0.0075 | 0 | 1 | 0.0032 | predicted protein |
| contig_25957_2 | 0.0011 | 0.0013 | 0.0021 | 0.0076 | 0 | 1 | 0.0033 | predicted protein |
| contig_22654_2 | 0.0011 | 0.0013 | 0.0021 | 0.0078 | 0 | 1 | 0.0033 | predicted protein |
| contig_15405_3 | 0.0011 | 0.0013 | 0.0022 | 0.0078 | 0 | 1 | 0.0033 | predicted protein |
| contig_38458_3 | 0.0011 | 0.0014 | 0.0022 | 0.0078 | 0 | 1 | 0.0033 | Proteasome subunit alpha/beta |
| contig_29871_5 | 0.0012 | 0.0014 | 0.0022 | 0.0081 | 0 | 1 | 0.0035 | lipoprotein type 6 |
| contig_6818_2 | 0.0012 | 0.0014 | 0.0022 | 0.0082 | 0 | 1 | 0.0035 | predicted protein |
| contig_29735_5 | 0.0012 | 0.0014 | 0.0022 | 0.0086 | 0 | 1 | 0.0036 | dihydrolipoamide succinyltransferase |
| contig_37722_1 | 0.0013 | 0.0014 | 0.0023 | 0.0086 | 0 | 1 | 0.0036 | hypothetical protein |
| contig_17554_4 | 0.0013 | 0.0014 | 0.0023 | 0.0089 | 0 | 1 | 0.0038 | predicted protein |
| contig_10341_1 | 0.0013 | 0.0015 | 0.0023 | 0.0089 | 0 | 1 | 0.0038 | predicted protein |
| contig_12144_4 | 0.0013 | 0.0015 | 0.0023 | 0.009 | 0 | 1 | 0.0038 | FAD-binding domain-containing protein |
| contig_40158_1 | 0.0014 | 0.0016 | 0.0023 | 0.0091 | 0 | 1 | 0.0039 | extrinsic protein in photosystem II |
| contig_27030_2 | 0.0014 | 0.0016 | 0.0023 | 0.0092 | 0 | 1 | 0.0039 | geranylgeranyl reductase |
| contig_9754_5 | 0.0014 | 0.0017 | 0.0024 | 0.0092 | 0 | 1 | 0.0039 | long-chain-fatty-acid-CoA ligase |

| | | | | | | | | |
|---|---|---|---|---|---|---|---|---|
| contig_33444_6 | 0.0014 | 0.0017 | 0.0024 | 0.0093 | 0 | 1 | 0.004 | predicted protein |
| contig_22705_1 | 0.0014 | 0.0017 | 0.0024 | 0.0095 | 0 | 1 | 0.004 | glyceraldehyde-3-phosphate dehydrogenase precursor |
| contig_9050_5 | 0.0014 | 0.0017 | 0.0025 | 0.0095 | 0 | 1 | 0.004 | triose phosphate/phosphate translocator |
| contig_3111_6 | 0.0015 | 0.0017 | 0.0025 | 0.0095 | 0 | 1 | 0.0041 | acyl-CoA dehydrogenase |
| contig_13757_6 | 0.0015 | 0.0017 | 0.0025 | 0.0097 | 0 | 1 | 0.0041 | predicted protein |
| contig_17409_1 | 0.0016 | 0.0017 | 0.0025 | 0.0102 | 0 | 1 | 0.0043 | vesicle-associated membrane protein 7b |
| contig_23545_2 | 0.0016 | 0.0018 | 0.0027 | 0.0103 | 0 | 1 | 0.0044 | predicted protein |
| contig_28730_5 | 0.0017 | 0.0019 | 0.0027 | 0.0105 | 0 | 1 | 0.0045 | predicted protein |
| contig_12511_6 | 0.0017 | 0.0019 | 0.0027 | 0.0105 | 0 | 1 | 0.0045 | predicted protein |
| contig_31795_6 | 0.0017 | 0.0019 | 0.0027 | 0.0105 | 0 | 1 | 0.0045 | glyceraldehyde-3-phosphate dehydrogenase |
| contig_10583_5 | 0.0017 | 0.0019 | 0.0028 | 0.0105 | 0 | 1 | 0.0045 | photosystem II stability/assembly factor HCF136 |
| contig_6012_4 | 0.0017 | 0.0019 | 0.0028 | 0.0107 | 0 | 1 | 0.0045 | V-type proton ATPase subunit B isoform |
| contig_9759_1 | 0.0017 | 0.002 | 0.0029 | 0.0108 | 0 | 1 | 0.0046 | clathrin heavy chain |
| contig_41157_2 | 0.0017 | 0.002 | 0.003 | 0.0109 | 0 | 1 | 0.0046 | predicted protein |
| contig_25846_1 | 0.0018 | 0.002 | 0.003 | 0.011 | 0 | 1 | 0.0047 | glyceraldehyde-3-phosphate dehydrogenase |
| contig_4841_4 | 0.0018 | 0.002 | 0.003 | 0.0113 | 0 | 1 | 0.0048 | 40S ribosomal protein S14 |
| contig_33643_5 | 0.0019 | 0.002 | 0.003 | 0.0115 | 0 | 1 | 0.0049 | predicted protein |
| contig_5813_3 | 0.0019 | 0.002 | 0.0031 | 0.0115 | 0 | 1 | 0.0049 | transketolase |
| contig_29387_4 | 0.0019 | 0.002 | 0.0031 | 0.0118 | 0 | 1 | 0.005 | predicted protein |
| contig_7643_5 | 0.0019 | 0.0021 | 0.0032 | 0.0118 | 0 | 1 | 0.005 | ATP synthase CF1 beta subunit (chloroplast) |
| contig_38543_1 | 0.0019 | 0.0021 | 0.0032 | 0.0118 | 0 | 1 | 0.005 | predicted protein |
| contig_23961_3 | 0.002 | 0.0021 | 0.0033 | 0.0118 | 0 | 1 | 0.005 | ribosomal protein S15 |
| contig_30840_2 | 0.002 | 0.0021 | 0.0033 | 0.0118 | 0 | 1 | 0.005 | 60S ribosomal protein L18a |
| contig_28448_5 | 0.002 | 0.0021 | 0.0034 | 0.0119 | 0 | 1 | 0.005 | glycine dehydrogenase |
| contig_20200_5 | 0.002 | 0.0022 | 0.0034 | 0.0119 | 0 | 1 | 0.005 | 60s ribosomal protein l18a |
| contig_30073_5 | 0.002 | 0.0022 | 0.0035 | 0.0119 | 0 | 1 | 0.005 | transaldolase |
| contig_28753_4 | 0.002 | 0.0022 | 0.0035 | 0.0119 | 0 | 1 | 0.0051 | FKBP-type peptidyl-prolyl cis-trans isomerase |

| | | | | | | | | |
|---|---|---|---|---|---|---|---|---|
| contig_9372_6 | 0.002 | 0.0022 | 0.0035 | 0.01 19 | 0 | 1 | 0.00 51 | predicted protein |
| contig_9486_1 | 0.0021 | 0.0023 | 0.0037 | 0.01 21 | 0 | 1 | 0.00 51 | thiamine biosynthesis protein G (chloroplast) |
| contig_24180_ 5 | 0.0021 | 0.0023 | 0.0037 | 0.01 21 | 0 | 1 | 0.00 51 | predicted protein |
| contig_23061_ 6 | 0.0021 | 0.0023 | 0.0038 | 0.01 22 | 0 | 1 | 0.00 52 | predicted protein |
| contig_36190_ 5 | 0.0021 | 0.0023 | 0.0038 | 0.01 22 | 0 | 1 | 0.00 52 | long chain acyl-coa synthetase |
| contig_12115_ 3 | 0.0021 | 0.0023 | 0.0039 | 0.01 25 | 0 | 1 | 0.00 53 | predicted protein |
| contig_28093_ 5 | 0.0022 | 0.0023 | 0.0039 | 0.01 26 | 0 | 1 | 0.00 54 | protein dispatched homolog 1-like |
| contig_25343_ 2 | 0.0022 | 0.0024 | 0.0039 | 0.01 26 | 0 | 1 | 0.00 54 | predicted protein |
| contig_9047_5 | 0.0022 | 0.0024 | 0.004 | 0.01 26 | 0 | 1 | 0.00 54 | predicted protein |
| contig_28483_ 6 | 0.0022 | 0.0024 | 0.0041 | 0.01 26 | 0 | 1 | 0.00 54 | predicted protein |
| contig_10829_ 2 | 0.0022 | 0.0024 | 0.0042 | 0.01 26 | 0 | 1 | 0.00 54 | plastid fructose 1 6-bisphosphatase isoform 1 |
| contig_27608_ 2 | 0.0023 | 0.0025 | 0.0042 | 0.01 29 | 0 | 1 | 0.00 55 | lysine-tRNA ligase |
| contig_5989_3 | 0.0023 | 0.0025 | 0.0042 | 0.01 29 | 0 | 1 | 0.00 55 | valyl-tRNA synthetase |
| contig_11702_ 1 | 0.0023 | 0.0025 | 0.0043 | 0.01 29 | 0 | 1 | 0.00 55 | glutamine synthetase catalytic domain protein protein |
| contig_30068_ 2 | 0.0023 | 0.0026 | 0.0043 | 0.01 29 | 0 | 1 | 0.00 55 | 40s ribosomal protein s4 |
| contig_8049_3 | 0.0023 | 0.0027 | 0.0044 | 0.01 29 | 0 | 1 | 0.00 55 | predicted protein |
| contig_9486_3 | 0.0023 | 0.0027 | 0.0046 | 0.01 3 | 0 | 1 | 0.00 55 | thiamine biosynthesis protein G (chloroplast) |
| contig_24254_ 6 | 0.0024 | 0.0028 | 0.0046 | 0.01 31 | 0 | 1 | 0.00 56 | predicted protein |
| contig_36895_ 3 | 0.0024 | 0.0028 | 0.0047 | 0.01 31 | 0 | 1 | 0.00 56 | phenylalanine-tRNA ligase beta subunit (chloroplast) |
| contig_24908_ 5 | 0.0024 | 0.0028 | 0.0048 | 0.01 32 | 0 | 1 | 0.00 56 | transitional endoplasmic reticulum ATPase TER94 |
| contig_10732_ 2 | 0.0024 | 0.0029 | 0.0048 | 0.01 33 | 0 | 1 | 0.00 57 | transaldolase |
| contig_18476_ 4 | 0.0025 | 0.003 | 0.0049 | 0.01 37 | 0 | 1 | 0.00 58 | transmembrane 9 superfamily member 4 |
| contig_13922_ 3 | 0.0025 | 0.003 | 0.0049 | 0.01 37 | 0 | 1 | 0.00 58 | annexin |
| contig_21620_ 6 | 0.0025 | 0.003 | 0.0049 | 0.01 38 | 0 | 1 | 0.00 59 | predicted protein |
| contig_12148_ 3 | 0.0026 | 0.003 | 0.0049 | 0.01 38 | 0 | 1 | 0.00 59 | predicted protein |
| contig_27843_ 1 | 0.0027 | 0.0032 | 0.005 | 0.01 44 | 0 | 1 | 0.00 61 | Fe-S cluster assembly protein SufD |
| contig_8921_1 | 0.0027 | 0.0032 | 0.005 | 0.01 44 | 0 | 1 | 0.00 61 | predicted protein |

| | | | | | | | | |
|---|---|---|---|---|---|---|---|---|
| contig_25014_3 | 0.0027 | 0.0032 | 0.005 | 0.0147 | 0 | 1 | 0.0063 | predicted protein |
| contig_28430_6 | 0.0028 | 0.0033 | 0.005 | 0.0148 | 0 | 1 | 0.0063 | predicted protein |
| contig_24813_3 | 0.0028 | 0.0033 | 0.005 | 0.015 | 0 | 1 | 0.0064 | predicted protein |
| contig_15626_6 | 0.0028 | 0.0033 | 0.0051 | 0.015 | 0 | 1 | 0.0064 | predicted protein |
| contig_24825_6 | 0.0029 | 0.0034 | 0.0052 | 0.0152 | 0 | 1 | 0.0065 | predicted protein |
| contig_28068_1 | 0.003 | 0.0034 | 0.0053 | 0.0157 | 0 | 1 | 0.0067 | UDP-glucose 6-dehydrogenase |
| contig_8868_5 | 0.003 | 0.0035 | 0.0054 | 0.0158 | 0 | 1 | 0.0067 | Enoyl-CoA hydratase |
| contig_32742_5 | 0.003 | 0.0035 | 0.0055 | 0.0158 | 0 | 1 | 0.0067 | predicted protein |
| contig_23510_1 | 0.003 | 0.0037 | 0.0055 | 0.0159 | 0 | 1 | 0.0067 | predicted protein |
| contig_40753_3 | 0.0031 | 0.0037 | 0.0055 | 0.0161 | 0 | 1 | 0.0068 | V-type proton ATPase subunit E |
| contig_40765_3 | 0.0032 | 0.0038 | 0.0056 | 0.0166 | 0 | 1 | 0.007 | triosephosphate isomerase/glyceraldehyde-3-phosphate dehydrogenase precursor |
| contig_40949_4 | 0.0032 | 0.0038 | 0.0056 | 0.0167 | 0 | 1 | 0.0071 | U3 small nucleolar RNA-associated protein 18 homolog |
| contig_9049_5 | 0.0032 | 0.0039 | 0.0058 | 0.0167 | 0 | 1 | 0.0071 | enolase |
| contig_23352_4 | 0.0033 | 0.0039 | 0.0059 | 0.0167 | 0 | 1 | 0.0071 | malate dehydrogenase |
| contig_28327_6 | 0.0033 | 0.0039 | 0.0059 | 0.0168 | 0 | 1 | 0.0072 | rab GDP dissociation inhibitor alpha |
| contig_30298_4 | 0.0033 | 0.004 | 0.0059 | 0.0168 | 0 | 1 | 0.0072 | predicted protein |
| contig_13811_4 | 0.0034 | 0.0042 | 0.006 | 0.0171 | 0 | 1 | 0.0073 | fucoxanthin chlorophyll a/c protein 8 |
| contig_5607_4 | 0.0034 | 0.0042 | 0.006 | 0.0173 | 0 | 1 | 0.0074 | protease SohB |
| contig_29863_5 | 0.0035 | 0.0042 | 0.0061 | 0.0174 | 0 | 1 | 0.0074 | Homoserine dehydrogenase [ |
| contig_26180_3 | 0.0035 | 0.0042 | 0.0062 | 0.0174 | 0 | 1 | 0.0074 | predicted protein |
| contig_32986_4 | 0.0035 | 0.0043 | 0.0062 | 0.0177 | 0 | 1 | 0.0075 | protochlorophyllide reductase magnesium chelatase subunit (chloroplast) |
| contig_25653_6 | 0.0037 | 0.0043 | 0.0063 | 0.0184 | 0 | 1 | 0.0078 | 40S ribosomal protein S8 |
| contig_25730_2 | 0.0037 | 0.0044 | 0.0063 | 0.0184 | 0 | 1 | 0.0078 | predicted protein |
| contig_27387_6 | 0.0038 | 0.0046 | 0.0065 | 0.0188 | 0 | 1 | 0.008 | GTP-binding protein YPTC1 variant |
| contig_29569_5 | 0.0038 | 0.0047 | 0.0066 | 0.0188 | 0 | 1 | 0.008 | Sedoheptulose-1 7-bisphosphatase chloroplastic |
| contig_24617_4 | 0.0039 | 0.0048 | 0.0067 | 0.019 | 0 | 1 | 0.0081 | predicted protein |

| | | | | | | | | |
|---|---|---|---|---|---|---|---|---|
| contig_33369_2 | 0.0039 | 0.0048 | 0.0067 | 0.0191 | 0 | 1 | 0.0081 | F0F1 ATP synthase subunit gamma |
| contig_30644_2 | 0.0039 | 0.0049 | 0.0067 | 0.0192 | 0 | 1 | 0.0082 | predicted protein |
| contig_40249_6 | 0.004 | 0.0049 | 0.0069 | 0.0194 | 0 | 1 | 0.0083 | translation elongation factor-like protein |
| contig_24496_1 | 0.0041 | 0.0049 | 0.0071 | 0.0199 | 0 | 1 | 0.0084 | protein RCC2 homolog |
| contig_10661_6 | 0.0042 | 0.0049 | 0.0071 | 0.0201 | 0 | 1 | 0.0086 | glutaredoxin 2 |
| contig_3037_1 | 0.0042 | 0.0049 | 0.0071 | 0.0201 | 0 | 1 | 0.0086 | ribosomal protein S4 (chloroplast) |
| contig_9858_4 | 0.0042 | 0.005 | 0.0072 | 0.0201 | 0 | 1 | 0.0086 | argininosuccinate synthase |
| contig_28237_2 | 0.0042 | 0.005 | 0.0072 | 0.0202 | 0 | 1 | 0.0086 | plant UBX domain-containing protein 10-like |
| contig_40448_3 | 0.0043 | 0.005 | 0.0072 | 0.0204 | 0 | 1 | 0.0087 | RNA-binding transcriptional accessory protein |
| contig_5881_4 | 0.0043 | 0.005 | 0.0073 | 0.0204 | 0 | 1 | 0.0087 | elongation factor Tu (chloroplast) |
| contig_13893_5 | 0.0044 | 0.0051 | 0.0076 | 0.0208 | 0 | 1 | 0.0089 | NADH dehydrogenase subunit 11 (mitochondrion) |
| contig_25275_2 | 0.0046 | 0.0052 | 0.0077 | 0.0217 | 0 | 1 | 0.0092 | histidine-tRNA ligase |
| contig_7641_5 | 0.0046 | 0.0053 | 0.0079 | 0.0218 | 0 | 1 | 0.0093 | precursor cytochrome c6 |
| contig_11304_3 | 0.0047 | 0.0054 | 0.0079 | 0.0224 | 0 | 1 | 0.0095 | predicted protein |
| contig_23649_2 | 0.0048 | 0.0055 | 0.008 | 0.0225 | 0 | 1 | 0.0096 | heat shock protein 90-2 |
| contig_16696_5 | 0.0048 | 0.0055 | 0.008 | 0.0225 | 0 | 1 | 0.0096 | pyruvate kinase |
| contig_34814_4 | 0.0049 | 0.0056 | 0.0082 | 0.0227 | 0 | 1 | 0.0097 | predicted protein |
| contig_26554_3 | 0.0049 | 0.0056 | 0.0082 | 0.0228 | 0 | 1 | 0.0097 | predicted protein |
| contig_25971_2 | 0.0049 | 0.0057 | 0.0082 | 0.0228 | 0 | 1 | 0.0097 | purine nucleoside phosphorylase I inosine and guanosine-specific |
| contig_25670_6 | 0.0049 | 0.0058 | 0.0084 | 0.0228 | 0 | 1 | 0.0097 | predicted protein |
| contig_29484_4 | 0.0049 | 0.0059 | 0.0085 | 0.0228 | 0 | 1 | 0.0097 | serine hydroxymethyltransferase |
| contig_28209_6 | 0.005 | 0.0059 | 0.0088 | 0.0228 | 0 | 1 | 0.0097 | dihydrolipoamide acetyltransferase |
| contig_6528_1 | 0.005 | 0.0059 | 0.009 | 0.0228 | 0 | 1 | 0.0097 | 40S ribosomal protein S19-3 |
| contig_24980_4 | 0.005 | 0.006 | 0.0091 | 0.0228 | 0 | 1 | 0.0097 | Bystin |
| contig_28795_5 | 0.005 | 0.006 | 0.0091 | 0.0229 | 0 | 1 | 0.0097 | protein transport protein SEC23 |
| contig_19536_5 | 0.005 | 0.0062 | 0.0093 | 0.0229 | 0 | 1 | 0.0097 | methionine S-adenosyl transferase |
| contig_6178_4 | 0.0051 | 0.0062 | 0.0093 | 0.0229 | 0 | 1 | 0.0098 | rubisco expression protein (chloroplast) |

| | | | | | | | | |
|---|---|---|---|---|---|---|---|---|
| contig_25157_1 | 0.0052 | 0.0063 | 0.0093 | 0.0237 | 0 | 1 | 0.0101 | magnesium chelatase subunit D |
| contig_27994_5 | 0.0053 | 0.0063 | 0.0096 | 0.0238 | 0 | 1 | 0.0101 | nitrate transporter |
| contig_28089_3 | 0.0054 | 0.0066 | 0.0097 | 0.0243 | 0 | 1 | 0.0103 | threonine synthase |
| contig_17027_5 | 0.0055 | 0.0066 | 0.0098 | 0.0244 | 0 | 1 | 0.0104 | predicted protein |
| contig_17744_4 | 0.0055 | 0.0067 | 0.0098 | 0.0246 | 0 | 1 | 0.0105 | integral membrane protein |
| contig_9284_2 | 0.0055 | 0.0067 | 0.01 | 0.0247 | 0 | 1 | 0.0105 | 60S ribosomal protein L12 |
| contig_38239_5 | 0.0056 | 0.0067 | 0.01 | 0.0247 | 0 | 1 | 0.0105 | predicted protein |
| contig_22850_1 | 0.0056 | 0.0069 | 0.0101 | 0.0247 | 0 | 1 | 0.0105 | nitrate transporter |
| contig_21794_2 | 0.0057 | 0.0071 | 0.0102 | 0.0253 | 0 | 1 | 0.0108 | fucoxanthin chlorophyll a/c protein deviant |
| contig_27013_6 | 0.0058 | 0.0071 | 0.0104 | 0.0253 | 0 | 1 | 0.0108 | sulfite reductase ferredoxin dependent |
| contig_39855_1 | 0.0059 | 0.0072 | 0.0104 | 0.0258 | 0 | 1 | 0.011 | predicted protein |
| contig_39759_2 | 0.0059 | 0.0072 | 0.0104 | 0.0259 | 0 | 1 | 0.011 | antibiotic biosynthesis monooxygenase |
| contig_13881_2 | 0.0059 | 0.0072 | 0.0105 | 0.0259 | 0 | 1 | 0.011 | predicted protein |
| contig_35584_4 | 0.006 | 0.0073 | 0.0108 | 0.0259 | 0 | 1 | 0.011 | nitrate transporter |
| contig_6613_6 | 0.006 | 0.0073 | 0.0109 | 0.0259 | 0 | 1 | 0.011 | sodium-coupled neutral amino acid transporter |
| contig_34797_4 | 0.0061 | 0.0076 | 0.0113 | 0.0263 | 0 | 1 | 0.0112 | GTP-binding protein SAR1 |
| contig_37899_2 | 0.0062 | 0.0079 | 0.0113 | 0.0266 | 0 | 1 | 0.0113 | ribosomal protein L2 (chloroplast) |
| contig_38142_5 | 0.0062 | 0.0079 | 0.0115 | 0.0268 | 0 | 1 | 0.0114 | ATP synthase F1 subunit alpha (mitochondrion) |
| contig_16670_4 | 0.0063 | 0.008 | 0.0115 | 0.027 | 0 | 1 | 0.0115 | porphobilinogen deaminase |
| contig_15137_6 | 0.0063 | 0.008 | 0.0115 | 0.0271 | 0 | 1 | 0.0115 | aspartate aminotransferase |
| contig_8803_5 | 0.0065 | 0.0082 | 0.0117 | 0.0279 | 0 | 1 | 0.0119 | predicted protein |
| contig_26157_4 | 0.0066 | 0.0082 | 0.0117 | 0.0279 | 0 | 1 | 0.0119 | predicted protein |
| contig_40093_3 | 0.0066 | 0.0082 | 0.0118 | 0.0281 | 0 | 1 | 0.012 | predicted protein |
| contig_26589_3 | 0.0067 | 0.0084 | 0.0118 | 0.0281 | 0 | 1 | 0.012 | predicted protein |
| contig_34578_1 | 0.0067 | 0.009 | 0.0119 | 0.0281 | 0 | 1 | 0.012 | predicted protein |
| contig_27826_5 | 0.0067 | 0.009 | 0.012 | 0.0281 | 0 | 1 | 0.012 | predicted protein |
| contig_34524_6 | 0.0069 | 0.0091 | 0.012 | 0.029 | 0 | 1 | 0.0123 | ---NA--- |

| | | | | | | | | |
|---|---|---|---|---|---|---|---|---|
| contig_27073_1 | 0.0071 | 0.0091 | 0.012 | 0.0296 | 0 | 1 | 0.0126 | Phospholipid hydroperoxide glutathione peroxidase |
| contig_30021_2 | 0.0071 | 0.0092 | 0.012 | 0.0296 | 0 | 1 | 0.0126 | predicted protein |
| contig_7686_2 | 0.0071 | 0.0093 | 0.0122 | 0.0296 | 0 | 1 | 0.0126 | glutathione-disulfide reductase |
| contig_8539_5 | 0.0072 | 0.0093 | 0.0125 | 0.0299 | 0 | 1 | 0.0127 | SET domain-containing protein |
| contig_30052_6 | 0.0072 | 0.0097 | 0.0127 | 0.0299 | 0 | 1 | 0.0127 | predicted protein |
| contig_35394_6 | 0.0072 | 0.0097 | 0.0128 | 0.0299 | 0 | 1 | 0.0127 | cytochrome c |
| contig_20131_5 | 0.0073 | 0.0098 | 0.013 | 0.0299 | 0 | 1 | 0.0127 | 60S ribosomal protein L21-A |
| contig_29420_3 | 0.0073 | 0.01 | 0.013 | 0.0301 | 0 | 1 | 0.0128 | thylakoid lumenal 15.0 kDa protein 2 chloroplastic |
| contig_10116_3 | 0.0076 | 0.01 | 0.013 | 0.031 | 0 | 1 | 0.0132 | peptidyl-prolyl cis-trans isomerase CYP19-4 |
| contig_11625_6 | 0.0077 | 0.01 | 0.0131 | 0.0315 | 0 | 1 | 0.0134 | T9SS type A sorting domain-containing protein |
| contig_11343_2 | 0.0079 | 0.0101 | 0.0133 | 0.0322 | 0 | 1 | 0.0137 | probable aspartyl aminopeptidase |
| contig_33347_6 | 0.0079 | 0.0102 | 0.0134 | 0.0322 | 0 | 1 | 0.0137 | V-type ATPase A subunit |
| contig_9651_3 | 0.008 | 0.0104 | 0.0137 | 0.0322 | 0 | 1 | 0.0137 | ribosomal protein RPL17 |
| contig_24729_6 | 0.008 | 0.0104 | 0.0137 | 0.0322 | 0 | 1 | 0.0137 | predicted protein |
| contig_24353_3 | 0.0082 | 0.0105 | 0.0137 | 0.0328 | 0 | 1 | 0.014 | hemagglutinin/amebocyte aggregation factor |
| contig_35362_1 | 0.0082 | 0.0108 | 0.0138 | 0.0329 | 0 | 1 | 0.014 | nitrate transporter |
| contig_22533_3 | 0.0082 | 0.0108 | 0.0151 | 0.0329 | 0 | 1 | 0.014 | predicted protein |
| contig_27705_5 | 0.0082 | 0.0113 | 0.0152 | 0.0329 | 0 | 1 | 0.014 | predicted protein |
| contig_6700_4 | 0.0084 | 0.0115 | 0.0154 | 0.0336 | 0 | 1 | 0.0143 | agmatinase |
| contig_18233_1 | 0.0085 | 0.0115 | 0.0154 | 0.034 | 0 | 1 | 0.0144 | dihydrolipoamide dehydrogenase |
| contig_24978_5 | 0.0088 | 0.0115 | 0.0154 | 0.0349 | 0 | 1 | 0.0148 | predicted protein |
| contig_27172_4 | 0.009 | 0.0117 | 0.0156 | 0.0357 | 0 | 1 | 0.0152 | peptidyl-prolyl cis-trans isomerase |
| contig_30065_5 | 0.0091 | 0.0117 | 0.0157 | 0.0358 | 0 | 1 | 0.0152 | importin subunit alpha-1-like isoform X2 |
| contig_24050_3 | 0.0091 | 0.0117 | 0.0158 | 0.0358 | 0 | 1 | 0.0152 | enolase-phosphatase E1 |
| contig_15357_4 | 0.0092 | 0.0118 | 0.0162 | 0.0361 | 0 | 1 | 0.0153 | calreticulin |
| contig_40568_2 | 0.0093 | 0.0118 | 0.0162 | 0.0363 | 0 | 1 | 0.0154 | glutathione reductase |
| contig_20792_5 | 0.0093 | 0.012 | 0.0165 | 0.0363 | 0 | 1 | 0.0155 | ---NA--- |

| | | | | | | | | |
|---|---|---|---|---|---|---|---|---|
| contig_30700_3 | 0.0093 | 0.012 | 0.0169 | 0.0364 | 0 | 1 | 0.0155 | predicted protein |
| contig_9105_2 | 0.0096 | 0.012 | 0.0169 | 0.0372 | 0 | 1 | 0.0158 | SufE-like protein 1 chloroplastic/mitochondrial |
| contig_27261_5 | 0.0097 | 0.012 | 0.0177 | 0.0376 | 0 | 1 | 0.016 | enoyl-CoA hydratase |
| contig_9136_5 | 0.0098 | 0.0122 | 0.0178 | 0.0378 | 0 | 1 | 0.0161 | predicted protein |
| contig_28761_2 | 0.0098 | 0.0123 | 0.0178 | 0.0378 | 0 | 1 | 0.0161 | protein dispatched homolog 3-like |
| contig_29454_5 | 0.01 | 0.0125 | 0.0181 | 0.0381 | 0 | 1 | 0.0162 | peptidylprolyl isomerase |
| contig_16230_2 | 0.01 | 0.013 | 0.0184 | 0.0381 | 0 | 1 | 0.0162 | phosphoglycerate mutase |
| contig_6711_4 | 0.01 | 0.013 | 0.0185 | 0.0381 | 0 | 1 | 0.0162 | predicted protein |
| contig_5740_5 | 0.0101 | 0.013 | 0.0187 | 0.0385 | 0 | 1 | 0.0164 | predicted protein |
| contig_16905_4 | 0.0102 | 0.013 | 0.0189 | 0.0389 | 0 | 1 | 0.0166 | predicted protein |
| contig_32280_2 | 0.0104 | 0.013 | 0.0189 | 0.0394 | 0 | 1 | 0.0167 | plastid sedoheptulose-1 7-bisphosphatase |
| contig_28172_4 | 0.0104 | 0.0133 | 0.0196 | 0.0394 | 0 | 1 | 0.0167 | 60S ribosomal protein L5 |
| contig_29442_5 | 0.0104 | 0.0134 | 0.0197 | 0.0394 | 0 | 1 | 0.0167 | eukaryotic translation initiation factor 2 subunit alpha |
| contig_5908_5 | 0.0105 | 0.0137 | 0.0198 | 0.0397 | 0 | 1 | 0.0169 | predicted protein |
| contig_15399_2 | 0.0108 | 0.0137 | 0.0198 | 0.0404 | 0 | 1 | 0.0172 | predicted protein |
| contig_33426_1 | 0.0108 | 0.0138 | 0.0199 | 0.0407 | 0 | 1 | 0.0173 | predicted protein |
| contig_20909_4 | 0.0109 | 0.0143 | 0.0201 | 0.0408 | 0 | 1 | 0.0174 | 40S ribosomal protein |
| contig_9059_3 | 0.0113 | 0.0152 | 0.0203 | 0.0422 | 0 | 1 | 0.0179 | predicted protein |
| contig_8365_3 | 0.0113 | 0.0152 | 0.0203 | 0.0422 | 0 | 1 | 0.018 | pyruvate dehydrogenase |
| contig_26535_2 | 0.0115 | 0.0154 | 0.0205 | 0.0424 | 0 | 1 | 0.018 | tubulin alpha chain |
| contig_9753_3 | 0.0115 | 0.0154 | 0.0207 | 0.0424 | 0 | 1 | 0.018 | predicted protein |
| contig_14932_3 | 0.0115 | 0.0156 | 0.0207 | 0.0424 | 0 | 1 | 0.018 | predicted protein |
| contig_40102_2 | 0.0117 | 0.0157 | 0.0208 | 0.0431 | 0 | 1 | 0.0183 | predicted protein |
| contig_19770_6 | 0.0117 | 0.0158 | 0.021 | 0.0431 | 0 | 1 | 0.0183 | predicted protein |
| contig_34506_1 | 0.0117 | 0.0162 | 0.021 | 0.0431 | 0 | 1 | 0.0183 | cell division protein FtsH |
| contig_17663_5 | 0.0118 | 0.0165 | 0.0213 | 0.0432 | 0 | 1 | 0.0184 | pyruvate dehydrogenase (acetyl-transferring) E1 component alpha subunit |

| | | | | | | | | |
|---|---|---|---|---|---|---|---|---|
| contig_37776_1 | 0.0118 | 0.0169 | 0.0213 | 0.0432 | 0 | 1 | 0.0184 | DNA polymerase |
| contig_5469_5 | 0.0119 | 0.0169 | 0.0213 | 0.0434 | 0 | 1 | 0.0185 | ribosomal protein S2 (chloroplast) |
| contig_30325_5 | 0.012 | 0.0177 | 0.0217 | 0.0434 | 0 | 1 | 0.0185 | predicted protein |
| contig_5781_3 | 0.012 | 0.0178 | 0.0218 | 0.0434 | 0 | 1 | 0.0185 | acetyl-coenzyme a synthetase |
| contig_28094_2 | 0.012 | 0.0178 | 0.0221 | 0.0434 | 0 | 1 | 0.0185 | agmatinase |
| contig_25890_5 | 0.012 | 0.0184 | 0.0225 | 0.0434 | 0 | 1 | 0.0185 | acetolactate synthase catalytic subunit |
| contig_8125_5 | 0.0122 | 0.0184 | 0.023 | 0.0439 | 0 | 1 | 0.0187 | adenosylhomocysteinase |
| contig_15161_3 | 0.0123 | 0.0187 | 0.0233 | 0.0442 | 0 | 1 | 0.0188 | serine/threonine-protein phosphatase PP1-beta catalytic subunit |
| contig_39771_2 | 0.0125 | 0.0188 | 0.0234 | 0.0448 | 0 | 1 | 0.0191 | hypothetical protein THAOC_36769 |
| contig_6246_4 | 0.0127 | 0.0188 | 0.0234 | 0.0455 | 0 | 1 | 0.0193 | predicted protein |
| contig_34816_5 | 0.0128 | 0.0189 | 0.0236 | 0.0455 | 0 | 1 | 0.0194 | 40S ribosomal protein S12 |
| contig_35273_1 | 0.013 | 0.0198 | 0.0237 | 0.0461 | 0 | 1 | 0.0196 | Dethiobiotin synthase |
| contig_24237_6 | 0.013 | 0.0198 | 0.024 | 0.0461 | 0 | 1 | 0.0196 | predicted protein |
| contig_14784_2 | 0.013 | 0.0198 | 0.0241 | 0.0462 | 0 | 1 | 0.0196 | atp-dependent RNA helicase |
| contig_32836_4 | 0.0131 | 0.0199 | 0.0245 | 0.0462 | 0 | 1 | 0.0196 | predicted protein |
| contig_5475_2 | 0.0132 | 0.0201 | 0.0245 | 0.0466 | 0 | 1 | 0.0198 | coatomer subunit epsilon |
| contig_22608_4 | 0.0133 | 0.0204 | 0.0246 | 0.0468 | 0 | 1 | 0.0199 | predicted protein |
| contig_24923_5 | 0.0134 | 0.0205 | 0.0247 | 0.047 | 0 | 1 | 0.02 | voltage activated chloride channel CLC7 type |
| contig_29933_6 | 0.0137 | 0.0205 | 0.0247 | 0.0478 | 0 | 1 | 0.0204 | 30S ribosomal protein S1 |
| contig_6039_6 | 0.0137 | 0.0207 | 0.025 | 0.0478 | 0 | 1 | 0.0204 | large subunit ribosomal protein L27Ae |
| contig_39521_4 | 0.0137 | 0.0207 | 0.0251 | 0.0478 | 0 | 1 | 0.0204 | predicted protein |
| contig_25113_4 | 0.0138 | 0.021 | 0.0251 | 0.048 | 0 | 1 | 0.0204 | YEATS domain-containing protein 4 |
| contig_25363_4 | 0.0143 | 0.021 | 0.0252 | 0.0495 | 0 | 1 | 0.0211 | ABC-transporter like protein |
| contig_32611_6 | 0.0149 | 0.0212 | 0.0252 | 0.0517 | 0 | 1 | 0.022 | predicted protein |
| contig_40284_6 | 0.0151 | 0.0213 | 0.0252 | 0.0521 | 0 | 1 | 0.0222 | large subunit ribosomal protein L13e |
| contig_24234_3 | 0.0152 | 0.0213 | 0.0253 | 0.0524 | 0 | 1 | 0.0223 | predicted protein |

| | | | | | | | | |
|---|---|---|---|---|---|---|---|---|
| contig_28019_5 | 0.0154 | 0.0217 | 0.0258 | 0.0527 | 0 | 1 | 0.0224 | NAD+ synthetase |
| contig_30107_6 | 0.0154 | 0.0217 | 0.0258 | 0.0527 | 0 | 1 | 0.0224 | ATP-dependent protease HslVU ATPase subunit |
| contig_2430_4 | 0.0154 | 0.022 | 0.026 | 0.0527 | 0 | 1 | 0.0224 | ubiquinol-cytochrome c r |
| contig_29925_5 | 0.0156 | 0.0225 | 0.0265 | 0.0531 | 0 | 1 | 0.0226 | glutamine synthetase |
| contig_37814_1 | 0.0156 | 0.023 | 0.0268 | 0.0533 | 0 | 1 | 0.0227 | ribosomal protein S3 (chloroplast) |
| contig_10817_6 | 0.0157 | 0.0234 | 0.0269 | 0.0533 | 0 | 1 | 0.0227 | silicon efflux transporter LSI2 |
| contig_28745_4 | 0.0158 | 0.0234 | 0.0274 | 0.0534 | 0 | 1 | 0.0227 | glutamyl-trna reductase |
| contig_40835_6 | 0.0162 | 0.0236 | 0.0275 | 0.0546 | 0 | 1 | 0.0232 | 60S ribosomal protein L35a-4 |
| contig_22730_1 | 0.0162 | 0.0236 | 0.0282 | 0.0546 | 0 | 1 | 0.0232 | predicted protein |
| contig_35315_5 | 0.0164 | 0.0236 | 0.0282 | 0.0553 | 0 | 1 | 0.0235 | ---NA--- |
| contig_14542_6 | 0.0165 | 0.0237 | 0.0282 | 0.0556 | 0 | 1 | 0.0236 | predicted protein |
| contig_24490_6 | 0.0169 | 0.0245 | 0.0284 | 0.0565 | 0 | 1 | 0.0241 | predicted protein |
| contig_27293_4 | 0.0173 | 0.0245 | 0.0289 | 0.058 | 0 | 1 | 0.0247 | predicted protein |
| contig_14496_5 | 0.0176 | 0.0246 | 0.0289 | 0.0589 | 0 | 1 | 0.0251 | Diaminopimelate epimerase |
| contig_35287_3 | 0.0177 | 0.0247 | 0.0291 | 0.059 | 0 | 1 | 0.0251 | photosystem II PsbU protein |
| contig_34823_4 | 0.0178 | 0.0249 | 0.0311 | 0.059 | 0 | 1 | 0.0251 | 60S ribosomal protein L18-2 |
| contig_5829_1 | 0.0178 | 0.025 | 0.0315 | 0.059 | 0 | 1 | 0.0251 | gametolysin peptidase M11 |
| contig_17520_3 | 0.0181 | 0.0251 | 0.0321 | 0.0599 | 0 | 1 | 0.0255 | methyltransferase FkbM family (macronuclear) |
| contig_39623_1 | 0.0181 | 0.0251 | 0.0321 | 0.0599 | 0 | 1 | 0.0255 | predicted protein |
| contig_7261_5 | 0.0184 | 0.0252 | 0.0323 | 0.0605 | 0 | 1 | 0.0257 | ubiquitin-conjugating enzyme E2 2 |
| contig_29950_5 | 0.0185 | 0.0252 | 0.0324 | 0.0609 | 0 | 1 | 0.0259 | Fe-S cluster assembly protein SufD |
| contig_32695_4 | 0.0187 | 0.0252 | 0.0328 | 0.0614 | 0 | 1 | 0.0261 | predicted protein |
| contig_30091_6 | 0.0188 | 0.0253 | 0.0328 | 0.0617 | 0 | 1 | 0.0262 | predicted protein |
| contig_26083_5 | 0.0189 | 0.0258 | 0.0328 | 0.0617 | 0 | 1 | 0.0262 | phosphatidylinositol transfer protein 3-like |
| contig_8012_5 | 0.0196 | 0.0259 | 0.0331 | 0.0638 | 0 | 1 | 0.0271 | predicted protein |
| contig_27789_6 | 0.0196 | 0.026 | 0.0335 | 0.0638 | 0 | 1 | 0.0271 | Mitochondrial-processing peptidase subunit alpha |
| contig_11460_2 | 0.0197 | 0.026 | 0.0335 | 0.0638 | 0 | 1 | 0.0272 | predicted protein |

| | | | | | | | | |
|---|---|---|---|---|---|---|---|---|
| contig_22469_6 | 0.0197 | 0.0265 | 0.0336 | 0.0638 | 0 | 1 | 0.0272 | predicted protein |
| contig_15395_3 | 0.0198 | 0.0273 | 0.0347 | 0.0639 | 0 | 1 | 0.0272 | predicted protein |
| contig_35069_1 | 0.0198 | 0.0274 | 0.0353 | 0.0639 | 0 | 1 | 0.0272 | translation elongation factor-like protein |
| contig_23045_4 | 0.0199 | 0.0282 | 0.0362 | 0.064 | 0 | 1 | 0.0272 | amino acid/polyamine transporter |
| contig_37814_3 | 0.0201 | 0.0282 | 0.0364 | 0.0646 | 0 | 1 | 0.0275 | ribosomal protein S3 (chloroplast) |
| contig_8461_2 | 0.0203 | 0.0282 | 0.0368 | 0.0651 | 0 | 1 | 0.0277 | SDR family oxidoreductase |
| contig_17360_5 | 0.0204 | 0.0284 | 0.0369 | 0.0651 | 0 | 1 | 0.0277 | Annexin protein |
| contig_16000_5 | 0.0205 | 0.0289 | 0.0373 | 0.0654 | 0 | 1 | 0.0278 | FAD-dependent oxidoreductase |
| contig_27905_6 | 0.0205 | 0.0289 | 0.0373 | 0.0654 | 0 | 1 | 0.0278 | predicted protein |
| contig_16388_6 | 0.0207 | 0.0291 | 0.0396 | 0.0657 | 0 | 1 | 0.028 | predicted protein |
| contig_7129_5 | 0.0207 | 0.0311 | 0.0396 | 0.0657 | 0 | 1 | 0.028 | T9SS type A sorting domain-containing protein |
| contig_22875_4 | 0.0208 | 0.0318 | 0.04 | 0.066 | 0 | 1 | 0.0281 | calcium-translocating P-type ATPase PMCA-type |
| contig_6155_1 | 0.021 | 0.0322 | 0.0401 | 0.0662 | 0 | 1 | 0.0282 | light harvesting protein |
| contig_39450_1 | 0.021 | 0.0323 | 0.0403 | 0.0662 | 0 | 1 | 0.0282 | CTL-like protein DDB_G0288717 |
| contig_23548_3 | 0.0212 | 0.0324 | 0.0403 | 0.0666 | 0 | 1 | 0.0284 | ATP synthase subunit beta mitochondrial |
| contig_20045_5 | 0.0213 | 0.0328 | 0.0407 | 0.0666 | 0 | 1 | 0.0284 | AP-1 complex subunit sigma-1 |
| contig_28144_3 | 0.0213 | 0.0328 | 0.041 | 0.0666 | 0 | 1 | 0.0284 | ---NA--- |
| contig_24655_5 | 0.0215 | 0.0328 | 0.041 | 0.0673 | 0 | 1 | 0.0286 | predicted protein |
| contig_10461_4 | 0.0217 | 0.0334 | 0.0412 | 0.0676 | 0 | 1 | 0.0288 | ATP synthase subunit beta mitochondrial |
| contig_19765_6 | 0.0218 | 0.0335 | 0.0418 | 0.0679 | 0 | 1 | 0.0289 | predicted protein |
| contig_12419_1 | 0.022 | 0.0336 | 0.0418 | 0.0683 | 0 | 1 | 0.0291 | T-complex protein 1 subunit delta |
| contig_24447_4 | 0.0221 | 0.0347 | 0.0421 | 0.0686 | 0 | 1 | 0.0292 | diaminopimelate decarboxylase |
| contig_10429_5 | 0.0225 | 0.0353 | 0.0426 | 0.0697 | 0 | 1 | 0.0297 | hypothetical protein T265_11290 partial |
| contig_37770_2 | 0.0229 | 0.0364 | 0.0435 | 0.0707 | 0 | 1 | 0.0301 | photosystem I P700 apoprotein A1 (chloroplast) |
| contig_5576_4 | 0.023 | 0.0364 | 0.0436 | 0.071 | 0 | 1 | 0.0302 | transketolase |
| contig_6178_5 | 0.0233 | 0.0364 | 0.0436 | 0.0717 | 0 | 1 | 0.0305 | rubisco expression protein (chloroplast) |
| contig_21411_6 | 0.0234 | 0.0369 | 0.0439 | 0.0717 | 0 | 1 | 0.0305 | ATP-dependent RNA helicase eIF4A |

| | | | | | | | | |
|---|---|---|---|---|---|---|---|---|
| contig_9065_4 | 0.0234 | 0.0373 | 0.0441 | 0.07 18 | 0 | 1 | 0.03 05 | tRNA cyclic N6-threonylcarbamoyladenosine(37) synthase TcdA |
| contig_18863_ 3 | 0.0236 | 0.0386 | 0.0446 | 0.07 21 | 0 | 1 | 0.03 07 | nucleoside diphosphate kinase |
| contig_30131_ 2 | 0.0236 | 0.0396 | 0.0458 | 0.07 21 | 0 | 1 | 0.03 07 | dihydrolipoyl dehydrogenase |
| contig_22476_ 2 | 0.0237 | 0.04 | 0.0466 | 0.07 22 | 0 | 1 | 0.03 07 | predicted protein |
| contig_14395_ 5 | 0.024 | 0.0401 | 0.0472 | 0.07 31 | 0 | 1 | 0.03 11 | manganese superoxide dismutase |
| contig_23995_ 6 | 0.0241 | 0.0403 | 0.0481 | 0.07 31 | 0 | 1 | 0.03 11 | predicted protein |
| contig_18892_ 6 | 0.0245 | 0.0403 | 0.0495 | 0.07 39 | 0 | 1 | 0.03 15 | predicted protein |
| contig_37718_ 5 | 0.0245 | 0.0403 | 0.0495 | 0.07 39 | 0 | 1 | 0.03 15 | predicted protein |
| contig_18966_ 2 | 0.0245 | 0.041 | 0.05 | 0.07 39 | 0 | 1 | 0.03 15 | predicted protein |
| contig_24988_ 5 | 0.0246 | 0.0411 | 0.0509 | 0.07 39 | 0 | 1 | 0.03 15 | Urea-proton symporter DUR3 |
| contig_6077_5 | 0.0247 | 0.0412 | 0.0535 | 0.07 43 | 0 | 1 | 0.03 16 | 60s ribosomal protein l9 |
| contig_38355_ 2 | 0.0249 | 0.0418 | 0.0553 | 0.07 46 | 0 | 1 | 0.03 17 | predicted protein |
| contig_25680_ 5 | 0.025 | 0.0421 | 0.0553 | 0.07 46 | 0 | 1 | 0.03 17 | glucose-6-phosphate isomerase |
| contig_5881_5 | 0.025 | 0.0422 | 0.0553 | 0.07 46 | 0 | 1 | 0.03 17 | elongation factor Tu (chloroplast) |
| contig_25854_ 4 | 0.0251 | 0.0435 | 0.0587 | 0.07 46 | 0 | 1 | 0.03 17 | L30e-like protein |
| contig_20747_ 6 | 0.0251 | 0.0435 | 0.0587 | 0.07 46 | 0 | 1 | 0.03 17 | translation initiation factor 4E |
| contig_33176_ 3 | 0.0252 | 0.0437 | 0.0598 | 0.07 46 | 0 | 1 | 0.03 17 | EF-hand domain-containing protein |
| contig_10410_ 5 | 0.0252 | 0.044 | 0.0606 | 0.07 46 | 0 | 1 | 0.03 17 | LL-diaminopimelate aminotransferase |
| contig_30185_ 1 | 0.0252 | 0.0441 | 0.0609 | 0.07 46 | 0 | 1 | 0.03 17 | signal peptidase I |
| contig_9849_4 | 0.0253 | 0.0446 | 0.061 | 0.07 46 | 0 | 1 | 0.03 18 | chloroplast sedoheptulose-1 7-bisphosphatase |
| contig_21628_ 5 | 0.0257 | 0.046 | 0.0625 | 0.07 58 | 0 | 1 | 0.03 22 | Tubulin alpha-5 chain |
| contig_24220_ 6 | 0.0258 | 0.0467 | 0.0633 | 0.07 58 | 0 | 1 | 0.03 22 | ATP-binding cassette superfamily |
| contig_11577_ 6 | 0.0259 | 0.0481 | 0.064 | 0.07 6 | 0 | 1 | 0.03 23 | Mitochondrial adenine nucleotide transporter ADNT1 |
| contig_18043_ 1 | 0.026 | 0.0497 | 0.0645 | 0.07 61 | 0 | 1 | 0.03 24 | Light-harvesting complex I LH38 protein |
| contig_14305_ 2 | 0.0265 | 0.0497 | 0.0652 | 0.07 74 | 0 | 1 | 0.03 29 | predicted protein |
| contig_34082_ 5 | 0.0265 | 0.05 | 0.0653 | 0.07 74 | 0 | 1 | 0.03 29 | predicted protein |

| | | | | | | | | |
|---|---|---|---|---|---|---|---|---|
| contig_29877_2 | 0.0268 | 0.0535 | 0.0656 | 0.0781 | 0 | 1 | 0.0332 | predicted protein |
| contig_33257_5 | 0.0269 | 0.0553 | 0.0657 | 0.0782 | 0 | 1 | 0.0333 | SulP family inorganic anion transporter |
| contig_17529_2 | 0.0273 | 0.0553 | 0.0658 | 0.0793 | 0 | 1 | 0.0337 | ribose-phosphate pyrophosphokinase |
| contig_23908_1 | 0.0274 | 0.0587 | 0.0676 | 0.0793 | 0 | 1 | 0.0337 | mitochondrial carrier protein |
| contig_30812_2 | 0.0275 | 0.059 | 0.0677 | 0.0795 | 0 | 1 | 0.0338 | citrate synthase |
| contig_33573_6 | 0.0282 | 0.0599 | 0.0678 | 0.081 | 0 | 1 | 0.0345 | ---NA--- |
| contig_29760_5 | 0.0282 | 0.0606 | 0.0698 | 0.081 | 0 | 1 | 0.0345 | predicted protein |
| contig_33090_6 | 0.0282 | 0.0609 | 0.0726 | 0.081 | 0 | 1 | 0.0345 | predicted protein |
| contig_5770_5 | 0.0282 | 0.061 | 0.0731 | 0.081 | 0 | 1 | 0.0345 | ketol-acid reductoisomerase |
| contig_21753_2 | 0.0284 | 0.0633 | 0.0732 | 0.0812 | 0 | 1 | 0.0346 | glyceraldehyde-3-phosphate dehydrogenase |
| contig_12107_3 | 0.0287 | 0.064 | 0.0737 | 0.0821 | 0 | 1 | 0.0349 | argininosuccinate synthase |
| contig_14563_3 | 0.0289 | 0.0645 | 0.0753 | 0.0823 | 0 | 1 | 0.035 | predicted protein |
| contig_38505_3 | 0.0289 | 0.0652 | 0.0777 | 0.0823 | 0 | 1 | 0.035 | serine/threonine-protein phosphatase 2A regulatory subunit B' |
| contig_25682_4 | 0.0291 | 0.0653 | 0.0785 | 0.0826 | 0 | 1 | 0.0351 | Immunoglobulin G-binding protein A |
| contig_38588_6 | 0.0306 | 0.0657 | 0.0797 | 0.0868 | 0 | 1 | 0.0369 | Ras-related protein Rab-8A |
| contig_28255_4 | 0.0308 | 0.0657 | 0.0809 | 0.0873 | 0 | 1 | 0.0372 | long chain acyl-coa synthetase |
| contig_39581_5 | 0.031 | 0.0658 | 0.0812 | 0.0875 | 0 | 1 | 0.0373 | cytochrome c1 |
| contig_8557_6 | 0.0311 | 0.0677 | 0.0815 | 0.0878 | 0 | 1 | 0.0373 | cytochrome b6 (chloroplast) |
| contig_22744_5 | 0.0315 | 0.0678 | 0.0821 | 0.0886 | 0 | 1 | 0.0377 | predicted protein |
| contig_26227_3 | 0.0318 | 0.0678 | 0.0824 | 0.0892 | 0 | 1 | 0.038 | 3-phosphoshikimate 1-carboxyvinyltransferase |
| contig_29269_4 | 0.0321 | 0.0726 | 0.0827 | 0.0899 | 0 | 1 | 0.0383 | predicted protein |
| contig_19001_2 | 0.0322 | 0.0731 | 0.0831 | 0.0901 | 0 | 1 | 0.0383 | predicted protein |
| contig_30861_4 | 0.0323 | 0.0732 | 0.0833 | 0.0901 | 0 | 1 | 0.0384 | fucoxanthin chlorophyll a/c protein 8 |
| contig_10045_6 | 0.0324 | 0.0737 | 0.0841 | 0.0904 | 0 | 1 | 0.0384 | dihydrolipoamide dehydrogenase |
| contig_27614_3 | 0.0325 | 0.0777 | 0.0845 | 0.0906 | 0 | 1 | 0.0386 | Regulator of nonsense transcripts 1-like protein |
| contig_34535_1 | 0.0328 | 0.0785 | 0.0854 | 0.0909 | 0 | 1 | 0.0387 | eukaryotic translation initiation factor 2 subunit 3 |

| | | | | | | | | |
|---|---|---|---|---|---|---|---|---|
| contig_5529_5 | 0.0328 | 0.0797 | 0.0861 | 0.09 09 | 0 | 1 | 0.03 87 | photosystem II protein D1 (chloroplast) |
| contig_16936_ 1 | 0.0328 | 0.0809 | 0.0877 | 0.09 09 | 0 | 1 | 0.03 87 | H/ACA ribonucleoprotein complex subunit 4 |
| contig_18025_ 5 | 0.0331 | 0.0814 | 0.0877 | 0.09 14 | 0 | 1 | 0.03 89 | histone H2B |
| contig_5318_4 | 0.0334 | 0.0815 | 0.0894 | 0.09 22 | 0 | 1 | 0.03 92 | 40S ribosomal protein S15a |
| contig_39034_ 5 | 0.0335 | 0.0821 | 0.0901 | 0.09 23 | 0 | 1 | 0.03 93 | predicted protein |
| contig_14405_ 3 | 0.0336 | 0.0827 | 0.0904 | 0.09 23 | 0 | 1 | 0.03 93 | predicted protein |
| contig_10966_ 3 | 0.0342 | 0.0829 | 0.0908 | 0.09 39 | 0 | 1 | 0.04 | predicted protein |
| contig_29776_ 5 | 0.0343 | 0.0833 | 0.0909 | 0.09 41 | 0 | 1 | 0.04 | fructose-bisphosphate aldolase |
| contig_15155_ 6 | 0.0347 | 0.0835 | 0.091 | 0.09 48 | 0 | 1 | 0.04 03 | predicted protein |
| contig_24410_ 6 | 0.035 | 0.0843 | 0.0916 | 0.09 56 | 0 | 1 | 0.04 07 | sodium/potassium/calcium exchanger 2-like |
| contig_16863_ 4 | 0.0353 | 0.085 | 0.0916 | 0.09 62 | 0 | 1 | 0.04 09 | D-3-phosphoglycerate dehydrogenase |
| contig_39943_ 5 | 0.0359 | 0.0861 | 0.0925 | 0.09 77 | 0 | 1 | 0.04 16 | mitochondrial alternative oxidase |
| contig_33687_ 4 | 0.0362 | 0.0877 | 0.0926 | 0.09 83 | 0 | 1 | 0.04 18 | predicted protein |
| contig_36067_ 5 | 0.0364 | 0.088 | 0.093 | 0.09 84 | 0 | 1 | 0.04 19 | silicon transporter |
| contig_36049_ 1 | 0.0364 | 0.0894 | 0.093 | 0.09 84 | 0 | 1 | 0.04 19 | predicted protein |
| contig_37059_ 1 | 0.0368 | 0.0901 | 0.0932 | 0.09 94 | 0 | 1 | 0.04 23 | Ura6p |
| contig_7582_3 | 0.0369 | 0.0907 | 0.0936 | 0.09 96 | 0 | 1 | 0.04 24 | vanadium-dependent haloperoxidase |
| contig_29365_ 1 | 0.0372 | 0.0908 | 0.0939 | 0.10 01 | 0 | 1 | 0.04 26 | predicted protein |
| contig_10866_ 1 | 0.0373 | 0.091 | 0.0939 | 0.10 03 | 0 | 1 | 0.04 27 | predicted protein |
| contig_34697_ 6 | 0.0377 | 0.0915 | 0.0973 | 0.10 12 | 0 | 1 | 0.04 31 | asparagine synthase |
| contig_7371_4 | 0.0385 | 0.0916 | 0.0984 | 0.10 3 | 0 | 1 | 0.04 38 | Lipocalin family protein |
| contig_29766_ 3 | 0.0386 | 0.092 | 0.0997 | 0.10 31 | 0 | 1 | 0.04 39 | GMP synthase (glutamine-hydrolyzing) domain-containing protein |
| contig_13819_ 5 | 0.0396 | 0.0926 | 0.0997 | 0.10 53 | 0 | 1 | 0.04 48 | 7-dehydrocholesterol reductase |
| contig_23765_ 6 | 0.0396 | 0.093 | 0.1022 | 0.10 53 | 0 | 1 | 0.04 48 | predicted protein |
| contig_25130_ 6 | 0.0396 | 0.093 | 0.1031 | 0.10 53 | 0 | 1 | 0.04 48 | methionine-tRNA ligase |
| contig_30367_ 5 | 0.04 | 0.0932 | 0.1032 | 0.10 6 | 0 | 1 | 0.04 51 | protein fucoxanthin chlorophyll a/c protein |

| | | | | | | | | |
|---|---|---|---|---|---|---|---|---|
| contig_25907_5 | 0.04 | 0.0936 | 0.1045 | 0.106 | 0 | 1 | 0.0451 | predicted protein |
| contig_41159_3 | 0.0401 | 0.0939 | 0.1071 | 0.106 | 0 | 1 | 0.0451 | predicted protein |
| contig_30585_6 | 0.0403 | 0.0939 | 0.1073 | 0.1063 | 0 | 1 | 0.0452 | ribosomal protein S2 |
| contig_21525_6 | 0.0403 | 0.0973 | 0.1074 | 0.1063 | 0 | 1 | 0.0452 | probable small nuclear ribonucleoprotein Sm D2 |
| contig_2955_5 | 0.0407 | 0.0985 | 0.1075 | 0.107 | 0 | 1 | 0.0455 | NADH dehydrogenase (ubiquinone) flavoprotein 2 |
| contig_27562_4 | 0.0408 | 0.0997 | 0.1169 | 0.1071 | 0 | 1 | 0.0456 | acyl-CoA dehydrogenase |
| contig_12161_5 | 0.041 | 0.1022 | 0.117 | 0.1074 | 0 | 1 | 0.0457 | predicted protein |
| contig_19474_5 | 0.0411 | 0.1031 | 0.1209 | 0.1074 | 0 | 1 | 0.0457 | aspartate aminotransferase |
| contig_34164_6 | 0.0412 | 0.1032 | 0.121 | 0.1076 | 0 | 1 | 0.0458 | multispanning membrane protein |
| contig_26152_4 | 0.0417 | 0.1045 | 0.1212 | 0.1087 | 0 | 1 | 0.0463 | inositol 5-phosphatase |
| contig_32749_4 | 0.0418 | 0.1071 | 0.1226 | 0.1087 | 0 | 1 | 0.0463 | leu phe-trna protein transferase |
| contig_6097_6 | 0.0418 | 0.1074 | 0.1227 | 0.1087 | 0 | 1 | 0.0463 | S-adenosylmethionine synthase 2 |
| contig_41167_2 | 0.0421 | 0.1079 | 0.1234 | 0.1092 | 0 | 1 | 0.0465 | oxygen-evolving enhancer protein 1 precursor |
| contig_8116_6 | 0.0422 | 0.1079 | 0.1263 | 0.1092 | 0 | 1 | 0.0465 | 40S ribosomal protein S11 |
| contig_38925_2 | 0.0426 | 0.117 | 0.127 | 0.1101 | 0 | 1 | 0.0469 | ---NA--- |
| contig_26568_4 | 0.0433 | 0.121 | 0.1276 | 0.1117 | 0 | 1 | 0.0475 | atpase family aaa domain-containing protein 3 |
| contig_34264_6 | 0.0435 | 0.121 | 0.129 | 0.112 | 0 | 1 | 0.0477 | probable small nuclear ribonucleoprotein E |
| contig_24205_6 | 0.0436 | 0.1212 | 0.1304 | 0.1122 | 0 | 1 | 0.0477 | protein fucoxanthin chlorophyll a/c protein |
| contig_19117_2 | 0.0437 | 0.1227 | 0.1306 | 0.1122 | 0 | 1 | 0.0477 | 60S ribosomal protein L21-A |
| contig_24983_1 | 0.0439 | 0.1234 | 0.1307 | 0.1126 | 0 | 1 | 0.0479 | GTP-binding nuclear protein Ran |
| contig_34938_2 | 0.044 | 0.127 | 0.1312 | 0.1126 | 0 | 1 | 0.0479 | predicted protein |
| contig_11078_3 | 0.0441 | 0.127 | 0.1314 | 0.1127 | 0 | 1 | 0.0479 | peroxiredoxin |
| contig_5526_6 | 0.0444 | 0.1276 | 0.1314 | 0.1133 | 0 | 1 | 0.0482 | pol protein |
| contig_13885_3 | 0.0446 | 0.1296 | 0.1349 | 0.1136 | 0 | 1 | 0.0484 | predicted protein |
| contig_25885_5 | 0.0451 | 0.1304 | 0.1363 | 0.1147 | 0 | 1 | 0.0488 | acetyl-coenzyme a synthetase |
| contig_12639_1 | 0.0458 | 0.1307 | 0.1366 | 0.1163 | 0 | 1 | 0.0495 | ADP-ribosylation factor-related protein 1 |
| contig_38660_3 | 0.046 | 0.131 | 0.141 | 0.1165 | 0 | 1 | 0.0496 | elongation factor 2 |

| | | | | | | | | |
|---|---|---|---|---|---|---|---|---|
| contig_23500_4 | 0.0466 | 0.1314 | 0.141 | 0.1178 | 0 | 1 | 0.0501 | proline-tRNA ligase |
| contig_24960_3 | 0.0467 | 0.1343 | 0.1417 | 0.118 | 0 | 1 | 0.0502 | predicted protein |
| contig_23912_3 | 0.0472 | 0.1349 | 0.1419 | 0.119 | 0 | 1 | 0.0506 | predicted protein |
| contig_9605_2 | 0.0476 | 0.1364 | 0.1426 | 0.1198 | 0 | 1 | 0.051 | arginine biosynthesis ArgJ mitochondrial |
| contig_23071_3 | 0.0481 | 0.1364 | 1 | 0.1208 | 0 | 1 | 0.0514 | D-3-phosphoglycerate dehydrogenase |
| contig_30517_5 | 0.0483 | 0.141 | 1 | 0.1212 | 0 | 1 | 0.0516 | predicted protein |
| contig_27521_6 | 0.0494 | 0.141 | 1 | 0.1237 | 0 | 1 | 0.0526 | predicted protein |
| contig_17246_6 | 0.0495 | 0.1417 | 1 | 0.1237 | 0 | 1 | 0.0526 | predicted protein |
| contig_32560_1 | 0.0497 | 0.1421 | 1 | 0.1239 | 0 | 1 | 0.0527 | predicted protein |
| contig_34763_6 | 0.0497 | 0.1428 | 1 | 0.1239 | 0 | 1 | 0.0527 | myosin light chain kinase |
| contig_38301_1 | 0.05 | 1 | 1 | 0.1245 | 0 | 1 | 0.053 | predicted protein |
| contig_28412_6 | 0.0509 | 1 | 1 | 0.1264 | 0 | 1 | 0.0538 | predicted protein |
| contig_9347_5 | 0.0509 | 1 | 1 | 0.1264 | 0 | 1 | 0.0538 | putative glycerol-3-phosphate transporter 1 |
| contig_21631_6 | 0.0517 | 1 | 1 | 0.128 | 0 | 1 | 0.0545 | predicted protein |
| contig_36399_5 | 0.0522 | 1 | 1 | 0.129 | 0 | 1 | 0.0549 | triosephosphate isomerase/glyceraldehyde-3-phosphate dehydrogenase precursor |
| contig_22684_3 | 0.0526 | 1 | 1 | 0.1299 | 0 | 1 | 0.0553 | predicted protein |
| contig_24628_4 | 0.0534 | 1 | 1 | 0.1317 | 0 | 1 | 0.056 | predicted protein |
| contig_27187_2 | 0.0535 | 1 | 1 | 0.1317 | 0 | 1 | 0.056 | CAAX prenyl protease 1 |
| contig_41162_4 | 0.0544 | 1 | 1 | 0.1336 | 0 | 1 | 0.0569 | 60S ribosomal protein L9-like |
| contig_28624_5 | 0.055 | 1 | 1 | 0.1348 | 0 | 1 | 0.0574 | eukaryotic peptide chain release factor subunit 1 |
| contig_19778_5 | 0.0553 | 1 | 1 | 0.135 | 0 | 1 | 0.0574 | voltage-gated chloride channel |
| contig_23099_5 | 0.0553 | 1 | 1 | 0.135 | 0 | 1 | 0.0574 | NAD(P)-dependent alcohol dehydrogenase |
| contig_36104_5 | 0.0553 | 1 | 1 | 0.135 | 0 | 1 | 0.0574 | 40S ribosomal protein S2 |
| contig_28948_1 | 0.0567 | 1 | 1 | 0.1382 | 0 | 1 | 0.0588 | geranylgeranyl reductase |
| contig_22479_1 | 0.0573 | 1 | 1 | 0.1394 | 0 | 1 | 0.0593 | predicted protein |
| contig_6179_6 | 0.0574 | 1 | 1 | 0.1394 | 1E-04 | 1 | 0.0593 | Long-chain-fatty-acid--CoA ligase putative |

| | | | | | | | | |
|---|---|---|---|---|---|---|---|---|
| contig_28302_4 | 0.0579 | 1 | 1 | 0.1404 | 1E-04 | 1 | 0.0598 | predicted protein |
| contig_34741_6 | 0.0585 | 1 | 1 | 0.1416 | 1E-04 | 1 | 0.0603 | 60S ribosomal protein |
| contig_38474_6 | 0.0587 | 1 | 1 | 0.1418 | 1E-04 | 1 | 0.0604 | phosphoserine transaminase |
| contig_8757_2 | 0.059 | 1 | 1 | 0.1424 | 1E-04 | 1 | 0.0606 | ABC transporter |
| contig_23946_2 | 0.0597 | 1 | 1 | 0.1438 | 2E-04 | 1 | 0.0612 | ATP-binding cassette subfamily F member 3 |
| contig_34246_6 | 0.0598 | 1 | 1 | 0.1438 | 2E-04 | 1 | 0.0612 | homoserine kinase |
| contig_14719_6 | 0.0599 | 1 | 1 | 0.1439 | 2E-04 | 1 | 0.0612 | thioredoxin-like |
| contig_15891_1 | 0.0606 | 1 | 1 | 0.1452 | 3E-04 | 1 | 0.0618 | predicted protein |
| contig_33594_1 | 0.0606 | 1 | 1 | 0.1452 | 4E-04 | 1 | 0.0618 | CULlin protein 1 |
| contig_22969_5 | 0.0609 | 1 | 1 | 0.1455 | 4E-04 | 1 | 0.0619 | molecular chaperone DnaJ |
| contig_28923_1 | 0.061 | 1 | 1 | 0.1455 | 5E-04 | 1 | 0.0619 | predicted protein |
| contig_35278_2 | 0.0617 | 1 | 1 | 0.1471 | 6E-04 | 1 | 0.0626 | light harvesting protein |
| contig_25552_1 | 0.062 | 1 | 1 | 0.1475 | 8E-04 | 1 | 0.0628 | predicted protein |
| contig_26661_4 | 0.0625 | 1 | 1 | 0.1486 | 9E-04 | 1 | 0.0632 | photosystem II oxygen evolving complex protein PsbP |
| contig_39585_3 | 0.0632 | 1 | 1 | 0.1499 | 0.0011 | 1 | 0.0638 | predicted protein |
| contig_8988_3 | 0.0633 | 1 | 1 | 0.15 | 0.0013 | 1 | 0.0638 | DUF1295 domain-containing protein |
| contig_27634_3 | 0.0639 | 1 | 1 | 0.151 | 0.0016 | 1 | 0.0643 | eukaryotic translation initiation factor 3 subunit E |
| contig_6151_5 | 0.064 | 1 | 1 | 0.1511 | 0.0019 | 1 | 0.0643 | CD209 antigen-like protein 2 isoform X1 |
| contig_29033_5 | 0.0645 | 1 | 1 | 0.1519 | 0.0022 | 1 | 0.0646 | EF-hand domain-containing protein |
| contig_15545_5 | 0.0645 | 1 | 1 | 0.1519 | 0.0026 | 1 | 0.0646 | Small nuclear ribonucleoprotein G |
| contig_22734_3 | 0.0649 | 1 | 1 | 0.1526 | 0.0031 | 1 | 0.0649 | predicted protein |
| contig_27953_1 | 0.0652 | 1 | 1 | 0.1531 | 0.0037 | 1 | 0.0651 | ribosomal protein L23 putative |
| contig_37721_1 | 0.0653 | 1 | 1 | 0.1531 | 0.0043 | 1 | 0.0651 | ---NA--- |
| contig_17375_3 | 0.0656 | 1 | 1 | 0.1532 | 0.005 | 1 | 0.0652 | cysteine synthase |
| contig_27477_6 | 0.0657 | 1 | 1 | 0.1532 | 0.0058 | 1 | 0.0652 | gdp-mannose 4 6-dehydratase |
| contig_15954_4 | 0.0657 | 1 | 1 | 0.1532 | 0.0068 | 1 | 0.0652 | elongation factor-like |
| contig_19427_5 | 0.0658 | 1 | 1 | 0.1533 | 0.0079 | 1 | 0.0652 | 60S ribosomal protein L18a |

| | | | | | | | | |
|---|---|---|---|---|---|---|---|---|
| contig_12505_5 | 0.0665 | 1 | 1 | 0.1546 | 0.0092 | 1 | 0.0658 | threonine ammonia-lyase |
| contig_38597_3 | 0.0675 | 1 | 1 | 0.1565 | 0.0106 | 1 | 0.0666 | ---NA--- |
| contig_18202_5 | 0.0676 | 1 | 1 | 0.1565 | 0.0122 | 1 | 0.0666 | protein fucoxanthin chlorophyll a/c protein |
| contig_19330_5 | 0.0677 | 1 | 1 | 0.1565 | 0.014 | 1 | 0.0666 | predicted protein |
| contig_9050_3 | 0.0678 | 1 | 1 | 0.1565 | 0.016 | 1 | 0.0666 | triose phosphate/phosphate translocator |
| contig_30315_5 | 0.0678 | 1 | 1 | 0.1565 | 0.0183 | 1 | 0.0666 | thioredoxin H4-1-like |
| contig_3793_6 | 0.069 | 1 | 1 | 0.1589 | 0.0208 | 1 | 0.0676 | ribosomal protein L20 (chloroplast) |
| contig_39895_1 | 0.0698 | 1 | 1 | 0.1604 | 0.0236 | 1 | 0.0683 | protein fucoxanthin chlorophyll a/c protein |
| contig_34416_5 | 0.0698 | 1 | 1 | 0.1604 | 0.0267 | 1 | 0.0683 | predicted protein |
| contig_27112_5 | 0.0708 | 1 | 1 | 0.1624 | 0.0302 | 1 | 0.0691 | calreticulin |
| contig_27033_6 | 0.0711 | 1 | 1 | 0.1628 | 0.034 | 1 | 0.0693 | glucokinase |
| contig_14621_1 | 0.0721 | 1 | 1 | 0.1649 | 0.0382 | 1 | 0.0702 | predicted protein |
| contig_28238_5 | 0.0726 | 1 | 1 | 0.1655 | 0.0428 | 1 | 0.0704 | ubiquinol-cytochrome c reductase iron-sulfur subunit |
| contig_35157_2 | 0.0726 | 1 | 1 | 0.1655 | 0.0478 | 1 | 0.0704 | predicted protein |
| contig_23694_6 | 0.073 | 1 | 1 | 0.1662 | 0.0532 | 1 | 0.0707 | cell division cycle protein 123 homolog |
| contig_33639_6 | 0.0731 | 1 | 1 | 0.1662 | 0.0592 | 1 | 0.0707 | ---NA--- |
| contig_22496_6 | 0.0732 | 1 | 1 | 0.1662 | 0.0656 | 1 | 0.0707 | predicted protein |
| contig_29967_5 | 0.0736 | 1 | 1 | 0.1668 | 0.0726 | 1 | 0.071 | protein dispatched homolog 1 isoform X2 |
| contig_24783_5 | 0.0737 | 1 | 1 | 0.1668 | 0.0801 | 1 | 0.071 | mitochondrial protein translocase MPT family |
| contig_32841_4 | 0.0749 | 1 | 1 | 0.1692 | 0.0883 | 1 | 0.072 | predicted protein |
| contig_26948_5 | 0.0752 | 1 | 1 | 0.1698 | 0.0969 | 1 | 0.0722 | predicted protein |
| contig_29330_2 | 0.0753 | 1 | 1 | 0.1698 | 0.1062 | 1 | 0.0722 | pilus assembly protein |
| contig_22578_5 | 0.0761 | 1 | 1 | 0.1712 | 0.1161 | 1 | 0.0728 | Urea-proton symporter DUR3 |
| contig_10388_3 | 0.0772 | 1 | 1 | 0.1735 | 0.1266 | 1 | 0.0738 | light-harvesting protein |
| contig_25123_5 | 0.0774 | 1 | 1 | 0.1737 | 0.1377 | 1 | 0.0739 | ubiquitin-activating enzyme E1 protein 3 |
| contig_30020_5 | 0.0777 | 1 | 1 | 0.1741 | 0.1494 | 1 | 0.0741 | chaperonin GroEL |
| contig_27824_2 | 0.0783 | 1 | 1 | 0.175 | 0.1617 | 1 | 0.0745 | protein phosphatase 2C putative |

| | | | | | | | | |
|---|---|---|---|---|---|---|---|---|
| contig_25835_2 | 0.0785 | 1 | 1 | 0.1752 | 0.1747 | 1 | 0.0745 | histidinol dehydrogenase |
| contig_39705_1 | 0.0788 | 1 | 1 | 0.1757 | 0.1883 | 1 | 0.0747 | inorganic pyrophosphatase |
| contig_5781_6 | 0.0795 | 1 | 1 | 0.1769 | 0.2026 | 1 | 0.0753 | acetyl-coenzyme a synthetase |
| contig_33536_4 | 0.0797 | 1 | 1 | 0.1771 | 0.2174 | 1 | 0.0754 | sec1 family transporter sly1 |
| contig_34219_5 | 0.0804 | 1 | 1 | 0.1784 | 0.2328 | 1 | 0.0759 | Two component regulator three Y domain protein |
| contig_37972_6 | 0.0807 | 1 | 1 | 0.179 | 0.2488 | 1 | 0.0762 | predicted protein |
| contig_32692_5 | 0.0809 | 1 | 1 | 0.1792 | 0.2653 | 1 | 0.0762 | predicted protein |
| contig_9464_1 | 0.0812 | 1 | 1 | 0.1794 | 0.2824 | 1 | 0.0763 | cytochrome c heme attachment protein (chloroplast) |
| contig_23660_5 | 0.0814 | 1 | 1 | 0.1797 | 0.3 | 1 | 0.0765 | predicted protein |
| contig_24297_3 | 0.0815 | 1 | 1 | 0.1797 | 0.3181 | 1 | 0.0765 | elongation factor Tu |
| contig_35081_4 | 0.0821 | 1 | 1 | 0.1805 | 0.3367 | 1 | 0.0768 | ubiquinol oxidase 4 chloroplastic/chromoplastic |
| contig_20020_4 | 0.0821 | 1 | 1 | 0.1805 | 0.3556 | 1 | 0.0768 | predicted protein |
| contig_12843_3 | 0.0824 | 1 | 1 | 0.1809 | 0.375 | 1 | 0.077 | proteasome subunit beta type-3 |
| contig_33755_6 | 0.0827 | 1 | 1 | 0.1813 | 0.3946 | 1 | 0.0771 | predicted protein |
| contig_25589_3 | 0.0829 | 1 | 1 | 0.1814 | 0.4145 | 1 | 0.0772 | small subunit ribosomal protein S3Ae |
| contig_25881_5 | 0.0831 | 1 | 1 | 0.1815 | 0.4347 | 1 | 0.0772 | predicted protein |
| contig_28787_6 | 0.0833 | 1 | 1 | 0.1818 | 0.455 | 1 | 0.0774 | tryptophan synthase |
| contig_10727_3 | 0.0835 | 1 | 1 | 0.182 | 0.4755 | 1 | 0.0775 | eukaryotic translation initiation factor 3 subunit G |
| contig_36385_4 | 0.0841 | 1 | 1 | 0.1829 | 0.4961 | 1 | 0.0778 | ---NA--- |
| contig_7030_6 | 0.0843 | 1 | 1 | 0.1832 | 0.5166 | 1 | 0.078 | ---NA--- |
| contig_28073_5 | 0.0845 | 1 | 1 | 0.1833 | 0.5371 | 1 | 0.078 | NAD-dependent epimerase/dehydratase family protein |
| contig_26111_3 | 0.085 | 1 | 1 | 0.1841 | 0.5575 | 1 | 0.0783 | predicted protein |
| contig_17354_1 | 0.0854 | 1 | 1 | 0.1848 | 0.5778 | 1 | 0.0786 | T9SS type A sorting domain-containing protein |
| contig_29429_4 | 0.0858 | 1 | 1 | 0.1854 | 0.5978 | 1 | 0.0789 | predicted protein |
| contig_40101_4 | 0.0861 | 1 | 1 | 0.1858 | 0.6175 | 1 | 0.079 | 40S ribosomal protein S13 |
| contig_16671_2 | 0.0868 | 1 | 1 | 0.187 | 0.637 | 1 | 0.0796 | ribosomal protein L21 (chloroplast) |

| | | | | | | | | |
|---|---|---|---|---|---|---|---|---|
| contig_19309_1 | 0.0874 | 1 | 1 | 0.1881 | 0.656 | 1 | 0.08 | 60S ribosomal protein L9-like |
| contig_25207_1 | 0.0877 | 1 | 1 | 0.1881 | 0.6746 | 1 | 0.08 | predicted protein |
| contig_38859_2 | 0.0877 | 1 | 1 | 0.1881 | 0.6928 | 1 | 0.08 | hypersensitive-induced response protein 1-like |
| contig_27280_6 | 0.088 | 1 | 1 | 0.1885 | 0.7106 | 1 | 0.0802 | dihydrolipoamide dehydrogenase |
| contig_24750_1 | 0.0889 | 1 | 1 | 0.1901 | 0.7279 | 1 | 0.0809 | AP-1 complex subunit beta-1 |
| contig_27547_5 | 0.0892 | 1 | 1 | 0.1906 | 0.7445 | 1 | 0.0811 | ribulose-phosphate 3-epimerase |
| contig_14481_1 | 0.0894 | 1 | 1 | 0.1908 | 0.7606 | 1 | 0.0812 | CDP-diacylglycerol--glycerol-3-phosphate 3-phosphatidyltransferase |
| contig_23603_4 | 0.0899 | 1 | 1 | 0.1914 | 0.7762 | 1 | 0.0815 | serine protease |
| contig_24544_3 | 0.0901 | 1 | 1 | 0.1916 | 0.7911 | 1 | 0.0815 | predicted protein |
| contig_14030_3 | 0.0904 | 1 | 1 | 0.192 | 0.8055 | 1 | 0.0817 | S-adenosylmethionine carrier 1 chloroplastic/mitochondrial |
| contig_24913_6 | 0.0907 | 1 | 1 | 0.1921 | 0.8192 | 1 | 0.0818 | glutamate synthase subunit alpha |
| contig_27487_4 | 0.0908 | 1 | 1 | 0.1921 | 0.8324 | 1 | 0.0818 | ---NA--- |
| contig_41187_5 | 0.0909 | 1 | 1 | 0.1921 | 0.8449 | 1 | 0.0818 | proteasome subunit beta type 4 |
| contig_5519_5 | 0.091 | 1 | 1 | 0.1921 | 0.8568 | 1 | 0.0818 | protein tesmin/TSO1-like CXC 5 |
| contig_5548_1 | 0.0915 | 1 | 1 | 0.1928 | 0.8681 | 1 | 0.082 | geranylgeranyl pyrophosphate synthase |
| contig_25733_5 | 0.0916 | 1 | 1 | 0.1928 | 0.8787 | 1 | 0.082 | predicted protein |
| contig_21520_1 | 0.0916 | 1 | 1 | 0.1928 | 0.8887 | 1 | 0.082 | phosphoglycerate mutase 1 family |
| contig_22528_2 | 0.092 | 1 | 1 | 0.1933 | 0.8982 | 1 | 0.0823 | predicted protein |
| contig_5617_6 | 0.0925 | 1 | 1 | 0.194 | 0.907 | 1 | 0.0825 | sarcoplasmic calcium-binding protein |
| contig_13609_2 | 0.0926 | 1 | 1 | 0.194 | 0.9153 | 1 | 0.0825 | U5 small nuclear ribonucleoprotein 200 kDa helicase |
| contig_28929_5 | 0.0929 | 1 | 1 | 0.194 | 0.923 | 1 | 0.0826 | 1-deoxy-D-xylulose 5-phosphate reductoisomerase |
| contig_23921_2 | 0.093 | 1 | 1 | 0.194 | 0.9302 | 1 | 0.0826 | predicted protein |
| contig_24766_6 | 0.093 | 1 | 1 | 0.194 | 0.9368 | 1 | 0.0826 | nitrate reductase |
| contig_40524_3 | 0.0932 | 1 | 1 | 0.1941 | 0.943 | 1 | 0.0826 | light harvesting protein |
| contig_5607_2 | 0.0935 | 1 | 1 | 0.1944 | 0.9487 | 1 | 0.0827 | protease SohB |
| contig_38999_3 | 0.0936 | 1 | 1 | 0.1944 | 0.9539 | 1 | 0.0827 | predicted protein |

| | | | | | | | | |
|---|---|---|---|---|---|---|---|---|
| contig_25649_3 | 0.0939 | 1 | 1 | 0.1946 | 0.9587 | 1 | 0.0828 | tRNA pseudouridine(55) synthase |
| contig_29992_5 | 0.0939 | 1 | 1 | 0.1946 | 0.9631 | 1 | 0.0828 | ---NA--- |
| contig_30596_2 | 0.0949 | 1 | 1 | 0.1965 | 0.9671 | 1 | 0.0836 | predicted protein |
| contig_27659_6 | 0.0965 | 1 | 1 | 0.1993 | 0.9707 | 1 | 0.0848 | predicted protein |
| contig_33557_6 | 0.0966 | 1 | 1 | 0.1993 | 0.974 | 1 | 0.0848 | phosphoglycerate kinase |
| contig_28744_4 | 0.0968 | 1 | 1 | 0.1994 | 0.9769 | 1 | 0.0849 | ascorbate peroxidase |
| contig_27144_6 | 0.0973 | 1 | 1 | 0.2001 | 0.9796 | 1 | 0.0851 | predicted protein |
| contig_13426_4 | 0.0973 | 1 | 1 | 0.2001 | 0.982 | 1 | 0.0851 | long-chain fatty acid--CoA ligase |
| contig_12588_5 | 0.0982 | 1 | 1 | 0.2015 | 0.9842 | 1 | 0.0857 | predicted protein |
| contig_25252_3 | 0.0984 | 1 | 1 | 0.2016 | 0.9861 | 1 | 0.0858 | Annexin protein |
| contig_25953_4 | 0.0985 | 1 | 1 | 0.2016 | 0.9878 | 1 | 0.0858 | predicted protein |
| contig_37814_2 | 0.0991 | 1 | 1 | 0.2026 | 0.9893 | 1 | 0.0862 | ribosomal protein S3 (chloroplast) |
| contig_23988_3 | 0.0997 | 1 | 1 | 0.2033 | 0.9907 | 1 | 0.0865 | predicted protein |
| contig_27236_4 | 0.0997 | 1 | 1 | 0.2033 | 0.9919 | 1 | 0.0865 | translation initiation factor aif-2 |
| contig_19585_5 | 0.1002 | 1 | 1 | 0.2039 | 0.993 | 1 | 0.0868 | ---NA--- |
| contig_33399_2 | 0.1011 | 1 | 1 | 0.2055 | 0.9939 | 1 | 0.0875 | predicted protein |
| contig_35860_4 | 0.1018 | 1 | 1 | 0.2067 | 0.9947 | 1 | 0.0879 | mitochondrial manganese superoxide dismutase |
| contig_19000_1 | 0.1022 | 1 | 1 | 0.2071 | 0.9955 | 1 | 0.0881 | predicted protein |
| contig_22590_5 | 0.1022 | 1 | 1 | 0.2071 | 0.9961 | 1 | 0.0881 | glutathione reductase |
| contig_25763_5 | 0.103 | 1 | 1 | 0.2081 | 0.9967 | 1 | 0.0886 | Dihydroflavonol 4-reductase |
| contig_30187_4 | 0.1031 | 1 | 1 | 0.2081 | 0.9971 | 1 | 0.0886 | adenylosuccinate lyase |
| contig_1081_1 | 0.1032 | 1 | 1 | 0.2081 | 0.9976 | 1 | 0.0886 | hypothetical protein CBC21_09400 |
| contig_7021_2 | 0.1036 | 1 | 1 | 0.2088 | 0.9979 | 1 | 0.0889 | protein phosphatase 2C putative |
| contig_40922_3 | 0.1042 | 1 | 1 | 0.2096 | 0.9982 | 1 | 0.0892 | V-type H-ATPase subunit C |
| contig_6442_4 | 0.1045 | 1 | 1 | 0.21 | 0.9985 | 1 | 0.0894 | putative ATP-dependent RNA helicase DHX30 isoform X3 |
| contig_5826_3 | 0.106 | 1 | 1 | 0.2126 | 0.9987 | 1 | 0.0905 | 60S ribosomal protein L2 |
| contig_22324_4 | 0.1067 | 1 | 1 | 0.2137 | 0.9989 | 1 | 0.091 | ---NA--- |

| | | | | | | | | |
|---|---|---|---|---|---|---|---|---|
| contig_34549_1 | 0.1069 | 1 | 1 | 0.2137 | 0.9991 | 1 | 0.091 | pirin family protein |
| contig_30830_5 | 0.107 | 1 | 1 | 0.2137 | 0.9992 | 1 | 0.091 | PGR5-like protein 1A chloroplastic |
| contig_33797_1 | 0.1071 | 1 | 1 | 0.2137 | 0.9994 | 1 | 0.091 | glycine-tRNA ligase |
| contig_17918_5 | 0.1073 | 1 | 1 | 0.2137 | 0.9995 | 1 | 0.091 | glutamate-1-semialdehyde 2 1-aminomutase |
| contig_33304_4 | 0.1074 | 1 | 1 | 0.2137 | 0.9996 | 1 | 0.091 | predicted protein |
| contig_15461_5 | 0.1075 | 1 | 1 | 0.2138 | 0.9996 | 1 | 0.091 | predicted protein |
| contig_40270_3 | 0.1079 | 1 | 1 | 0.2142 | 0.9997 | 1 | 0.0912 | predicted protein |
| contig_24322_5 | 0.1088 | 1 | 1 | 0.2157 | 0.9997 | 1 | 0.0918 | ARM repeat superfamily protein |
| contig_22513_4 | 0.1095 | 1 | 1 | 0.2168 | 0.9998 | 1 | 0.0923 | predicted protein |
| contig_11299_6 | 0.1104 | 1 | 1 | 0.2183 | 0.9998 | 1 | 0.0929 | predicted protein |
| contig_3444_2 | 0.111 | 1 | 1 | 0.2193 | 0.9999 | 1 | 0.0933 | protoporphyrin IX methyltransferase |
| contig_24884_4 | 0.113 | 1 | 1 | 0.2229 | 0.9999 | 1 | 0.0949 | predicted protein |
| contig_11923_6 | 0.1139 | 1 | 1 | 0.2243 | 0.9999 | 1 | 0.0955 | aspartate aminotransferase |
| contig_10198_5 | 0.1148 | 1 | 1 | 0.2259 | 0.9999 | 1 | 0.0961 | predicted protein |
| contig_39819_3 | 0.1155 | 1 | 1 | 0.2269 | 0.9999 | 1 | 0.0966 | predicted protein |
| contig_24221_5 | 0.1156 | 1 | 1 | 0.2269 | 0.9999 | 1 | 0.0966 | ATP-binding cassette superfamily |
| contig_32942_6 | 0.116 | 1 | 1 | 0.2274 | 1 | 1 | 0.0968 | predicted protein |
| contig_38368_6 | 0.1166 | 1 | 1 | 0.2279 | 1 | 1 | 0.097 | glutathione S-transferase |
| contig_41171_6 | 0.1167 | 1 | 1 | 0.2279 | 1 | 1 | 0.097 | ADP/ATP transporter on adenylate translocase putative |
| contig_24282_5 | 0.1168 | 1 | 1 | 0.2279 | 1 | 1 | 0.097 | predicted protein |
| contig_30780_6 | 0.1169 | 1 | 1 | 0.2279 | 1 | 1 | 0.097 | possible transporter |
| contig_21789_5 | 0.117 | 1 | 1 | 0.2279 | 1 | 1 | 0.097 | Mitochondrial Carrier (MC) Family |
| contig_15866_5 | 0.1176 | 1 | 1 | 0.2286 | 1 | 1 | 0.0973 | hypothetical protein THAOC_02924 |
| contig_6495_6 | 0.1192 | 1 | 1 | 0.2315 | 1 | 1 | 0.0985 | predicted protein |
| contig_22939_2 | 0.12 | 1 | 1 | 0.2327 | 1 | 1 | 0.099 | DNA damage-binding protein 1 |
| contig_9471_3 | 0.1201 | 1 | 1 | 0.2327 | 1 | 1 | 0.099 | photosystem II CP43 chlorophyll apoprotein (chloroplast) |
| contig_8346_5 | 0.1206 | 1 | 1 | 0.2333 | 1 | 1 | 0.0993 | putative plastid division protein (chloroplast) |

| | | | | | | | | |
|---|---|---|---|---|---|---|---|---|
| contig_22768_6 | 0.1209 | 1 | 1 | 0.2333 | 1 | 1 | 0.0993 | putative LOV domain-containing protein |
| contig_22808_5 | 0.121 | 1 | 1 | 0.2333 | 1 | 1 | 0.0993 | predicted protein |
| contig_9719_3 | 0.121 | 1 | 1 | 0.2333 | 1 | 1 | 0.0993 | Co2+/Mg2+ efflux protein ApaG |
| contig_29868_5 | 0.1212 | 1 | 1 | 0.2333 | 1 | 1 | 0.0993 | adp-ribosylation factor 1 |
| contig_25969_5 | 0.1219 | 1 | 1 | 0.2345 | 1 | 1 | 0.0998 | Tic20-like protein |
| contig_16462_2 | 0.1226 | 1 | 1 | 0.2351 | 1 | 1 | 0.1 | chorismate synthase |
| contig_9464_6 | 0.1226 | 1 | 1 | 0.2351 | 1 | 1 | 0.1 | cytochrome c heme attachment protein (chloroplast) |
| contig_28775_6 | 0.1227 | 1 | 1 | 0.2351 | 1 | 1 | 0.1 | cullin-associated NEDD8-dissociated protein 1 |
| contig_27603_4 | 0.1234 | 1 | 1 | 0.2358 | 1 | 1 | 0.1003 | predicted protein |
| contig_28186_6 | 0.1234 | 1 | 1 | 0.2358 | 1 | 1 | 0.1003 | predicted protein |
| contig_26865_4 | 0.1244 | 1 | 1 | 0.2374 | 1 | 1 | 0.101 | ---NA--- |
| contig_21965_5 | 0.1246 | 1 | 1 | 0.2374 | 1 | 1 | 0.101 | predicted protein |
| contig_30693_4 | 0.1253 | 1 | 1 | 0.2385 | 1 | 1 | 0.1015 | ubiquitin-conjugating enzyme E2-16 kDa |
| contig_8226_3 | 0.1261 | 1 | 1 | 0.2398 | 1 | 1 | 0.1021 | NAD(P) transhydrogenase mitochondrial |
| contig_16707_4 | 0.1263 | 1 | 1 | 0.2399 | 1 | 1 | 0.1021 | nitrate reductase |
| contig_15734_6 | 0.127 | 1 | 1 | 0.2406 | 1 | 1 | 0.1024 | ALA dehydratase |
| contig_15258_2 | 0.127 | 1 | 1 | 0.2406 | 1 | 1 | 0.1024 | predicted protein |
| contig_10572_1 | 0.1275 | 1 | 1 | 0.2411 | 1 | 1 | 0.1026 | NADH dehydrogenase [ubiquinone] flavoprotein 1 mitochondrial |
| contig_25119_2 | 0.1276 | 1 | 1 | 0.2412 | 1 | 1 | 0.1026 | 3-phosphoshikimate 1-carboxyvinyltransferase |
| contig_15393_2 | 0.128 | 1 | 1 | 0.2416 | 1 | 1 | 0.1028 | predicted protein |
| contig_25394_4 | 0.1287 | 1 | 1 | 0.2426 | 1 | 1 | 0.1032 | prohibitin putative |
| contig_17183_2 | 0.129 | 1 | 1 | 0.2428 | 1 | 1 | 0.1033 | 40S ribosomal protein S17 |
| contig_33583_4 | 0.1296 | 1 | 1 | 0.2437 | 1 | 1 | 0.1037 | hypersensitive-induced response protein 1-like |
| contig_28714_6 | 0.1303 | 1 | 1 | 0.2445 | 1 | 1 | 0.1041 | predicted protein |
| contig_13636_6 | 0.1304 | 1 | 1 | 0.2445 | 1 | 1 | 0.1041 | acyl-CoA dehydrogenase-like protein |
| contig_30668_1 | 0.1306 | 1 | 1 | 0.2446 | 1 | 1 | 0.1041 | predicted protein |
| contig_19673_5 | 0.1307 | 1 | 1 | 0.2446 | 1 | 1 | 0.1041 | predicted protein |

| | | | | | | | | |
|---|---|---|---|---|---|---|---|---|
| contig_6511_2 | 0.131 | 1 | 1 | 0.24 47 | 1 | 1 | 0.10 41 | glutamyl-tRNA synthetase |
| contig_12625_ 2 | 0.1312 | 1 | 1 | 0.24 47 | 1 | 1 | 0.10 41 | 26S proteasome non-ATPase regulatory subunit 7 |
| contig_25208_ 3 | 0.1314 | 1 | 1 | 0.24 47 | 1 | 1 | 0.10 41 | predicted protein |
| contig_6054_4 | 0.1314 | 1 | 1 | 0.24 47 | 1 | 1 | 0.10 41 | ---NA--- |
| contig_5425_1 | 0.1325 | 1 | 1 | 0.24 64 | 1 | 1 | 0.10 49 | sulfate adenylyltransferase |
| contig_32739_ 1 | 0.133 | 1 | 1 | 0.24 7 | 1 | 1 | 0.10 51 | fibronectin type III domain-containing protein |
| contig_29806_ 5 | 0.1341 | 1 | 1 | 0.24 88 | 1 | 1 | 0.10 59 | predicted protein |
| contig_17096_ 5 | 0.1343 | 1 | 1 | 0.24 88 | 1 | 1 | 0.10 59 | predicted protein |
| contig_11292_ 5 | 0.1348 | 1 | 1 | 0.24 93 | 1 | 1 | 0.10 61 | heat shock protein belonging to the HSP70 family |
| contig_14919_ 6 | 0.1349 | 1 | 1 | 0.24 93 | 1 | 1 | 0.10 61 | predicted protein |
| contig_14990_ 5 | 0.1358 | 1 | 1 | 0.25 07 | 1 | 1 | 0.10 67 | heme oxygenase 1 chloroplastic-like |
| contig_20218_ 5 | 0.1361 | 1 | 1 | 0.25 09 | 1 | 1 | 0.10 67 | 40S ribosomal protein S26 |
| contig_18884_ 5 | 0.1363 | 1 | 1 | 0.25 09 | 1 | 1 | 0.10 68 | predicted protein |
| contig_39090_ 5 | 0.1364 | 1 | 1 | 0.25 09 | 1 | 1 | 0.10 68 | Glycerol-3-phosphate dehydrogenase |
| contig_9968_3 | 0.1366 | 1 | 1 | 0.25 1 | 1 | 1 | 0.10 68 | predicted protein |
| contig_16777_ 4 | 0.1382 | 1 | 1 | 0.25 36 | 1 | 1 | 0.10 79 | H - or Na -translocating F-type V-type and A-type ATPase (F-ATPase) Superfamily |
| contig_3138_3 | 0.1387 | 1 | 1 | 0.25 41 | 1 | 1 | 0.10 81 | predicted protein |
| contig_25820_ 1 | 0.14 | 1 | 1 | 0.25 6 | 1 | 1 | 0.10 89 | predicted protein |
| contig_23722_ 3 | 0.14 | 1 | 1 | 0.25 6 | 1 | 1 | 0.10 89 | light harvesting protein |
| contig_6069_1 | 0.1402 | 1 | 1 | 0.25 6 | 1 | 1 | 0.10 89 | GLNA glutamine synthase |
| contig_39464_ 1 | 0.1407 | 1 | 1 | 0.25 66 | 1 | 1 | 0.10 92 | histone deacetylase 1 |
| contig_27423_ 4 | 0.141 | 1 | 1 | 0.25 66 | 1 | 1 | 0.10 92 | predicted protein |
| contig_18936_ 6 | 0.141 | 1 | 1 | 0.25 66 | 1 | 1 | 0.10 92 | 40S ribosomal protein S3a |
| contig_37899_ 1 | 0.1414 | 1 | 1 | 0.25 7 | 1 | 1 | 0.10 94 | ribosomal protein L2 (chloroplast) |
| contig_16347_ 5 | 0.1417 | 1 | 1 | 0.25 72 | 1 | 1 | 0.10 94 | predicted protein |
| contig_25211_ 4 | 0.1419 | 1 | 1 | 0.25 72 | 1 | 1 | 0.10 95 | silicon transporter |

| | | | | | | | | |
|---|---|---|---|---|---|---|---|---|
| contig_5591_4 | 0.1421 | 1 | 1 | 0.2573 | 1 | 1 | 0.1095 | predicted protein |
| contig_23311_6 | 0.1426 | 1 | 1 | 0.2579 | 1 | 1 | 0.1097 | predicted protein |
| contig_6981_5 | 0.1428 | 1 | 1 | 0.258 | 1 | 1 | 0.1098 | predicted protein |
| contig_24874_1 | 0.1438 | 1 | 1 | 0.2594 | 1 | 1 | 0.1104 | predicted protein |
| contig_23827_5 | 0.1461 | 1 | 1 | 0.2633 | 1 | 1 | 0.112 | predicted protein |
| contig_25962_4 | 0.1472 | 1 | 1 | 0.2649 | 1 | 1 | 0.1127 | predicted protein |
| contig_11877_6 | 0.1483 | 1 | 1 | 0.2666 | 1 | 1 | 0.1134 | 30S ribosomal protein S5 |
| contig_30582_1 | 0.1548 | 1 | 1 | 0.278 | 1 | 1 | 0.1183 | silicon transporter |
| contig_25782_5 | 0.1555 | 1 | 1 | 0.2789 | 1 | 1 | 0.1187 | predicted protein |
| contig_6255_2 | 0.1559 | 1 | 1 | 0.2792 | 1 | 1 | 0.1188 | Nuclear transport factor 2 |
| contig_28743_4 | 0.1561 | 1 | 1 | 0.2792 | 1 | 1 | 0.1188 | chlorophyll a/b-binding protein |
| contig_16919_4 | 0.1562 | 1 | 1 | 0.2792 | 1 | 1 | 0.1188 | predicted protein |
| contig_37712_6 | 0.1571 | 1 | 1 | 0.2805 | 1 | 1 | 0.1194 | predicted protein |
| contig_29278_3 | 0.1589 | 1 | 1 | 0.2834 | 1 | 1 | 0.1206 | predicted protein |
| contig_22773_6 | 0.1603 | 1 | 1 | 0.2852 | 1 | 1 | 0.1214 | phosphoglycerate kinase |
| contig_22418_5 | 0.1603 | 1 | 1 | 0.2852 | 1 | 1 | 0.1214 | protein fucoxanthin chlorophyll a/c protein |
| contig_33184_2 | 0.161 | 1 | 1 | 0.2861 | 1 | 1 | 0.1218 | predicted protein |
| contig_28323_4 | 0.1615 | 1 | 1 | 0.2865 | 1 | 1 | 0.1219 | leucyl aminopeptidase |
| contig_28224_2 | 0.1616 | 1 | 1 | 0.2865 | 1 | 1 | 0.1219 | alpha tubulin |
| contig_25921_4 | 0.1619 | 1 | 1 | 0.2867 | 1 | 1 | 0.122 | predicted protein |
| contig_19038_3 | 0.1635 | 1 | 1 | 0.2892 | 1 | 1 | 0.123 | peptidylprolyl isomerase |
| contig_27523_3 | 0.1637 | 1 | 1 | 0.2892 | 1 | 1 | 0.123 | plastid protoporphyrinogen oxidase |
| contig_34911_5 | 0.1639 | 1 | 1 | 0.2892 | 1 | 1 | 0.123 | predicted protein |
| contig_22327_4 | 0.1642 | 1 | 1 | 0.2894 | 1 | 1 | 0.1232 | gamma-aminobutyric-acid receptor subunit beta |
| contig_23942_6 | 0.1645 | 1 | 1 | 0.2896 | 1 | 1 | 0.1232 | sister-chromatid cohesion protein 3 |
| contig_10451_3 | 0.1648 | 1 | 1 | 0.2899 | 1 | 1 | 0.1234 | predicted protein |
| contig_7935_3 | 0.1658 | 1 | 1 | 0.2912 | 1 | 1 | 0.1239 | predicted protein |

| | | | | | | | | |
|---|---|---|---|---|---|---|---|---|
| contig_6933_4 | 0.1662 | 1 | 1 | 0.2917 | 1 | 1 | 0.1241 | methylmalonate-semialdehyde dehydrogenase (acylating) |
| contig_23093_4 | 0.1674 | 1 | 1 | 0.2934 | 1 | 1 | 0.1248 | predicted protein |
| contig_1045_6 | 0.1676 | 1 | 1 | 0.2934 | 1 | 1 | 0.1248 | iron-sulfur cluster formation ABC transporter (chloroplast) |
| contig_28913_5 | 0.168 | 1 | 1 | 0.2938 | 1 | 1 | 0.125 | predicted protein |
| contig_6766_1 | 0.1684 | 1 | 1 | 0.2941 | 1 | 1 | 0.1251 | N-carbamoylputrescine amidase |
| contig_9535_1 | 0.1708 | 1 | 1 | 0.298 | 1 | 1 | 0.1268 | transitional endoplasmic reticulum ATPase |
| contig_26693_5 | 0.1713 | 1 | 1 | 0.2985 | 1 | 1 | 0.127 | UMP-CMP kinase |
| contig_30561_2 | 0.1728 | 1 | 1 | 0.3008 | 1 | 1 | 0.128 | cytochrome b6-f complex iron-sulfur subunit |
| contig_8482_6 | 0.1736 | 1 | 1 | 0.3018 | 1 | 1 | 0.1284 | UV excision repair protein RAD23 |
| contig_5001_1 | 0.1747 | 1 | 1 | 0.3034 | 1 | 1 | 0.1291 | membrane guanylyl cyclase |
| contig_27420_4 | 0.175 | 1 | 1 | 0.3035 | 1 | 1 | 0.1291 | polyhydroxybutyrate depolymerase |
| contig_21850_6 | 0.1753 | 1 | 1 | 0.3038 | 1 | 1 | 0.1293 | 40S ribosomal protein S14 |
| contig_26293_4 | 0.1781 | 1 | 1 | 0.3082 | 1 | 1 | 0.1311 | predicted protein |
| contig_10169_3 | 0.1784 | 1 | 1 | 0.3082 | 1 | 1 | 0.1312 | NADH-cytochrome b5 reductase 3-like |
| contig_34331_5 | 0.1785 | 1 | 1 | 0.3082 | 1 | 1 | 0.1312 | Light-harvesting complex I LH38 protein |
| contig_28392_3 | 0.179 | 1 | 1 | 0.3085 | 1 | 1 | 0.1313 | predicted protein |
| contig_23006_1 | 0.1791 | 1 | 1 | 0.3085 | 1 | 1 | 0.1313 | predicted protein |
| contig_36134_4 | 0.1795 | 1 | 1 | 0.309 | 1 | 1 | 0.1315 | fucoxanthin chlorophyll a/c protein |
| contig_36089_2 | 0.1814 | 1 | 1 | 0.3118 | 1 | 1 | 0.1327 | archaeal ribosomal protein S17P |
| contig_12446_4 | 0.1839 | 1 | 1 | 0.3158 | 1 | 1 | 0.1344 | UPF0187 protein |
| contig_28046_4 | 0.1847 | 1 | 1 | 0.3168 | 1 | 1 | 0.1348 | light harvesting protein |
| contig_33470_2 | 0.1863 | 1 | 1 | 0.3192 | 1 | 1 | 0.1358 | farnesyl pyrophosphate synthase |
| contig_30690_3 | 0.1867 | 1 | 1 | 0.3196 | 1 | 1 | 0.136 | phosphoglycerate mutase |
| contig_22932_5 | 0.1876 | 1 | 1 | 0.3206 | 1 | 1 | 0.1364 | predicted protein |
| contig_28171_1 | 0.1882 | 1 | 1 | 0.3214 | 1 | 1 | 0.1368 | predicted protein |
| contig_25214_5 | 0.1887 | 1 | 1 | 0.3217 | 1 | 1 | 0.1369 | predicted protein |
| contig_3511_5 | 0.1889 | 1 | 1 | 0.3217 | 1 | 1 | 0.1369 | Ycf89 (chloroplast) |

| | | | | | | | | |
|---|---|---|---|---|---|---|---|---|
| contig_16732_6 | 0.1905 | 1 | 1 | 0.3242 | 1 | 1 | 0.1379 | membrane spanning protein of the nonaspanin family |
| contig_27069_6 | 0.191 | 1 | 1 | 0.3246 | 1 | 1 | 0.1381 | predicted protein |
| contig_10149_5 | 0.1923 | 1 | 1 | 0.3264 | 1 | 1 | 0.1389 | phosphoribosylformylglycinamidine synthase |
| contig_27544_6 | 0.1934 | 1 | 1 | 0.3279 | 1 | 1 | 0.1395 | Long-chain-fatty-acid--CoA ligase putative |
| contig_30338_1 | 0.1939 | 1 | 1 | 0.3285 | 1 | 1 | 0.1398 | 60S ribosomal protein L5 |
| contig_23336_4 | 0.1949 | 1 | 1 | 0.3297 | 1 | 1 | 0.1403 | 1-hydroxy-2-methyl-2-(E)-butenyl-4-diphosphate synthase |
| contig_10991_6 | 0.195 | 1 | 1 | 0.3297 | 1 | 1 | 0.1403 | ABC transporter |
| contig_40323_4 | 0.1963 | 1 | 1 | 0.3314 | 1 | 1 | 0.141 | ---NA--- |
| contig_39484_1 | 0.1968 | 1 | 1 | 0.3317 | 1 | 1 | 0.1412 | probable aspartyl aminopeptidase |
| contig_7982_4 | 0.1969 | 1 | 1 | 0.3317 | 1 | 1 | 0.1412 | photosystem II protein V (chloroplast) |
| contig_22654_5 | 0.1974 | 1 | 1 | 0.3323 | 1 | 1 | 0.1414 | predicted protein |
| contig_19155_3 | 0.2006 | 1 | 1 | 0.3372 | 1 | 1 | 0.1435 | geranylgeranyl reductase |
| contig_25429_4 | 0.2023 | 1 | 1 | 0.3396 | 1 | 1 | 0.1445 | V-type proton ATPase proteolipid subunit |
| contig_27839_5 | 0.2027 | 1 | 1 | 0.3398 | 1 | 1 | 0.1446 | predicted protein |
| contig_33729_4 | 0.2028 | 1 | 1 | 0.3398 | 1 | 1 | 0.1446 | secreted protein |
| contig_15455_3 | 0.2033 | 1 | 1 | 0.3403 | 1 | 1 | 0.1448 | 1 2-dihydroxy-3-keto-5-methylthiopentene dioxygenase |
| contig_11344_5 | 0.2045 | 1 | 1 | 0.3419 | 1 | 1 | 0.1455 | Iron-sulfur cluster co-chaperone protein HscB mitochondrial |
| contig_11742_4 | 0.2054 | 1 | 1 | 0.3429 | 1 | 1 | 0.1459 | predicted protein |
| contig_31285_5 | 0.2057 | 1 | 1 | 0.3431 | 1 | 1 | 0.146 | threonine ammonia-lyase biosynthetic |
| contig_30510_4 | 0.2059 | 1 | 1 | 0.3431 | 1 | 1 | 0.146 | 60S ribosomal protein |
| contig_34843_5 | 0.2078 | 1 | 1 | 0.3456 | 1 | 1 | 0.1471 | Major Facilitator Superfamily (MFS) |
| contig_17153_5 | 0.2079 | 1 | 1 | 0.3456 | 1 | 1 | 0.1471 | predicted protein |
| contig_28947_4 | 0.2119 | 1 | 1 | 0.3519 | 1 | 1 | 0.1497 | predicted protein |
| contig_24784_5 | 0.2132 | 1 | 1 | 0.3533 | 1 | 1 | 0.1504 | DUF423-domain-containing protein |
| contig_20174_5 | 0.2132 | 1 | 1 | 0.3533 | 1 | 1 | 0.1504 | glutathione S-transferase |
| contig_23471_1 | 0.2154 | 1 | 1 | 0.3566 | 1 | 1 | 0.1517 | predicted protein |
| contig_20846_4 | 0.2159 | 1 | 1 | 0.3568 | 1 | 1 | 0.1518 | fructose-bisphosphate aldolase |

| | | | | | | | | |
|---|---|---|---|---|---|---|---|---|
| contig_12711_6 | 0.216 | 1 | 1 | 0.3568 | 1 | 1 | 0.1518 | predicted protein |
| contig_22753_2 | 0.2168 | 1 | 1 | 0.3577 | 1 | 1 | 0.1522 | nicotinate phosphoribosyltransferase 2-like |
| contig_25351_2 | 0.217 | 1 | 1 | 0.3577 | 1 | 1 | 0.1522 | predicted protein |
| contig_6058_5 | 0.2173 | 1 | 1 | 0.3578 | 1 | 1 | 0.1522 | predicted protein |
| contig_24756_6 | 0.2176 | 1 | 1 | 0.3578 | 1 | 1 | 0.1522 | tropinone reductase |
| contig_30824_4 | 0.2178 | 1 | 1 | 0.3578 | 1 | 1 | 0.1522 | predicted protein |
| contig_16381_6 | 0.2182 | 1 | 1 | 0.3582 | 1 | 1 | 0.1524 | predicted protein |
| contig_33515_6 | 0.2187 | 1 | 1 | 0.3586 | 1 | 1 | 0.1526 | predicted protein |
| contig_30573_3 | 0.2194 | 1 | 1 | 0.3594 | 1 | 1 | 0.1529 | predicted protein |
| contig_33090_4 | 0.2197 | 1 | 1 | 0.3594 | 1 | 1 | 0.1529 | predicted protein |
| contig_39564_2 | 0.221 | 1 | 1 | 0.3611 | 1 | 1 | 0.1536 | predicted protein |
| contig_29395_5 | 0.2221 | 1 | 1 | 0.3625 | 1 | 1 | 0.1542 | triosephosphate isomerase |
| contig_28707_5 | 0.2244 | 1 | 1 | 0.3659 | 1 | 1 | 0.1557 | V-type H+-transporting ATPase subunit a |
| contig_7643_4 | 0.2265 | 1 | 1 | 0.3686 | 1 | 1 | 0.1569 | ATP synthase CF1 beta subunit (chloroplast) |
| contig_9513_6 | 0.2265 | 1 | 1 | 0.3686 | 1 | 1 | 0.1569 | predicted protein |
| contig_28392_1 | 0.2274 | 1 | 1 | 0.3696 | 1 | 1 | 0.1573 | predicted protein |
| contig_28331_5 | 0.2289 | 1 | 1 | 0.3717 | 1 | 1 | 0.1582 | transketolase |
| contig_15868_5 | 0.2294 | 1 | 1 | 0.372 | 1 | 1 | 0.1583 | fucoxanthin-chlorophyll a-c binding protein |
| contig_28439_5 | 0.2296 | 1 | 1 | 0.372 | 1 | 1 | 0.1583 | 3-isopropylmalate dehydrogenase |
| contig_24581_4 | 0.2332 | 1 | 1 | 0.3774 | 1 | 1 | 0.1606 | predicted protein |
| contig_40422_5 | 0.2334 | 1 | 1 | 0.3774 | 1 | 1 | 0.1606 | light harvesting protein |
| contig_23787_5 | 0.2361 | 1 | 1 | 0.3814 | 1 | 1 | 0.1623 | predicted protein |
| contig_17212_4 | 0.2364 | 1 | 1 | 0.3814 | 1 | 1 | 0.1623 | uroporphyrinogen decarboxylase |
| contig_13739_4 | 0.2366 | 1 | 1 | 0.3814 | 1 | 1 | 0.1623 | protein DETOXIFICATION 46 chloroplastic |
| contig_24334_5 | 0.2387 | 1 | 1 | 0.3843 | 1 | 1 | 0.1635 | predicted protein |
| contig_27168_5 | 0.2403 | 1 | 1 | 0.3864 | 1 | 1 | 0.1644 | predicted protein |
| contig_8511_4 | 0.2407 | 1 | 1 | 0.3868 | 1 | 1 | 0.1646 | O-acetylhomoserine aminocarboxypropyltransferase |

| | | | | | | | | |
|---|---|---|---|---|---|---|---|---|
| contig_11147_6 | 0.2412 | 1 | 1 | 0.387 | 1 | 1 | 0.1647 | predicted protein |
| contig_23416_1 | 0.2419 | 1 | 1 | 0.3878 | 1 | 1 | 0.165 | predicted protein |
| contig_5402_3 | 0.2424 | 1 | 1 | 0.3882 | 1 | 1 | 0.1652 | NAD(P)-dependent oxidoreductase |
| contig_38523_3 | 0.2434 | 1 | 1 | 0.3893 | 1 | 1 | 0.1657 | h+-translocating pyrophosphatase family |
| contig_35155_5 | 0.2444 | 1 | 1 | 0.3905 | 1 | 1 | 0.1662 | fucoxanthin chlorophyll a/c protein 8 |
| contig_39064_3 | 0.2449 | 1 | 1 | 0.3906 | 1 | 1 | 0.1662 | Glycerophosphoryl diester phosphodiesterase |
| contig_39369_3 | 0.2449 | 1 | 1 | 0.3906 | 1 | 1 | 0.1662 | hypothetical protein FRACYDRAFT_269748 |
| contig_19181_5 | 0.2457 | 1 | 1 | 0.3915 | 1 | 1 | 0.1666 | ribosomal protein S4 |
| contig_27330_6 | 0.247 | 1 | 1 | 0.393 | 1 | 1 | 0.1672 | predicted protein |
| contig_6155_6 | 0.2473 | 1 | 1 | 0.393 | 1 | 1 | 0.1672 | light harvesting protein |
| contig_27320_3 | 0.2475 | 1 | 1 | 0.393 | 1 | 1 | 0.1672 | transketolase |
| contig_9471_1 | 0.2486 | 1 | 1 | 0.3945 | 1 | 1 | 0.1679 | photosystem II CP43 chlorophyll apoprotein (chloroplast) |
| contig_27589_4 | 0.25 | 1 | 1 | 0.3962 | 1 | 1 | 0.1686 | nucleolar protein 4 |
| contig_24818_1 | 0.2504 | 1 | 1 | 0.3965 | 1 | 1 | 0.1687 | UBC-like protein |
| contig_14363_4 | 0.2509 | 1 | 1 | 0.3968 | 1 | 1 | 0.1689 | predicted protein |
| contig_40248_2 | 0.2522 | 1 | 1 | 0.3984 | 1 | 1 | 0.1695 | predicted protein |
| contig_6009_6 | 0.2538 | 1 | 1 | 0.4005 | 1 | 1 | 0.1704 | 40S ribosomal protein S5 |
| contig_9153_4 | 0.2543 | 1 | 1 | 0.4009 | 1 | 1 | 0.1706 | predicted protein |
| contig_39202_4 | 0.2549 | 1 | 1 | 0.4015 | 1 | 1 | 0.1708 | Bystin |
| contig_21617_6 | 0.2575 | 1 | 1 | 0.4052 | 1 | 1 | 0.1724 | predicted protein |
| contig_39949_2 | 0.258 | 1 | 1 | 0.4054 | 1 | 1 | 0.1725 | predicted protein |
| contig_30410_4 | 0.2589 | 1 | 1 | 0.4065 | 1 | 1 | 0.173 | predicted protein |
| contig_18343_1 | 0.2592 | 1 | 1 | 0.4066 | 1 | 1 | 0.173 | glutamine synthetase |
| contig_29362_6 | 0.2622 | 1 | 1 | 0.4109 | 1 | 1 | 0.1748 | Stimulated by retinoic acid gene 6 protein |
| contig_30727_4 | 0.2642 | 1 | 1 | 0.4136 | 1 | 1 | 0.176 | predicted protein |
| contig_27872_3 | 0.2645 | 1 | 1 | 0.4136 | 1 | 1 | 0.176 | predicted protein |
| contig_17562_5 | 0.2654 | 1 | 1 | 0.4146 | 1 | 1 | 0.1764 | patched domain-containing protein 3-like |

| | | | | | | | | |
|---|---|---|---|---|---|---|---|---|
| contig_24083_4 | 0.2668 | 1 | 1 | 0.4159 | 1 | 1 | 0.177 | predicted protein |
| contig_24278_4 | 0.2668 | 1 | 1 | 0.4159 | 1 | 1 | 0.177 | predicted protein |
| contig_10600_6 | 0.2677 | 1 | 1 | 0.4166 | 1 | 1 | 0.1773 | predicted protein |
| contig_28105_5 | 0.2678 | 1 | 1 | 0.4166 | 1 | 1 | 0.1773 | regulatory proteasome non-atpase subunit 3 |
| contig_23768_5 | 0.2688 | 1 | 1 | 0.4177 | 1 | 1 | 0.1777 | Flavocytochrome c |
| contig_17548_1 | 0.269 | 1 | 1 | 0.4177 | 1 | 1 | 0.1777 | DUF1772 domain-containing protein |
| contig_28354_5 | 0.2695 | 1 | 1 | 0.418 | 1 | 1 | 0.1779 | hypothetical protein FRACYDRAFT_169912 |
| contig_30832_6 | 0.2719 | 1 | 1 | 0.4212 | 1 | 1 | 0.1792 | predicted protein |
| contig_39337_5 | 0.2728 | 1 | 1 | 0.4222 | 1 | 1 | 0.1797 | ---NA--- |
| contig_28484_6 | 0.275 | 1 | 1 | 0.4252 | 1 | 1 | 0.1809 | carbamoyl-phosphate synthase (glutamine-hydrolyzing) large subunit |
| contig_28304_6 | 0.2762 | 1 | 1 | 0.4267 | 1 | 1 | 0.1816 | predicted protein |
| contig_28195_3 | 0.2779 | 1 | 1 | 0.4288 | 1 | 1 | 0.1824 | predicted protein |
| contig_30555_4 | 0.2786 | 1 | 1 | 0.4294 | 1 | 1 | 0.1827 | fructose-1 6-bisphosphate aldolase precursor |
| contig_9763_5 | 0.2818 | 1 | 1 | 0.4339 | 1 | 1 | 0.1847 | predicted protein |
| contig_13388_5 | 0.283 | 1 | 1 | 0.4353 | 1 | 1 | 0.1852 | ornithine cyclodeaminase |
| contig_30444_6 | 0.2847 | 1 | 1 | 0.4376 | 1 | 1 | 0.1862 | NADH dehydrogenase (ubiquinone) 1 beta subcomplex subunit 9 |
| contig_26683_6 | 0.2866 | 1 | 1 | 0.4401 | 1 | 1 | 0.1873 | predicted protein |
| contig_32573_2 | 0.2901 | 1 | 1 | 0.4449 | 1 | 1 | 0.1893 | ATP synthase CF1 alpha subunit (chloroplast) |
| contig_35016_2 | 0.2932 | 1 | 1 | 0.4493 | 1 | 1 | 0.1912 | predicted protein |
| contig_30343_1 | 0.2937 | 1 | 1 | 0.4496 | 1 | 1 | 0.1913 | fucoxanthin chl a/c light-harvesting protein |
| contig_23117_2 | 0.2958 | 1 | 1 | 0.4522 | 1 | 1 | 0.1924 | peroxisomal (S)-2-hydroxy-acid oxidase GLO5-like |
| contig_10416_5 | 0.2964 | 1 | 1 | 0.4527 | 1 | 1 | 0.1927 | ATP synthase F0 subunit 6 (mitochondrion) |
| contig_21557_3 | 0.2986 | 1 | 1 | 0.4557 | 1 | 1 | 0.1939 | 40S ribosomal protein S2 |
| contig_37939_4 | 0.3005 | 1 | 1 | 0.4581 | 1 | 1 | 0.1949 | predicted protein |
| contig_20315_6 | 0.3021 | 1 | 1 | 0.46 | 1 | 1 | 0.1958 | NADH dehydrogenase subunit 5 (mitochondrion) |
| contig_41409_3 | 0.3039 | 1 | 1 | 0.4625 | 1 | 1 | 0.1968 | ribosomal protein L13 |
| contig_22487_6 | 0.3074 | 1 | 1 | 0.4673 | 1 | 1 | 0.1988 | predicted protein |

| | | | | | | | | |
|---|---|---|---|---|---|---|---|---|
| contig_37765_ 6 | 0.3093 | 1 | 1 | 0.46 96 | 1 | 1 | 0.19 98 | serine/threonine-protein phosphatase 2A regulatory subunit B' |
| contig_33628_ 1 | 0.3099 | 1 | 1 | 0.47 01 | 1 | 1 | 0.2 | glycine-rich domain-containing protein 2-like |
| contig_5869_1 | 0.311 | 1 | 1 | 0.47 09 | 1 | 1 | 0.20 04 | threonine-tRNA ligase |
| contig_17882_ 6 | 0.3112 | 1 | 1 | 0.47 09 | 1 | 1 | 0.20 04 | light harvesting protein |
| contig_16859_ 4 | 0.3116 | 1 | 1 | 0.47 09 | 1 | 1 | 0.20 04 | DNA-directed RNA polymerase II 135 kDa polypeptide |
| contig_15153_ 4 | 0.3117 | 1 | 1 | 0.47 09 | 1 | 1 | 0.20 04 | Ras-like GTP-binding protein YPT1 |
| contig_15354_ 3 | 0.3123 | 1 | 1 | 0.47 14 | 1 | 1 | 0.20 06 | predicted protein |
| contig_28369_ 6 | 0.3144 | 1 | 1 | 0.47 41 | 1 | 1 | 0.20 18 | predicted protein |
| contig_9649_2 | 0.3162 | 1 | 1 | 0.47 64 | 1 | 1 | 0.20 27 | predicted protein |
| contig_27341_ 1 | 0.3168 | 1 | 1 | 0.47 68 | 1 | 1 | 0.20 29 | protein patched homolog 2 |
| contig_27417_ 6 | 0.3189 | 1 | 1 | 0.47 94 | 1 | 1 | 0.20 4 | ferredoxin-nitrite reductase |
| contig_23301_ 2 | 0.3198 | 1 | 1 | 0.47 94 | 1 | 1 | 0.20 4 | protein fucoxanthin chlorophyll a/c protein |
| contig_30406_ 3 | 0.3203 | 1 | 1 | 0.47 94 | 1 | 1 | 0.20 4 | Glycylpeptide N-tetradecanoyltransferase 2 |
| contig_6119_2 | 0.3204 | 1 | 1 | 0.47 94 | 1 | 1 | 0.20 4 | putative tRNA (cytidine(32)/guanosine(34)-2'-O)-methyltransferase |
| contig_22809_ 5 | 0.3204 | 1 | 1 | 0.47 94 | 1 | 1 | 0.20 4 | predicted protein |
| contig_5949_4 | 0.3204 | 1 | 1 | 0.47 94 | 1 | 1 | 0.20 4 | predicted protein |
| contig_21707_ 3 | 0.3256 | 1 | 1 | 0.48 66 | 1 | 1 | 0.20 71 | 40S ribosomal protein S4 |
| contig_17402_ 3 | 0.3259 | 1 | 1 | 0.48 66 | 1 | 1 | 0.20 71 | hypothetical protein FRACYDRAFT_278091 |
| contig_22713_ 3 | 0.3287 | 1 | 1 | 0.48 9 | 1 | 1 | 0.20 81 | predicted protein |
| contig_22476_ 4 | 0.3287 | 1 | 1 | 0.48 9 | 1 | 1 | 0.20 81 | predicted protein |
| contig_30749_ 6 | 0.3287 | 1 | 1 | 0.48 9 | 1 | 1 | 0.20 81 | fucoxanthin-chlorophyll a/c light-harvesting protein |
| contig_27060_ 3 | 0.3288 | 1 | 1 | 0.48 9 | 1 | 1 | 0.20 81 | calcium-binding protein |
| contig_28112_ 2 | 0.3318 | 1 | 1 | 0.49 3 | 1 | 1 | 0.20 98 | predicted protein |
| contig_6511_6 | 0.3326 | 1 | 1 | 0.49 34 | 1 | 1 | 0.20 99 | glutamyl-tRNA synthetase |
| contig_32725_ 2 | 0.3327 | 1 | 1 | 0.49 34 | 1 | 1 | 0.20 99 | predicted protein |
| contig_24590_ 4 | 0.3335 | 1 | 1 | 0.49 41 | 1 | 1 | 0.21 02 | acyl-CoA dehydrogenase NM domain-like protein |

| | | | | | | | | |
|---|---|---|---|---|---|---|---|---|
| contig_24802_3 | 0.3352 | 1 | 1 | 0.4962 | 1 | 1 | 0.2111 | predicted protein |
| contig_9038_6 | 0.3374 | 1 | 1 | 0.4982 | 1 | 1 | 0.212 | methionine-tRNA ligase beta subunit |
| contig_14552_5 | 0.3377 | 1 | 1 | 0.4982 | 1 | 1 | 0.212 | phosphomannomutase |
| contig_39611_1 | 0.3377 | 1 | 1 | 0.4982 | 1 | 1 | 0.212 | methionine-tRNA ligase |
| contig_26806_4 | 0.3379 | 1 | 1 | 0.4982 | 1 | 1 | 0.212 | rab11 protein |
| contig_15547_4 | 0.3396 | 1 | 1 | 0.5002 | 1 | 1 | 0.2129 | hypothetical protein FRACYDRAFT_234640 |
| contig_8371_4 | 0.3414 | 1 | 1 | 0.5024 | 1 | 1 | 0.2138 | Rab family other |
| contig_24804_5 | 0.342 | 1 | 1 | 0.5028 | 1 | 1 | 0.214 | ribosomal protein L13 |
| contig_33544_3 | 0.3424 | 1 | 1 | 0.5028 | 1 | 1 | 0.214 | predicted protein |
| contig_40230_1 | 0.3428 | 1 | 1 | 0.5029 | 1 | 1 | 0.214 | predicted protein |
| contig_17076_5 | 0.3444 | 1 | 1 | 0.5048 | 1 | 1 | 0.2148 | predicted protein |
| contig_16920_6 | 0.3448 | 1 | 1 | 0.5049 | 1 | 1 | 0.2148 | predicted protein |
| contig_9617_4 | 0.3464 | 1 | 1 | 0.5057 | 1 | 1 | 0.2152 | bifunctional methylenetetrahydrofolate dehydrogenase/methenyltetrahydrofolate cyclohydrolase FolD |
| contig_38515_1 | 0.3465 | 1 | 1 | 0.5057 | 1 | 1 | 0.2152 | predicted protein |
| contig_33419_4 | 0.3467 | 1 | 1 | 0.5057 | 1 | 1 | 0.2152 | light harvesting protein |
| contig_8369_4 | 0.3468 | 1 | 1 | 0.5057 | 1 | 1 | 0.2152 | Clp protease ATP binding subunit (chloroplast) |
| contig_29346_4 | 0.347 | 1 | 1 | 0.5057 | 1 | 1 | 0.2152 | homogentisate solanesyltransferase |
| contig_17493_4 | 0.3478 | 1 | 1 | 0.5062 | 1 | 1 | 0.2154 | lamin tail domain-containing protein |
| contig_6467_5 | 0.348 | 1 | 1 | 0.5062 | 1 | 1 | 0.2154 | predicted protein |
| contig_21612_4 | 0.3485 | 1 | 1 | 0.5065 | 1 | 1 | 0.2155 | 60S ribosomal protein L27 |
| contig_8422_4 | 0.3498 | 1 | 1 | 0.5078 | 1 | 1 | 0.2161 | predicted protein |
| contig_28911_6 | 0.3551 | 1 | 1 | 0.5151 | 1 | 1 | 0.2192 | eukaryotic translation initiation factor 6 |
| contig_15995_6 | 0.3555 | 1 | 1 | 0.5152 | 1 | 1 | 0.2192 | cyclopropane-fatty-acyl-phospholipid synthase |
| contig_11948_3 | 0.3561 | 1 | 1 | 0.5156 | 1 | 1 | 0.2194 | aldo/keto reductase |
| contig_28340_4 | 0.3585 | 1 | 1 | 0.5184 | 1 | 1 | 0.2206 | Ras-like protein Rab-11B |
| contig_25423_6 | 0.3603 | 1 | 1 | 0.5206 | 1 | 1 | 0.2215 | 2 5-diketo-D-gluconic acid reductase A putative |

| | | | | | | | | |
|---|---|---|---|---|---|---|---|---|
| contig_5961_5 | 0.3653 | 1 | 1 | 0.5274 | 1 | 1 | 0.2244 | cytochrome oxidase subunit III (mitochondrion) |
| contig_11835_4 | 0.367 | 1 | 1 | 0.5293 | 1 | 1 | 0.2252 | ion transporter |
| contig_28140_1 | 0.3708 | 1 | 1 | 0.5342 | 1 | 1 | 0.2273 | citrate (Si)-synthase eukaryotic |
| contig_9759_2 | 0.3715 | 1 | 1 | 0.5347 | 1 | 1 | 0.2275 | clathrin heavy chain |
| contig_23621_6 | 0.3719 | 1 | 1 | 0.5348 | 1 | 1 | 0.2276 | predicted protein |
| contig_29986_5 | 0.3728 | 1 | 1 | 0.535 | 1 | 1 | 0.2277 | H - or Na -translocating F-type V-type and A-type ATPase (F-ATPase) Superfamily |
| contig_33374_5 | 0.3737 | 1 | 1 | 0.535 | 1 | 1 | 0.2277 | predicted protein |
| contig_6339_4 | 0.3738 | 1 | 1 | 0.535 | 1 | 1 | 0.2277 | predicted protein |
| contig_5862_2 | 0.3742 | 1 | 1 | 0.535 | 1 | 1 | 0.2277 | arylsulfatase |
| contig_24778_4 | 0.3745 | 1 | 1 | 0.535 | 1 | 1 | 0.2277 | predicted protein |
| contig_28951_4 | 0.3747 | 1 | 1 | 0.535 | 1 | 1 | 0.2277 | predicted protein |
| contig_33723_4 | 0.3747 | 1 | 1 | 0.535 | 1 | 1 | 0.2277 | predicted protein |
| contig_28208_6 | 0.3748 | 1 | 1 | 0.535 | 1 | 1 | 0.2277 | SPFH/Band 7/PHB domain protein |
| contig_31457_1 | 0.3758 | 1 | 1 | 0.536 | 1 | 1 | 0.2281 | predicted protein |
| contig_30540_5 | 0.38 | 1 | 1 | 0.5413 | 1 | 1 | 0.2304 | unnamed protein product |
| contig_34883_6 | 0.3804 | 1 | 1 | 0.5414 | 1 | 1 | 0.2304 | Fructose-bisphosphate aldolase |
| contig_17114_6 | 0.3809 | 1 | 1 | 0.5417 | 1 | 1 | 0.2305 | predicted protein |
| contig_28740_4 | 0.3814 | 1 | 1 | 0.5418 | 1 | 1 | 0.2306 | tubulin alpha chain |
| contig_23244_4 | 0.3839 | 1 | 1 | 0.545 | 1 | 1 | 0.2319 | predicted protein |
| contig_35847_5 | 0.3847 | 1 | 1 | 0.5452 | 1 | 1 | 0.232 | serine/threonine-protein phosphatase PP1-beta catalytic subunit |
| contig_25034_6 | 0.3848 | 1 | 1 | 0.5452 | 1 | 1 | 0.232 | predicted protein |
| contig_6296_5 | 0.3882 | 1 | 1 | 0.5494 | 1 | 1 | 0.2338 | predicted protein |
| contig_22941_4 | 0.3897 | 1 | 1 | 0.5509 | 1 | 1 | 0.2344 | mitochondrial carrier |
| contig_5668_1 | 0.3899 | 1 | 1 | 0.5509 | 1 | 1 | 0.2344 | 40s ribosomal protein s20 |
| contig_24888_1 | 0.3938 | 1 | 1 | 0.5558 | 1 | 1 | 0.2365 | chromodomain-helicase-DNA-binding protein 7 |
| contig_8256_2 | 0.3969 | 1 | 1 | 0.5587 | 1 | 1 | 0.2378 | importin subunit beta putative |

| | | | | | | | | |
|---|---|---|---|---|---|---|---|---|
| contig_32615_1 | 0.3969 | 1 | 1 | 0.5587 | 1 | 1 | 0.2378 | predicted protein |
| contig_3408_5 | 0.397 | 1 | 1 | 0.5587 | 1 | 1 | 0.2378 | fucoxanthin-chlorophyll a-c binding protein |
| contig_26990_2 | 0.3976 | 1 | 1 | 0.5591 | 1 | 1 | 0.2379 | uracil phosphoribosyltransferase |
| contig_25874_5 | 0.3982 | 1 | 1 | 0.5595 | 1 | 1 | 0.2381 | predicted protein |
| contig_28185_5 | 0.3998 | 1 | 1 | 0.5607 | 1 | 1 | 0.2386 | UMP-CMP kinase |
| contig_14140_6 | 0.3999 | 1 | 1 | 0.5607 | 1 | 1 | 0.2386 | predicted protein |
| contig_16247_6 | 0.4002 | 1 | 1 | 0.5607 | 1 | 1 | 0.2386 | imidazoleglycerol-phosphate dehydratase |
| contig_27149_6 | 0.401 | 1 | 1 | 0.5613 | 1 | 1 | 0.2388 | homogentisate phytyltransferase |
| contig_19241_5 | 0.4018 | 1 | 1 | 0.5616 | 1 | 1 | 0.239 | ---NA--- |
| contig_22787_6 | 0.4019 | 1 | 1 | 0.5616 | 1 | 1 | 0.239 | bifunctional aspartate kinase/homoserine dehydrogenase I |
| contig_6105_6 | 0.4051 | 1 | 1 | 0.5655 | 1 | 1 | 0.2406 | photosystem II CP47 chlorophyll apoprotein (chloroplast) |
| contig_19141_5 | 0.4064 | 1 | 1 | 0.5667 | 1 | 1 | 0.2412 | photosystem I subunit IV (chloroplast) |
| contig_7075_5 | 0.4103 | 1 | 1 | 0.5718 | 1 | 1 | 0.2433 | predicted protein |
| contig_24941_5 | 0.4114 | 1 | 1 | 0.5725 | 1 | 1 | 0.2436 | triosephosphate isomerase |
| contig_17279_6 | 0.4116 | 1 | 1 | 0.5725 | 1 | 1 | 0.2436 | vacuolar protein sorting-associated protein 26 |
| contig_36986_1 | 0.4134 | 1 | 1 | 0.5744 | 1 | 1 | 0.2444 | ---NA--- |
| contig_35523_6 | 0.4159 | 1 | 1 | 0.5774 | 1 | 1 | 0.2457 | 60S ribosomal protein L10a |
| contig_23607_2 | 0.418 | 1 | 1 | 0.5798 | 1 | 1 | 0.2467 | alanyl-tRNA synthetase |
| contig_18307_6 | 0.4186 | 1 | 1 | 0.5801 | 1 | 1 | 0.2469 | predicted protein |
| contig_25721_4 | 0.4202 | 1 | 1 | 0.5817 | 1 | 1 | 0.2475 | putative manganese superoxide dismutase protein |
| contig_38364_2 | 0.4236 | 1 | 1 | 0.586 | 1 | 1 | 0.2494 | predicted protein |
| contig_28468_2 | 0.424 | 1 | 1 | 0.586 | 1 | 1 | 0.2494 | ribosome recycling factor |
| contig_38581_3 | 0.4273 | 1 | 1 | 0.59 | 1 | 1 | 0.2511 | predicted protein |
| contig_9468_6 | 0.4293 | 1 | 1 | 0.5923 | 1 | 1 | 0.252 | anaphase-promoting complex subunit 1 |
| contig_33488_4 | 0.4315 | 1 | 1 | 0.5947 | 1 | 1 | 0.2531 | transcription elongation factor SPT6 |
| contig_40734_1 | 0.432 | 1 | 1 | 0.5949 | 1 | 1 | 0.2532 | predicted protein |
| contig_23548_5 | 0.4328 | 1 | 1 | 0.5955 | 1 | 1 | 0.2534 | ATP synthase subunit beta mitochondrial |

| | | | | | | | | |
|---|---|---|---|---|---|---|---|---|
| contig_26174_3 | 0.4334 | 1 | 1 | 0.5958 | 1 | 1 | 0.2535 | predicted protein |
| contig_32710_4 | 0.4351 | 1 | 1 | 0.5975 | 1 | 1 | 0.2543 | class I SAM-dependent methyltransferase |
| contig_37718_6 | 0.4371 | 1 | 1 | 0.5997 | 1 | 1 | 0.2552 | predicted protein |
| contig_24921_6 | 0.4382 | 1 | 1 | 0.6007 | 1 | 1 | 0.2556 | nitrate transporter |
| contig_22050_5 | 0.4417 | 1 | 1 | 0.6049 | 1 | 1 | 0.2574 | 60S ribosomal protein L9 |
| contig_7031_3 | 0.4421 | 1 | 1 | 0.6049 | 1 | 1 | 0.2574 | transmembrane 9 superfamily member 3 |
| contig_27279_6 | 0.4429 | 1 | 1 | 0.6055 | 1 | 1 | 0.2577 | adenosine kinase |
| contig_38793_1 | 0.446 | 1 | 1 | 0.6092 | 1 | 1 | 0.2592 | predicted protein |
| contig_32599_2 | 0.4468 | 1 | 1 | 0.6097 | 1 | 1 | 0.2595 | ATP-dependent DNA helicase PIF1-like |
| contig_11202_6 | 0.4492 | 1 | 1 | 0.6124 | 1 | 1 | 0.2606 | predicted protein |
| contig_37899_3 | 0.4522 | 1 | 1 | 0.6156 | 1 | 1 | 0.262 | ribosomal protein L2 (chloroplast) |
| contig_26135_5 | 0.4523 | 1 | 1 | 0.6156 | 1 | 1 | 0.262 | Mitochondrial Carrier (MC) Family |
| contig_26792_2 | 0.4535 | 1 | 1 | 0.6167 | 1 | 1 | 0.2624 | predicted protein |
| contig_28207_5 | 0.4543 | 1 | 1 | 0.6173 | 1 | 1 | 0.2627 | mannose-P-dolichol utilization defect 1 protein |
| contig_7122_5 | 0.4555 | 1 | 1 | 0.6182 | 1 | 1 | 0.2631 | ---NA--- |
| contig_6303_5 | 0.4566 | 1 | 1 | 0.619 | 1 | 1 | 0.2634 | acyl carrier protein (chloroplast) |
| contig_34472_2 | 0.4568 | 1 | 1 | 0.619 | 1 | 1 | 0.2634 | histone h4 |
| contig_26815_5 | 0.4576 | 1 | 1 | 0.6195 | 1 | 1 | 0.2636 | AGC/PKA protein kinase |
| contig_24961_6 | 0.4582 | 1 | 1 | 0.6197 | 1 | 1 | 0.2637 | mitochondrial carrier |
| contig_9857_5 | 0.4598 | 1 | 1 | 0.6213 | 1 | 1 | 0.2644 | ---NA--- |
| contig_6653_5 | 0.4602 | 1 | 1 | 0.6213 | 1 | 1 | 0.2644 | chorismate mutase |
| contig_8596_6 | 0.462 | 1 | 1 | 0.6232 | 1 | 1 | 0.2652 | predicted protein |
| contig_15022_4 | 0.4627 | 1 | 1 | 0.6236 | 1 | 1 | 0.2653 | ---NA--- |
| contig_21238_5 | 0.4635 | 1 | 1 | 0.6242 | 1 | 1 | 0.2656 | fucoxanthin chlorophyll a/c protein 8 |
| contig_38910_1 | 0.4689 | 1 | 1 | 0.6308 | 1 | 1 | 0.2684 | predicted protein |
| contig_25602_3 | 0.4708 | 1 | 1 | 0.6329 | 1 | 1 | 0.2693 | protein transport protein SEC61 subunit alpha |
| contig_23751_6 | 0.4717 | 1 | 1 | 0.6332 | 1 | 1 | 0.2694 | high mobility group protein B2 |

| | | | | | | | | |
|---|---|---|---|---|---|---|---|---|
| contig_17463_2 | 0.4719 | 1 | 1 | 0.6332 | 1 | 1 | 0.2694 | glucose-6-phosphate isomerase |
| contig_27061_1 | 0.4739 | 1 | 1 | 0.6353 | 1 | 1 | 0.2703 | acetyl-CoA carboxylase biotin carboxylase subunit |
| contig_5503_5 | 0.4749 | 1 | 1 | 0.6361 | 1 | 1 | 0.2707 | predicted protein |
| contig_27039_5 | 0.4767 | 1 | 1 | 0.6379 | 1 | 1 | 0.2715 | hypothetical protein THAOC_36140 |
| contig_34760_6 | 0.485 | 1 | 1 | 0.6486 | 1 | 1 | 0.276 | low molecular mass early light-inducible protein HV60 |
| contig_9265_3 | 0.4859 | 1 | 1 | 0.6491 | 1 | 1 | 0.2762 | mitochondrial carrier family |
| contig_11658_2 | 0.4879 | 1 | 1 | 0.6513 | 1 | 1 | 0.2772 | pyrimidine/purine nucleoside phosphorylase |
| contig_26772_5 | 0.4891 | 1 | 1 | 0.6523 | 1 | 1 | 0.2776 | predicted protein |
| contig_30838_5 | 0.4901 | 1 | 1 | 0.6531 | 1 | 1 | 0.2779 | histone H4 |
| contig_33542_2 | 0.4915 | 1 | 1 | 0.6543 | 1 | 1 | 0.2784 | nucleoside-diphosphate kinase |
| contig_30686_5 | 0.493 | 1 | 1 | 0.6554 | 1 | 1 | 0.2789 | chlorophyll synthase chloroplastic |
| contig_28383_6 | 0.4936 | 1 | 1 | 0.6554 | 1 | 1 | 0.2789 | predicted protein |
| contig_25876_4 | 0.4936 | 1 | 1 | 0.6554 | 1 | 1 | 0.2789 | predicted protein |
| contig_15143_1 | 0.4958 | 1 | 1 | 0.6573 | 1 | 1 | 0.2797 | CK1/CK1/CK1-D protein kinase variant |
| contig_5809_2 | 0.4962 | 1 | 1 | 0.6573 | 1 | 1 | 0.2797 | ---NA--- |
| contig_13878_2 | 0.4963 | 1 | 1 | 0.6573 | 1 | 1 | 0.2797 | glycine--tRNA ligase |
| contig_11813_1 | 0.4968 | 1 | 1 | 0.6573 | 1 | 1 | 0.2797 | predicted protein |
| contig_38406_2 | 0.4973 | 1 | 1 | 0.6574 | 1 | 1 | 0.2797 | predicted protein |
| contig_30486_2 | 0.4977 | 1 | 1 | 0.6574 | 1 | 1 | 0.2797 | signal peptidase I |
| contig_35608_4 | 0.4986 | 1 | 1 | 0.658 | 1 | 1 | 0.28 | chaperone dnaj-like protein |
| contig_13588_3 | 0.4993 | 1 | 1 | 0.6584 | 1 | 1 | 0.2802 | aarF domain-containing kinase |
| contig_38299_1 | 0.5015 | 1 | 1 | 0.6594 | 1 | 1 | 0.2806 | ATP-binding cassette superfamily |
| contig_39493_2 | 0.5017 | 1 | 1 | 0.6594 | 1 | 1 | 0.2806 | 60s ribosomal protein |
| contig_33008_5 | 0.5019 | 1 | 1 | 0.6594 | 1 | 1 | 0.2806 | protein fucoxanthin chlorophyll a/c protein |
| contig_26362_4 | 0.5021 | 1 | 1 | 0.6594 | 1 | 1 | 0.2806 | predicted protein |
| contig_30747_4 | 0.5022 | 1 | 1 | 0.6594 | 1 | 1 | 0.2806 | bile acid:Na+ symporter BASS family |
| contig_10800_3 | 0.503 | 1 | 1 | 0.6599 | 1 | 1 | 0.2808 | predicted protein |

| | | | | | | | | |
|---|---|---|---|---|---|---|---|---|
| contig_5385_3 | 0.5037 | 1 | 1 | 0.6602 | 1 | 1 | 0.281 | glutathione S-transferase Mu 3-like |
| contig_11991_5 | 0.5079 | 1 | 1 | 0.6643 | 1 | 1 | 0.2827 | ---NA--- |
| contig_25825_6 | 0.508 | 1 | 1 | 0.6643 | 1 | 1 | 0.2827 | predicted protein |
| contig_8332_1 | 0.5082 | 1 | 1 | 0.6643 | 1 | 1 | 0.2827 | oxoglutarate dehydrogenase (succinyl-transferring) E1 component |
| contig_41371_1 | 0.5087 | 1 | 1 | 0.6644 | 1 | 1 | 0.2827 | 40s ribosomal protein |
| contig_5747_6 | 0.5104 | 1 | 1 | 0.6661 | 1 | 1 | 0.2835 | photosystem I assembly protein Ycf4 (chloroplast) |
| contig_11002_5 | 0.5117 | 1 | 1 | 0.6671 | 1 | 1 | 0.2839 | peroxiredoxin |
| contig_38390_3 | 0.5123 | 1 | 1 | 0.6671 | 1 | 1 | 0.2839 | predicted protein |
| contig_41308_6 | 0.5125 | 1 | 1 | 0.6671 | 1 | 1 | 0.2839 | nucleoside diphosphate kinase |
| contig_25076_6 | 0.5136 | 1 | 1 | 0.668 | 1 | 1 | 0.2842 | phosphoglucomutase |
| contig_8878_1 | 0.5144 | 1 | 1 | 0.6684 | 1 | 1 | 0.2844 | predicted protein |
| contig_29801_5 | 0.5149 | 1 | 1 | 0.6686 | 1 | 1 | 0.2845 | triosephosphate isomerase |
| contig_27972_6 | 0.5158 | 1 | 1 | 0.6687 | 1 | 1 | 0.2845 | atp-dependent rna helicase dbp5 |
| contig_27313_6 | 0.5159 | 1 | 1 | 0.6687 | 1 | 1 | 0.2845 | predicted protein |
| contig_10238_6 | 0.5164 | 1 | 1 | 0.6687 | 1 | 1 | 0.2846 | ras-like gtp-binding protein ypt1 |
| contig_7026_3 | 0.5168 | 1 | 1 | 0.6687 | 1 | 1 | 0.2846 | 26S proteasome non-ATPase regulatory subunit 2 |
| contig_29498_6 | 0.5173 | 1 | 1 | 0.6688 | 1 | 1 | 0.2846 | aldehyde dehydrogenase |
| contig_9710_5 | 0.5179 | 1 | 1 | 0.6689 | 1 | 1 | 0.2847 | predicted protein |
| contig_32573_1 | 0.5183 | 1 | 1 | 0.6689 | 1 | 1 | 0.2847 | ATP synthase CF1 alpha subunit (chloroplast) |
| contig_28970_6 | 0.5196 | 1 | 1 | 0.6701 | 1 | 1 | 0.2851 | peptide methionine sulfoxide reductase |
| contig_5663_5 | 0.5208 | 1 | 1 | 0.6711 | 1 | 1 | 0.2856 | tubulin beta chain |
| contig_33218_4 | 0.5222 | 1 | 1 | 0.6719 | 1 | 1 | 0.2859 | predicted protein |
| contig_28796_6 | 0.5227 | 1 | 1 | 0.6719 | 1 | 1 | 0.2859 | predicted protein |
| contig_38064_1 | 0.5228 | 1 | 1 | 0.6719 | 1 | 1 | 0.2859 | phosphoenolpyruvate carboxylase |
| contig_6525_2 | 0.5239 | 1 | 1 | 0.6723 | 1 | 1 | 0.2861 | probable aminopeptidase NPEPL1 |
| contig_27862_5 | 0.5241 | 1 | 1 | 0.6723 | 1 | 1 | 0.2861 | predicted protein |

| | | | | | | | | |
|---|---|---|---|---|---|---|---|---|
| contig_27983_4 | 0.5244 | 1 | 1 | 0.6723 | 1 | 1 | 0.2861 | alpha/beta hydrolase |
| contig_23501_5 | 0.5253 | 1 | 1 | 0.6728 | 1 | 1 | 0.2863 | bile acid:sodium symporter |
| contig_18033_5 | 0.5265 | 1 | 1 | 0.6738 | 1 | 1 | 0.2867 | mitochondrial-processing peptidase subunit beta |
| contig_39259_1 | 0.5302 | 1 | 1 | 0.6781 | 1 | 1 | 0.2885 | preprotein-translocase subunit a (chloroplast) |
| contig_29172_6 | 0.5308 | 1 | 1 | 0.6781 | 1 | 1 | 0.2886 | arginyl-tRNA synthetase |
| contig_33877_5 | 0.532 | 1 | 1 | 0.6792 | 1 | 1 | 0.289 | ---NA--- |
| contig_5530_1 | 0.5361 | 1 | 1 | 0.6838 | 1 | 1 | 0.291 | NAD(P)-dependent steroid dehydrogenase-like protein |
| contig_10909_2 | 0.5366 | 1 | 1 | 0.6839 | 1 | 1 | 0.291 | Glycerol-3-phosphate dehydrogenase mitochondrial |
| contig_12035_1 | 0.538 | 1 | 1 | 0.6851 | 1 | 1 | 0.2915 | predicted protein |
| contig_34263_4 | 0.5401 | 1 | 1 | 0.6872 | 1 | 1 | 0.2924 | nucleolar protein 56 |
| contig_39293_4 | 0.5423 | 1 | 1 | 0.6892 | 1 | 1 | 0.2933 | predicted protein |
| contig_17991_3 | 0.5426 | 1 | 1 | 0.6892 | 1 | 1 | 0.2933 | light-harvesting protein |
| contig_24459_5 | 0.5441 | 1 | 1 | 0.6905 | 1 | 1 | 0.2938 | fact complex subunit ssrp1 |
| contig_15835_5 | 0.5451 | 1 | 1 | 0.6913 | 1 | 1 | 0.2942 | ---NA--- |
| contig_9532_4 | 0.5462 | 1 | 1 | 0.692 | 1 | 1 | 0.2945 | NADH dehydrogenase [ubiquinone] iron-sulfur protein 8-A mitochondrial |
| contig_3295_5 | 0.5467 | 1 | 1 | 0.6921 | 1 | 1 | 0.2945 | V-type ATPase F subunit |
| contig_34431_6 | 0.5473 | 1 | 1 | 0.6923 | 1 | 1 | 0.2946 | predicted protein |
| contig_9855_4 | 0.5487 | 1 | 1 | 0.6931 | 1 | 1 | 0.2949 | S-adenosyl-L-methionine-dependent methyltransferase |
| contig_6218_5 | 0.5488 | 1 | 1 | 0.6931 | 1 | 1 | 0.2949 | ferredoxin-nitrite reductase |
| contig_7217_3 | 0.551 | 1 | 1 | 0.6952 | 1 | 1 | 0.2958 | P-loop containing nucleoside triphosphate hydrolase protein |
| contig_41228_1 | 0.5516 | 1 | 1 | 0.6952 | 1 | 1 | 0.2958 | light-harvesting protein |
| contig_10952_4 | 0.5519 | 1 | 1 | 0.6952 | 1 | 1 | 0.2958 | diacylglycerol O-acyltransferase 2D |
| contig_41396_3 | 0.5551 | 1 | 1 | 0.6987 | 1 | 1 | 0.2973 | predicted protein |
| contig_24751_3 | 0.5573 | 1 | 1 | 0.7009 | 1 | 1 | 0.2983 | predicted protein |
| contig_16774_4 | 0.562 | 1 | 1 | 0.7055 | 1 | 1 | 0.3002 | probable microsomal very long chain fatty acid elongase |
| contig_13446_5 | 0.5622 | 1 | 1 | 0.7055 | 1 | 1 | 0.3002 | predicted protein |
| contig_12155_4 | 0.5624 | 1 | 1 | 0.7055 | 1 | 1 | 0.3002 | predicted protein |

| | | | | | | | | |
|---|---|---|---|---|---|---|---|---|
| contig_9129_6 | 0.5638 | 1 | 1 | 0.7068 | 1 | 1 | 0.3008 | predicted protein |
| contig_21644_6 | 0.5662 | 1 | 1 | 0.7091 | 1 | 1 | 0.3018 | putative manganese superoxide dismutase protein |
| contig_9571_3 | 0.5668 | 1 | 1 | 0.7094 | 1 | 1 | 0.3019 | ATP:ADP antiporter AAA family |
| contig_16087_6 | 0.5684 | 1 | 1 | 0.7107 | 1 | 1 | 0.3024 | predicted protein |
| contig_24829_2 | 0.571 | 1 | 1 | 0.7134 | 1 | 1 | 0.3036 | ornithine carbamoyltransferase |
| contig_32529_3 | 0.5727 | 1 | 1 | 0.7149 | 1 | 1 | 0.3042 | GDT1-like protein 2 chloroplastic |
| contig_25557_4 | 0.5733 | 1 | 1 | 0.7151 | 1 | 1 | 0.3043 | nucleolar complex protein 2 |
| contig_9214_2 | 0.5744 | 1 | 1 | 0.7156 | 1 | 1 | 0.3045 | AMP-dependent synthetase |
| contig_37607_6 | 0.5746 | 1 | 1 | 0.7156 | 1 | 1 | 0.3045 | 2-isopropylmalate synthase |
| contig_11653_4 | 0.5794 | 1 | 1 | 0.7209 | 1 | 1 | 0.3068 | COP9 signalosome complex subunit 3 |
| contig_25806_5 | 0.5817 | 1 | 1 | 0.7232 | 1 | 1 | 0.3077 | 2-C-methyl-D-erythritol 2 4-cyclodiphosphate synthase |
| contig_33296_6 | 0.5846 | 1 | 1 | 0.7262 | 1 | 1 | 0.309 | light-harvesting protein |
| contig_9488_1 | 0.5863 | 1 | 1 | 0.7275 | 1 | 1 | 0.3096 | 26s protease regulatory subunit 8 |
| contig_17799_3 | 0.5866 | 1 | 1 | 0.7275 | 1 | 1 | 0.3096 | light-harvesting protein |
| contig_27520_2 | 0.5874 | 1 | 1 | 0.7279 | 1 | 1 | 0.3097 | predicted protein |
| contig_33115_6 | 0.5882 | 1 | 1 | 0.7284 | 1 | 1 | 0.3099 | predicted protein |
| contig_9387_4 | 0.5892 | 1 | 1 | 0.729 | 1 | 1 | 0.3102 | DUF791-domain-containing protein |
| contig_16700_1 | 0.5914 | 1 | 1 | 0.7311 | 1 | 1 | 0.3111 | serine decarboxylase |
| contig_28417_5 | 0.6001 | 1 | 1 | 0.7413 | 1 | 1 | 0.3154 | predicted protein |
| contig_30562_5 | 0.6011 | 1 | 1 | 0.7419 | 1 | 1 | 0.3157 | cytochrome b6-f complex iron-sulfur subunit |
| contig_26112_6 | 0.602 | 1 | 1 | 0.7424 | 1 | 1 | 0.3159 | adenylate kinase 5 chloroplastic isoform X1 |
| contig_30717_5 | 0.6028 | 1 | 1 | 0.7427 | 1 | 1 | 0.3161 | predicted protein |
| contig_23843_1 | 0.6051 | 1 | 1 | 0.7448 | 1 | 1 | 0.3169 | methyltransferase domain-containing protein |
| contig_6572_6 | 0.6054 | 1 | 1 | 0.7448 | 1 | 1 | 0.3169 | catalase |
| contig_36071_6 | 0.6063 | 1 | 1 | 0.7453 | 1 | 1 | 0.3171 | predicted protein |
| contig_30482_3 | 0.6077 | 1 | 1 | 0.7464 | 1 | 1 | 0.3176 | predicted protein |
| contig_26788_4 | 0.6092 | 1 | 1 | 0.7474 | 1 | 1 | 0.3181 | regulatory proteasome non-atpase subunit 2 |

| | | | | | | | | |
|---|---|---|---|---|---|---|---|---|
| contig_29915_2 | 0.6095 | 1 | 1 | 0.7474 | 1 | 1 | 0.3181 | nucleolar protein nop56 |
| contig_30867_1 | 0.6108 | 1 | 1 | 0.7484 | 1 | 1 | 0.3185 | Light-harvesting complex I LH38 protein |
| contig_17202_2 | 0.6119 | 1 | 1 | 0.7486 | 1 | 1 | 0.3186 | predicted protein |
| contig_9006_3 | 0.612 | 1 | 1 | 0.7486 | 1 | 1 | 0.3186 | thiol-activated cytolysin |
| contig_37856_1 | 0.6137 | 1 | 1 | 0.7501 | 1 | 1 | 0.3192 | 3-phosphoserine/phosphohydroxythreonine transaminase |
| contig_38702_2 | 0.6175 | 1 | 1 | 0.7538 | 1 | 1 | 0.3208 | predicted protein |
| contig_22940_5 | 0.6177 | 1 | 1 | 0.7538 | 1 | 1 | 0.3208 | PH domain and leucine-rich repeat-containing protein phosphatase |
| contig_16580_4 | 0.62 | 1 | 1 | 0.756 | 1 | 1 | 0.3217 | casein kinase I |
| contig_25500_5 | 0.6231 | 1 | 1 | 0.7592 | 1 | 1 | 0.3231 | predicted protein |
| contig_29698_3 | 0.6238 | 1 | 1 | 0.7594 | 1 | 1 | 0.3232 | Light-harvesting complex I LH38 protein |
| contig_16105_5 | 0.6251 | 1 | 1 | 0.7604 | 1 | 1 | 0.3236 | hypothetical protein FRACYDRAFT_264519 |
| contig_40265_2 | 0.627 | 1 | 1 | 0.7618 | 1 | 1 | 0.3242 | predicted protein |
| contig_9646_5 | 0.6279 | 1 | 1 | 0.7618 | 1 | 1 | 0.3242 | cell differentiation protein |
| contig_37779_2 | 0.6289 | 1 | 1 | 0.7618 | 1 | 1 | 0.3242 | protein fucoxanthin chlorophyll a/c protein |
| contig_22796_6 | 0.6294 | 1 | 1 | 0.7618 | 1 | 1 | 0.3242 | predicted protein |
| contig_24919_4 | 0.6296 | 1 | 1 | 0.7618 | 1 | 1 | 0.3242 | predicted protein |
| contig_30385_5 | 0.6297 | 1 | 1 | 0.7618 | 1 | 1 | 0.3242 | myo-inositol-1(or 4)-monophosphatase |
| contig_24765_5 | 0.6298 | 1 | 1 | 0.7618 | 1 | 1 | 0.3242 | ABC transporter ATP-binding protein/permease |
| contig_41244_3 | 0.6323 | 1 | 1 | 0.7637 | 1 | 1 | 0.325 | class II fructose-bisphosphate aldolase |
| contig_5445_2 | 0.6323 | 1 | 1 | 0.7637 | 1 | 1 | 0.325 | asparagine synthase B |
| contig_23551_6 | 0.6378 | 1 | 1 | 0.7697 | 1 | 1 | 0.3275 | predicted protein |
| contig_28636_6 | 0.6391 | 1 | 1 | 0.7705 | 1 | 1 | 0.3279 | Small nuclear ribonucleoprotein G |
| contig_22874_2 | 0.6397 | 1 | 1 | 0.7705 | 1 | 1 | 0.3279 | inositol 5-phosphatase |
| contig_5361_1 | 0.6399 | 1 | 1 | 0.7705 | 1 | 1 | 0.3279 | ---NA--- |
| contig_23369_5 | 0.6408 | 1 | 1 | 0.7707 | 1 | 1 | 0.328 | predicted protein |
| contig_24715_2 | 0.6412 | 1 | 1 | 0.7707 | 1 | 1 | 0.328 | predicted protein |

| | | | | | | | | |
|---|---|---|---|---|---|---|---|---|
| contig_22602_4 | 0.6421 | 1 | 1 | 0.7713 | 1 | 1 | 0.3282 | putative all-trans-retinol 13 14-reductase |
| contig_8945_4 | 0.6439 | 1 | 1 | 0.7727 | 1 | 1 | 0.3288 | delta-9 acyl-desaturase |
| contig_27634_6 | 0.647 | 1 | 1 | 0.7758 | 1 | 1 | 0.3301 | eukaryotic translation initiation factor 3 subunit E |
| contig_30620_5 | 0.6474 | 1 | 1 | 0.7758 | 1 | 1 | 0.3301 | Transmembrane protein 87B |
| contig_19964_6 | 0.6482 | 1 | 1 | 0.7761 | 1 | 1 | 0.3303 | predicted protein |
| contig_24114_3 | 0.6507 | 1 | 1 | 0.7785 | 1 | 1 | 0.3313 | predicted protein |
| contig_22675_1 | 0.6525 | 1 | 1 | 0.7801 | 1 | 1 | 0.3319 | predicted protein |
| contig_13656_4 | 0.6544 | 1 | 1 | 0.7813 | 1 | 1 | 0.3325 | predicted protein |
| contig_40102_3 | 0.6552 | 1 | 1 | 0.7813 | 1 | 1 | 0.3325 | predicted protein |
| contig_36096_1 | 0.6554 | 1 | 1 | 0.7813 | 1 | 1 | 0.3325 | predicted protein |
| contig_11453_6 | 0.6556 | 1 | 1 | 0.7813 | 1 | 1 | 0.3325 | major facilitator transporter |
| contig_22967_5 | 0.6594 | 1 | 1 | 0.7852 | 1 | 1 | 0.3341 | predicted protein |
| contig_38768_2 | 0.6609 | 1 | 1 | 0.7853 | 1 | 1 | 0.3342 | ---NA--- |
| contig_16980_6 | 0.6611 | 1 | 1 | 0.7853 | 1 | 1 | 0.3342 | translation initiation factor 3 subunit A |
| contig_36080_5 | 0.6612 | 1 | 1 | 0.7853 | 1 | 1 | 0.3342 | predicted protein |
| contig_18485_3 | 0.6618 | 1 | 1 | 0.7853 | 1 | 1 | 0.3342 | Mitochondrial-processing peptidase subunit beta |
| contig_20147_4 | 0.6621 | 1 | 1 | 0.7853 | 1 | 1 | 0.3342 | fucoxanthin chlorophyl a/c protein |
| contig_6257_1 | 0.6633 | 1 | 1 | 0.7858 | 1 | 1 | 0.3344 | predicted protein |
| contig_24876_6 | 0.6635 | 1 | 1 | 0.7858 | 1 | 1 | 0.3344 | predicted protein |
| contig_25833_1 | 0.6644 | 1 | 1 | 0.7863 | 1 | 1 | 0.3346 | predicted protein |
| contig_23311_3 | 0.6657 | 1 | 1 | 0.7871 | 1 | 1 | 0.3349 | predicted protein |
| contig_39062_4 | 0.667 | 1 | 1 | 0.7877 | 1 | 1 | 0.3352 | 2-isopropylmalate synthase |
| contig_40291_3 | 0.6674 | 1 | 1 | 0.7877 | 1 | 1 | 0.3352 | predicted protein |
| contig_28722_5 | 0.6677 | 1 | 1 | 0.7877 | 1 | 1 | 0.3352 | predicted protein |
| contig_10657_4 | 0.6692 | 1 | 1 | 0.7888 | 1 | 1 | 0.3357 | atp-dependent RNA helicase |
| contig_38608_1 | 0.6707 | 1 | 1 | 0.79 | 1 | 1 | 0.3362 | cytochrome 450 |
| contig_11320_6 | 0.6714 | 1 | 1 | 0.7902 | 1 | 1 | 0.3363 | predicted protein |

| | | | | | | | | |
|---|---|---|---|---|---|---|---|---|
| contig_38881_1 | 0.6743 | 1 | 1 | 0.7912 | 1 | 1 | 0.3367 | SulP family inorganic anion transporter |
| contig_9982_6 | 0.6744 | 1 | 1 | 0.7912 | 1 | 1 | 0.3367 | light harvesting protein |
| contig_33127_4 | 0.6745 | 1 | 1 | 0.7912 | 1 | 1 | 0.3367 | Not1-domain-containing protein |
| contig_27473_6 | 0.6747 | 1 | 1 | 0.7912 | 1 | 1 | 0.3367 | peptidylprolyl isomerase |
| contig_36065_2 | 0.6752 | 1 | 1 | 0.7912 | 1 | 1 | 0.3367 | Translation elongation factor IF5A |
| contig_7288_5 | 0.6755 | 1 | 1 | 0.7912 | 1 | 1 | 0.3367 | protein fucoxanthin chlorophyll a/c protein |
| contig_15864_3 | 0.6759 | 1 | 1 | 0.7912 | 1 | 1 | 0.3367 | 50S ribosomal protein L15 |
| contig_11186_4 | 0.6789 | 1 | 1 | 0.7936 | 1 | 1 | 0.3377 | predicted protein |
| contig_7506_5 | 0.679 | 1 | 1 | 0.7936 | 1 | 1 | 0.3377 | apocytochrome b (mitochondrion) |
| contig_6079_5 | 0.6797 | 1 | 1 | 0.7938 | 1 | 1 | 0.3378 | predicted protein |
| contig_24815_3 | 0.6834 | 1 | 1 | 0.7976 | 1 | 1 | 0.3394 | MFS transporter |
| contig_25647_1 | 0.6845 | 1 | 1 | 0.7982 | 1 | 1 | 0.3397 | putative all-trans-retinol 13 14-reductase |
| contig_9755_4 | 0.6872 | 1 | 1 | 0.8005 | 1 | 1 | 0.3407 | valyl-tRNA synthetase |
| contig_25515_1 | 0.6875 | 1 | 1 | 0.8005 | 1 | 1 | 0.3407 | cysteine desulfurase |
| contig_34200_5 | 0.6883 | 1 | 1 | 0.8007 | 1 | 1 | 0.3407 | fructose-1 6-bisphosphatase |
| contig_13952_4 | 0.6887 | 1 | 1 | 0.8007 | 1 | 1 | 0.3407 | predicted protein |
| contig_30013_1 | 0.6896 | 1 | 1 | 0.8011 | 1 | 1 | 0.3409 | ribonucleoside-diphosphate reductase subunit alpha |
| contig_8378_3 | 0.6907 | 1 | 1 | 0.8017 | 1 | 1 | 0.3412 | phosphoglyceromutase |
| contig_39705_5 | 0.6924 | 1 | 1 | 0.8032 | 1 | 1 | 0.3418 | inorganic pyrophosphatase |
| contig_29898_4 | 0.6935 | 1 | 1 | 0.8038 | 1 | 1 | 0.342 | mitochondrial 2-oxoglutarate/malate carrier protein |
| contig_13497_3 | 0.6943 | 1 | 1 | 0.8038 | 1 | 1 | 0.342 | predicted protein |
| contig_22256_3 | 0.6946 | 1 | 1 | 0.8038 | 1 | 1 | 0.342 | 30S ribosomal protein S15 |
| contig_6643_5 | 0.6951 | 1 | 1 | 0.8038 | 1 | 1 | 0.342 | phosphoglycolate phosphatase |
| contig_38073_1 | 0.6961 | 1 | 1 | 0.8043 | 1 | 1 | 0.3422 | ---NA--- |
| contig_33904_2 | 0.6966 | 1 | 1 | 0.8043 | 1 | 1 | 0.3422 | pectate lyase |
| contig_29914_2 | 0.6977 | 1 | 1 | 0.805 | 1 | 1 | 0.3426 | predicted protein |
| contig_25927_6 | 0.6995 | 1 | 1 | 0.8061 | 1 | 1 | 0.343 | light harvesting protein |

| | | | | | | | | |
|---|---|---|---|---|---|---|---|---|
| contig_25458_4 | 0.7005 | 1 | 1 | 0.8061 | 1 | 1 | 0.343 | succinate dehydrogenase [ubiquinone] iron-sulfur subunit mitochondrial |
| contig_9641_3 | 0.7005 | 1 | 1 | 0.8061 | 1 | 1 | 0.343 | predicted protein |
| contig_22713_2 | 0.7008 | 1 | 1 | 0.8061 | 1 | 1 | 0.343 | predicted protein |
| contig_6474_4 | 0.7024 | 1 | 1 | 0.8073 | 1 | 1 | 0.3436 | FAD-dependent monooxygenase |
| contig_36054_1 | 0.7033 | 1 | 1 | 0.8078 | 1 | 1 | 0.3437 | protein fucoxanthin chlorophyll a/c protein |
| contig_27545_4 | 0.7059 | 1 | 1 | 0.8102 | 1 | 1 | 0.3448 | predicted protein |
| contig_14974_2 | 0.7071 | 1 | 1 | 0.8109 | 1 | 1 | 0.3451 | dnaj subfamily b member 4 |
| contig_35046_3 | 0.7081 | 1 | 1 | 0.8115 | 1 | 1 | 0.3453 | CK1/CK1/CK1-D protein kinase |
| contig_32515_5 | 0.7134 | 1 | 1 | 0.8169 | 1 | 1 | 0.3476 | phosphoenolpyruvate synthase |
| contig_7406_5 | 0.7174 | 1 | 1 | 0.8208 | 1 | 1 | 0.3493 | predicted protein |
| contig_30128_6 | 0.7194 | 1 | 1 | 0.8225 | 1 | 1 | 0.35 | predicted protein |
| contig_34277_5 | 0.7201 | 1 | 1 | 0.8227 | 1 | 1 | 0.3501 | 2-isopropylmalate synthase |
| contig_25676_6 | 0.7266 | 1 | 1 | 0.8296 | 1 | 1 | 0.353 | predicted protein |
| contig_7732_1 | 0.7272 | 1 | 1 | 0.8296 | 1 | 1 | 0.353 | predicted protein |
| contig_15379_5 | 0.7292 | 1 | 1 | 0.8312 | 1 | 1 | 0.3537 | nucleolar gtp-binding protein 1 |
| contig_25106_6 | 0.7303 | 1 | 1 | 0.8317 | 1 | 1 | 0.3539 | predicted protein |
| contig_6576_3 | 0.7307 | 1 | 1 | 0.8317 | 1 | 1 | 0.3539 | protein COFACTOR ASSEMBLY OF COMPLEX C SUBUNIT B CCB1 chloroplastic |
| contig_5545_2 | 0.7324 | 1 | 1 | 0.8326 | 1 | 1 | 0.3543 | NarK/NasA family nitrate transporter |
| contig_30420_3 | 0.7331 | 1 | 1 | 0.8326 | 1 | 1 | 0.3543 | predicted protein |
| contig_14343_6 | 0.7336 | 1 | 1 | 0.8326 | 1 | 1 | 0.3543 | riboflavin synthase alpha subunit |
| contig_26089_5 | 0.7341 | 1 | 1 | 0.8326 | 1 | 1 | 0.3543 | exportin-like protein |
| contig_15603_6 | 0.7347 | 1 | 1 | 0.8326 | 1 | 1 | 0.3543 | predicted protein |
| contig_10808_6 | 0.7347 | 1 | 1 | 0.8326 | 1 | 1 | 0.3543 | predicted protein |
| contig_5980_5 | 0.7412 | 1 | 1 | 0.839 | 1 | 1 | 0.357 | sensor kinase protein 3 |
| contig_38591_1 | 0.7415 | 1 | 1 | 0.839 | 1 | 1 | 0.357 | 40S ribosomal protein S18 |
| contig_35543_6 | 0.7503 | 1 | 1 | 0.8483 | 1 | 1 | 0.361 | ferredoxin--NADP+ reductase |

| | | | | | | | | |
|---|---|---|---|---|---|---|---|---|
| contig_7536_5 | 0.752 | 1 | 1 | 0.84 96 | 1 | 1 | 0.36 15 | predicted protein |
| contig_25375_ 6 | 0.7528 | 1 | 1 | 0.84 99 | 1 | 1 | 0.36 17 | predicted protein |
| contig_5563_5 | 0.755 | 1 | 1 | 0.85 17 | 1 | 1 | 0.36 24 | predicted protein |
| contig_26646_ 2 | 0.7582 | 1 | 1 | 0.85 47 | 1 | 1 | 0.36 37 | short-chain dehydrogenase/reductase acting with NAD or NADP as acceptor |
| contig_28322_ 4 | 0.7626 | 1 | 1 | 0.85 91 | 1 | 1 | 0.36 56 | farnesyl pyrophosphate synthase |
| contig_14977_ 5 | 0.7635 | 1 | 1 | 0.85 94 | 1 | 1 | 0.36 57 | predicted protein |
| contig_25573_ 4 | 0.764 | 1 | 1 | 0.85 94 | 1 | 1 | 0.36 57 | predicted protein |
| contig_14595_ 4 | 0.7651 | 1 | 1 | 0.86 | 1 | 1 | 0.36 6 | predicted protein |
| contig_8557_5 | 0.7666 | 1 | 1 | 0.86 1 | 1 | 1 | 0.36 64 | cytochrome b6 (chloroplast) |
| contig_14784_ 5 | 0.7684 | 1 | 1 | 0.86 24 | 1 | 1 | 0.36 7 | atp-dependent RNA helicase |
| contig_33495_ 6 | 0.7791 | 1 | 1 | 0.87 28 | 1 | 1 | 0.37 14 | triose or hexose phosphate/phosphate translocator |
| contig_23306_ 1 | 0.7796 | 1 | 1 | 0.87 28 | 1 | 1 | 0.37 14 | ATP:ADP antiporter AAA family |
| contig_20820_ 5 | 0.7797 | 1 | 1 | 0.87 28 | 1 | 1 | 0.37 14 | photosystem II protein D1 (chloroplast) |
| contig_13099_ 5 | 0.7799 | 1 | 1 | 0.87 28 | 1 | 1 | 0.37 14 | predicted protein |
| contig_5934_3 | 0.7817 | 1 | 1 | 0.87 4 | 1 | 1 | 0.37 19 | bifunctional aconitate hydratase 2/2-methylisocitrate dehydratase |
| contig_17493_ 5 | 0.7822 | 1 | 1 | 0.87 4 | 1 | 1 | 0.37 19 | lamin tail domain-containing protein |
| contig_9980_5 | 0.7833 | 1 | 1 | 0.87 46 | 1 | 1 | 0.37 22 | protein fucoxanthin chlorophyll a/c protein |
| contig_38336_ 2 | 0.785 | 1 | 1 | 0.87 59 | 1 | 1 | 0.37 27 | class I SAM-dependent methyltransferase |
| contig_27237_ 6 | 0.7874 | 1 | 1 | 0.87 79 | 1 | 1 | 0.37 36 | translation initiation factor 5B |
| contig_34233_ 5 | 0.7896 | 1 | 1 | 0.87 87 | 1 | 1 | 0.37 39 | predicted protein |
| contig_31150_ 6 | 0.7899 | 1 | 1 | 0.87 87 | 1 | 1 | 0.37 39 | protein fucoxanthin chlorophyll a/c protein |
| contig_10321_ 2 | 0.7905 | 1 | 1 | 0.87 87 | 1 | 1 | 0.37 39 | beta-hydroxyacyl-acyl carrier protein dehydratase (FABZ) |
| contig_23323_ 6 | 0.7906 | 1 | 1 | 0.87 87 | 1 | 1 | 0.37 39 | predicted protein |
| contig_39157_ 3 | 0.791 | 1 | 1 | 0.87 87 | 1 | 1 | 0.37 39 | predicted protein |
| contig_9226_3 | 0.793 | 1 | 1 | 0.87 93 | 1 | 1 | 0.37 42 | 2-isopropylmalate synthase |
| contig_27171_ 6 | 0.7931 | 1 | 1 | 0.87 93 | 1 | 1 | 0.37 42 | ribosome production factor 2 homolog |

| | | | | | | | | |
|---|---|---|---|---|---|---|---|---|
| contig_38019_3 | 0.7943 | 1 | 1 | 0.8793 | 1 | 1 | 0.3742 | ferredoxin--NADP+ reductase |
| contig_28535_5 | 0.7946 | 1 | 1 | 0.8793 | 1 | 1 | 0.3742 | 26S proteasome non-ATPase regulatory subunit 2 |
| contig_29765_5 | 0.7949 | 1 | 1 | 0.8793 | 1 | 1 | 0.3742 | predicted protein |
| contig_27832_2 | 0.7951 | 1 | 1 | 0.8793 | 1 | 1 | 0.3742 | ---NA--- |
| contig_25050_4 | 0.7959 | 1 | 1 | 0.8793 | 1 | 1 | 0.3742 | predicted protein |
| contig_21839_5 | 0.7962 | 1 | 1 | 0.8793 | 1 | 1 | 0.3742 | 1-deoxy-d-xylulose-5-phosphate reductoisomerase |
| contig_38571_1 | 0.7975 | 1 | 1 | 0.8796 | 1 | 1 | 0.3743 | photosystem I subunit III (chloroplast) |
| contig_39431_2 | 0.7981 | 1 | 1 | 0.8796 | 1 | 1 | 0.3743 | GDP-mannose transporter |
| contig_30481_5 | 0.7982 | 1 | 1 | 0.8796 | 1 | 1 | 0.3743 | ornithine cyclodeaminase |
| contig_26837_3 | 0.7991 | 1 | 1 | 0.8799 | 1 | 1 | 0.3744 | AP-1 complex subunit gamma-1 |
| contig_7979_4 | 0.7996 | 1 | 1 | 0.8799 | 1 | 1 | 0.3744 | protein fucoxanthin chlorophyll a/c protein |
| contig_12287_5 | 0.8032 | 1 | 1 | 0.8832 | 1 | 1 | 0.3758 | predicted protein |
| contig_19428_2 | 0.8045 | 1 | 1 | 0.8837 | 1 | 1 | 0.3761 | chloroplast beta-keto acyl reductase |
| contig_15845_2 | 0.8053 | 1 | 1 | 0.8837 | 1 | 1 | 0.3761 | predicted protein |
| contig_21581_3 | 0.8054 | 1 | 1 | 0.8837 | 1 | 1 | 0.3761 | predicted protein |
| contig_27804_5 | 0.8075 | 1 | 1 | 0.8854 | 1 | 1 | 0.3768 | Coatomer subunit alpha-1 |
| contig_33602_5 | 0.8114 | 1 | 1 | 0.889 | 1 | 1 | 0.3783 | FAD-binding domain-containing protein |
| contig_33315_3 | 0.816 | 1 | 1 | 0.8935 | 1 | 1 | 0.3802 | gametolysin peptidase M11 |
| contig_18889_1 | 0.8179 | 1 | 1 | 0.8948 | 1 | 1 | 0.3808 | ---NA--- |
| contig_9499_6 | 0.8229 | 1 | 1 | 0.8997 | 1 | 1 | 0.3828 | predicted protein |
| contig_7101_4 | 0.8236 | 1 | 1 | 0.8999 | 1 | 1 | 0.3829 | DNA-directed RNA polymerase II 19 kDa polypeptide |
| contig_17080_5 | 0.8253 | 1 | 1 | 0.8999 | 1 | 1 | 0.3829 | predicted protein |
| contig_24202_5 | 0.8257 | 1 | 1 | 0.8999 | 1 | 1 | 0.3829 | 2-isopropylmalate synthase |
| contig_12042_4 | 0.8257 | 1 | 1 | 0.8999 | 1 | 1 | 0.3829 | coproporphyrinogen III oxidase |
| contig_30519_2 | 0.826 | 1 | 1 | 0.8999 | 1 | 1 | 0.3829 | predicted protein |
| contig_3085_5 | 0.8286 | 1 | 1 | 0.902 | 1 | 1 | 0.3838 | ferredoxin (chloroplast) |
| contig_32503_1 | 0.8297 | 1 | 1 | 0.9025 | 1 | 1 | 0.3841 | predicted protein |

| | | | | | | | | |
|---|---|---|---|---|---|---|---|---|
| contig_6836_6 | 0.8303 | 1 | 1 | 0.9025 | 1 | 1 | 0.3841 | predicted protein |
| contig_17128_2 | 0.8309 | 1 | 1 | 0.9026 | 1 | 1 | 0.3841 | predicted protein |
| contig_22140_2 | 0.8331 | 1 | 1 | 0.9043 | 1 | 1 | 0.3848 | predicted protein |
| contig_33453_2 | 0.8367 | 1 | 1 | 0.9077 | 1 | 1 | 0.3862 | predicted protein |
| contig_14735_6 | 0.8375 | 1 | 1 | 0.9077 | 1 | 1 | 0.3862 | proteasome subunit beta type-5-like |
| contig_22660_5 | 0.8379 | 1 | 1 | 0.9077 | 1 | 1 | 0.3862 | predicted protein |
| contig_27847_4 | 0.8394 | 1 | 1 | 0.9086 | 1 | 1 | 0.3866 | unnamed protein product |
| contig_10682_1 | 0.8432 | 1 | 1 | 0.9118 | 1 | 1 | 0.388 | predicted protein |
| contig_6450_1 | 0.8436 | 1 | 1 | 0.9118 | 1 | 1 | 0.388 | glutamine-fructose-6-phosphate transaminase |
| contig_7847_6 | 0.8447 | 1 | 1 | 0.9122 | 1 | 1 | 0.3882 | predicted protein |
| contig_14363_5 | 0.846 | 1 | 1 | 0.9122 | 1 | 1 | 0.3882 | predicted protein |
| contig_39446_2 | 0.8471 | 1 | 1 | 0.9122 | 1 | 1 | 0.3882 | U6 snRNA-associated Sm-like protein LSm2 |
| contig_7506_4 | 0.848 | 1 | 1 | 0.9122 | 1 | 1 | 0.3882 | apocytochrome b (mitochondrion) |
| contig_36394_4 | 0.8487 | 1 | 1 | 0.9122 | 1 | 1 | 0.3882 | Fucoxanthin-chlorophyll a-c binding protein F chloroplastic |
| contig_18489_1 | 0.8501 | 1 | 1 | 0.9122 | 1 | 1 | 0.3882 | predicted protein |
| contig_23752_4 | 0.8503 | 1 | 1 | 0.9122 | 1 | 1 | 0.3882 | predicted protein |
| contig_26596_5 | 0.8504 | 1 | 1 | 0.9122 | 1 | 1 | 0.3882 | leucyl aminopeptidase |
| contig_29804_5 | 0.8508 | 1 | 1 | 0.9122 | 1 | 1 | 0.3882 | S-adenosylmethionine synthetase |
| contig_40197_2 | 0.8513 | 1 | 1 | 0.9122 | 1 | 1 | 0.3882 | protein fucoxanthin chlorophyll a/c protein |
| contig_24539_2 | 0.8518 | 1 | 1 | 0.9122 | 1 | 1 | 0.3882 | phosphomannomutase/phosphoglucomutase |
| contig_31306_6 | 0.8519 | 1 | 1 | 0.9122 | 1 | 1 | 0.3882 | predicted protein |
| contig_17633_4 | 0.8521 | 1 | 1 | 0.9122 | 1 | 1 | 0.3882 | Adenylyl cyclase-associated protein |
| contig_24223_6 | 0.8526 | 1 | 1 | 0.9122 | 1 | 1 | 0.3882 | predicted protein |
| contig_32617_4 | 0.8529 | 1 | 1 | 0.9122 | 1 | 1 | 0.3882 | VPLPA-CTERM protein sorting domain-containing protein |
| contig_9342_6 | 0.8541 | 1 | 1 | 0.9128 | 1 | 1 | 0.3884 | predicted protein |
| contig_17187_5 | 0.8549 | 1 | 1 | 0.9131 | 1 | 1 | 0.3885 | small ubiquitin-like modifier |
| contig_25573_5 | 0.8555 | 1 | 1 | 0.9131 | 1 | 1 | 0.3885 | predicted protein |

| | | | | | | | | |
|---|---|---|---|---|---|---|---|---|
| contig_11033_5 | 0.8575 | 1 | 1 | 0.9146 | 1 | 1 | 0.3892 | predicted protein |
| contig_27384_5 | 0.8594 | 1 | 1 | 0.9153 | 1 | 1 | 0.3895 | predicted protein |
| contig_33655_6 | 0.8598 | 1 | 1 | 0.9153 | 1 | 1 | 0.3895 | predicted protein |
| contig_20295_6 | 0.86 | 1 | 1 | 0.9153 | 1 | 1 | 0.3895 | cytochrome c oxidase subunit II (mitochondrion) |
| contig_10034_3 | 0.8607 | 1 | 1 | 0.9154 | 1 | 1 | 0.3895 | 60S ribosomal protein L10 |
| contig_33443_2 | 0.8637 | 1 | 1 | 0.9169 | 1 | 1 | 0.3902 | predicted protein |
| contig_34791_3 | 0.8639 | 1 | 1 | 0.9169 | 1 | 1 | 0.3902 | Regulator of chromosome condensation 1/beta-lactamase-inhibitor protein II |
| contig_30863_3 | 0.864 | 1 | 1 | 0.9169 | 1 | 1 | 0.3902 | predicted protein |
| contig_23416_6 | 0.867 | 1 | 1 | 0.9188 | 1 | 1 | 0.391 | predicted protein |
| contig_5863_5 | 0.8672 | 1 | 1 | 0.9188 | 1 | 1 | 0.391 | predicted protein |
| contig_27998_6 | 0.8683 | 1 | 1 | 0.9188 | 1 | 1 | 0.391 | 2 3-bisphosphoglycerate-dependent phosphoglycerate mutase |
| contig_40133_3 | 0.8686 | 1 | 1 | 0.9188 | 1 | 1 | 0.391 | 30S ribosomal protein S15 |
| contig_20096_5 | 0.8688 | 1 | 1 | 0.9188 | 1 | 1 | 0.391 | predicted protein |
| contig_26282_6 | 0.8707 | 1 | 1 | 0.9202 | 1 | 1 | 0.3916 | dihydrodipicolinate reductase |
| contig_18211_4 | 0.8712 | 1 | 1 | 0.9202 | 1 | 1 | 0.3916 | light harvesting protein |
| contig_6242_5 | 0.8728 | 1 | 1 | 0.9212 | 1 | 1 | 0.392 | predicted protein |
| contig_23620_5 | 0.8737 | 1 | 1 | 0.9214 | 1 | 1 | 0.3921 | nitrate transporter |
| contig_12101_5 | 0.8742 | 1 | 1 | 0.9214 | 1 | 1 | 0.3921 | ---NA--- |
| contig_38903_2 | 0.8751 | 1 | 1 | 0.9217 | 1 | 1 | 0.3922 | ---NA--- |
| contig_6146_1 | 0.8775 | 1 | 1 | 0.9224 | 1 | 1 | 0.3925 | aliphatic amidase |
| contig_14698_3 | 0.8781 | 1 | 1 | 0.9224 | 1 | 1 | 0.3925 | AP-1 complex subunit gamma-1 |
| contig_29793_4 | 0.8782 | 1 | 1 | 0.9224 | 1 | 1 | 0.3925 | predicted protein |
| contig_27823_2 | 0.8785 | 1 | 1 | 0.9224 | 1 | 1 | 0.3925 | predicted protein |
| contig_26918_5 | 0.8792 | 1 | 1 | 0.9224 | 1 | 1 | 0.3925 | predicted protein |
| contig_14126_5 | 0.8794 | 1 | 1 | 0.9224 | 1 | 1 | 0.3925 | predicted protein |
| contig_16833_4 | 0.8826 | 1 | 1 | 0.9252 | 1 | 1 | 0.3937 | 60S ribosomal protein |

| | | | | | | | | |
|---|---|---|---|---|---|---|---|---|
| contig_39533_1 | 0.8848 | 1 | 1 | 0.9268 | 1 | 1 | 0.3944 | sodium/proton antiporter 1 |
| contig_25457_5 | 0.887 | 1 | 1 | 0.9285 | 1 | 1 | 0.3951 | farnesyl-diphosphate farnesyltransferase |
| contig_33750_4 | 0.8891 | 1 | 1 | 0.93 | 1 | 1 | 0.3957 | Violaxanthin de-epoxidase |
| contig_29535_5 | 0.8901 | 1 | 1 | 0.9305 | 1 | 1 | 0.3959 | predicted protein |
| contig_27901_2 | 0.8957 | 1 | 1 | 0.9341 | 1 | 1 | 0.3975 | methionyl-trna formyltransferase |
| contig_28443_2 | 0.8959 | 1 | 1 | 0.9341 | 1 | 1 | 0.3975 | nad-dependent epimerase/dehydratase |
| contig_29713_5 | 0.8961 | 1 | 1 | 0.9341 | 1 | 1 | 0.3975 | probable prefoldin subunit 2 |
| contig_38239_2 | 0.8961 | 1 | 1 | 0.9341 | 1 | 1 | 0.3975 | predicted protein |
| contig_33784_6 | 0.8976 | 1 | 1 | 0.9346 | 1 | 1 | 0.3977 | predicted protein |
| contig_9827_1 | 0.8978 | 1 | 1 | 0.9346 | 1 | 1 | 0.3977 | predicted protein |
| contig_15516_6 | 0.9049 | 1 | 1 | 0.9411 | 1 | 1 | 0.4005 | pyrimidine/purine nucleoside phosphorylase |
| contig_38574_3 | 0.9052 | 1 | 1 | 0.9411 | 1 | 1 | 0.4005 | predicted protein |
| contig_7412_1 | 0.907 | 1 | 1 | 0.9422 | 1 | 1 | 0.4009 | E3 ubiquitin-protein ligase RING2-like |
| contig_33528_6 | 0.9087 | 1 | 1 | 0.9434 | 1 | 1 | 0.4014 | SPX domain-containing membrane protein At4g22990-like |
| contig_39210_6 | 0.9113 | 1 | 1 | 0.9454 | 1 | 1 | 0.4023 | predicted protein |
| contig_6937_2 | 0.9141 | 1 | 1 | 0.9469 | 1 | 1 | 0.4029 | predicted protein |
| contig_5926_5 | 0.9144 | 1 | 1 | 0.9469 | 1 | 1 | 0.4029 | predicted protein |
| contig_16552_4 | 0.9145 | 1 | 1 | 0.9469 | 1 | 1 | 0.4029 | predicted protein |
| contig_19373_4 | 0.9173 | 1 | 1 | 0.949 | 1 | 1 | 0.4038 | fucoxanthin chlorophyll a/c-binding protein precursor |
| contig_12026_5 | 0.9188 | 1 | 1 | 0.95 | 1 | 1 | 0.4042 | predicted protein |
| contig_7125_2 | 0.92 | 1 | 1 | 0.9506 | 1 | 1 | 0.4045 | cysteine proteinase 3-like |
| contig_28316_5 | 0.9214 | 1 | 1 | 0.9508 | 1 | 1 | 0.4046 | hypothetical protein FRACYDRAFT_234640 |
| contig_27657_5 | 0.9215 | 1 | 1 | 0.9508 | 1 | 1 | 0.4046 | solute carrier family 35 member E1 |
| contig_40960_3 | 0.9226 | 1 | 1 | 0.9512 | 1 | 1 | 0.4048 | predicted protein |
| contig_23910_2 | 0.9241 | 1 | 1 | 0.9512 | 1 | 1 | 0.4048 | ARM repeat-containing protein |
| contig_37879_1 | 0.9243 | 1 | 1 | 0.9512 | 1 | 1 | 0.4048 | UPF0187 protein |
| contig_40882_2 | 0.9243 | 1 | 1 | 0.9512 | 1 | 1 | 0.4048 | predicted protein |

| | | | | | | | | |
|---|---|---|---|---|---|---|---|---|
| contig_28215_4 | 0.9264 | 1 | 1 | 0.9526 | 1 | 1 | 0.4054 | light harvesting complex protein |
| contig_37256_3 | 0.9317 | 1 | 1 | 0.9571 | 1 | 1 | 0.4073 | histone H3 |
| contig_38536_2 | 0.9322 | 1 | 1 | 0.9571 | 1 | 1 | 0.4073 | predicted protein |
| contig_38023_2 | 0.9326 | 1 | 1 | 0.9571 | 1 | 1 | 0.4073 | predicted protein |
| contig_32601_3 | 0.9337 | 1 | 1 | 0.9576 | 1 | 1 | 0.4075 | predicted protein |
| contig_3491_1 | 0.9367 | 1 | 1 | 0.9594 | 1 | 1 | 0.4083 | predicted protein |
| contig_28995_4 | 0.9368 | 1 | 1 | 0.9594 | 1 | 1 | 0.4083 | predicted protein |
| contig_30564_1 | 0.9396 | 1 | 1 | 0.9617 | 1 | 1 | 0.4092 | predicted protein |
| contig_36999_3 | 0.9411 | 1 | 1 | 0.9626 | 1 | 1 | 0.4096 | predicted protein |
| contig_6884_5 | 0.9455 | 1 | 1 | 0.9665 | 1 | 1 | 0.4113 | putative ATP-dependent RNA helicase DHX30 isoform X3 |
| contig_7315_2 | 0.9477 | 1 | 1 | 0.968 | 1 | 1 | 0.4119 | predicted protein |
| contig_14080_4 | 0.9544 | 1 | 1 | 0.9739 | 1 | 1 | 0.4144 | dihydroorotase |
| contig_33151_3 | 0.9549 | 1 | 1 | 0.9739 | 1 | 1 | 0.4144 | acetylglutamate kinase |
| contig_25696_5 | 0.9555 | 1 | 1 | 0.9739 | 1 | 1 | 0.4144 | predicted protein |
| contig_36050_1 | 0.956 | 1 | 1 | 0.9739 | 1 | 1 | 0.4144 | Light-harvesting complex I LH38 protein |
| contig_26114_2 | 0.9584 | 1 | 1 | 0.9755 | 1 | 1 | 0.4151 | ATP phosphoribosyltransferase |
| contig_26198_6 | 0.9597 | 1 | 1 | 0.9755 | 1 | 1 | 0.4151 | ribosomal protein L20 |
| contig_35135_5 | 0.9597 | 1 | 1 | 0.9755 | 1 | 1 | 0.4151 | peptidyl-prolyl cis-trans isomerase A |
| contig_34378_5 | 0.9602 | 1 | 1 | 0.9755 | 1 | 1 | 0.4151 | predicted protein |
| contig_30869_3 | 0.9608 | 1 | 1 | 0.9755 | 1 | 1 | 0.4151 | predicted protein |
| contig_36079_3 | 0.9614 | 1 | 1 | 0.9755 | 1 | 1 | 0.4151 | photosystem II PsbU protein |
| contig_10491_6 | 0.9654 | 1 | 1 | 0.9782 | 1 | 1 | 0.4163 | glutamine synthetase catalytic domain protein protein |
| contig_29342_2 | 0.9666 | 1 | 1 | 0.9782 | 1 | 1 | 0.4163 | leukotriene A-4 hydrolase/aminopeptidase |
| contig_30461_3 | 0.9669 | 1 | 1 | 0.9782 | 1 | 1 | 0.4163 | 6 7-dimethyl-8-ribityllumazine synthase |
| contig_19158_3 | 0.9672 | 1 | 1 | 0.9782 | 1 | 1 | 0.4163 | small ubiquitin-like modifier (SUMO) |
| contig_34883_3 | 0.9673 | 1 | 1 | 0.9782 | 1 | 1 | 0.4163 | Fructose-bisphosphate aldolase |
| contig_16053_5 | 0.9686 | 1 | 1 | 0.9789 | 1 | 1 | 0.4165 | predicted protein |

| | | | | | | | | |
|---|---|---|---|---|---|---|---|---|
| contig_16841_6 | 0.9702 | 1 | 1 | 0.9794 | 1 | 1 | 0.4168 | predicted protein |
| contig_15496_6 | 0.9704 | 1 | 1 | 0.9794 | 1 | 1 | 0.4168 | catalase |
| contig_10162_6 | 0.9728 | 1 | 1 | 0.9811 | 1 | 1 | 0.4175 | predicted protein |
| contig_9256_6 | 0.9749 | 1 | 1 | 0.9822 | 1 | 1 | 0.418 | predicted protein |
| contig_40017_2 | 0.9751 | 1 | 1 | 0.9822 | 1 | 1 | 0.418 | predicted protein |
| contig_37905_5 | 0.9769 | 1 | 1 | 0.9831 | 1 | 1 | 0.4183 | ---NA--- |
| contig_21568_5 | 0.9773 | 1 | 1 | 0.9831 | 1 | 1 | 0.4183 | predicted protein |
| contig_30516_6 | 0.9786 | 1 | 1 | 0.9831 | 1 | 1 | 0.4183 | aspartate--ammonia ligase |
| contig_25871_6 | 0.9786 | 1 | 1 | 0.9831 | 1 | 1 | 0.4183 | RNA-binding domain-containing protein |
| contig_28732_1 | 0.98 | 1 | 1 | 0.9839 | 1 | 1 | 0.4187 | light harvesting protein |
| contig_15445_5 | 0.9891 | 1 | 1 | 0.9924 | 1 | 1 | 0.4223 | zeaxanthin epoxidase |
| contig_35278_1 | 0.991 | 1 | 1 | 0.9936 | 1 | 1 | 0.4228 | light harvesting protein |
| contig_10138_6 | 0.9941 | 1 | 1 | 0.9961 | 1 | 1 | 0.4239 | adenylate kinase chloroplastic |
| contig_28061_5 | 0.9975 | 1 | 1 | 0.9979 | 1 | 1 | 0.4246 | apospory-associated protein C |
| contig_33083_6 | 0.9978 | 1 | 1 | 0.9979 | 1 | 1 | 0.4246 | Glucose-6-phosphate isomerase cytosolic |
| contig_25855_2 | 0.9979 | 1 | 1 | 0.9979 | 0.9979 | 1 | 0.4246 | protein transport protein Sec61 subunit gamma-like |

## Nt25 – Tt34

| proteinAccession | ttest_pvals | Binomial_SGoF | Conservative_SGoF | BH | SFisher | Bonferroni | q_val | Description |
|---|---|---|---|---|---|---|---|---|
| contig_6708_5 | 0 | 0 | 2E-04 | 0 | 0 | 0 | 0 | hemagglutinin/amebocyte aggregation factor |
| contig_5982_4 | 0 | 1E-04 | 2E-04 | 0.0083 | 0 | 0.0219 | 0.0037 | abnormal spindle-like microcephaly-associated protein homolog |
| contig_32146_2 | 0 | 1E-04 | 2E-04 | 0.0083 | 0 | 0.0249 | 0.0037 | silicon transporter |
| contig_15786_6 | 1E-04 | 1E-04 | 3E-04 | 0.0155 | 0 | 0.0821 | 0.0068 | TIGR00297 family protein |
| contig_24655_5 | 1E-04 | 1E-04 | 3E-04 | 0.0155 | 0 | 0.0827 | 0.0068 | predicted protein |
| contig_14853_4 | 1E-04 | 1E-04 | 3E-04 | 0.0155 | 0 | 0.1162 | 0.0068 | peptidyl-prolyl cis-trans isomerase FKBP19 chloroplastic |
| contig_24967_5 | 1E-04 | 1E-04 | 3E-04 | 0.0155 | 0 | 0.1411 | 0.0068 | PGR5-like protein 1A chloroplastic |
| contig_22589_6 | 1E-04 | 1E-04 | 3E-04 | 0.0155 | 0 | 0.1457 | 0.0068 | Peptidase_M16-domain-containing protein |

| | | | | | | | | |
|---|---|---|---|---|---|---|---|---|
| contig_6709_6 | 1E-04 | 1E-04 | 3E-04 | 0.0155 | 0 | 0.1733 | 0.0068 | predicted protein |
| contig_28171_1 | 1E-04 | 1E-04 | 4E-04 | 0.0155 | 0 | 0.1767 | 0.0068 | predicted protein |
| contig_13715_6 | 1E-04 | 2E-04 | 4E-04 | 0.0155 | 0 | 0.1772 | 0.0068 | dnaJ homolog subfamily B member 6 isoform X2 |
| contig_25730_2 | 1E-04 | 2E-04 | 4E-04 | 0.0155 | 0 | 0.1846 | 0.0068 | predicted protein |
| contig_26303_6 | 1E-04 | 2E-04 | 4E-04 | 0.0174 | 0 | 0.225 | 0.0077 | DEAD-domain-containing protein |
| contig_13757_6 | 2E-04 | 3E-04 | 5E-04 | 0.0191 | 0 | 0.2783 | 0.0084 | predicted protein |
| contig_28781_6 | 2E-04 | 3E-04 | 5E-04 | 0.0191 | 0 | 0.2838 | 0.0084 | heat shock protein 90 |
| contig_11304_3 | 2E-04 | 3E-04 | 5E-04 | 0.0197 | 0 | 0.3188 | 0.0087 | predicted protein |
| contig_30060_4 | 2E-04 | 3E-04 | 6E-04 | 0.0197 | 0 | 0.3696 | 0.0087 | nitrite transporter NAR1 |
| contig_39519_2 | 3E-04 | 3E-04 | 6E-04 | 0.0197 | 0 | 0.377 | 0.0087 | predicted protein |
| contig_34524_6 | 3E-04 | 3E-04 | 7E-04 | 0.0197 | 0 | 0.4197 | 0.0087 | ---NA--- |
| contig_15405_3 | 3E-04 | 3E-04 | 7E-04 | 0.0197 | 0 | 0.4239 | 0.0087 | predicted protein |
| contig_29723_5 | 3E-04 | 3E-04 | 7E-04 | 0.0197 | 0 | 0.4275 | 0.0087 | predicted protein |
| contig_37567_5 | 3E-04 | 3E-04 | 7E-04 | 0.0197 | 0 | 0.4541 | 0.0087 | ATP synthase gamma subunit |
| contig_29346_4 | 3E-04 | 4E-04 | 8E-04 | 0.0197 | 0 | 0.4649 | 0.0087 | homogentisate solanesyltransferase |
| contig_20441_2 | 3E-04 | 4E-04 | 8E-04 | 0.0197 | 0 | 0.4655 | 0.0087 | putative glyoxalase I |
| contig_24784_5 | 3E-04 | 4E-04 | 0.001 | 0.02 | 0 | 0.4929 | 0.0088 | DUF423-domain-containing protein |
| contig_21794_2 | 3E-04 | 4E-04 | 0.001 | 0.0201 | 0 | 0.5154 | 0.0089 | fucoxanthin chlorophyll a/c protein deviant |
| contig_32615_1 | 4E-04 | 5E-04 | 0.001 | 0.0201 | 0 | 0.5337 | 0.0089 | predicted protein |
| contig_23988_3 | 4E-04 | 5E-04 | 0.0013 | 0.0213 | 0 | 0.5857 | 0.0094 | predicted protein |
| contig_30367_5 | 4E-04 | 5E-04 | 0.0013 | 0.0213 | 0 | 0.6061 | 0.0094 | protein fucoxanthin chlorophyll a/c protein |
| contig_22744_5 | 4E-04 | 5E-04 | 0.0013 | 0.0217 | 0 | 0.6471 | 0.0095 | predicted protein |
| contig_12144_4 | 4E-04 | 5E-04 | 0.0014 | 0.0217 | 0 | 0.6582 | 0.0095 | FAD-binding domain-containing protein |
| contig_21620_6 | 5E-04 | 6E-04 | 0.0014 | 0.0218 | 0 | 0.697 | 0.0096 | predicted protein |
| contig_28501_6 | 5E-04 | 6E-04 | 0.0015 | 0.0218 | 0 | 0.7178 | 0.0096 | predicted protein |
| contig_27073_1 | 5E-04 | 7E-04 | 0.0015 | 0.0218 | 0 | 0.734 | 0.0096 | Phospholipid hydroperoxide glutathione peroxidase |
| contig_33184_2 | 5E-04 | 7E-04 | 0.0015 | 0.0218 | 0 | 0.7448 | 0.0096 | predicted protein |

| | | | | | | | | |
|---|---|---|---|---|---|---|---|---|
| contig_33369_2 | 5E-04 | 7E-04 | 0.0017 | 0.0223 | 0 | 0.7832 | 0.0098 | F0F1 ATP synthase subunit gamma |
| contig_23510_1 | 6E-04 | 7E-04 | 0.0017 | 0.0245 | 0 | 0.8868 | 0.0108 | predicted protein |
| contig_24254_6 | 6E-04 | 8E-04 | 0.0018 | 0.0248 | 0 | 0.9307 | 0.011 | predicted protein |
| contig_6795_4 | 7E-04 | 8E-04 | 0.0018 | 0.0248 | 0 | 0.9641 | 0.011 | WSC domain protein putative |
| contig_6069_2 | 7E-04 | 8E-04 | 0.0019 | 0.0248 | 0 | 0.9681 | 0.011 | GLNA glutamine synthase |
| contig_32573_3 | 7E-04 | 0.001 | 0.0019 | 0.0256 | 0 | 1 | 0.0113 | ATP synthase CF1 alpha subunit (chloroplast) |
| contig_28430_6 | 7E-04 | 0.001 | 0.0022 | 0.0264 | 0 | 1 | 0.0116 | predicted protein |
| contig_38925_2 | 7E-04 | 0.0013 | 0.0023 | 0.0264 | 0 | 1 | 0.0116 | ---NA--- |
| contig_16024_6 | 8E-04 | 0.0013 | 0.0023 | 0.028 | 0 | 1 | 0.0123 | Chaperonin GroL |
| contig_29863_5 | 8E-04 | 0.0013 | 0.0024 | 0.028 | 0 | 1 | 0.0123 | Homoserine dehydrogenase [ |
| contig_27387_6 | 8E-04 | 0.0013 | 0.0024 | 0.028 | 0 | 1 | 0.0123 | GTP-binding protein YPTC1 variant |
| contig_6911_6 | 0.001 | 0.0014 | 0.0025 | 0.0309 | 0 | 1 | 0.0136 | pyrophosphate-dependent phosphofructose kinase |
| contig_6700_4 | 0.001 | 0.0015 | 0.0025 | 0.0317 | 0 | 1 | 0.014 | agmatinase |
| contig_34416_5 | 0.001 | 0.0015 | 0.0026 | 0.0322 | 0 | 1 | 0.0142 | predicted protein |
| contig_21753_2 | 0.0013 | 0.0015 | 0.0026 | 0.0377 | 0 | 1 | 0.0166 | glyceraldehyde-3-phosphate dehydrogenase |
| contig_13811_4 | 0.0013 | 0.0015 | 0.0027 | 0.0377 | 0 | 1 | 0.0166 | fucoxanthin chlorophyll a/c protein 8 |
| contig_33607_5 | 0.0013 | 0.0017 | 0.0029 | 0.0377 | 0 | 1 | 0.0166 | S-adenosyl-L-methionine-dependent methyltransferase |
| contig_9050_5 | 0.0013 | 0.0017 | 0.003 | 0.0383 | 0 | 1 | 0.0169 | triose phosphate/phosphate translocator |
| contig_29365_1 | 0.0014 | 0.0017 | 0.003 | 0.0383 | 0 | 1 | 0.0169 | predicted protein |
| contig_27069_6 | 0.0014 | 0.0018 | 0.003 | 0.0383 | 0 | 1 | 0.0169 | predicted protein |
| contig_22939_2 | 0.0015 | 0.0019 | 0.0031 | 0.0388 | 0 | 1 | 0.0171 | DNA damage-binding protein 1 |
| contig_6179_6 | 0.0015 | 0.0019 | 0.0031 | 0.0388 | 0 | 1 | 0.0171 | Long-chain-fatty-acid--CoA ligase putative |
| contig_30298_4 | 0.0015 | 0.002 | 0.0032 | 0.0388 | 0 | 1 | 0.0171 | predicted protein |
| contig_12148_3 | 0.0015 | 0.0022 | 0.0032 | 0.0388 | 0 | 1 | 0.0171 | predicted protein |
| contig_12511_6 | 0.0017 | 0.0023 | 0.0033 | 0.0423 | 0 | 1 | 0.0187 | predicted protein |
| contig_6836_6 | 0.0017 | 0.0024 | 0.0033 | 0.0423 | 0 | 1 | 0.0187 | predicted protein |
| contig_7129_5 | 0.0017 | 0.0024 | 0.0033 | 0.0423 | 0 | 1 | 0.0187 | T9SS type A sorting domain-containing protein |

| | | | | | | | | |
|---|---|---|---|---|---|---|---|---|
| contig_16388_6 | 0.0018 | 0.0024 | 0.0034 | 0.0434 | 0 | 1 | 0.0191 | predicted protein |
| contig_8232_1 | 0.0018 | 0.0025 | 0.0037 | 0.0435 | 0 | 1 | 0.0192 | alanine--glyoxylate aminotransferase family protein |
| contig_30765_6 | 0.0019 | 0.0025 | 0.0037 | 0.0441 | 0 | 1 | 0.0194 | dynactin subunit 1 |
| contig_23318_6 | 0.0019 | 0.0026 | 0.0038 | 0.0447 | 0 | 1 | 0.0197 | putative LOV domain-containing protein |
| contig_23093_4 | 0.002 | 0.0027 | 0.0038 | 0.0453 | 0 | 1 | 0.02 | predicted protein |
| contig_23912_3 | 0.0022 | 0.0027 | 0.004 | 0.0483 | 0 | 1 | 0.0213 | predicted protein |
| contig_22850_1 | 0.0023 | 0.0029 | 0.004 | 0.0504 | 0 | 1 | 0.0222 | nitrate transporter |
| contig_27313_6 | 0.0023 | 0.003 | 0.004 | 0.0507 | 0 | 1 | 0.0224 | predicted protein |
| contig_17463_2 | 0.0024 | 0.003 | 0.004 | 0.0507 | 0 | 1 | 0.0224 | glucose-6-phosphate isomerase |
| contig_28094_2 | 0.0024 | 0.003 | 0.0042 | 0.0507 | 0 | 1 | 0.0224 | agmatinase |
| contig_20792_5 | 0.0024 | 0.0031 | 0.0042 | 0.0508 | 0 | 1 | 0.0224 | ---NA--- |
| contig_27826_5 | 0.0025 | 0.0032 | 0.0043 | 0.0508 | 0 | 1 | 0.0224 | predicted protein |
| contig_7380_4 | 0.0025 | 0.0032 | 0.0043 | 0.0508 | 0 | 1 | 0.0224 | predicted protein |
| contig_6050_1 | 0.0026 | 0.0032 | 0.0043 | 0.0517 | 0 | 1 | 0.0228 | heat shock protein 70 (chloroplast) |
| contig_14326_6 | 0.0026 | 0.0033 | 0.0044 | 0.0519 | 0 | 1 | 0.0229 | peroxiredoxin |
| contig_27112_5 | 0.0027 | 0.0033 | 0.0044 | 0.0529 | 0 | 1 | 0.0233 | calreticulin |
| contig_27994_5 | 0.0027 | 0.0033 | 0.0046 | 0.0529 | 0 | 1 | 0.0233 | nitrate transporter |
| contig_35584_4 | 0.0029 | 0.0033 | 0.0046 | 0.0546 | 0 | 1 | 0.0241 | nitrate transporter |
| contig_37059_1 | 0.003 | 0.0034 | 0.0046 | 0.0548 | 0 | 1 | 0.0242 | Ura6p |
| contig_30052_6 | 0.003 | 0.0037 | 0.0047 | 0.0548 | 0 | 1 | 0.0242 | predicted protein |
| contig_35362_1 | 0.003 | 0.0037 | 0.0047 | 0.0548 | 0 | 1 | 0.0242 | nitrate transporter |
| contig_22539_3 | 0.0031 | 0.0038 | 0.0048 | 0.0551 | 0 | 1 | 0.0243 | sulfate adenylyltransferase |
| contig_22875_4 | 0.0031 | 0.0038 | 0.0048 | 0.0551 | 0 | 1 | 0.0243 | calcium-translocating P-type ATPase PMCA-type |
| contig_38406_6 | 0.0032 | 0.004 | 0.0051 | 0.0551 | 0 | 1 | 0.0243 | predicted protein |
| contig_24766_6 | 0.0032 | 0.004 | 0.0052 | 0.0551 | 0 | 1 | 0.0243 | nitrate reductase |
| contig_15866_5 | 0.0032 | 0.004 | 0.0053 | 0.0551 | 0 | 1 | 0.0243 | hypothetical protein THAOC_02924 |
| contig_30865_4 | 0.0033 | 0.0042 | 0.0053 | 0.0552 | 0 | 1 | 0.0243 | 2-oxo acid dehydrogenase subunit E2 |

| | | | | | | | | |
|---|---|---|---|---|---|---|---|---|
| contig_24606_4 | 0.0033 | 0.0042 | 0.0055 | 0.0552 | 0 | 1 | 0.0243 | beta-ketoacyl-ACP synthase I |
| contig_30091_6 | 0.0033 | 0.0042 | 0.0055 | 0.0552 | 0 | 1 | 0.0243 | predicted protein |
| contig_10583_5 | 0.0033 | 0.0043 | 0.0058 | 0.0552 | 0 | 1 | 0.0243 | photosystem II stability/assembly factor HCF136 |
| contig_27341_1 | 0.0034 | 0.0043 | 0.006 | 0.0553 | 0 | 1 | 0.0244 | protein patched homolog 2 |
| contig_24874_1 | 0.0037 | 0.0043 | 0.0062 | 0.0591 | 0 | 1 | 0.026 | predicted protein |
| contig_28347_6 | 0.0037 | 0.0044 | 0.0062 | 0.0591 | 0 | 1 | 0.026 | DEAD-box ATP-dependent RNA helicase 8 |
| contig_17096_5 | 0.0037 | 0.0044 | 0.0063 | 0.0591 | 0 | 1 | 0.026 | predicted protein |
| contig_9471_3 | 0.0038 | 0.0044 | 0.0067 | 0.0594 | 0 | 1 | 0.0262 | photosystem II CP43 chlorophyll apoprotein (chloroplast) |
| contig_10388_3 | 0.0038 | 0.0046 | 0.0067 | 0.0594 | 0 | 1 | 0.0262 | light-harvesting protein |
| contig_12115_3 | 0.004 | 0.0047 | 0.0067 | 0.0599 | 0 | 1 | 0.0264 | predicted protein |
| contig_34750_5 | 0.004 | 0.0047 | 0.0067 | 0.0599 | 0 | 1 | 0.0264 | thioredoxin-disulfide reductase |
| contig_6151_5 | 0.004 | 0.0047 | 0.0069 | 0.0599 | 0 | 1 | 0.0264 | CD209 antigen-like protein 2 isoform X1 |
| contig_33257_5 | 0.004 | 0.0047 | 0.0071 | 0.0599 | 0 | 1 | 0.0264 | SulP family inorganic anion transporter |
| contig_36067_5 | 0.0041 | 0.0047 | 0.0071 | 0.0599 | 0 | 1 | 0.0264 | silicon transporter |
| contig_30185_1 | 0.0042 | 0.0048 | 0.0073 | 0.0604 | 0 | 1 | 0.0266 | signal peptidase I |
| contig_30486_2 | 0.0042 | 0.0052 | 0.0076 | 0.0604 | 0 | 1 | 0.0266 | signal peptidase I |
| contig_19340_6 | 0.0043 | 0.0053 | 0.0077 | 0.0604 | 0 | 1 | 0.0266 | acetyl-coa carboxylase |
| contig_39521_4 | 0.0043 | 0.0053 | 0.0077 | 0.0604 | 0 | 1 | 0.0266 | predicted protein |
| contig_29484_4 | 0.0043 | 0.0053 | 0.0077 | 0.0604 | 0 | 1 | 0.0266 | serine hydroxymethyltransferase |
| contig_6525_2 | 0.0044 | 0.0055 | 0.0078 | 0.0604 | 0 | 1 | 0.0266 | probable aminopeptidase NPEPL1 |
| contig_29420_3 | 0.0044 | 0.0057 | 0.0078 | 0.0604 | 0 | 1 | 0.0266 | thylakoid lumenal 15.0 kDa protein 2 chloroplastic |
| contig_6613_6 | 0.0044 | 0.0058 | 0.008 | 0.0604 | 0 | 1 | 0.0266 | sodium-coupled neutral amino acid transporter |
| contig_30021_2 | 0.0046 | 0.0062 | 0.0081 | 0.0612 | 0 | 1 | 0.027 | predicted protein |
| contig_7261_5 | 0.0046 | 0.0062 | 0.0083 | 0.0612 | 0 | 1 | 0.027 | ubiquitin-conjugating enzyme E2 2 |
| contig_8557_6 | 0.0046 | 0.0062 | 0.0083 | 0.0612 | 0 | 1 | 0.027 | cytochrome b6 (chloroplast) |
| contig_21617_6 | 0.0047 | 0.0067 | 0.0086 | 0.0615 | 0 | 1 | 0.0271 | predicted protein |
| contig_9464_1 | 0.0047 | 0.0067 | 0.009 | 0.0615 | 0 | 1 | 0.0271 | cytochrome c heme attachment protein (chloroplast) |

| | | | | | | | | |
|---|---|---|---|---|---|---|---|---|
| contig_14379_2 | 0.0047 | 0.0067 | 0.009 | 0.0615 | 0 | 1 | 0.0271 | probable aspartyl aminopeptidase |
| contig_30881_4 | 0.0048 | 0.0067 | 0.0091 | 0.0615 | 0 | 1 | 0.0271 | pyruvate carboxylase |
| contig_8803_5 | 0.0048 | 0.0068 | 0.0091 | 0.0615 | 0 | 1 | 0.0271 | predicted protein |
| contig_38239_5 | 0.0051 | 0.0069 | 0.0091 | 0.0653 | 0 | 1 | 0.0288 | predicted protein |
| contig_24914_6 | 0.0052 | 0.0073 | 0.0092 | 0.0654 | 0 | 1 | 0.0288 | bifunctional dTDP-4-dehydrorhamnose 3 5-epimerase/dTDP-4-dehydrorhamnose reductase-like |
| contig_8921_1 | 0.0053 | 0.0073 | 0.0093 | 0.0654 | 0 | 1 | 0.0288 | predicted protein |
| contig_26568_4 | 0.0053 | 0.0075 | 0.0094 | 0.0654 | 0 | 1 | 0.0288 | atpase family aaa domain-containing protein 3 |
| contig_34578_1 | 0.0053 | 0.0076 | 0.0094 | 0.0654 | 0 | 1 | 0.0288 | predicted protein |
| contig_28789_2 | 0.0055 | 0.0076 | 0.0094 | 0.0665 | 0 | 1 | 0.0293 | predicted protein |
| contig_6155_1 | 0.0055 | 0.0077 | 0.0095 | 0.0665 | 0 | 1 | 0.0293 | light harvesting protein |
| contig_16671_2 | 0.0057 | 0.0078 | 0.0097 | 0.0688 | 0 | 1 | 0.0303 | ribosomal protein L21 (chloroplast) |
| contig_30517_5 | 0.0058 | 0.0078 | 0.0097 | 0.0695 | 0 | 1 | 0.0306 | predicted protein |
| contig_33628_1 | 0.006 | 0.0078 | 0.0099 | 0.0709 | 0 | 1 | 0.0313 | glycine-rich domain-containing protein 2-like |
| contig_3111_6 | 0.0062 | 0.008 | 0.0099 | 0.0724 | 0 | 1 | 0.0319 | acyl-CoA dehydrogenase |
| contig_25695_5 | 0.0062 | 0.0081 | 0.0101 | 0.0724 | 0 | 1 | 0.0319 | 40S ribosomal protein S7 |
| contig_13616_6 | 0.0063 | 0.0084 | 0.0102 | 0.0724 | 0 | 1 | 0.0319 | hsp70-like protein |
| contig_17134_1 | 0.0065 | 0.0085 | 0.0107 | 0.0747 | 0 | 1 | 0.0329 | 60S ribosomal protein L22 |
| contig_9065_4 | 0.0067 | 0.0086 | 0.011 | 0.0749 | 0 | 1 | 0.033 | tRNA cyclic N6-threonylcarbamoyladenosine(37) synthase TcdA |
| contig_30834_3 | 0.0067 | 0.009 | 0.0113 | 0.0749 | 0 | 1 | 0.033 | phosphoglycerate kinase |
| contig_40448_3 | 0.0067 | 0.0091 | 0.0116 | 0.0749 | 0 | 1 | 0.033 | RNA-binding transcriptional accessory protein |
| contig_26248_5 | 0.0067 | 0.0091 | 0.0116 | 0.0749 | 0 | 1 | 0.033 | predicted protein |
| contig_15521_3 | 0.0068 | 0.0091 | 0.0116 | 0.0753 | 0 | 1 | 0.0332 | predicted protein |
| contig_28103_5 | 0.0069 | 0.0091 | 0.0117 | 0.0753 | 0 | 1 | 0.0332 | magnesium chelatase subunit H |
| contig_39484_1 | 0.0071 | 0.0092 | 0.0118 | 0.077 | 0 | 1 | 0.034 | probable aspartyl aminopeptidase |
| contig_17394_3 | 0.0073 | 0.0094 | 0.0118 | 0.0784 | 0 | 1 | 0.0346 | proliferating cell nuclear antigen |
| contig_23548_3 | 0.0073 | 0.0094 | 0.0119 | 0.0784 | 0 | 1 | 0.0346 | ATP synthase subunit beta mitochondrial |

| | | | | | | | | |
|---|---|---|---|---|---|---|---|---|
| contig_5740_5 | 0.0075 | 0.0094 | 0.0119 | 0.07 95 | 0 | 1 | 0.03 51 | predicted protein |
| contig_30343_ 1 | 0.0076 | 0.0095 | 0.0123 | 0.07 95 | 0 | 1 | 0.03 51 | fucoxanthin chl a/c light-harvesting protein |
| contig_31795_ 6 | 0.0077 | 0.0095 | 0.0125 | 0.07 95 | 0 | 1 | 0.03 51 | glyceraldehyde-3-phosphate dehydrogenase |
| contig_27562_ 4 | 0.0077 | 0.0097 | 0.0126 | 0.07 95 | 0 | 1 | 0.03 51 | acyl-CoA dehydrogenase |
| contig_22705_ 1 | 0.0077 | 0.0097 | 0.0128 | 0.07 95 | 0 | 1 | 0.03 51 | glyceraldehyde-3-phosphate dehydrogenase precursor |
| contig_13426_ 4 | 0.0078 | 0.0099 | 0.0128 | 0.07 95 | 0 | 1 | 0.03 51 | long-chain fatty acid--CoA ligase |
| contig_12843_ 3 | 0.0078 | 0.01 | 0.0128 | 0.07 95 | 0 | 1 | 0.03 51 | proteasome subunit beta type-3 |
| contig_37452_ 3 | 0.0078 | 0.0101 | 0.0129 | 0.07 95 | 0 | 1 | 0.03 51 | ADP-ribosylation factor |
| contig_27293_ 4 | 0.008 | 0.0102 | 0.0129 | 0.08 11 | 0 | 1 | 0.03 57 | predicted protein |
| contig_35081_ 4 | 0.0081 | 0.0107 | 0.0132 | 0.08 13 | 0 | 1 | 0.03 58 | ubiquinol oxidase 4 chloroplastic/chromoplastic |
| contig_25894_ 5 | 0.0083 | 0.0113 | 0.0132 | 0.08 23 | 0 | 1 | 0.03 63 | predicted protein |
| contig_23416_ 6 | 0.0084 | 0.0114 | 0.0134 | 0.08 33 | 0 | 1 | 0.03 67 | predicted protein |
| contig_9976_6 | 0.0085 | 0.0116 | 0.0134 | 0.08 37 | 0 | 1 | 0.03 69 | poly(U)-binding-splicing factor PUF60 |
| contig_19673_ 5 | 0.0086 | 0.0116 | 0.0135 | 0.08 4 | 0 | 1 | 0.03 7 | predicted protein |
| contig_39738_ 3 | 0.009 | 0.0117 | 0.0135 | 0.08 58 | 0 | 1 | 0.03 78 | Asparaginyl-tRNA synthetase |
| contig_27905_ 6 | 0.009 | 0.0118 | 0.0136 | 0.08 58 | 0 | 1 | 0.03 78 | predicted protein |
| contig_24496_ 1 | 0.0091 | 0.0118 | 0.0137 | 0.08 58 | 0 | 1 | 0.03 78 | protein RCC2 homolog |
| contig_26782_ 3 | 0.0091 | 0.0119 | 0.0141 | 0.08 58 | 0 | 1 | 0.03 78 | predicted protein |
| contig_26787_ 4 | 0.0091 | 0.0119 | 0.0141 | 0.08 58 | 0 | 1 | 0.03 78 | DUF4336 domain-containing protein |
| contig_25969_ 5 | 0.0091 | 0.0119 | 0.0143 | 0.08 58 | 0 | 1 | 0.03 78 | Tic20-like protein |
| contig_22808_ 5 | 0.0092 | 0.0125 | 0.0144 | 0.08 58 | 0 | 1 | 0.03 78 | predicted protein |
| contig_5881_6 | 0.0093 | 0.0125 | 0.0147 | 0.08 58 | 0 | 1 | 0.03 78 | elongation factor Tu (chloroplast) |
| contig_1081_1 | 0.0094 | 0.0128 | 0.015 | 0.08 58 | 0 | 1 | 0.03 78 | hypothetical protein CBC21_09400 |
| contig_23995_ 6 | 0.0094 | 0.0128 | 0.015 | 0.08 58 | 0 | 1 | 0.03 78 | predicted protein |
| contig_10661_ 6 | 0.0094 | 0.0128 | 0.015 | 0.08 58 | 0 | 1 | 0.03 78 | glutaredoxin 2 |
| contig_29442_ 5 | 0.0095 | 0.0129 | 0.0151 | 0.08 58 | 0 | 1 | 0.03 78 | eukaryotic translation initiation factor 2 subunit alpha |
| contig_8784_1 | 0.0095 | 0.0129 | 0.0152 | 0.08 58 | 0 | 1 | 0.03 78 | 40S ribosomal protein S10 |

| | | | | | | | | |
|---|---|---|---|---|---|---|---|---|
| contig_6927_4 | 0.0097 | 0.0131 | 0.0152 | 0.08 61 | 0 | 1 | 0.03 8 | predicted protein |
| contig_25619_ 1 | 0.0097 | 0.0132 | 0.0153 | 0.08 61 | 0 | 1 | 0.03 8 | ornithine--oxo-acid transaminase |
| contig_29760_ 5 | 0.0099 | 0.0134 | 0.0154 | 0.08 74 | 0 | 1 | 0.03 85 | predicted protein |
| contig_25394_ 4 | 0.0099 | 0.0134 | 0.0155 | 0.08 74 | 0 | 1 | 0.03 85 | prohibitin putative |
| contig_23722_ 3 | 0.01 | 0.0135 | 0.0156 | 0.08 74 | 0 | 1 | 0.03 85 | light harvesting protein |
| contig_5475_2 | 0.0101 | 0.0135 | 0.0156 | 0.08 74 | 0 | 1 | 0.03 85 | coatomer subunit epsilon |
| contig_29278_ 3 | 0.0102 | 0.0135 | 0.0157 | 0.08 81 | 0 | 1 | 0.03 89 | predicted protein |
| contig_23256_ 4 | 0.0107 | 0.0136 | 0.0161 | 0.09 18 | 0 | 1 | 0.04 05 | fumarate hydratase mitochondrial |
| contig_17554_ 4 | 0.0107 | 0.0137 | 0.0162 | 0.09 18 | 0 | 1 | 0.04 05 | predicted protein |
| contig_9347_5 | 0.011 | 0.0141 | 0.0164 | 0.09 35 | 0 | 1 | 0.04 12 | putative glycerol-3-phosphate transporter 1 |
| contig_35648_ 5 | 0.0113 | 0.0141 | 0.0164 | 0.09 55 | 0 | 1 | 0.04 21 | ubiquitin / ribosomal protein CEP52 fusion protein putative |
| contig_30131_ 2 | 0.0114 | 0.0144 | 0.0165 | 0.09 55 | 0 | 1 | 0.04 21 | dihydrolipoyl dehydrogenase |
| contig_23352_ 4 | 0.0116 | 0.0147 | 0.0166 | 0.09 55 | 0 | 1 | 0.04 21 | malate dehydrogenase |
| contig_25286_ 5 | 0.0116 | 0.0147 | 0.0168 | 0.09 55 | 0 | 1 | 0.04 21 | ap-2 complex subunit mu |
| contig_28302_ 4 | 0.0116 | 0.015 | 0.0168 | 0.09 55 | 0 | 1 | 0.04 21 | predicted protein |
| contig_25924_ 5 | 0.0117 | 0.015 | 0.0171 | 0.09 55 | 0 | 1 | 0.04 21 | ATP-dependent Clp protease proteolytic subunit |
| contig_8461_2 | 0.0118 | 0.0151 | 0.0171 | 0.09 55 | 0 | 1 | 0.04 21 | SDR family oxidoreductase |
| contig_10415_ 5 | 0.0118 | 0.0152 | 0.0173 | 0.09 55 | 0 | 1 | 0.04 21 | DNA-binding protein 42 kDa |
| contig_14563_ 3 | 0.0119 | 0.0152 | 0.0174 | 0.09 55 | 0 | 1 | 0.04 21 | predicted protein |
| contig_30640_ 5 | 0.0119 | 0.0153 | 0.0177 | 0.09 55 | 0 | 1 | 0.04 21 | predicted protein |
| contig_5989_3 | 0.0119 | 0.0153 | 0.0178 | 0.09 56 | 0 | 1 | 0.04 21 | valyl-tRNA synthetase |
| contig_15161_ 3 | 0.0123 | 0.0154 | 0.0178 | 0.09 83 | 0 | 1 | 0.04 33 | serine/threonine-protein phosphatase PP1-beta catalytic subunit |
| contig_23563_ 5 | 0.0125 | 0.0156 | 0.0179 | 0.09 86 | 0 | 1 | 0.04 35 | FK506-binding protein 2 isoform X2 |
| contig_33008_ 5 | 0.0125 | 0.0157 | 0.0179 | 0.09 86 | 0 | 1 | 0.04 35 | protein fucoxanthin chlorophyll a/c protein |
| contig_28396_ 1 | 0.0126 | 0.0157 | 0.018 | 0.09 86 | 0 | 1 | 0.04 35 | chaperone DnaK |
| contig_25343_ 2 | 0.0128 | 0.0157 | 0.018 | 0.09 86 | 0 | 1 | 0.04 35 | predicted protein |

| | | | | | | | | |
|---|---|---|---|---|---|---|---|---|
| contig_39450_1 | 0.0128 | 0.0162 | 0.0185 | 0.0986 | 0 | 1 | 0.0435 | CTL-like protein DDB_G0288717 |
| contig_5427_6 | 0.0128 | 0.0164 | 0.0192 | 0.0986 | 0 | 1 | 0.0435 | tyrosyl-tRNA synthetase |
| contig_14556_3 | 0.0129 | 0.0164 | 0.0193 | 0.0986 | 0 | 1 | 0.0435 | anion-transporting ATPase |
| contig_24360_5 | 0.0129 | 0.0165 | 0.0193 | 0.0986 | 0 | 1 | 0.0435 | splicing factor 3B subunit 3-like |
| contig_15864_3 | 0.0131 | 0.0165 | 0.0194 | 0.0993 | 0 | 1 | 0.0438 | 50S ribosomal protein L15 |
| contig_23649_2 | 0.0132 | 0.0166 | 0.0194 | 0.0993 | 0 | 1 | 0.0438 | heat shock protein 90-2 |
| contig_32718_6 | 0.0132 | 0.017 | 0.0197 | 0.0993 | 0 | 1 | 0.0438 | aspartate-semialdehyde dehydrogenase |
| contig_15357_4 | 0.0134 | 0.017 | 0.0197 | 0.0993 | 0 | 1 | 0.0438 | calreticulin |
| contig_17958_6 | 0.0134 | 0.0171 | 0.0198 | 0.0993 | 0 | 1 | 0.0438 | luminal-binding protein |
| contig_9376_2 | 0.0135 | 0.0171 | 0.0199 | 0.0993 | 0 | 1 | 0.0438 | predicted protein |
| contig_24221_5 | 0.0135 | 0.0171 | 0.0199 | 0.0993 | 0 | 1 | 0.0438 | ATP-binding cassette superfamily |
| contig_17853_5 | 0.0135 | 0.0177 | 0.0203 | 0.0993 | 0 | 1 | 0.0438 | predicted protein |
| contig_27521_6 | 0.0136 | 0.0177 | 0.0204 | 0.0993 | 0 | 1 | 0.0438 | predicted protein |
| contig_28995_4 | 0.0137 | 0.0178 | 0.0204 | 0.0998 | 0 | 1 | 0.044 | predicted protein |
| contig_34814_4 | 0.0141 | 0.0178 | 0.0205 | 0.1015 | 0 | 1 | 0.0447 | predicted protein |
| contig_38523_3 | 0.0141 | 0.018 | 0.0206 | 0.1015 | 0 | 1 | 0.0447 | h+-translocating pyrophosphatase family |
| contig_28181_3 | 0.0141 | 0.018 | 0.0207 | 0.1015 | 0 | 1 | 0.0447 | dihydroxy-acid dehydratase |
| contig_5591_4 | 0.0143 | 0.0181 | 0.0207 | 0.1022 | 0 | 1 | 0.0451 | predicted protein |
| contig_13636_6 | 0.0144 | 0.0183 | 0.0208 | 0.1022 | 0 | 1 | 0.0451 | acyl-CoA dehydrogenase-like protein |
| contig_16696_4 | 0.0147 | 0.0192 | 0.0214 | 0.1038 | 0 | 1 | 0.0458 | pyruvate kinase |
| contig_26535_2 | 0.0147 | 0.0193 | 0.0218 | 0.1038 | 0 | 1 | 0.0458 | tubulin alpha chain |
| contig_27608_2 | 0.015 | 0.0193 | 0.0218 | 0.1038 | 0 | 1 | 0.0458 | lysine-tRNA ligase |
| contig_24180_5 | 0.015 | 0.0194 | 0.0222 | 0.1038 | 0 | 1 | 0.0458 | predicted protein |
| contig_25971_2 | 0.015 | 0.0194 | 0.0222 | 0.1038 | 0 | 1 | 0.0458 | purine nucleoside phosphorylase I inosine and guanosine-specific |
| contig_28073_5 | 0.0151 | 0.0197 | 0.0224 | 0.1038 | 0 | 1 | 0.0458 | NAD-dependent epimerase/dehydratase family protein |
| contig_18892_6 | 0.0152 | 0.0197 | 0.0225 | 0.1038 | 0 | 1 | 0.0458 | predicted protein |

| | | | | | | | | |
|---|---|---|---|---|---|---|---|---|
| contig_11292_5 | 0.0152 | 0.0198 | 0.0227 | 0.1038 | 0 | 1 | 0.0458 | heat shock protein belonging to the HSP70 family |
| contig_7582_3 | 0.0153 | 0.0199 | 0.0232 | 0.1038 | 0 | 1 | 0.0458 | vanadium-dependent haloperoxidase |
| contig_9342_6 | 0.0153 | 0.02 | 0.0237 | 0.1038 | 0 | 1 | 0.0458 | predicted protein |
| contig_13922_3 | 0.0154 | 0.0202 | 0.0238 | 0.1042 | 0 | 1 | 0.0459 | annexin |
| contig_9617_4 | 0.0155 | 0.0204 | 0.0242 | 0.1042 | 0 | 1 | 0.0459 | bifunctional methylenetetrahydrofolate dehydrogenase/methenyltetrahydrofolate cyclohydrolase FolD |
| contig_35016_2 | 0.0156 | 0.0204 | 0.0246 | 0.1042 | 0 | 1 | 0.0459 | predicted protein |
| contig_11658_2 | 0.0156 | 0.0204 | 0.0246 | 0.1042 | 0 | 1 | 0.0459 | pyrimidine/purine nucleoside phosphorylase |
| contig_23664_6 | 0.0157 | 0.0206 | 0.0249 | 0.1042 | 0 | 1 | 0.0459 | enoyl-acp reductase |
| contig_26180_3 | 0.0158 | 0.0207 | 0.025 | 0.1045 | 0 | 1 | 0.0461 | predicted protein |
| contig_22513_4 | 0.0161 | 0.0207 | 0.025 | 0.1062 | 0 | 1 | 0.0468 | predicted protein |
| contig_6178_5 | 0.0162 | 0.0207 | 0.0252 | 0.1062 | 0 | 1 | 0.0468 | rubisco expression protein (chloroplast) |
| contig_33347_4 | 0.0164 | 0.0214 | 0.0252 | 0.1062 | 0 | 1 | 0.0468 | V-type ATPase A subunit |
| contig_34331_5 | 0.0164 | 0.0218 | 0.0262 | 0.1062 | 0 | 1 | 0.0468 | Light-harvesting complex I LH38 protein |
| contig_29943_6 | 0.0165 | 0.022 | 0.0263 | 0.1062 | 0 | 1 | 0.0468 | Got1-domain-containing protein |
| contig_29735_5 | 0.0165 | 0.0222 | 0.0263 | 0.1062 | 0 | 1 | 0.0468 | dihydrolipoamide succinyltransferase |
| contig_33723_4 | 0.0166 | 0.0222 | 0.0267 | 0.1068 | 0 | 1 | 0.0471 | predicted protein |
| contig_19000_1 | 0.0168 | 0.0224 | 0.0267 | 0.1072 | 0 | 1 | 0.0473 | predicted protein |
| contig_17693_5 | 0.0168 | 0.0224 | 0.0267 | 0.1072 | 0 | 1 | 0.0473 | elongation factor-like |
| contig_25874_5 | 0.017 | 0.023 | 0.0271 | 0.1076 | 0 | 1 | 0.0475 | predicted protein |
| contig_19197_6 | 0.0171 | 0.0232 | 0.0273 | 0.1076 | 0 | 1 | 0.0475 | casein kinase I isoform X1 |
| contig_27950_6 | 0.0171 | 0.0237 | 0.0273 | 0.1077 | 0 | 1 | 0.0475 | aspartyl-tRNA synthetase |
| contig_37879_1 | 0.0173 | 0.0238 | 0.0273 | 0.1081 | 0 | 1 | 0.0477 | UPF0187 protein |
| contig_5576_4 | 0.0174 | 0.0242 | 0.0277 | 0.1086 | 0 | 1 | 0.0479 | transketolase |
| contig_11625_6 | 0.0177 | 0.0247 | 0.0279 | 0.1089 | 0 | 1 | 0.048 | T9SS type A sorting domain-containing protein |
| contig_9980_5 | 0.0177 | 0.0249 | 0.028 | 0.1089 | 0 | 1 | 0.048 | protein fucoxanthin chlorophyll a/c protein |
| contig_39499_3 | 0.0178 | 0.0249 | 0.0282 | 0.1089 | 0 | 1 | 0.048 | adenylosuccinate synthetase |

| | | | | | | | | |
|---|---|---|---|---|---|---|---|---|
| contig_10138_6 | 0.0178 | 0.025 | 0.0284 | 0.1089 | 0 | 1 | 0.048 | adenylate kinase chloroplastic |
| contig_22726_1 | 0.0179 | 0.025 | 0.0287 | 0.1089 | 0 | 1 | 0.048 | Valine-tRNA ligase |
| contig_18202_5 | 0.0179 | 0.025 | 0.0287 | 0.1089 | 0 | 1 | 0.048 | protein fucoxanthin chlorophyll a/c protein |
| contig_20080_5 | 0.018 | 0.0256 | 0.0293 | 0.109 | 0 | 1 | 0.0481 | AP-1 complex subunit beta-1-like |
| contig_30325_5 | 0.0181 | 0.0262 | 0.0293 | 0.1093 | 0 | 1 | 0.0482 | predicted protein |
| contig_40753_3 | 0.0183 | 0.0263 | 0.0297 | 0.1099 | 0 | 1 | 0.0485 | V-type proton ATPase subunit E |
| contig_19778_5 | 0.0185 | 0.0267 | 0.0298 | 0.1108 | 0 | 1 | 0.0488 | voltage-gated chloride channel |
| contig_27261_5 | 0.0191 | 0.0267 | 0.0298 | 0.1137 | 0 | 1 | 0.0501 | enoyl-CoA hydratase |
| contig_7982_5 | 0.0192 | 0.0271 | 0.0299 | 0.1137 | 0 | 1 | 0.0501 | photosystem II protein V (chloroplast) |
| contig_17991_3 | 0.0193 | 0.0271 | 0.0299 | 0.1137 | 0 | 1 | 0.0501 | light-harvesting protein |
| contig_15626_6 | 0.0193 | 0.0273 | 0.0304 | 0.1137 | 0 | 1 | 0.0501 | predicted protein |
| contig_21931_4 | 0.0194 | 0.0273 | 0.0304 | 0.1137 | 0 | 1 | 0.0501 | chaperonin GroEL (chloroplast) |
| contig_30065_5 | 0.0194 | 0.0273 | 0.0306 | 0.1137 | 0 | 1 | 0.0501 | importin subunit alpha-1-like isoform X2 |
| contig_26554_3 | 0.0197 | 0.0275 | 0.0307 | 0.114 | 0 | 1 | 0.0503 | predicted protein |
| contig_38301_1 | 0.0197 | 0.0279 | 0.0307 | 0.114 | 0 | 1 | 0.0503 | predicted protein |
| contig_40094_2 | 0.0197 | 0.028 | 0.031 | 0.114 | 0 | 1 | 0.0503 | 40S ribosomal protein S14 |
| contig_40524_3 | 0.0198 | 0.0282 | 0.0316 | 0.114 | 0 | 1 | 0.0503 | light harvesting protein |
| contig_11702_1 | 0.0199 | 0.0284 | 0.0317 | 0.1144 | 0 | 1 | 0.0504 | glutamine synthetase catalytic domain protein protein |
| contig_33265_1 | 0.02 | 0.0287 | 0.0318 | 0.1145 | 0 | 1 | 0.0505 | phenylalanine-tRNA ligase beta subunit |
| contig_25846_1 | 0.0202 | 0.0289 | 0.032 | 0.1148 | 0 | 1 | 0.0506 | glyceraldehyde-3-phosphate dehydrogenase |
| contig_22734_3 | 0.0203 | 0.0293 | 0.032 | 0.1148 | 0 | 1 | 0.0506 | predicted protein |
| contig_24490_6 | 0.0204 | 0.0293 | 0.0324 | 0.1148 | 0 | 1 | 0.0506 | predicted protein |
| contig_28450_6 | 0.0204 | 0.0297 | 0.0324 | 0.1148 | 0 | 1 | 0.0506 | transketolase |
| contig_9968_3 | 0.0205 | 0.0298 | 0.0325 | 0.1148 | 0 | 1 | 0.0506 | predicted protein |
| contig_28903_4 | 0.0206 | 0.0298 | 0.0326 | 0.1148 | 0 | 1 | 0.0506 | predicted protein |
| contig_25835_2 | 0.0207 | 0.0299 | 0.0326 | 0.1148 | 0 | 1 | 0.0506 | histidinol dehydrogenase |
| contig_16887_1 | 0.0207 | 0.0299 | 0.0338 | 0.1148 | 0 | 1 | 0.0506 | predicted protein |

| | | | | | | | | |
|---|---|---|---|---|---|---|---|---|
| contig_40308_5 | 0.0207 | 0.0304 | 0.0339 | 0.1148 | 0 | 1 | 0.0506 | 40S ribosomal protein S6 |
| contig_8751_4 | 0.0208 | 0.0304 | 0.034 | 0.1148 | 0 | 1 | 0.0506 | predicted protein |
| contig_27614_3 | 0.0214 | 0.0307 | 0.0343 | 0.1172 | 0 | 1 | 0.0517 | Regulator of nonsense transcripts 1-like protein |
| contig_28448_5 | 0.0217 | 0.0307 | 0.0344 | 0.1187 | 0 | 1 | 0.0523 | glycine dehydrogenase |
| contig_12195_1 | 0.0218 | 0.0307 | 0.0346 | 0.1188 | 0 | 1 | 0.0524 | thioredoxin-like protein 4A isoform X1 |
| contig_5529_5 | 0.022 | 0.031 | 0.0347 | 0.1192 | 0 | 1 | 0.0526 | photosystem II protein D1 (chloroplast) |
| contig_14030_3 | 0.0221 | 0.0316 | 0.0351 | 0.1194 | 0 | 1 | 0.0527 | S-adenosylmethionine carrier 1 chloroplastic/mitochondrial |
| contig_26348_5 | 0.0222 | 0.0317 | 0.0352 | 0.1194 | 0 | 1 | 0.0527 | heat shock protein 70 |
| contig_37313_1 | 0.0222 | 0.0318 | 0.0352 | 0.1194 | 0 | 1 | 0.0527 | predicted protein |
| contig_30107_6 | 0.0224 | 0.032 | 0.0352 | 0.1201 | 0 | 1 | 0.0529 | ATP-dependent protease HslVU ATPase subunit |
| contig_7643_5 | 0.0225 | 0.0321 | 0.0352 | 0.1203 | 0 | 1 | 0.0531 | ATP synthase CF1 beta subunit (chloroplast) |
| contig_6818_2 | 0.0227 | 0.0324 | 0.0361 | 0.1206 | 0 | 1 | 0.0532 | predicted protein |
| contig_9754_5 | 0.023 | 0.0324 | 0.0361 | 0.1219 | 0 | 1 | 0.0537 | long-chain-fatty-acid-CoA ligase |
| contig_28046_4 | 0.0232 | 0.0326 | 0.0366 | 0.1227 | 0 | 1 | 0.0541 | light harvesting protein |
| contig_29569_5 | 0.0234 | 0.0326 | 0.0367 | 0.1234 | 0 | 1 | 0.0544 | Sedoheptulose-1 7-bisphosphatase chloroplastic |
| contig_37779_2 | 0.0237 | 0.0336 | 0.0372 | 0.1242 | 0 | 1 | 0.0548 | protein fucoxanthin chlorophyll a/c protein |
| contig_34506_1 | 0.0238 | 0.0339 | 0.0374 | 0.1242 | 0 | 1 | 0.0548 | cell division protein FtsH |
| contig_33643_5 | 0.0242 | 0.034 | 0.0374 | 0.1258 | 0 | 1 | 0.0555 | predicted protein |
| contig_24581_4 | 0.0242 | 0.0341 | 0.0377 | 0.1258 | 0 | 1 | 0.0555 | predicted protein |
| contig_16700_6 | 0.0246 | 0.0343 | 0.0378 | 0.127 | 0 | 1 | 0.056 | serine decarboxylase |
| contig_32841_4 | 0.0247 | 0.0346 | 0.0383 | 0.127 | 0 | 1 | 0.056 | predicted protein |
| contig_27487_4 | 0.0249 | 0.0347 | 0.0383 | 0.127 | 0 | 1 | 0.056 | ---NA--- |
| contig_7406_5 | 0.0249 | 0.035 | 0.0384 | 0.127 | 0 | 1 | 0.056 | predicted protein |
| contig_22324_4 | 0.025 | 0.0352 | 0.0385 | 0.127 | 0 | 1 | 0.056 | ---NA--- |
| contig_16604_4 | 0.025 | 0.0352 | 0.0386 | 0.127 | 0 | 1 | 0.056 | myo-inositol 2-dehydrogenase putative |
| contig_10672_3 | 0.0252 | 0.0352 | 0.0387 | 0.1272 | 0 | 1 | 0.0561 | ATP synthase subunit |
| contig_39564_2 | 0.0252 | 0.0352 | 0.0391 | 0.1272 | 0 | 1 | 0.0561 | predicted protein |

| | | | | | | | | |
|---|---|---|---|---|---|---|---|---|
| contig_23545_2 | 0.0256 | 0.0354 | 0.0397 | 0.1291 | 0 | 1 | 0.0569 | predicted protein |
| contig_14305_2 | 0.0262 | 0.0361 | 0.0398 | 0.1308 | 0 | 1 | 0.0577 | predicted protein |
| contig_32836_4 | 0.0262 | 0.0361 | 0.0399 | 0.1308 | 0 | 1 | 0.0577 | predicted protein |
| contig_33755_6 | 0.0263 | 0.0367 | 0.0404 | 0.1308 | 0 | 1 | 0.0577 | predicted protein |
| contig_6246_4 | 0.0263 | 0.0372 | 0.0404 | 0.1308 | 0 | 1 | 0.0577 | predicted protein |
| contig_8347_5 | 0.0267 | 0.0374 | 0.0413 | 0.1319 | 0 | 1 | 0.0581 | elongation factor Ts |
| contig_8988_3 | 0.0267 | 0.0374 | 0.0413 | 0.1319 | 0 | 1 | 0.0581 | DUF1295 domain-containing protein |
| contig_27839_5 | 0.0269 | 0.0377 | 0.0419 | 0.1323 | 0 | 1 | 0.0583 | predicted protein |
| contig_18966_2 | 0.0271 | 0.0378 | 0.042 | 0.1323 | 0 | 1 | 0.0583 | predicted protein |
| contig_5747_6 | 0.0271 | 0.0383 | 0.0421 | 0.1323 | 0 | 1 | 0.0583 | photosystem I assembly protein Ycf4 (chloroplast) |
| contig_38571_1 | 0.0273 | 0.0383 | 0.0422 | 0.1323 | 0 | 1 | 0.0583 | photosystem I subunit III (chloroplast) |
| contig_14719_6 | 0.0273 | 0.0384 | 0.0423 | 0.1323 | 0 | 1 | 0.0583 | thioredoxin-like |
| contig_9535_1 | 0.0273 | 0.0385 | 0.0429 | 0.1323 | 0 | 1 | 0.0583 | transitional endoplasmic reticulum ATPase |
| contig_9105_2 | 0.0275 | 0.0386 | 0.0435 | 0.1327 | 0 | 1 | 0.0585 | SufE-like protein 1 chloroplastic/mitochondrial |
| contig_26987_1 | 0.0277 | 0.0387 | 0.0439 | 0.1335 | 0 | 1 | 0.0588 | predicted protein |
| contig_32695_4 | 0.0279 | 0.0396 | 0.0442 | 0.1337 | 0 | 1 | 0.0589 | predicted protein |
| contig_17354_1 | 0.028 | 0.0397 | 0.0459 | 0.1338 | 0 | 1 | 0.059 | T9SS type A sorting domain-containing protein |
| contig_23843_1 | 0.0282 | 0.0398 | 0.046 | 0.1341 | 0 | 1 | 0.0591 | methyltransferase domain-containing protein |
| contig_24751_3 | 0.0282 | 0.0399 | 0.0471 | 0.1341 | 0 | 1 | 0.0591 | predicted protein |
| contig_27062_5 | 0.0284 | 0.0404 | 0.0472 | 0.1348 | 0 | 1 | 0.0594 | heat shock protein 70 |
| contig_5809_2 | 0.0287 | 0.0404 | 0.0474 | 0.1355 | 0 | 1 | 0.0597 | ---NA--- |
| contig_11453_6 | 0.0289 | 0.0413 | 0.0477 | 0.1355 | 0 | 1 | 0.0598 | major facilitator transporter |
| contig_30861_4 | 0.0289 | 0.0419 | 0.0484 | 0.1355 | 0 | 1 | 0.0598 | fucoxanthin chlorophyll a/c protein 8 |
| contig_33639_6 | 0.0293 | 0.0419 | 0.0485 | 0.1366 | 0 | 1 | 0.0602 | ---NA--- |
| contig_16707_4 | 0.0293 | 0.0421 | 0.0485 | 0.1366 | 0 | 1 | 0.0602 | nitrate reductase |
| contig_18571_6 | 0.0296 | 0.0422 | 0.0489 | 0.1369 | 0 | 1 | 0.0604 | phosphoribulokinase |
| contig_22960_6 | 0.0297 | 0.0423 | 0.0491 | 0.1369 | 0 | 1 | 0.0604 | urease alpha subunit |

| | | | | | | | | |
|---|---|---|---|---|---|---|---|---|
| contig_24459_5 | 0.0298 | 0.0423 | 0.0494 | 0.1369 | 0 | 1 | 0.0604 | fact complex subunit ssrp1 |
| contig_15223_4 | 0.0298 | 0.0429 | 0.0496 | 0.1369 | 0 | 1 | 0.0604 | predicted protein |
| contig_16919_4 | 0.0299 | 0.0438 | 0.0497 | 0.1369 | 0 | 1 | 0.0604 | predicted protein |
| contig_23620_5 | 0.0299 | 0.0442 | 0.0497 | 0.1369 | 0 | 1 | 0.0604 | nitrate transporter |
| contig_38768_2 | 0.0302 | 0.0459 | 0.0498 | 0.1379 | 0 | 1 | 0.0608 | ---NA--- |
| contig_34219_5 | 0.0304 | 0.046 | 0.0498 | 0.1379 | 0 | 1 | 0.0608 | Two component regulator three Y domain protein |
| contig_22629_4 | 0.0304 | 0.0472 | 0.0499 | 0.1379 | 0 | 1 | 0.0608 | acetyl-coa carboxylase |
| contig_35069_1 | 0.0306 | 0.0473 | 0.0506 | 0.1379 | 0 | 1 | 0.0608 | translation elongation factor-like protein |
| contig_21628_5 | 0.0307 | 0.0477 | 0.052 | 0.1379 | 0 | 1 | 0.0608 | Tubulin alpha-5 chain |
| contig_18043_1 | 0.0307 | 0.0484 | 0.052 | 0.1379 | 0 | 1 | 0.0608 | Light-harvesting complex I LH38 protein |
| contig_30700_3 | 0.0307 | 0.0485 | 0.0537 | 0.1379 | 0 | 1 | 0.0608 | predicted protein |
| contig_5813_3 | 0.031 | 0.0485 | 0.0551 | 0.1386 | 0 | 1 | 0.0611 | transketolase |
| contig_30585_6 | 0.0316 | 0.0489 | 0.0553 | 0.1404 | 0 | 1 | 0.0619 | ribosomal protein S2 |
| contig_22686_4 | 0.0316 | 0.0489 | 0.0553 | 0.1404 | 0 | 1 | 0.0619 | predicted protein |
| contig_39759_2 | 0.0317 | 0.0493 | 0.0555 | 0.1404 | 0 | 1 | 0.0619 | antibiotic biosynthesis monooxygenase |
| contig_29871_5 | 0.0318 | 0.0494 | 0.0558 | 0.1405 | 0 | 1 | 0.0619 | lipoprotein type 6 |
| contig_23908_1 | 0.032 | 0.0497 | 0.0559 | 0.1405 | 0 | 1 | 0.0619 | mitochondrial carrier protein |
| contig_39493_2 | 0.032 | 0.0497 | 0.0562 | 0.1405 | 0 | 1 | 0.0619 | 60s ribosomal protein |
| contig_24628_4 | 0.0321 | 0.0498 | 0.0566 | 0.1405 | 0 | 1 | 0.0619 | predicted protein |
| contig_30564_1 | 0.0324 | 0.0498 | 0.0573 | 0.1411 | 0 | 1 | 0.0622 | predicted protein |
| contig_30020_5 | 0.0324 | 0.0499 | 0.0575 | 0.1411 | 0 | 1 | 0.0622 | chaperonin GroEL |
| contig_29776_5 | 0.0325 | 0.0499 | 0.058 | 0.1411 | 0 | 1 | 0.0622 | fructose-bisphosphate aldolase |
| contig_6296_5 | 0.0326 | 0.052 | 0.0581 | 0.1411 | 0 | 1 | 0.0622 | predicted protein |
| contig_24783_5 | 0.0326 | 0.0521 | 0.0592 | 0.1411 | 0 | 1 | 0.0622 | mitochondrial protein translocase MPT family |
| contig_37802_5 | 0.0334 | 0.0537 | 0.0594 | 0.144 | 0 | 1 | 0.0635 | predicted protein |
| contig_19001_2 | 0.0336 | 0.0551 | 0.0598 | 0.1442 | 0 | 1 | 0.0636 | predicted protein |
| contig_25881_5 | 0.0338 | 0.0553 | 0.0599 | 0.1446 | 0 | 1 | 0.0638 | predicted protein |

| | | | | | | | | |
|---|---|---|---|---|---|---|---|---|
| contig_33602_5 | 0.0339 | 0.0555 | 0.06 | 0.1448 | 0 | 1 | 0.0638 | FAD-binding domain-containing protein |
| contig_38702_2 | 0.034 | 0.0558 | 0.06 | 0.1448 | 0 | 1 | 0.0638 | predicted protein |
| contig_30444_6 | 0.0341 | 0.0558 | 0.0607 | 0.1448 | 0 | 1 | 0.0638 | NADH dehydrogenase (ubiquinone) 1 beta subcomplex subunit 9 |
| contig_10451_3 | 0.0343 | 0.0562 | 0.061 | 0.1454 | 0 | 1 | 0.0641 | predicted protein |
| contig_40249_6 | 0.0344 | 0.0564 | 0.0616 | 0.1455 | 0 | 1 | 0.0641 | translation elongation factor-like protein |
| contig_22590_5 | 0.0346 | 0.0573 | 0.0623 | 0.1456 | 0 | 1 | 0.0642 | glutathione reductase |
| contig_30141_2 | 0.0347 | 0.0575 | 0.0625 | 0.1456 | 0 | 1 | 0.0642 | Lipoamide acyltransferase component of branched-chain alpha-keto acid dehydrogenase complex mitochondrial |
| contig_27187_2 | 0.035 | 0.0578 | 0.0632 | 0.1456 | 0 | 1 | 0.0642 | CAAX prenyl protease 1 |
| contig_34140_1 | 0.0351 | 0.059 | 0.0634 | 0.1456 | 0 | 1 | 0.0642 | predicted protein |
| contig_14441_4 | 0.0352 | 0.0591 | 0.0637 | 0.1456 | 0 | 1 | 0.0642 | predicted protein |
| contig_39493_5 | 0.0352 | 0.0594 | 0.0639 | 0.1456 | 0 | 1 | 0.0642 | 60s ribosomal protein |
| contig_28453_2 | 0.0352 | 0.0598 | 0.064 | 0.1456 | 0 | 1 | 0.0642 | methionine-dependent methyltransferase |
| contig_22418_5 | 0.0352 | 0.0599 | 0.0699 | 0.1456 | 0 | 1 | 0.0642 | protein fucoxanthin chlorophyll a/c protein |
| contig_34823_4 | 0.0354 | 0.06 | 0.0701 | 0.1458 | 0 | 1 | 0.0643 | 60S ribosomal protein L18-2 |
| contig_25113_4 | 0.0361 | 0.06 | 0.071 | 0.1477 | 0 | 1 | 0.0651 | YEATS domain-containing protein 4 |
| contig_25862_1 | 0.0361 | 0.061 | 0.071 | 0.1477 | 0 | 1 | 0.0651 | predicted protein |
| contig_28761_2 | 0.0361 | 0.061 | 0.0715 | 0.1477 | 0 | 1 | 0.0651 | protein dispatched homolog 3-like |
| contig_30735_2 | 0.0366 | 0.0616 | 0.0731 | 0.1491 | 0 | 1 | 0.0657 | myo-inositol 2-dehydrogenase putative |
| contig_28237_2 | 0.0367 | 0.0623 | 0.0736 | 0.1491 | 0 | 1 | 0.0657 | plant UBX domain-containing protein 10-like |
| contig_18233_1 | 0.0369 | 0.0627 | 0.0753 | 0.1495 | 0 | 1 | 0.0659 | dihydrolipoamide dehydrogenase |
| contig_5402_3 | 0.0372 | 0.0633 | 0.0789 | 0.1505 | 0 | 1 | 0.0663 | NAD(P)-dependent oxidoreductase |
| contig_29531_5 | 0.0374 | 0.0634 | 0.0794 | 0.1506 | 0 | 1 | 0.0664 | oxidoreductase putative |
| contig_7935_3 | 0.0374 | 0.0639 | 0.0795 | 0.1506 | 0 | 1 | 0.0664 | predicted protein |
| contig_13885_3 | 0.0375 | 0.064 | 0.08 | 0.1506 | 0 | 1 | 0.0664 | predicted protein |
| contig_38142_5 | 0.0377 | 0.064 | 0.0801 | 0.1508 | 0 | 1 | 0.0665 | ATP synthase F1 subunit alpha (mitochondrion) |
| contig_12161_5 | 0.0378 | 0.0701 | 0.0806 | 0.1508 | 0 | 1 | 0.0665 | predicted protein |

| | | | | | | | | |
|---|---|---|---|---|---|---|---|---|
| contig_30073_5 | 0.0382 | 0.071 | 0.0808 | 0.1515 | 0 | 1 | 0.0668 | transaldolase |
| contig_39157_3 | 0.0383 | 0.0713 | 0.0808 | 0.1515 | 0 | 1 | 0.0668 | predicted protein |
| contig_17114_6 | 0.0383 | 0.0731 | 0.081 | 0.1515 | 0 | 1 | 0.0668 | predicted protein |
| contig_7234_4 | 0.0384 | 0.0736 | 0.0813 | 0.1515 | 0 | 1 | 0.0668 | ras-related gtp-binding protein |
| contig_10561_5 | 0.0385 | 0.0739 | 0.0814 | 0.1515 | 0 | 1 | 0.0668 | 6-phosphogluconate dehydrogenase |
| contig_30780_6 | 0.0386 | 0.0789 | 0.0814 | 0.1515 | 0 | 1 | 0.0668 | possible transporter |
| contig_36968_1 | 0.0387 | 0.0795 | 0.0826 | 0.1515 | 0 | 1 | 0.0668 | 60S ribosomal protein L38 |
| contig_37899_1 | 0.039 | 0.0796 | 0.0827 | 0.1526 | 0 | 1 | 0.0673 | ribosomal protein L2 (chloroplast) |
| contig_40230_1 | 0.0391 | 0.0796 | 0.0827 | 0.1526 | 0 | 1 | 0.0673 | predicted protein |
| contig_23500_4 | 0.0396 | 0.0806 | 0.0833 | 0.154 | 0 | 1 | 0.0679 | proline-tRNA ligase |
| contig_9330_5 | 0.0397 | 0.0808 | 0.0839 | 0.154 | 0 | 1 | 0.0679 | short-chain alcohol dehydrogenase with NAD or NADP as acceptor |
| contig_38881_1 | 0.0398 | 0.0808 | 0.0848 | 0.154 | 0 | 1 | 0.0679 | SulP family inorganic anion transporter |
| contig_37722_1 | 0.0399 | 0.081 | 0.0869 | 0.154 | 0 | 1 | 0.0679 | hypothetical protein |
| contig_28327_6 | 0.0403 | 0.081 | 0.0869 | 0.1548 | 0 | 1 | 0.0683 | rab GDP dissociation inhibitor alpha |
| contig_24921_6 | 0.0404 | 0.0814 | 0.0875 | 0.1548 | 0 | 1 | 0.0683 | nitrate transporter |
| contig_41330_2 | 0.0404 | 0.0814 | 0.0877 | 0.1548 | 0 | 1 | 0.0683 | 60S ribosomal protein L23a |
| contig_10429_5 | 0.0412 | 0.0816 | 0.0892 | 0.157 | 0 | 1 | 0.0692 | hypothetical protein T265_11290 partial |
| contig_5617_6 | 0.0413 | 0.0827 | 0.0897 | 0.157 | 0 | 1 | 0.0692 | sarcoplasmic calcium-binding protein |
| contig_21030_6 | 0.0413 | 0.0829 | 0.0907 | 0.157 | 0 | 1 | 0.0692 | 60S ribosomal protein L39 |
| contig_8539_5 | 0.0417 | 0.0829 | 0.0907 | 0.1583 | 0 | 1 | 0.0698 | SET domain-containing protein |
| contig_6012_4 | 0.0419 | 0.0839 | 0.0917 | 0.1583 | 0 | 1 | 0.0698 | V-type proton ATPase subunit B isoform |
| contig_17027_5 | 0.042 | 0.0848 | 0.0926 | 0.1583 | 0 | 1 | 0.0698 | predicted protein |
| contig_40093_3 | 0.0421 | 0.0869 | 0.0931 | 0.1583 | 0 | 1 | 0.0698 | predicted protein |
| contig_15603_6 | 0.0422 | 0.0869 | 0.0933 | 0.1583 | 0 | 1 | 0.0698 | predicted protein |
| contig_14697_5 | 0.0423 | 0.0875 | 0.0938 | 0.1583 | 0 | 1 | 0.0698 | fructose 1 6-bisphosphatase |
| contig_25275_2 | 0.0423 | 0.0877 | 0.0961 | 0.1583 | 0 | 1 | 0.0698 | histidine-tRNA ligase |
| contig_27832_2 | 0.0429 | 0.0893 | 0.0966 | 0.1597 | 0 | 1 | 0.0704 | ---NA--- |

| | | | | | | | | |
|---|---|---|---|---|---|---|---|---|
| contig_35026_4 | 0.0429 | 0.0903 | 0.0966 | 0.1597 | 0 | 1 | 0.0704 | glyceraldehyde 3-phosphate dehydrogenase |
| contig_13566_4 | 0.0435 | 0.0907 | 0.0969 | 0.1616 | 0 | 1 | 0.0712 | 60S ribosomal protein L35a-4 |
| contig_29330_2 | 0.0438 | 0.0915 | 0.0969 | 0.162 | 0 | 1 | 0.0714 | pilus assembly protein |
| contig_5781_6 | 0.0439 | 0.0915 | 0.0969 | 0.162 | 0 | 1 | 0.0714 | acetyl-coenzyme a synthetase |
| contig_16381_6 | 0.0442 | 0.0928 | 0.0987 | 0.1626 | 0 | 1 | 0.0717 | predicted protein |
| contig_17799_3 | 0.0449 | 0.0933 | 0.0989 | 0.165 | 0 | 1 | 0.0728 | light-harvesting protein |
| contig_25763_5 | 0.0451 | 0.0934 | 0.0989 | 0.1654 | 0 | 1 | 0.0729 | Dihydroflavonol 4-reductase |
| contig_29386_5 | 0.0458 | 0.0961 | 0.0996 | 0.1673 | 0 | 1 | 0.0738 | predicted protein |
| contig_24941_5 | 0.0459 | 0.0966 | 0.1 | 0.1673 | 0 | 1 | 0.0738 | triosephosphate isomerase |
| contig_17409_1 | 0.046 | 0.0966 | 0.1008 | 0.1673 | 0 | 1 | 0.0738 | vesicle-associated membrane protein 7b |
| contig_28093_5 | 0.0466 | 0.0969 | 0.1014 | 0.1692 | 0 | 1 | 0.0746 | protein dispatched homolog 1-like |
| contig_22768_6 | 0.0471 | 0.0969 | 0.1015 | 0.1703 | 0 | 1 | 0.0751 | putative LOV domain-containing protein |
| contig_14919_6 | 0.0472 | 0.097 | 0.1016 | 0.1703 | 0 | 1 | 0.0751 | predicted protein |
| contig_28919_4 | 0.0473 | 0.0987 | 0.1029 | 0.1703 | 0 | 1 | 0.0751 | adenosylhomocysteinase |
| contig_28354_5 | 0.0474 | 0.0987 | 0.1029 | 0.1704 | 0 | 1 | 0.0751 | hypothetical protein FRACYDRAFT_169912 |
| contig_23946_2 | 0.0477 | 0.0993 | 0.1061 | 0.1712 | 0 | 1 | 0.0755 | ATP-binding cassette subfamily F member 3 |
| contig_24813_3 | 0.0482 | 0.0999 | 0.1087 | 0.1722 | 0 | 1 | 0.0759 | predicted protein |
| contig_7686_2 | 0.0484 | 0.1 | 0.1091 | 0.1722 | 0 | 1 | 0.0759 | glutathione-disulfide reductase |
| contig_35278_2 | 0.0485 | 0.1008 | 0.1097 | 0.1722 | 0 | 1 | 0.0759 | light harvesting protein |
| contig_32692_5 | 0.0485 | 0.1014 | 0.1101 | 0.1722 | 0 | 1 | 0.0759 | predicted protein |
| contig_9499_6 | 0.0489 | 0.1015 | 0.1115 | 0.1726 | 0 | 1 | 0.0761 | predicted protein |
| contig_35135_5 | 0.0489 | 0.1027 | 0.1121 | 0.1726 | 0 | 1 | 0.0761 | peptidyl-prolyl cis-trans isomerase A |
| contig_28412_6 | 0.0491 | 0.1035 | 0.1148 | 0.1726 | 0 | 1 | 0.0761 | predicted protein |
| contig_28951_4 | 0.0493 | 0.106 | 0.1148 | 0.1726 | 0 | 1 | 0.0761 | predicted protein |
| contig_9006_3 | 0.0494 | 0.1061 | 0.1164 | 0.1726 | 0 | 1 | 0.0761 | thiol-activated cytolysin |
| contig_34911_5 | 0.0496 | 0.1061 | 0.1164 | 0.1726 | 0 | 1 | 0.0761 | predicted protein |
| contig_15547_4 | 0.0497 | 0.1097 | 0.1212 | 0.1726 | 0 | 1 | 0.0761 | hypothetical protein FRACYDRAFT_234640 |

| | | | | | | | | |
|---|---|---|---|---|---|---|---|---|
| contig_29967_ 5 | 0.0497 | 0.1107 | 0.1229 | 0.17 26 | 0 | 1 | 0.07 61 | protein dispatched homolog 1 isoform X2 |
| contig_38390_ 3 | 0.0498 | 0.112 | 0.1232 | 0.17 26 | 0 | 1 | 0.07 61 | predicted protein |
| contig_9763_5 | 0.0498 | 0.1139 | 0.1233 | 0.17 26 | 0 | 1 | 0.07 61 | predicted protein |
| contig_22969_ 5 | 0.0499 | 0.1148 | 0.1253 | 0.17 26 | 0 | 1 | 0.07 61 | molecular chaperone DnaJ |
| contig_22753_ 2 | 0.0499 | 0.116 | 0.1274 | 0.17 26 | 0 | 1 | 0.07 61 | nicotinate phosphoribosyltransferase 2-like |
| contig_40017_ 2 | 0.0506 | 0.1165 | 0.13 | 0.17 46 | 0 | 1 | 0.07 7 | predicted protein |
| contig_32742_ 5 | 0.0513 | 0.1179 | 0.1303 | 0.17 66 | 0 | 1 | 0.07 79 | predicted protein |
| contig_38550_ 1 | 0.0515 | 0.1229 | 0.1306 | 0.17 69 | 0 | 1 | 0.07 8 | polyadenylate-binding protein 1-B |
| contig_7030_6 | 0.052 | 0.1232 | 0.1314 | 0.17 77 | 0 | 1 | 0.07 83 | ---NA--- |
| contig_39623_ 1 | 0.052 | 0.1232 | 0.1314 | 0.17 77 | 0 | 1 | 0.07 83 | predicted protein |
| contig_38505_ 3 | 0.0521 | 0.1256 | 0.1359 | 0.17 77 | 0 | 1 | 0.07 83 | serine/threonine-protein phosphatase 2A regulatory subunit B' |
| contig_19373_ 4 | 0.0529 | 0.13 | 0.1361 | 0.17 99 | 0 | 1 | 0.07 93 | fucoxanthin chlorophyll a/c-binding protein precursor |
| contig_28714_ 6 | 0.0535 | 0.1303 | 0.1365 | 0.18 17 | 0 | 1 | 0.08 01 | predicted protein |
| contig_33687_ 4 | 0.0537 | 0.1305 | 0.1372 | 0.18 19 | 0 | 1 | 0.08 02 | predicted protein |
| contig_24923_ 5 | 0.0548 | 0.1306 | 0.1389 | 0.18 53 | 0 | 1 | 0.08 17 | voltage activated chloride channel CLC7 type |
| contig_9513_6 | 0.055 | 0.1359 | 0.139 | 0.18 54 | 0 | 1 | 0.08 17 | predicted protein |
| contig_20174_ 5 | 0.0551 | 0.1359 | 0.1393 | 0.18 54 | 0 | 1 | 0.08 17 | glutathione S-transferase |
| contig_29868_ 5 | 0.0553 | 0.1359 | 0.1416 | 0.18 54 | 0 | 1 | 0.08 17 | adp-ribosylation factor 1 |
| contig_28463_ 3 | 0.0553 | 0.1365 | 0.1459 | 0.18 54 | 0 | 1 | 0.08 17 | aspartate kinase |
| contig_38064_ 1 | 0.0555 | 0.1372 | 0.1459 | 0.18 54 | 0 | 1 | 0.08 17 | phosphoenolpyruvate carboxylase |
| contig_12446_ 4 | 0.0557 | 0.1372 | 0.1459 | 0.18 57 | 0 | 1 | 0.08 19 | UPF0187 protein |
| contig_35366_ 6 | 0.0558 | 0.139 | 0.1462 | 0.18 57 | 0 | 1 | 0.08 19 | chaperonin GroEL (chloroplast) |
| contig_27719_ 4 | 0.0559 | 0.1405 | 0.1466 | 0.18 57 | 0 | 1 | 0.08 19 | predicted protein |
| contig_21768_ 6 | 0.0562 | 0.1416 | 0.1466 | 0.18 61 | 0 | 1 | 0.08 21 | 40S ribosomal protein |
| contig_10817_ 6 | 0.0564 | 0.1459 | 0.1469 | 0.18 64 | 0 | 1 | 0.08 22 | silicon efflux transporter LSI2 |
| contig_23501_ 5 | 0.0566 | 0.1459 | 0.147 | 0.18 68 | 0 | 1 | 0.08 23 | bile acid:sodium symporter |

| | | | | | | | | |
|---|---|---|---|---|---|---|---|---|
| contig_27196_4 | 0.0572 | 0.1459 | 0.1536 | 0.1881 | 0 | 1 | 0.0829 | predicted protein |
| contig_17402_3 | 0.0573 | 0.1465 | 0.1563 | 0.1881 | 0 | 1 | 0.0829 | hypothetical protein FRACYDRAFT_278091 |
| contig_9372_6 | 0.0575 | 0.1466 | 0.1564 | 0.1885 | 0 | 1 | 0.0831 | predicted protein |
| contig_24919_4 | 0.0578 | 0.1469 | 0.1759 | 0.189 | 0 | 1 | 0.0833 | predicted protein |
| contig_25490_5 | 0.058 | 0.147 | 0.2008 | 0.1891 | 0 | 1 | 0.0834 | ribonucleoside-diphosphate reductase large subunit |
| contig_21155_4 | 0.0581 | 0.147 | 0.2008 | 0.1891 | 0 | 1 | 0.0834 | heat shock 70 kDa protein |
| contig_28186_6 | 0.059 | 0.1563 | 0.2039 | 0.1915 | 0 | 1 | 0.0844 | predicted protein |
| contig_33557_6 | 0.0591 | 0.1564 | 0.2042 | 0.1915 | 0 | 1 | 0.0844 | phosphoglycerate kinase |
| contig_6511_2 | 0.0592 | 0.2038 | 0.2049 | 0.1915 | 0 | 1 | 0.0844 | glutamyl-tRNA synthetase |
| contig_30582_1 | 0.0594 | 0.2039 | 0.2054 | 0.1918 | 0 | 1 | 0.0846 | silicon transporter |
| contig_13881_2 | 0.0598 | 0.2046 | 0.2064 | 0.1918 | 0 | 1 | 0.0846 | predicted protein |
| contig_38598_5 | 0.0598 | 0.2052 | 0.2064 | 0.1918 | 0 | 1 | 0.0846 | Adenylyl cyclase-associated protein |
| contig_38023_2 | 0.0599 | 0.2054 | 1 | 0.1918 | 0 | 1 | 0.0846 | predicted protein |
| contig_34164_6 | 0.06 | 0.2069 | 1 | 0.1918 | 0 | 1 | 0.0846 | multispanning membrane protein |
| contig_4611_6 | 0.06 | 1 | 1 | 0.1918 | 0 | 1 | 0.0846 | long chain acyl-coa synthetase |
| contig_24988_5 | 0.0606 | 1 | 1 | 0.1932 | 1E-04 | 1 | 0.0852 | Urea-proton symporter DUR3 |
| contig_25500_5 | 0.0607 | 1 | 1 | 0.1932 | 1E-04 | 1 | 0.0852 | predicted protein |
| contig_33630_6 | 0.061 | 1 | 1 | 0.1936 | 1E-04 | 1 | 0.0854 | actin |
| contig_23751_6 | 0.0616 | 1 | 1 | 0.1947 | 1E-04 | 1 | 0.0858 | high mobility group protein B2 |
| contig_40859_3 | 0.0616 | 1 | 1 | 0.1947 | 1E-04 | 1 | 0.0858 | predicted protein |
| contig_24799_5 | 0.062 | 1 | 1 | 0.1957 | 1E-04 | 1 | 0.0863 | predicted protein |
| contig_7075_5 | 0.0623 | 1 | 1 | 0.1962 | 2E-04 | 1 | 0.0865 | predicted protein |
| contig_6933_4 | 0.0625 | 1 | 1 | 0.1963 | 2E-04 | 1 | 0.0865 | methylmalonate-semialdehyde dehydrogenase (acylating) |
| contig_27544_6 | 0.0627 | 1 | 1 | 0.1967 | 3E-04 | 1 | 0.0867 | Long-chain-fatty-acid--CoA ligase putative |
| contig_16000_5 | 0.0632 | 1 | 1 | 0.1976 | 3E-04 | 1 | 0.0871 | FAD-dependent oxidoreductase |
| contig_36049_1 | 0.0633 | 1 | 1 | 0.1976 | 4E-04 | 1 | 0.0871 | predicted protein |
| contig_12155_4 | 0.0634 | 1 | 1 | 0.1976 | 4E-04 | 1 | 0.0871 | predicted protein |

| | | | | | | | | |
|---|---|---|---|---|---|---|---|---|
| contig_39034_5 | 0.0637 | 1 | 1 | 0.198 | 5E-04 | 1 | 0.0873 | predicted protein |
| contig_10461_4 | 0.0639 | 1 | 1 | 0.198 | 6E-04 | 1 | 0.0873 | ATP synthase subunit beta mitochondrial |
| contig_30013_1 | 0.064 | 1 | 1 | 0.198 | 7E-04 | 1 | 0.0873 | ribonucleoside-diphosphate reductase subunit alpha |
| contig_28089_3 | 0.064 | 1 | 1 | 0.198 | 9E-04 | 1 | 0.0873 | threonine synthase |
| contig_40270_3 | 0.0669 | 1 | 1 | 0.2063 | 0.001 | 1 | 0.091 | predicted protein |
| contig_26589_3 | 0.0672 | 1 | 1 | 0.2069 | 0.0012 | 1 | 0.0912 | predicted protein |
| contig_34452_4 | 0.0674 | 1 | 1 | 0.2071 | 0.0014 | 1 | 0.0913 | glyceraldehyde-3-phosphate dehydrogenase precursor |
| contig_27705_5 | 0.0676 | 1 | 1 | 0.2072 | 0.0017 | 1 | 0.0914 | predicted protein |
| contig_38361_6 | 0.068 | 1 | 1 | 0.208 | 0.0019 | 1 | 0.0917 | 60S ribosomal protein L7a |
| contig_27589_4 | 0.069 | 1 | 1 | 0.2106 | 0.0023 | 1 | 0.0929 | nucleolar protein 4 |
| contig_29981_4 | 0.0696 | 1 | 1 | 0.2121 | 0.0026 | 1 | 0.0935 | 20S proteasome subunit alpha 4 |
| contig_39202_4 | 0.0697 | 1 | 1 | 0.2121 | 0.003 | 1 | 0.0935 | Bystin |
| contig_25542_6 | 0.0699 | 1 | 1 | 0.2123 | 0.0035 | 1 | 0.0936 | ---NA--- |
| contig_23910_2 | 0.0701 | 1 | 1 | 0.2124 | 0.004 | 1 | 0.0937 | ARM repeat-containing protein |
| contig_25957_2 | 0.0705 | 1 | 1 | 0.2132 | 0.0047 | 1 | 0.094 | predicted protein |
| contig_25014_3 | 0.071 | 1 | 1 | 0.2138 | 0.0053 | 1 | 0.0943 | predicted protein |
| contig_15954_4 | 0.071 | 1 | 1 | 0.2138 | 0.0061 | 1 | 0.0943 | elongation factor-like |
| contig_9649_2 | 0.0713 | 1 | 1 | 0.2144 | 0.007 | 1 | 0.0945 | predicted protein |
| contig_15496_6 | 0.0715 | 1 | 1 | 0.2144 | 0.008 | 1 | 0.0945 | catalase |
| contig_6981_5 | 0.0721 | 1 | 1 | 0.2159 | 0.0091 | 1 | 0.0952 | predicted protein |
| contig_27675_6 | 0.0728 | 1 | 1 | 0.2175 | 0.0103 | 1 | 0.0959 | inosine-5'-monophosphate dehydrogenase |
| contig_27595_5 | 0.0731 | 1 | 1 | 0.2179 | 0.0117 | 1 | 0.0961 | transitional endoplasmic reticulum ATPase |
| contig_8049_3 | 0.0734 | 1 | 1 | 0.2184 | 0.0133 | 1 | 0.0963 | predicted protein |
| contig_40568_2 | 0.0736 | 1 | 1 | 0.2187 | 0.015 | 1 | 0.0964 | glutathione reductase |
| contig_33594_1 | 0.0739 | 1 | 1 | 0.2191 | 0.0168 | 1 | 0.0966 | CULlin protein 1 |
| contig_32601_3 | 0.0748 | 1 | 1 | 0.2214 | 0.0189 | 1 | 0.0976 | predicted protein |
| contig_25871_6 | 0.0753 | 1 | 1 | 0.2223 | 0.0212 | 1 | 0.098 | RNA-binding domain-containing protein |

| | | | | | | | | |
|---|---|---|---|---|---|---|---|---|
| contig_10223_6 | 0.0755 | 1 | 1 | 0.2225 | 0.0237 | 1 | 0.0981 | predicted protein |
| contig_22140_2 | 0.0767 | 1 | 1 | 0.2256 | 0.0265 | 1 | 0.0995 | predicted protein |
| contig_24114_3 | 0.0773 | 1 | 1 | 0.2271 | 0.0295 | 1 | 0.1001 | predicted protein |
| contig_22651_1 | 0.0781 | 1 | 1 | 0.2287 | 0.0327 | 1 | 0.1009 | glutamate synthase subunit alpha |
| contig_38588_6 | 0.0782 | 1 | 1 | 0.2289 | 0.0363 | 1 | 0.1009 | Ras-related protein Rab-8A |
| contig_32710_4 | 0.0789 | 1 | 1 | 0.23 | 0.0401 | 1 | 0.1014 | class I SAM-dependent methyltransferase |
| contig_40102_2 | 0.0789 | 1 | 1 | 0.23 | 0.0442 | 1 | 0.1014 | predicted protein |
| contig_34173_4 | 0.0794 | 1 | 1 | 0.2307 | 0.0487 | 1 | 0.1017 | succinate dehydrogenase flavoprotein subunit |
| contig_25429_4 | 0.0795 | 1 | 1 | 0.2307 | 0.0536 | 1 | 0.1017 | V-type proton ATPase proteolipid subunit |
| contig_8868_5 | 0.0796 | 1 | 1 | 0.2307 | 0.0588 | 1 | 0.1017 | Enoyl-CoA hydratase |
| contig_29925_5 | 0.08 | 1 | 1 | 0.231 | 0.0644 | 1 | 0.1019 | glutamine synthetase |
| contig_25890_5 | 0.0801 | 1 | 1 | 0.231 | 0.0704 | 1 | 0.1019 | acetolactate synthase catalytic subunit |
| contig_9050_3 | 0.0805 | 1 | 1 | 0.2311 | 0.0769 | 1 | 0.1019 | triose phosphate/phosphate translocator |
| contig_40261_6 | 0.0806 | 1 | 1 | 0.2311 | 0.0837 | 1 | 0.1019 | predicted protein |
| contig_30644_2 | 0.0808 | 1 | 1 | 0.2311 | 0.0911 | 1 | 0.1019 | predicted protein |
| contig_25987_5 | 0.0808 | 1 | 1 | 0.2311 | 0.0989 | 1 | 0.1019 | polyadenylate-binding protein 1-B |
| contig_28753_4 | 0.081 | 1 | 1 | 0.2311 | 0.1072 | 1 | 0.1019 | FKBP-type peptidyl-prolyl cis-trans isomerase |
| contig_34246_6 | 0.081 | 1 | 1 | 0.2311 | 0.116 | 1 | 0.1019 | homoserine kinase |
| contig_22713_2 | 0.0813 | 1 | 1 | 0.2311 | 0.1253 | 1 | 0.1019 | predicted protein |
| contig_9038_6 | 0.0814 | 1 | 1 | 0.2311 | 0.1351 | 1 | 0.1019 | methionine-tRNA ligase beta subunit |
| contig_25670_6 | 0.0814 | 1 | 1 | 0.2311 | 0.1454 | 1 | 0.1019 | predicted protein |
| contig_11877_6 | 0.0816 | 1 | 1 | 0.2312 | 0.1563 | 1 | 0.1019 | 30S ribosomal protein S5 |
| contig_29625_1 | 0.0825 | 1 | 1 | 0.233 | 0.1677 | 1 | 0.1027 | rRNA 2'-O-methyltransferase fibrillarin |
| contig_15153_4 | 0.0826 | 1 | 1 | 0.233 | 0.1796 | 1 | 0.1027 | Ras-like GTP-binding protein YPT1 |
| contig_22636_2 | 0.0827 | 1 | 1 | 0.233 | 0.192 | 1 | 0.1027 | 60s acidic ribosomal protein p0 |
| contig_5987_6 | 0.0829 | 1 | 1 | 0.233 | 0.2049 | 1 | 0.1027 | predicted protein |
| contig_32774_1 | 0.0831 | 1 | 1 | 0.2333 | 0.2183 | 1 | 0.1028 | 3-deoxy-7-phosphoheptulonate synthase |

| | | | | | | | | |
|---|---|---|---|---|---|---|---|---|
| contig_23694_6 | 0.0833 | 1 | 1 | 0.2333 | 0.2323 | 1 | 0.1028 | cell division cycle protein 123 homolog |
| contig_34535_1 | 0.0838 | 1 | 1 | 0.2343 | 0.2467 | 1 | 0.1033 | eukaryotic translation initiation factor 2 subunit 3 |
| contig_33536_4 | 0.0839 | 1 | 1 | 0.2343 | 0.2615 | 1 | 0.1033 | sec1 family transporter sly1 |
| contig_37607_6 | 0.0847 | 1 | 1 | 0.2359 | 0.2768 | 1 | 0.104 | 2-isopropylmalate synthase |
| contig_34979_6 | 0.0848 | 1 | 1 | 0.2359 | 0.2925 | 1 | 0.104 | ubiquitin-like protein |
| contig_9641_3 | 0.0857 | 1 | 1 | 0.238 | 0.3086 | 1 | 0.1049 | predicted protein |
| contig_24220_6 | 0.0866 | 1 | 1 | 0.2395 | 0.3249 | 1 | 0.1056 | ATP-binding cassette superfamily |
| contig_789_5 | 0.0867 | 1 | 1 | 0.2395 | 0.3416 | 1 | 0.1056 | ribulose-1 5-bisphosphate carboxylase/oxygenase large subunit (chloroplast) |
| contig_16696_5 | 0.0869 | 1 | 1 | 0.2395 | 0.3585 | 1 | 0.1056 | pyruvate kinase |
| contig_22474_6 | 0.0869 | 1 | 1 | 0.2395 | 0.3758 | 1 | 0.1056 | succinyl-CoA ligase beta-chain mitochondrial precursor |
| contig_41222_3 | 0.0872 | 1 | 1 | 0.2399 | 0.3933 | 1 | 0.1058 | predicted protein |
| contig_7641_5 | 0.0875 | 1 | 1 | 0.2402 | 0.4111 | 1 | 0.1059 | precursor cytochrome c6 |
| contig_6069_1 | 0.0877 | 1 | 1 | 0.2404 | 0.429 | 1 | 0.106 | GLNA glutamine synthase |
| contig_11948_3 | 0.0885 | 1 | 1 | 0.2422 | 0.4471 | 1 | 0.1068 | aldo/keto reductase |
| contig_27259_5 | 0.0888 | 1 | 1 | 0.2424 | 0.4652 | 1 | 0.1069 | ATP synthase F1 gamma subunit |
| contig_33444_6 | 0.0892 | 1 | 1 | 0.2429 | 0.4834 | 1 | 0.1071 | predicted protein |
| contig_3037_3 | 0.0893 | 1 | 1 | 0.2429 | 0.5016 | 1 | 0.1071 | ribosomal protein S4 (chloroplast) |
| contig_39771_2 | 0.0897 | 1 | 1 | 0.2437 | 0.5198 | 1 | 0.1074 | hypothetical protein THAOC_36769 |
| contig_10116_3 | 0.0903 | 1 | 1 | 0.2449 | 0.538 | 1 | 0.108 | peptidyl-prolyl cis-trans isomerase CYP19-4 |
| contig_24447_4 | 0.0907 | 1 | 1 | 0.2451 | 0.556 | 1 | 0.1081 | diaminopimelate decarboxylase |
| contig_22011_5 | 0.0907 | 1 | 1 | 0.2451 | 0.5739 | 1 | 0.1081 | argininosuccinate synthase |
| contig_23369_5 | 0.0911 | 1 | 1 | 0.2457 | 0.5917 | 1 | 0.1083 | predicted protein |
| contig_12101_5 | 0.0915 | 1 | 1 | 0.2463 | 0.6092 | 1 | 0.1086 | ---NA--- |
| contig_5862_2 | 0.0917 | 1 | 1 | 0.2463 | 0.6266 | 1 | 0.1086 | arylsulfatase |
| contig_17285_6 | 0.0924 | 1 | 1 | 0.248 | 0.6437 | 1 | 0.1093 | 40S ribosomal protein S27-4 putative |
| contig_14481_1 | 0.0926 | 1 | 1 | 0.2481 | 0.6604 | 1 | 0.1094 | CDP-diacylglycerol--glycerol-3-phosphate 3-phosphatidyltransferase |

| | | | | | | | | |
|---|---|---|---|---|---|---|---|---|
| contig_33426_<br>1 | 0.0928 | 1 | 1 | 0.24<br>81 | 0.676<br>9 | 1 | 0.10<br>94 | predicted protein |
| contig_16777_<br>4 | 0.0931 | 1 | 1 | 0.24<br>84 | 0.693 | 1 | 0.10<br>95 | H - or Na -translocating F-type V-type and A-type ATPase (F-ATPase) Superfamily |
| contig_21909_<br>6 | 0.0933 | 1 | 1 | 0.24<br>84 | 0.708<br>7 | 1 | 0.10<br>95 | 26S proteasome beta type 7 subunit |
| contig_23921_<br>2 | 0.0934 | 1 | 1 | 0.24<br>84 | 0.724<br>1 | 1 | 0.10<br>95 | predicted protein |
| contig_4841_4 | 0.0938 | 1 | 1 | 0.24<br>89 | 0.739<br>1 | 1 | 0.10<br>97 | 40S ribosomal protein S14 |
| contig_15891_<br>1 | 0.0943 | 1 | 1 | 0.25 | 0.753<br>6 | 1 | 0.11<br>02 | predicted protein |
| contig_27659_<br>6 | 0.0956 | 1 | 1 | 0.25<br>26 | 0.767<br>6 | 1 | 0.11<br>14 | predicted protein |
| contig_29172_<br>6 | 0.0961 | 1 | 1 | 0.25<br>26 | 0.781<br>1 | 1 | 0.11<br>14 | arginyl-tRNA synthetase |
| contig_24139_<br>1 | 0.0961 | 1 | 1 | 0.25<br>26 | 0.794<br>1 | 1 | 0.11<br>14 | chaperonin GroEL |
| contig_32986_<br>4 | 0.0965 | 1 | 1 | 0.25<br>26 | 0.806<br>7 | 1 | 0.11<br>14 | protochlorophyllide reductase magnesium chelatase subunit (chloroplast) |
| contig_11991_<br>5 | 0.0966 | 1 | 1 | 0.25<br>26 | 0.818<br>8 | 1 | 0.11<br>14 | ---NA--- |
| contig_24980_<br>4 | 0.0966 | 1 | 1 | 0.25<br>26 | 0.830<br>4 | 1 | 0.11<br>14 | Bystin |
| contig_16094_<br>5 | 0.0969 | 1 | 1 | 0.25<br>26 | 0.841<br>5 | 1 | 0.11<br>14 | 40S ribosomal protein S23 |
| contig_25552_<br>1 | 0.0969 | 1 | 1 | 0.25<br>26 | 0.852<br>2 | 1 | 0.11<br>14 | predicted protein |
| contig_37718_<br>5 | 0.0969 | 1 | 1 | 0.25<br>26 | 0.862<br>4 | 1 | 0.11<br>14 | predicted protein |
| contig_27384_<br>5 | 0.097 | 1 | 1 | 0.25<br>26 | 0.872<br>1 | 1 | 0.11<br>14 | predicted protein |
| contig_32502_<br>6 | 0.0981 | 1 | 1 | 0.25<br>52 | 0.881<br>3 | 1 | 0.11<br>25 | predicted protein |
| contig_27149_<br>6 | 0.0987 | 1 | 1 | 0.25<br>55 | 0.89 | 1 | 0.11<br>27 | homogentisate phytyltransferase |
| contig_22479_<br>1 | 0.0987 | 1 | 1 | 0.25<br>55 | 0.898<br>2 | 1 | 0.11<br>27 | predicted protein |
| contig_27190_<br>6 | 0.0989 | 1 | 1 | 0.25<br>55 | 0.906 | 1 | 0.11<br>27 | 20S proteasome subunit alpha 2 |
| contig_36895_<br>3 | 0.0989 | 1 | 1 | 0.25<br>55 | 0.913<br>3 | 1 | 0.11<br>27 | phenylalanine-tRNA ligase beta subunit (chloroplast) |
| contig_30620_<br>5 | 0.0993 | 1 | 1 | 0.25<br>59 | 0.920<br>2 | 1 | 0.11<br>28 | Transmembrane protein 87B |
| contig_10727_<br>3 | 0.0996 | 1 | 1 | 0.25<br>63 | 0.926<br>7 | 1 | 0.11<br>3 | eukaryotic translation initiation factor 3 subunit G |
| contig_27592_<br>6 | 0.0999 | 1 | 1 | 0.25<br>66 | 0.932<br>8 | 1 | 0.11<br>31 | zeta-carotene desaturase |
| contig_28417_<br>5 | 0.1 | 1 | 1 | 0.25<br>66 | 0.938<br>4 | 1 | 0.11<br>31 | predicted protein |
| contig_28970_<br>6 | 0.1007 | 1 | 1 | 0.25<br>77 | 0.943<br>7 | 1 | 0.11<br>36 | peptide methionine sulfoxide reductase |

| | | | | | | | | |
|---|---|---|---|---|---|---|---|---|
| contig_6711_4 | 0.1008 | 1 | 1 | 0.2578 | 0.9487 | 1 | 0.1136 | predicted protein |
| contig_30318_6 | 0.1012 | 1 | 1 | 0.2579 | 0.9532 | 1 | 0.1137 | pyruvate dehydrogenase complex dihydrolipoamide acetyltransferase |
| contig_28743_4 | 0.1014 | 1 | 1 | 0.2579 | 0.9575 | 1 | 0.1137 | chlorophyll a/b-binding protein |
| contig_14595_3 | 0.1015 | 1 | 1 | 0.2579 | 0.9614 | 1 | 0.1137 | predicted protein |
| contig_11147_6 | 0.1016 | 1 | 1 | 0.2579 | 0.965 | 1 | 0.1137 | predicted protein |
| contig_27901_2 | 0.1026 | 1 | 1 | 0.2599 | 0.9684 | 1 | 0.1146 | methionyl-trna formyltransferase |
| contig_10952_4 | 0.1027 | 1 | 1 | 0.2599 | 0.9714 | 1 | 0.1146 | diacylglycerol O-acyltransferase 2D |
| contig_40308_6 | 0.1029 | 1 | 1 | 0.2599 | 0.9742 | 1 | 0.1146 | 40S ribosomal protein S6 |
| contig_30106_6 | 0.1035 | 1 | 1 | 0.261 | 0.9768 | 1 | 0.1151 | predicted protein |
| contig_8777_3 | 0.1041 | 1 | 1 | 0.2622 | 0.9792 | 1 | 0.1156 | elongation factor G |
| contig_12284_5 | 0.1047 | 1 | 1 | 0.2632 | 0.9813 | 1 | 0.116 | ribosomal protein S9 |
| contig_17187_5 | 0.1051 | 1 | 1 | 0.2637 | 0.9833 | 1 | 0.1163 | small ubiquitin-like modifier |
| contig_27039_5 | 0.1058 | 1 | 1 | 0.265 | 0.9851 | 1 | 0.1169 | hypothetical protein THAOC_36140 |
| contig_38543_1 | 0.106 | 1 | 1 | 0.265 | 0.9867 | 1 | 0.1169 | predicted protein |
| contig_8757_2 | 0.1061 | 1 | 1 | 0.265 | 0.9881 | 1 | 0.1169 | ABC transporter |
| contig_8125_5 | 0.1068 | 1 | 1 | 0.2663 | 0.9894 | 1 | 0.1174 | adenosylhomocysteinase |
| contig_9464_6 | 0.108 | 1 | 1 | 0.2689 | 0.9906 | 1 | 0.1185 | cytochrome c heme attachment protein (chloroplast) |
| contig_10045_6 | 0.1085 | 1 | 1 | 0.2695 | 0.9917 | 1 | 0.1188 | dihydrolipoamide dehydrogenase |
| contig_6766_1 | 0.1087 | 1 | 1 | 0.2697 | 0.9926 | 1 | 0.1189 | N-carbamoylputrescine amidase |
| contig_32739_1 | 0.109 | 1 | 1 | 0.2699 | 0.9935 | 1 | 0.119 | fibronectin type III domain-containing protein |
| contig_34816_5 | 0.1091 | 1 | 1 | 0.2699 | 0.9942 | 1 | 0.119 | 40S ribosomal protein S12 |
| contig_29498_6 | 0.1094 | 1 | 1 | 0.2702 | 0.9949 | 1 | 0.1191 | aldehyde dehydrogenase |
| contig_27547_5 | 0.1097 | 1 | 1 | 0.2705 | 0.9955 | 1 | 0.1192 | ribulose-phosphate 3-epimerase |
| contig_25733_5 | 0.1101 | 1 | 1 | 0.271 | 0.9961 | 1 | 0.1195 | predicted protein |
| contig_29877_2 | 0.1107 | 1 | 1 | 0.272 | 0.9966 | 1 | 0.1199 | predicted protein |
| contig_29387_4 | 0.1114 | 1 | 1 | 0.2731 | 0.997 | 1 | 0.1204 | predicted protein |
| contig_31457_1 | 0.1115 | 1 | 1 | 0.2731 | 0.9974 | 1 | 0.1204 | predicted protein |

| | | | | | | | | |
|---|---|---|---|---|---|---|---|---|
| contig_38793_1 | 0.112 | 1 | 1 | 0.2737 | 0.9977 | 1 | 0.1207 | predicted protein |
| contig_27237_6 | 0.1121 | 1 | 1 | 0.2737 | 0.998 | 1 | 0.1207 | translation initiation factor 5B |
| contig_25680_5 | 0.1136 | 1 | 1 | 0.2767 | 0.9983 | 1 | 0.122 | glucose-6-phosphate isomerase |
| contig_32573_1 | 0.1137 | 1 | 1 | 0.2767 | 0.9985 | 1 | 0.122 | ATP synthase CF1 alpha subunit (chloroplast) |
| contig_19964_6 | 0.1139 | 1 | 1 | 0.2767 | 0.9987 | 1 | 0.122 | predicted protein |
| contig_30668_1 | 0.1146 | 1 | 1 | 0.2776 | 0.9989 | 1 | 0.1224 | predicted protein |
| contig_13084_2 | 0.1148 | 1 | 1 | 0.2776 | 0.999 | 1 | 0.1224 | 60S ribosomal protein L14 putative |
| contig_37905_5 | 0.1148 | 1 | 1 | 0.2776 | 0.9992 | 1 | 0.1224 | ---NA--- |
| contig_9284_2 | 0.1158 | 1 | 1 | 0.2796 | 0.9993 | 1 | 0.1233 | 60S ribosomal protein L12 |
| contig_36394_4 | 0.116 | 1 | 1 | 0.2796 | 0.9994 | 1 | 0.1233 | Fucoxanthin-chlorophyll a-c binding protein F chloroplastic |
| contig_18580_6 | 0.1164 | 1 | 1 | 0.2798 | 0.9995 | 1 | 0.1234 | predicted protein |
| contig_26948_5 | 0.1165 | 1 | 1 | 0.2798 | 0.9995 | 1 | 0.1234 | predicted protein |
| contig_29801_5 | 0.117 | 1 | 1 | 0.2807 | 0.9996 | 1 | 0.1237 | triosephosphate isomerase |
| contig_19770_6 | 0.1176 | 1 | 1 | 0.2816 | 0.9997 | 1 | 0.1242 | predicted protein |
| contig_27634_3 | 0.1179 | 1 | 1 | 0.2819 | 0.9997 | 1 | 0.1243 | eukaryotic translation initiation factor 3 subunit E |
| contig_39276_6 | 0.1191 | 1 | 1 | 0.2844 | 0.9998 | 1 | 0.1254 | orotidine 5'-phosphate decarboxylase |
| contig_5445_2 | 0.1196 | 1 | 1 | 0.2852 | 0.9998 | 1 | 0.1258 | asparagine synthase B |
| contig_21411_6 | 0.1204 | 1 | 1 | 0.2866 | 0.9998 | 1 | 0.1264 | ATP-dependent RNA helicase eIF4A |
| contig_25207_1 | 0.1209 | 1 | 1 | 0.2872 | 0.9999 | 1 | 0.1266 | predicted protein |
| contig_6178_4 | 0.1212 | 1 | 1 | 0.2876 | 0.9999 | 1 | 0.1268 | rubisco expression protein (chloroplast) |
| contig_23603_4 | 0.1221 | 1 | 1 | 0.2893 | 0.9999 | 1 | 0.1275 | serine protease |
| contig_35278_1 | 0.1224 | 1 | 1 | 0.2896 | 0.9999 | 1 | 0.1277 | light harvesting protein |
| contig_30690_3 | 0.1227 | 1 | 1 | 0.2898 | 0.9999 | 1 | 0.1278 | phosphoglycerate mutase |
| contig_34797_4 | 0.1229 | 1 | 1 | 0.2898 | 0.9999 | 1 | 0.1278 | GTP-binding protein SAR1 |
| contig_6054_4 | 0.1232 | 1 | 1 | 0.2898 | 0.9999 | 1 | 0.1278 | ---NA--- |
| contig_24050_3 | 0.1233 | 1 | 1 | 0.2898 | 1 | 1 | 0.1278 | enolase-phosphatase E1 |
| contig_13388_5 | 0.1238 | 1 | 1 | 0.2906 | 1 | 1 | 0.1281 | ornithine cyclodeaminase |

| | | | | | | | | |
|---|---|---|---|---|---|---|---|---|
| contig_7979_4 | 0.1243 | 1 | 1 | 0.2913 | 1 | 1 | 0.1284 | protein fucoxanthin chlorophyll a/c protein |
| contig_24750_1 | 0.1251 | 1 | 1 | 0.2927 | 1 | 1 | 0.1291 | AP-1 complex subunit beta-1 |
| contig_8482_6 | 0.1253 | 1 | 1 | 0.2929 | 1 | 1 | 0.1291 | UV excision repair protein RAD23 |
| contig_26227_3 | 0.1256 | 1 | 1 | 0.2931 | 1 | 1 | 0.1292 | 3-phosphoshikimate 1-carboxyvinyltransferase |
| contig_10238_6 | 0.1264 | 1 | 1 | 0.2944 | 1 | 1 | 0.1298 | ras-like gtp-binding protein ypt1 |
| contig_16905_4 | 0.1272 | 1 | 1 | 0.2958 | 1 | 1 | 0.1304 | predicted protein |
| contig_10857_4 | 0.1274 | 1 | 1 | 0.2959 | 1 | 1 | 0.1305 | predicted protein |
| contig_21525_6 | 0.1283 | 1 | 1 | 0.2974 | 1 | 1 | 0.1311 | probable small nuclear ribonucleoprotein Sm D2 |
| contig_25907_5 | 0.1289 | 1 | 1 | 0.2984 | 1 | 1 | 0.1316 | predicted protein |
| contig_41159_3 | 0.1299 | 1 | 1 | 0.2996 | 1 | 1 | 0.1321 | predicted protein |
| contig_15393_2 | 0.13 | 1 | 1 | 0.2996 | 1 | 1 | 0.1321 | predicted protein |
| contig_41171_6 | 0.13 | 1 | 1 | 0.2996 | 1 | 1 | 0.1321 | ADP/ATP transporter on adenylate translocase putative |
| contig_3295_5 | 0.1303 | 1 | 1 | 0.2996 | 1 | 1 | 0.1321 | V-type ATPase F subunit |
| contig_17562_5 | 0.1305 | 1 | 1 | 0.2996 | 1 | 1 | 0.1321 | patched domain-containing protein 3-like |
| contig_25214_5 | 0.1306 | 1 | 1 | 0.2996 | 1 | 1 | 0.1321 | predicted protein |
| contig_40248_2 | 0.1313 | 1 | 1 | 0.3004 | 1 | 1 | 0.1325 | predicted protein |
| contig_32573_2 | 0.1314 | 1 | 1 | 0.3004 | 1 | 1 | 0.1325 | ATP synthase CF1 alpha subunit (chloroplast) |
| contig_30840_2 | 0.1328 | 1 | 1 | 0.3032 | 1 | 1 | 0.1337 | 60S ribosomal protein L18a |
| contig_15258_2 | 0.1333 | 1 | 1 | 0.304 | 1 | 1 | 0.134 | predicted protein |
| contig_22940_5 | 0.1348 | 1 | 1 | 0.3067 | 1 | 1 | 0.1352 | PH domain and leucine-rich repeat-containing protein phosphatase |
| contig_24297_3 | 0.1349 | 1 | 1 | 0.3067 | 1 | 1 | 0.1352 | elongation factor Tu |
| contig_14496_5 | 0.1351 | 1 | 1 | 0.3067 | 1 | 1 | 0.1352 | Diaminopimelate epimerase |
| contig_23942_6 | 0.1357 | 1 | 1 | 0.3067 | 1 | 1 | 0.1352 | sister-chromatid cohesion protein 3 |
| contig_27467_4 | 0.1359 | 1 | 1 | 0.3067 | 1 | 1 | 0.1352 | histidinol-phosphate aminotransferase |
| contig_17829_4 | 0.1359 | 1 | 1 | 0.3067 | 1 | 1 | 0.1352 | predicted protein |
| contig_28068_1 | 0.1359 | 1 | 1 | 0.3067 | 1 | 1 | 0.1352 | UDP-glucose 6-dehydrogenase |
| contig_7643_4 | 0.1361 | 1 | 1 | 0.3067 | 1 | 1 | 0.1352 | ATP synthase CF1 beta subunit (chloroplast) |

| | | | | | | | | |
|---|---|---|---|---|---|---|---|---|
| contig_35394_6 | 0.1365 | 1 | 1 | 0.3072 | 1 | 1 | 0.1354 | cytochrome c |
| contig_32280_2 | 0.1369 | 1 | 1 | 0.3075 | 1 | 1 | 0.1356 | plastid sedoheptulose-1 7-bisphosphatase |
| contig_16274_4 | 0.1372 | 1 | 1 | 0.3076 | 1 | 1 | 0.1356 | predicted protein |
| contig_19428_2 | 0.1373 | 1 | 1 | 0.3076 | 1 | 1 | 0.1356 | chloroplast beta-keto acyl reductase |
| contig_14784_2 | 0.1384 | 1 | 1 | 0.3095 | 1 | 1 | 0.1365 | atp-dependent RNA helicase |
| contig_35155_5 | 0.1389 | 1 | 1 | 0.3095 | 1 | 1 | 0.1365 | fucoxanthin chlorophyll a/c protein 8 |
| contig_38515_1 | 0.1389 | 1 | 1 | 0.3095 | 1 | 1 | 0.1365 | predicted protein |
| contig_24223_6 | 0.139 | 1 | 1 | 0.3095 | 1 | 1 | 0.1365 | predicted protein |
| contig_24960_3 | 0.1393 | 1 | 1 | 0.3098 | 1 | 1 | 0.1366 | predicted protein |
| contig_22594_4 | 0.1403 | 1 | 1 | 0.3114 | 1 | 1 | 0.1373 | magnesium chelatase subunit H |
| contig_15143_1 | 0.1405 | 1 | 1 | 0.3114 | 1 | 1 | 0.1373 | CK1/CK1/CK1-D protein kinase variant |
| contig_32617_4 | 0.141 | 1 | 1 | 0.3123 | 1 | 1 | 0.1377 | VPLPA-CTERM protein sorting domain-containing protein |
| contig_27953_1 | 0.1415 | 1 | 1 | 0.3127 | 1 | 1 | 0.1379 | ribosomal protein L23 putative |
| contig_23961_3 | 0.1416 | 1 | 1 | 0.3127 | 1 | 1 | 0.1379 | ribosomal protein S15 |
| contig_11114_6 | 0.1424 | 1 | 1 | 0.3139 | 1 | 1 | 0.1384 | predicted protein |
| contig_11344_5 | 0.1435 | 1 | 1 | 0.3159 | 1 | 1 | 0.1393 | Iron-sulfur cluster co-chaperone protein HscB mitochondrial |
| contig_23311_3 | 0.1439 | 1 | 1 | 0.3164 | 1 | 1 | 0.1395 | predicted protein |
| contig_11923_6 | 0.1451 | 1 | 1 | 0.3176 | 1 | 1 | 0.14 | aspartate aminotransferase |
| contig_39581_5 | 0.1453 | 1 | 1 | 0.3176 | 1 | 1 | 0.14 | cytochrome c1 |
| contig_26111_3 | 0.1457 | 1 | 1 | 0.3176 | 1 | 1 | 0.14 | predicted protein |
| contig_24825_6 | 0.1459 | 1 | 1 | 0.3176 | 1 | 1 | 0.14 | predicted protein |
| contig_30596_2 | 0.1459 | 1 | 1 | 0.3176 | 1 | 1 | 0.14 | predicted protein |
| contig_26174_3 | 0.1459 | 1 | 1 | 0.3176 | 1 | 1 | 0.14 | predicted protein |
| contig_35273_1 | 0.1462 | 1 | 1 | 0.3176 | 1 | 1 | 0.14 | Dethiobiotin synthase |
| contig_41167_2 | 0.1465 | 1 | 1 | 0.3176 | 1 | 1 | 0.14 | oxygen-evolving enhancer protein 1 precursor |
| contig_28795_5 | 0.1466 | 1 | 1 | 0.3176 | 1 | 1 | 0.14 | protein transport protein SEC23 |
| contig_12639_1 | 0.1466 | 1 | 1 | 0.3176 | 1 | 1 | 0.14 | ADP-ribosylation factor-related protein 1 |

| | | | | | | | | |
|---|---|---|---|---|---|---|---|---|
| contig_32942_6 | 0.1469 | 1 | 1 | 0.3176 | 1 | 1 | 0.14 | predicted protein |
| contig_28929_5 | 0.147 | 1 | 1 | 0.3176 | 1 | 1 | 0.14 | 1-deoxy-D-xylulose 5-phosphate reductoisomerase |
| contig_17246_6 | 0.1482 | 1 | 1 | 0.3197 | 1 | 1 | 0.1409 | predicted protein |
| contig_30461_3 | 0.1485 | 1 | 1 | 0.32 | 1 | 1 | 0.1411 | 6 7-dimethyl-8-ribityllumazine synthase |
| contig_27280_6 | 0.1491 | 1 | 1 | 0.3207 | 1 | 1 | 0.1414 | dihydrolipoamide dehydrogenase |
| contig_11813_1 | 0.1498 | 1 | 1 | 0.3219 | 1 | 1 | 0.1419 | predicted protein |
| contig_23323_6 | 0.1502 | 1 | 1 | 0.3222 | 1 | 1 | 0.142 | predicted protein |
| contig_19241_5 | 0.1516 | 1 | 1 | 0.3244 | 1 | 1 | 0.143 | ---NA--- |
| contig_30867_1 | 0.1517 | 1 | 1 | 0.3244 | 1 | 1 | 0.143 | Light-harvesting complex I LH38 protein |
| contig_15461_5 | 0.1523 | 1 | 1 | 0.3253 | 1 | 1 | 0.1434 | predicted protein |
| contig_9759_1 | 0.1531 | 1 | 1 | 0.3265 | 1 | 1 | 0.144 | clathrin heavy chain |
| contig_24729_6 | 0.1533 | 1 | 1 | 0.3265 | 1 | 1 | 0.144 | predicted protein |
| contig_23006_1 | 0.1536 | 1 | 1 | 0.3268 | 1 | 1 | 0.1441 | predicted protein |
| contig_7982_4 | 0.1541 | 1 | 1 | 0.3274 | 1 | 1 | 0.1444 | photosystem II protein V (chloroplast) |
| contig_36190_5 | 0.155 | 1 | 1 | 0.3288 | 1 | 1 | 0.145 | long chain acyl-coa synthetase |
| contig_9471_1 | 0.156 | 1 | 1 | 0.33 | 1 | 1 | 0.1455 | photosystem II CP43 chlorophyll apoprotein (chloroplast) |
| contig_26157_4 | 0.1562 | 1 | 1 | 0.33 | 1 | 1 | 0.1455 | predicted protein |
| contig_17548_1 | 0.1563 | 1 | 1 | 0.33 | 1 | 1 | 0.1455 | DUF1772 domain-containing protein |
| contig_15354_3 | 0.1564 | 1 | 1 | 0.33 | 1 | 1 | 0.1455 | predicted protein |
| contig_31150_6 | 0.1594 | 1 | 1 | 0.3357 | 1 | 1 | 0.148 | protein fucoxanthin chlorophyll a/c protein |
| contig_39090_5 | 0.1596 | 1 | 1 | 0.3357 | 1 | 1 | 0.148 | Glycerol-3-phosphate dehydrogenase |
| contig_33684_3 | 0.1604 | 1 | 1 | 0.337 | 1 | 1 | 0.1486 | nadh dehydrogenase |
| contig_13609_2 | 0.1611 | 1 | 1 | 0.3381 | 1 | 1 | 0.1491 | U5 small nuclear ribonucleoprotein 200 kDa helicase |
| contig_26583_1 | 0.1617 | 1 | 1 | 0.3389 | 1 | 1 | 0.1494 | predicted protein |
| contig_27030_2 | 0.1622 | 1 | 1 | 0.3393 | 1 | 1 | 0.1496 | geranylgeranyl reductase |
| contig_14405_3 | 0.1625 | 1 | 1 | 0.3393 | 1 | 1 | 0.1496 | predicted protein |
| contig_28911_6 | 0.1626 | 1 | 1 | 0.3393 | 1 | 1 | 0.1496 | eukaryotic translation initiation factor 6 |

| | | | | | | | | |
|---|---|---|---|---|---|---|---|---|
| contig_24818_1 | 0.1631 | 1 | 1 | 0.3399 | 1 | 1 | 0.1499 | UBC-like protein |
| contig_33218_4 | 0.166 | 1 | 1 | 0.3452 | 1 | 1 | 0.1522 | predicted protein |
| contig_17447_3 | 0.1661 | 1 | 1 | 0.3452 | 1 | 1 | 0.1522 | tetrapyrrole-binding protein |
| contig_32503_1 | 0.1684 | 1 | 1 | 0.3493 | 1 | 1 | 0.154 | predicted protein |
| contig_26083_5 | 0.1687 | 1 | 1 | 0.3493 | 1 | 1 | 0.154 | phosphatidylinositol transfer protein 3-like |
| contig_16247_6 | 0.1688 | 1 | 1 | 0.3493 | 1 | 1 | 0.154 | imidazoleglycerol-phosphate dehydratase |
| contig_7506_5 | 0.1694 | 1 | 1 | 0.3497 | 1 | 1 | 0.1542 | apocytochrome b (mitochondrion) |
| contig_24983_1 | 0.1696 | 1 | 1 | 0.3497 | 1 | 1 | 0.1542 | GTP-binding nuclear protein Ran |
| contig_28744_4 | 0.1697 | 1 | 1 | 0.3497 | 1 | 1 | 0.1542 | ascorbate peroxidase |
| contig_28209_6 | 0.1701 | 1 | 1 | 0.3501 | 1 | 1 | 0.1544 | dihydrolipoamide acetyltransferase |
| contig_6790_6 | 0.1707 | 1 | 1 | 0.3508 | 1 | 1 | 0.1547 | GNAT family N-acetyltransferase |
| contig_12026_5 | 0.1716 | 1 | 1 | 0.3523 | 1 | 1 | 0.1553 | predicted protein |
| contig_28207_5 | 0.1719 | 1 | 1 | 0.3524 | 1 | 1 | 0.1554 | mannose-P-dolichol utilization defect 1 protein |
| contig_37899_3 | 0.1725 | 1 | 1 | 0.3532 | 1 | 1 | 0.1557 | ribosomal protein L2 (chloroplast) |
| contig_9468_6 | 0.1743 | 1 | 1 | 0.3561 | 1 | 1 | 0.157 | anaphase-promoting complex subunit 1 |
| contig_24334_5 | 0.1744 | 1 | 1 | 0.3561 | 1 | 1 | 0.157 | predicted protein |
| contig_28927_1 | 0.1747 | 1 | 1 | 0.3561 | 1 | 1 | 0.157 | 60S ribosomal protein |
| contig_30723_5 | 0.1752 | 1 | 1 | 0.3566 | 1 | 1 | 0.1572 | 20S proteasome subunit beta 1 |
| contig_15137_6 | 0.1755 | 1 | 1 | 0.3566 | 1 | 1 | 0.1572 | aspartate aminotransferase |
| contig_38859_2 | 0.1756 | 1 | 1 | 0.3566 | 1 | 1 | 0.1572 | hypersensitive-induced response protein 1-like |
| contig_28340_4 | 0.1759 | 1 | 1 | 0.3567 | 1 | 1 | 0.1572 | Ras-like protein Rab-11B |
| contig_17153_5 | 0.1775 | 1 | 1 | 0.3593 | 1 | 1 | 0.1584 | predicted protein |
| contig_28369_6 | 0.1776 | 1 | 1 | 0.3593 | 1 | 1 | 0.1584 | predicted protein |
| contig_33399_2 | 0.1784 | 1 | 1 | 0.3604 | 1 | 1 | 0.1589 | predicted protein |
| contig_16833_4 | 0.1811 | 1 | 1 | 0.3652 | 1 | 1 | 0.161 | 60S ribosomal protein |
| contig_25351_2 | 0.1813 | 1 | 1 | 0.3652 | 1 | 1 | 0.161 | predicted protein |
| contig_23548_5 | 0.1817 | 1 | 1 | 0.3656 | 1 | 1 | 0.1612 | ATP synthase subunit beta mitochondrial |

| | | | | | | | | |
|---|---|---|---|---|---|---|---|---|
| contig_28331_5 | 0.1821 | 1 | 1 | 0.3656 | 1 | 1 | 0.1612 | transketolase |
| contig_28730_5 | 0.1822 | 1 | 1 | 0.3656 | 1 | 1 | 0.1612 | predicted protein |
| contig_22476_2 | 0.1832 | 1 | 1 | 0.3672 | 1 | 1 | 0.1619 | predicted protein |
| contig_9753_3 | 0.184 | 1 | 1 | 0.3683 | 1 | 1 | 0.1624 | predicted protein |
| contig_25927_6 | 0.1853 | 1 | 1 | 0.3703 | 1 | 1 | 0.1633 | light harvesting protein |
| contig_22608_4 | 0.1865 | 1 | 1 | 0.3722 | 1 | 1 | 0.1641 | predicted protein |
| contig_7847_6 | 0.1876 | 1 | 1 | 0.3739 | 1 | 1 | 0.1648 | predicted protein |
| contig_26596_5 | 0.1878 | 1 | 1 | 0.3739 | 1 | 1 | 0.1648 | leucyl aminopeptidase |
| contig_30824_4 | 0.188 | 1 | 1 | 0.3739 | 1 | 1 | 0.1648 | predicted protein |
| contig_27423_4 | 0.1887 | 1 | 1 | 0.3746 | 1 | 1 | 0.1652 | predicted protein |
| contig_27236_4 | 0.1892 | 1 | 1 | 0.3751 | 1 | 1 | 0.1654 | translation initiation factor aif-2 |
| contig_25653_6 | 0.1904 | 1 | 1 | 0.3765 | 1 | 1 | 0.166 | 40S ribosomal protein S8 |
| contig_29806_5 | 0.1905 | 1 | 1 | 0.3765 | 1 | 1 | 0.166 | predicted protein |
| contig_23061_6 | 0.1907 | 1 | 1 | 0.3765 | 1 | 1 | 0.166 | predicted protein |
| contig_19038_3 | 0.1908 | 1 | 1 | 0.3765 | 1 | 1 | 0.166 | peptidylprolyl isomerase |
| contig_36399_5 | 0.192 | 1 | 1 | 0.3783 | 1 | 1 | 0.1668 | triosephosphate isomerase/glyceraldehyde-3-phosphate dehydrogenase precursor |
| contig_11460_2 | 0.1933 | 1 | 1 | 0.3803 | 1 | 1 | 0.1677 | predicted protein |
| contig_36050_1 | 0.194 | 1 | 1 | 0.3812 | 1 | 1 | 0.1681 | Light-harvesting complex I LH38 protein |
| contig_25363_4 | 0.1942 | 1 | 1 | 0.3812 | 1 | 1 | 0.1681 | ABC-transporter like protein |
| contig_24234_3 | 0.1946 | 1 | 1 | 0.3815 | 1 | 1 | 0.1682 | predicted protein |
| contig_32749_4 | 0.196 | 1 | 1 | 0.3838 | 1 | 1 | 0.1692 | leu phe-trna protein transferase |
| contig_14080_4 | 0.1969 | 1 | 1 | 0.3841 | 1 | 1 | 0.1694 | dihydroorotase |
| contig_16700_1 | 0.1972 | 1 | 1 | 0.3841 | 1 | 1 | 0.1694 | serine decarboxylase |
| contig_28923_1 | 0.1972 | 1 | 1 | 0.3841 | 1 | 1 | 0.1694 | predicted protein |
| contig_10358_5 | 0.1972 | 1 | 1 | 0.3841 | 1 | 1 | 0.1694 | 60S ribosomal protein L12 |
| contig_40835_6 | 0.1985 | 1 | 1 | 0.386 | 1 | 1 | 0.1702 | 60S ribosomal protein L35a-4 |

| | | | | | | | | |
|---|---|---|---|---|---|---|---|---|
| contig_5526_6 | 0.1997 | 1 | 1 | 0.38 74 | 1 | 1 | 0.17 08 | pol protein |
| contig_37712_ 6 | 0.1997 | 1 | 1 | 0.38 74 | 1 | 1 | 0.17 08 | predicted protein |
| contig_7122_5 | 0.2003 | 1 | 1 | 0.38 76 | 1 | 1 | 0.17 09 | ---NA--- |
| contig_26646_ 2 | 0.2004 | 1 | 1 | 0.38 76 | 1 | 1 | 0.17 09 | short-chain dehydrogenase/reductase acting with NAD or NADP as acceptor |
| contig_30838_ 5 | 0.2008 | 1 | 1 | 0.38 76 | 1 | 1 | 0.17 09 | histone H4 |
| contig_8226_3 | 0.2008 | 1 | 1 | 0.38 76 | 1 | 1 | 0.17 09 | NAD(P) transhydrogenase mitochondrial |
| contig_28628_ 1 | 0.2021 | 1 | 1 | 0.38 94 | 1 | 1 | 0.17 17 | phosphoribosylaminoimidazolecarbo xamide formyltransferase/IMP cyclohydrolase |
| contig_24590_ 4 | 0.2022 | 1 | 1 | 0.38 94 | 1 | 1 | 0.17 17 | acyl-CoA dehydrogenase NM domain-like protein |
| contig_24961_ 6 | 0.2034 | 1 | 1 | 0.39 05 | 1 | 1 | 0.17 22 | mitochondrial carrier |
| contig_17882_ 6 | 0.2036 | 1 | 1 | 0.39 05 | 1 | 1 | 0.17 22 | light harvesting protein |
| contig_26792_ 2 | 0.2038 | 1 | 1 | 0.39 05 | 1 | 1 | 0.17 22 | predicted protein |
| contig_10732_ 2 | 0.2039 | 1 | 1 | 0.39 05 | 1 | 1 | 0.17 22 | transaldolase |
| contig_34472_ 2 | 0.2042 | 1 | 1 | 0.39 07 | 1 | 1 | 0.17 23 | histone h4 |
| contig_28215_ 4 | 0.2046 | 1 | 1 | 0.39 1 | 1 | 1 | 0.17 24 | light harvesting complex protein |
| contig_30830_ 5 | 0.2049 | 1 | 1 | 0.39 1 | 1 | 1 | 0.17 24 | PGR5-like protein 1A chloroplastic |
| contig_18025_ 5 | 0.2052 | 1 | 1 | 0.39 11 | 1 | 1 | 0.17 24 | histone H2B |
| contig_24888_ 1 | 0.2054 | 1 | 1 | 0.39 11 | 1 | 1 | 0.17 24 | chromodomain-helicase-DNA-binding protein 7 |
| contig_2430_4 | 0.2061 | 1 | 1 | 0.39 15 | 1 | 1 | 0.17 26 | ubiquinol-cytochrome c r |
| contig_6576_3 | 0.2064 | 1 | 1 | 0.39 15 | 1 | 1 | 0.17 26 | protein COFACTOR ASSEMBLY OF COMPLEX C SUBUNIT B CCB1 chloroplastic |
| contig_25854_ 4 | 0.2064 | 1 | 1 | 0.39 15 | 1 | 1 | 0.17 26 | L30e-like protein |
| contig_9651_3 | 0.2069 | 1 | 1 | 0.39 18 | 1 | 1 | 0.17 28 | ribosomal protein RPL17 |
| contig_33797_ 1 | 0.208 | 1 | 1 | 0.39 36 | 1 | 1 | 0.17 35 | glycine-tRNA ligase |
| contig_12107_ 3 | 0.2085 | 1 | 1 | 0.39 4 | 1 | 1 | 0.17 37 | argininosuccinate synthase |
| contig_7984_6 | 0.2096 | 1 | 1 | 0.39 56 | 1 | 1 | 0.17 44 | ribosomal protein L1 (chloroplast) |
| contig_28172_ 4 | 0.2099 | 1 | 1 | 0.39 56 | 1 | 1 | 0.17 44 | 60S ribosomal protein L5 |
| contig_14395_ 5 | 0.2111 | 1 | 1 | 0.39 74 | 1 | 1 | 0.17 52 | manganese superoxide dismutase |

| | | | | | | | | |
|---|---|---|---|---|---|---|---|---|
| contig_40158_1 | 0.2114 | 1 | 1 | 0.3975 | 1 | 1 | 0.1752 | extrinsic protein in photosystem II |
| contig_27301_5 | 0.2142 | 1 | 1 | 0.4013 | 1 | 1 | 0.177 | enolase |
| contig_28255_4 | 0.2143 | 1 | 1 | 0.4013 | 1 | 1 | 0.177 | long chain acyl-coa synthetase |
| contig_28304_6 | 0.2147 | 1 | 1 | 0.4013 | 1 | 1 | 0.177 | predicted protein |
| contig_16230_2 | 0.2148 | 1 | 1 | 0.4013 | 1 | 1 | 0.177 | phosphoglycerate mutase |
| contig_5001_1 | 0.2148 | 1 | 1 | 0.4013 | 1 | 1 | 0.177 | membrane guanylyl cyclase |
| contig_37939_4 | 0.2185 | 1 | 1 | 0.4078 | 1 | 1 | 0.1798 | predicted protein |
| contig_13656_4 | 0.2189 | 1 | 1 | 0.408 | 1 | 1 | 0.1799 | predicted protein |
| contig_19309_1 | 0.2212 | 1 | 1 | 0.4117 | 1 | 1 | 0.1815 | 60S ribosomal protein L9-like |
| contig_25921_4 | 0.2214 | 1 | 1 | 0.4117 | 1 | 1 | 0.1815 | predicted protein |
| contig_14621_1 | 0.223 | 1 | 1 | 0.4142 | 1 | 1 | 0.1826 | predicted protein |
| contig_26806_4 | 0.2238 | 1 | 1 | 0.4142 | 1 | 1 | 0.1826 | rab11 protein |
| contig_5881_4 | 0.2239 | 1 | 1 | 0.4142 | 1 | 1 | 0.1826 | elongation factor Tu (chloroplast) |
| contig_7536_5 | 0.2241 | 1 | 1 | 0.4142 | 1 | 1 | 0.1826 | predicted protein |
| contig_9049_5 | 0.2248 | 1 | 1 | 0.4142 | 1 | 1 | 0.1826 | enolase |
| contig_27862_5 | 0.2253 | 1 | 1 | 0.4142 | 1 | 1 | 0.1826 | predicted protein |
| contig_33347_6 | 0.2256 | 1 | 1 | 0.4142 | 1 | 1 | 0.1826 | V-type ATPase A subunit |
| contig_33528_6 | 0.2257 | 1 | 1 | 0.4142 | 1 | 1 | 0.1826 | SPX domain-containing membrane protein At4g22990-like |
| contig_19181_5 | 0.2258 | 1 | 1 | 0.4142 | 1 | 1 | 0.1826 | ribosomal protein S4 |
| contig_16347_5 | 0.226 | 1 | 1 | 0.4142 | 1 | 1 | 0.1826 | predicted protein |
| contig_17744_4 | 0.2261 | 1 | 1 | 0.4142 | 1 | 1 | 0.1826 | integral membrane protein |
| contig_3138_3 | 0.2262 | 1 | 1 | 0.4142 | 1 | 1 | 0.1826 | predicted protein |
| contig_35523_6 | 0.2263 | 1 | 1 | 0.4142 | 1 | 1 | 0.1826 | 60S ribosomal protein L10a |
| contig_28140_1 | 0.2276 | 1 | 1 | 0.4161 | 1 | 1 | 0.1835 | citrate (Si)-synthase eukaryotic |
| contig_10170_1 | 0.229 | 1 | 1 | 0.4181 | 1 | 1 | 0.1843 | predicted protein |
| contig_30068_2 | 0.2294 | 1 | 1 | 0.4184 | 1 | 1 | 0.1845 | 40s ribosomal protein s4 |
| contig_9153_4 | 0.2302 | 1 | 1 | 0.4193 | 1 | 1 | 0.1849 | predicted protein |

| | | | | | | | | |
|---|---|---|---|---|---|---|---|---|
| contig_28796_6 | 0.2316 | 1 | 1 | 0.4214 | 1 | 1 | 0.1858 | predicted protein |
| contig_30727_4 | 0.232 | 1 | 1 | 0.4216 | 1 | 1 | 0.1859 | predicted protein |
| contig_30693_4 | 0.234 | 1 | 1 | 0.4246 | 1 | 1 | 0.1872 | ubiquitin-conjugating enzyme E2-16 kDa |
| contig_17529_2 | 0.2345 | 1 | 1 | 0.4246 | 1 | 1 | 0.1872 | ribose-phosphate pyrophosphokinase |
| contig_29429_4 | 0.2345 | 1 | 1 | 0.4246 | 1 | 1 | 0.1872 | predicted protein |
| contig_36134_4 | 0.238 | 1 | 1 | 0.4305 | 1 | 1 | 0.1898 | fucoxanthin chlorophyll a/c protein |
| contig_11002_5 | 0.2383 | 1 | 1 | 0.4305 | 1 | 1 | 0.1898 | peroxiredoxin |
| contig_27033_6 | 0.2388 | 1 | 1 | 0.4308 | 1 | 1 | 0.1899 | glucokinase |
| contig_27523_3 | 0.2395 | 1 | 1 | 0.4316 | 1 | 1 | 0.1903 | plastid protoporphyrinogen oxidase |
| contig_40734_1 | 0.2401 | 1 | 1 | 0.4321 | 1 | 1 | 0.1905 | predicted protein |
| contig_18863_3 | 0.2412 | 1 | 1 | 0.4336 | 1 | 1 | 0.1912 | nucleoside diphosphate kinase |
| contig_9532_4 | 0.2423 | 1 | 1 | 0.435 | 1 | 1 | 0.1918 | NADH dehydrogenase [ubiquinone] iron-sulfur protein 8-A mitochondrial |
| contig_29419_6 | 0.2435 | 1 | 1 | 0.4363 | 1 | 1 | 0.1924 | leucyl-tRNA synthetase |
| contig_9858_4 | 0.2435 | 1 | 1 | 0.4363 | 1 | 1 | 0.1924 | argininosuccinate synthase |
| contig_24815_3 | 0.244 | 1 | 1 | 0.4367 | 1 | 1 | 0.1925 | MFS transporter |
| contig_16732_6 | 0.2443 | 1 | 1 | 0.4367 | 1 | 1 | 0.1925 | membrane spanning protein of the nonaspanin family |
| contig_15395_3 | 0.2456 | 1 | 1 | 0.4384 | 1 | 1 | 0.1933 | predicted protein |
| contig_28439_5 | 0.2478 | 1 | 1 | 0.4418 | 1 | 1 | 0.1948 | 3-isopropylmalate dehydrogenase |
| contig_6528_1 | 0.2488 | 1 | 1 | 0.4427 | 1 | 1 | 0.1952 | 40S ribosomal protein S19-3 |
| contig_5908_5 | 0.2489 | 1 | 1 | 0.4427 | 1 | 1 | 0.1952 | predicted protein |
| contig_21850_6 | 0.2493 | 1 | 1 | 0.443 | 1 | 1 | 0.1953 | 40S ribosomal protein S14 |
| contig_14784_5 | 0.2497 | 1 | 1 | 0.4431 | 1 | 1 | 0.1954 | atp-dependent RNA helicase |
| contig_33304_4 | 0.2501 | 1 | 1 | 0.4434 | 1 | 1 | 0.1955 | predicted protein |
| contig_25211_4 | 0.2519 | 1 | 1 | 0.4459 | 1 | 1 | 0.1966 | silicon transporter |
| contig_29073_1 | 0.2524 | 1 | 1 | 0.4462 | 1 | 1 | 0.1967 | 26S proteasome non-ATPase regulatory subunit 6 |
| contig_5563_5 | 0.2528 | 1 | 1 | 0.4462 | 1 | 1 | 0.1967 | predicted protein |
| contig_27634_6 | 0.2529 | 1 | 1 | 0.4462 | 1 | 1 | 0.1967 | eukaryotic translation initiation factor 3 subunit E |

| | | | | | | | | |
|---|---|---|---|---|---|---|---|---|
| contig_22730_1 | 0.2564 | 1 | 1 | 0.4514 | 1 | 1 | 0.199 | predicted protein |
| contig_10829_2 | 0.2566 | 1 | 1 | 0.4514 | 1 | 1 | 0.199 | plastid fructose 1 6-bisphosphatase isoform 1 |
| contig_40949_4 | 0.2567 | 1 | 1 | 0.4514 | 1 | 1 | 0.199 | U3 small nucleolar RNA-associated protein 18 homolog |
| contig_21631_6 | 0.2574 | 1 | 1 | 0.4522 | 1 | 1 | 0.1994 | predicted protein |
| contig_39895_1 | 0.2581 | 1 | 1 | 0.4527 | 1 | 1 | 0.1996 | protein fucoxanthin chlorophyll a/c protein |
| contig_14140_6 | 0.2588 | 1 | 1 | 0.4531 | 1 | 1 | 0.1998 | predicted protein |
| contig_24939_3 | 0.2589 | 1 | 1 | 0.4531 | 1 | 1 | 0.1998 | beta-ketoacyl-[acyl-carrier-protein] synthase II |
| contig_3085_5 | 0.2598 | 1 | 1 | 0.4542 | 1 | 1 | 0.2002 | ferredoxin (chloroplast) |
| contig_22773_6 | 0.2609 | 1 | 1 | 0.4551 | 1 | 1 | 0.2006 | phosphoglycerate kinase |
| contig_5318_4 | 0.261 | 1 | 1 | 0.4551 | 1 | 1 | 0.2006 | 40S ribosomal protein S15a |
| contig_30749_6 | 0.2612 | 1 | 1 | 0.4551 | 1 | 1 | 0.2006 | fucoxanthin-chlorophyll a/c light-harvesting protein |
| contig_24765_5 | 0.2622 | 1 | 1 | 0.4563 | 1 | 1 | 0.2012 | ABC transporter ATP-binding protein/permease |
| contig_16670_4 | 0.2626 | 1 | 1 | 0.4565 | 1 | 1 | 0.2013 | porphobilinogen deaminase |
| contig_32515_5 | 0.2642 | 1 | 1 | 0.4588 | 1 | 1 | 0.2023 | phosphoenolpyruvate synthase |
| contig_10657_4 | 0.2647 | 1 | 1 | 0.4591 | 1 | 1 | 0.2024 | atp-dependent RNA helicase |
| contig_14974_2 | 0.2662 | 1 | 1 | 0.4612 | 1 | 1 | 0.2033 | dnaj subfamily b member 4 |
| contig_38368_6 | 0.2668 | 1 | 1 | 0.4617 | 1 | 1 | 0.2036 | glutathione S-transferase |
| contig_27477_6 | 0.2677 | 1 | 1 | 0.4623 | 1 | 1 | 0.2038 | gdp-mannose 4 6-dehydratase |
| contig_28787_6 | 0.2678 | 1 | 1 | 0.4623 | 1 | 1 | 0.2038 | tryptophan synthase |
| contig_23765_6 | 0.2684 | 1 | 1 | 0.4624 | 1 | 1 | 0.2039 | predicted protein |
| contig_21520_1 | 0.2687 | 1 | 1 | 0.4624 | 1 | 1 | 0.2039 | phosphoglycerate mutase 1 family |
| contig_29454_5 | 0.2688 | 1 | 1 | 0.4624 | 1 | 1 | 0.2039 | peptidylprolyl isomerase |
| contig_40765_3 | 0.2694 | 1 | 1 | 0.4631 | 1 | 1 | 0.2042 | triosephosphate isomerase/glyceraldehyde-3-phosphate dehydrogenase precursor |
| contig_34938_2 | 0.2703 | 1 | 1 | 0.4637 | 1 | 1 | 0.2045 | predicted protein |
| contig_22932_5 | 0.2704 | 1 | 1 | 0.4637 | 1 | 1 | 0.2045 | predicted protein |
| contig_37972_6 | 0.2713 | 1 | 1 | 0.4643 | 1 | 1 | 0.2047 | predicted protein |

| | | | | | | | | |
|---|---|---|---|---|---|---|---|---|
| contig_9136_5 | 0.2715 | 1 | 1 | 0.4643 | 1 | 1 | 0.2047 | predicted protein |
| contig_8371_4 | 0.2717 | 1 | 1 | 0.4643 | 1 | 1 | 0.2047 | Rab family other |
| contig_22684_3 | 0.2721 | 1 | 1 | 0.4643 | 1 | 1 | 0.2047 | predicted protein |
| contig_22256_3 | 0.2723 | 1 | 1 | 0.4643 | 1 | 1 | 0.2047 | 30S ribosomal protein S15 |
| contig_33573_6 | 0.274 | 1 | 1 | 0.4662 | 1 | 1 | 0.2056 | ---NA--- |
| contig_10966_3 | 0.274 | 1 | 1 | 0.4662 | 1 | 1 | 0.2056 | predicted protein |
| contig_22496_6 | 0.2752 | 1 | 1 | 0.4678 | 1 | 1 | 0.2062 | predicted protein |
| contig_10491_6 | 0.2757 | 1 | 1 | 0.4679 | 1 | 1 | 0.2063 | glutamine synthetase catalytic domain protein protein |
| contig_24908_5 | 0.2761 | 1 | 1 | 0.4681 | 1 | 1 | 0.2064 | transitional endoplasmic reticulum ATPase TER94 |
| contig_6572_6 | 0.2769 | 1 | 1 | 0.4681 | 1 | 1 | 0.2064 | catalase |
| contig_38458_3 | 0.2772 | 1 | 1 | 0.4681 | 1 | 1 | 0.2064 | Proteasome subunit alpha/beta |
| contig_39062_4 | 0.2774 | 1 | 1 | 0.4681 | 1 | 1 | 0.2064 | 2-isopropylmalate synthase |
| contig_24617_4 | 0.2775 | 1 | 1 | 0.4681 | 1 | 1 | 0.2064 | predicted protein |
| contig_23768_5 | 0.278 | 1 | 1 | 0.4681 | 1 | 1 | 0.2064 | Flavocytochrome c |
| contig_6097_6 | 0.2782 | 1 | 1 | 0.4681 | 1 | 1 | 0.2064 | S-adenosylmethionine synthase 2 |
| contig_19474_5 | 0.2782 | 1 | 1 | 0.4681 | 1 | 1 | 0.2064 | aspartate aminotransferase |
| contig_11653_4 | 0.2788 | 1 | 1 | 0.4686 | 1 | 1 | 0.2066 | COP9 signalosome complex subunit 3 |
| contig_33090_6 | 0.2806 | 1 | 1 | 0.4711 | 1 | 1 | 0.2077 | predicted protein |
| contig_28323_4 | 0.281 | 1 | 1 | 0.4713 | 1 | 1 | 0.2078 | leucyl aminopeptidase |
| contig_20045_5 | 0.2818 | 1 | 1 | 0.4717 | 1 | 1 | 0.208 | AP-1 complex subunit sigma-1 |
| contig_41228_1 | 0.2819 | 1 | 1 | 0.4717 | 1 | 1 | 0.208 | light-harvesting protein |
| contig_28732_1 | 0.283 | 1 | 1 | 0.473 | 1 | 1 | 0.2085 | light harvesting protein |
| contig_23099_5 | 0.2836 | 1 | 1 | 0.4735 | 1 | 1 | 0.2088 | NAD(P)-dependent alcohol dehydrogenase |
| contig_24539_2 | 0.2845 | 1 | 1 | 0.4739 | 1 | 1 | 0.209 | phosphomannomutase/phosphoglucomutase |
| contig_20096_5 | 0.2845 | 1 | 1 | 0.4739 | 1 | 1 | 0.209 | predicted protein |
| contig_33470_2 | 0.2849 | 1 | 1 | 0.4741 | 1 | 1 | 0.209 | farnesyl pyrophosphate synthase |
| contig_27171_6 | 0.2854 | 1 | 1 | 0.4744 | 1 | 1 | 0.2092 | ribosome production factor 2 homolog |

| | | | | | | | | |
|---|---|---|---|---|---|---|---|---|
| contig_15022_4 | 0.2859 | 1 | 1 | 0.4747 | 1 | 1 | 0.2093 | ---NA--- |
| contig_28236_1 | 0.2872 | 1 | 1 | 0.4763 | 1 | 1 | 0.21 | tRNA splicing endonuclease putative |
| contig_28105_5 | 0.2888 | 1 | 1 | 0.4784 | 1 | 1 | 0.2109 | regulatory proteasome non-atpase subunit 3 |
| contig_24353_3 | 0.2898 | 1 | 1 | 0.4795 | 1 | 1 | 0.2114 | hemagglutinin/amebocyte aggregation factor |
| contig_8012_5 | 0.29 | 1 | 1 | 0.4795 | 1 | 1 | 0.2114 | predicted protein |
| contig_5881_5 | 0.292 | 1 | 1 | 0.4822 | 1 | 1 | 0.2126 | elongation factor Tu (chloroplast) |
| contig_33583_4 | 0.2926 | 1 | 1 | 0.4827 | 1 | 1 | 0.2128 | hypersensitive-induced response protein 1-like |
| contig_6442_4 | 0.2933 | 1 | 1 | 0.4834 | 1 | 1 | 0.2131 | putative ATP-dependent RNA helicase DHX30 isoform X3 |
| contig_3408_5 | 0.2937 | 1 | 1 | 0.4834 | 1 | 1 | 0.2132 | fucoxanthin-chlorophyll a-c binding protein |
| contig_18211_4 | 0.294 | 1 | 1 | 0.4834 | 1 | 1 | 0.2132 | light harvesting protein |
| contig_6511_6 | 0.2973 | 1 | 1 | 0.4873 | 1 | 1 | 0.2149 | glutamyl-tRNA synthetase |
| contig_26815_5 | 0.2973 | 1 | 1 | 0.4873 | 1 | 1 | 0.2149 | AGC/PKA protein kinase |
| contig_9710_5 | 0.2973 | 1 | 1 | 0.4873 | 1 | 1 | 0.2149 | predicted protein |
| contig_38999_3 | 0.2984 | 1 | 1 | 0.4885 | 1 | 1 | 0.2154 | predicted protein |
| contig_24410_6 | 0.2994 | 1 | 1 | 0.4892 | 1 | 1 | 0.2157 | sodium/potassium/calcium exchanger 2-like |
| contig_26661_4 | 0.2995 | 1 | 1 | 0.4892 | 1 | 1 | 0.2157 | photosystem II oxygen evolving complex protein PsbP |
| contig_38910_1 | 0.3018 | 1 | 1 | 0.4926 | 1 | 1 | 0.2172 | predicted protein |
| contig_33488_4 | 0.3022 | 1 | 1 | 0.4927 | 1 | 1 | 0.2172 | transcription elongation factor SPT6 |
| contig_41187_5 | 0.3038 | 1 | 1 | 0.4947 | 1 | 1 | 0.2181 | proteasome subunit beta type 4 |
| contig_30187_4 | 0.3055 | 1 | 1 | 0.497 | 1 | 1 | 0.2191 | adenylosuccinate lyase |
| contig_18485_3 | 0.3063 | 1 | 1 | 0.4978 | 1 | 1 | 0.2195 | Mitochondrial-processing peptidase subunit beta |
| contig_21644_6 | 0.3088 | 1 | 1 | 0.5012 | 1 | 1 | 0.221 | putative manganese superoxide dismutase protein |
| contig_10162_6 | 0.3094 | 1 | 1 | 0.5016 | 1 | 1 | 0.2211 | predicted protein |
| contig_5607_4 | 0.3096 | 1 | 1 | 0.5016 | 1 | 1 | 0.2211 | protease SohB |
| contig_20131_5 | 0.3124 | 1 | 1 | 0.5054 | 1 | 1 | 0.2228 | 60S ribosomal protein L21-A |
| contig_8365_3 | 0.3127 | 1 | 1 | 0.5054 | 1 | 1 | 0.2228 | pyruvate dehydrogenase |
| contig_34233_5 | 0.3143 | 1 | 1 | 0.5071 | 1 | 1 | 0.2236 | predicted protein |

| | | | | | | | | |
|---|---|---|---|---|---|---|---|---|
| contig_33419_4 | 0.3148 | 1 | 1 | 0.5071 | 1 | 1 | 0.2236 | light harvesting protein |
| contig_37770_2 | 0.3148 | 1 | 1 | 0.5071 | 1 | 1 | 0.2236 | photosystem I P700 apoprotein A1 (chloroplast) |
| contig_39855_1 | 0.315 | 1 | 1 | 0.5071 | 1 | 1 | 0.2236 | predicted protein |
| contig_29535_5 | 0.3176 | 1 | 1 | 0.5107 | 1 | 1 | 0.2252 | predicted protein |
| contig_35315_5 | 0.3184 | 1 | 1 | 0.5112 | 1 | 1 | 0.2254 | ---NA--- |
| contig_26198_6 | 0.3186 | 1 | 1 | 0.5112 | 1 | 1 | 0.2254 | ribosomal protein L20 |
| contig_22533_3 | 0.3202 | 1 | 1 | 0.5131 | 1 | 1 | 0.2262 | predicted protein |
| contig_15399_2 | 0.3205 | 1 | 1 | 0.5131 | 1 | 1 | 0.2262 | predicted protein |
| contig_38597_3 | 0.3224 | 1 | 1 | 0.5156 | 1 | 1 | 0.2273 | ---NA--- |
| contig_35287_3 | 0.3236 | 1 | 1 | 0.5164 | 1 | 1 | 0.2277 | photosystem II PsbU protein |
| contig_28392_1 | 0.324 | 1 | 1 | 0.5164 | 1 | 1 | 0.2277 | predicted protein |
| contig_8557_5 | 0.3242 | 1 | 1 | 0.5164 | 1 | 1 | 0.2277 | cytochrome b6 (chloroplast) |
| contig_39064_3 | 0.3243 | 1 | 1 | 0.5164 | 1 | 1 | 0.2277 | Glycerophosphoryl diester phosphodiesterase |
| contig_33997_4 | 0.3253 | 1 | 1 | 0.5175 | 1 | 1 | 0.2282 | predicted protein |
| contig_33090_4 | 0.3264 | 1 | 1 | 0.5187 | 1 | 1 | 0.2287 | predicted protein |
| contig_30516_6 | 0.3276 | 1 | 1 | 0.5201 | 1 | 1 | 0.2293 | aspartate--ammonia ligase |
| contig_30812_2 | 0.3291 | 1 | 1 | 0.5219 | 1 | 1 | 0.2301 | citrate synthase |
| contig_25953_4 | 0.3297 | 1 | 1 | 0.5224 | 1 | 1 | 0.2303 | predicted protein |
| contig_17279_6 | 0.3306 | 1 | 1 | 0.5229 | 1 | 1 | 0.2306 | vacuolar protein sorting-associated protein 26 |
| contig_40101_4 | 0.3307 | 1 | 1 | 0.5229 | 1 | 1 | 0.2306 | 40S ribosomal protein S13 |
| contig_22654_2 | 0.3333 | 1 | 1 | 0.5264 | 1 | 1 | 0.2321 | predicted protein |
| contig_33214_4 | 0.3342 | 1 | 1 | 0.5273 | 1 | 1 | 0.2325 | phosphoadenosine-phosphosulphate reductase |
| contig_20820_5 | 0.3351 | 1 | 1 | 0.5282 | 1 | 1 | 0.2329 | photosystem II protein D1 (chloroplast) |
| contig_9605_2 | 0.3378 | 1 | 1 | 0.5319 | 1 | 1 | 0.2345 | arginine biosynthesis ArgJ mitochondrial |
| contig_30747_4 | 0.3396 | 1 | 1 | 0.5341 | 1 | 1 | 0.2355 | bile acid:Na+ symporter BASS family |
| contig_29765_5 | 0.3401 | 1 | 1 | 0.5341 | 1 | 1 | 0.2355 | predicted protein |
| contig_28144_3 | 0.3403 | 1 | 1 | 0.5341 | 1 | 1 | 0.2355 | ---NA--- |

| | | | | | | | | |
|---|---|---|---|---|---|---|---|---|
| contig_12419_1 | 0.3429 | 1 | 1 | 0.5377 | 1 | 1 | 0.2371 | T-complex protein 1 subunit delta |
| contig_32611_6 | 0.3436 | 1 | 1 | 0.5382 | 1 | 1 | 0.2373 | predicted protein |
| contig_5545_2 | 0.3461 | 1 | 1 | 0.5415 | 1 | 1 | 0.2388 | NarK/NasA family nitrate transporter |
| contig_12711_6 | 0.3503 | 1 | 1 | 0.5471 | 1 | 1 | 0.2412 | predicted protein |
| contig_24322_5 | 0.3503 | 1 | 1 | 0.5471 | 1 | 1 | 0.2412 | ARM repeat superfamily protein |
| contig_20200_5 | 0.3513 | 1 | 1 | 0.548 | 1 | 1 | 0.2416 | 60s ribosomal protein l18a |
| contig_6119_2 | 0.3533 | 1 | 1 | 0.55 | 1 | 1 | 0.2425 | putative tRNA (cytidine(32)/guanosine(34)-2'-O)-methyltransferase |
| contig_28740_4 | 0.3536 | 1 | 1 | 0.55 | 1 | 1 | 0.2425 | tubulin alpha chain |
| contig_24205_6 | 0.3537 | 1 | 1 | 0.55 | 1 | 1 | 0.2425 | protein fucoxanthin chlorophyll a/c protein |
| contig_27060_3 | 0.3563 | 1 | 1 | 0.5536 | 1 | 1 | 0.2441 | calcium-binding protein |
| contig_38660_3 | 0.3567 | 1 | 1 | 0.5536 | 1 | 1 | 0.2441 | elongation factor 2 |
| contig_27872_3 | 0.3578 | 1 | 1 | 0.5539 | 1 | 1 | 0.2442 | predicted protein |
| contig_34763_6 | 0.3579 | 1 | 1 | 0.5539 | 1 | 1 | 0.2442 | myosin light chain kinase |
| contig_29713_5 | 0.3582 | 1 | 1 | 0.5539 | 1 | 1 | 0.2442 | probable prefoldin subunit 2 |
| contig_21839_5 | 0.3583 | 1 | 1 | 0.5539 | 1 | 1 | 0.2442 | 1-deoxy-d-xylulose-5-phosphate reductoisomerase |
| contig_22708_3 | 0.3592 | 1 | 1 | 0.5546 | 1 | 1 | 0.2445 | Laminin G domain protein |
| contig_12588_5 | 0.3616 | 1 | 1 | 0.5577 | 1 | 1 | 0.2459 | predicted protein |
| contig_7371_4 | 0.3663 | 1 | 1 | 0.5644 | 1 | 1 | 0.2488 | Lipocalin family protein |
| contig_26510_5 | 0.367 | 1 | 1 | 0.565 | 1 | 1 | 0.2491 | LL-diaminopimelate aminotransferase |
| contig_25573_4 | 0.3686 | 1 | 1 | 0.5666 | 1 | 1 | 0.2498 | predicted protein |
| contig_24202_5 | 0.369 | 1 | 1 | 0.5666 | 1 | 1 | 0.2498 | 2-isopropylmalate synthase |
| contig_10198_5 | 0.3693 | 1 | 1 | 0.5666 | 1 | 1 | 0.2498 | predicted protein |
| contig_22487_6 | 0.3699 | 1 | 1 | 0.5666 | 1 | 1 | 0.2498 | predicted protein |
| contig_6884_5 | 0.3699 | 1 | 1 | 0.5666 | 1 | 1 | 0.2498 | putative ATP-dependent RNA helicase DHX30 isoform X3 |
| contig_32599_2 | 0.3709 | 1 | 1 | 0.5675 | 1 | 1 | 0.2502 | ATP-dependent DNA helicase PIF1-like |
| contig_20846_4 | 0.3744 | 1 | 1 | 0.5724 | 1 | 1 | 0.2523 | fructose-bisphosphate aldolase |

| | | | | | | | | |
|---|---|---|---|---|---|---|---|---|
| contig_27013_6 | 0.3748 | 1 | 1 | 0.5724 | 1 | 1 | 0.2524 | sulfite reductase ferredoxin dependent |
| contig_5548_1 | 0.3763 | 1 | 1 | 0.574 | 1 | 1 | 0.2531 | geranylgeranyl pyrophosphate synthase |
| contig_18936_6 | 0.3769 | 1 | 1 | 0.5741 | 1 | 1 | 0.2531 | 40S ribosomal protein S3a |
| contig_27420_4 | 0.3772 | 1 | 1 | 0.5741 | 1 | 1 | 0.2531 | polyhydroxybutyrate depolymerase |
| contig_28948_1 | 0.3775 | 1 | 1 | 0.5741 | 1 | 1 | 0.2531 | geranylgeranyl reductase |
| contig_15868_5 | 0.3801 | 1 | 1 | 0.5776 | 1 | 1 | 0.2547 | fucoxanthin-chlorophyll a-c binding protein |
| contig_6077_5 | 0.3809 | 1 | 1 | 0.5782 | 1 | 1 | 0.2549 | 60s ribosomal protein l9 |
| contig_22578_5 | 0.3818 | 1 | 1 | 0.5789 | 1 | 1 | 0.2552 | Urea-proton symporter DUR3 |
| contig_19765_6 | 0.3838 | 1 | 1 | 0.5809 | 1 | 1 | 0.2561 | predicted protein |
| contig_25123_5 | 0.3839 | 1 | 1 | 0.5809 | 1 | 1 | 0.2561 | ubiquitin-activating enzyme E1 protein 3 |
| contig_20909_4 | 0.3843 | 1 | 1 | 0.5809 | 1 | 1 | 0.2561 | 40S ribosomal protein |
| contig_15445_5 | 0.3849 | 1 | 1 | 0.5809 | 1 | 1 | 0.2561 | zeaxanthin epoxidase |
| contig_18476_4 | 0.3851 | 1 | 1 | 0.5809 | 1 | 1 | 0.2561 | transmembrane 9 superfamily member 4 |
| contig_18884_5 | 0.3854 | 1 | 1 | 0.5809 | 1 | 1 | 0.2561 | predicted protein |
| contig_5829_1 | 0.3865 | 1 | 1 | 0.5818 | 1 | 1 | 0.2565 | gametolysin peptidase M11 |
| contig_25876_4 | 0.3867 | 1 | 1 | 0.5818 | 1 | 1 | 0.2565 | predicted protein |
| contig_24278_4 | 0.3906 | 1 | 1 | 0.5863 | 1 | 1 | 0.2585 | predicted protein |
| contig_16087_6 | 0.3908 | 1 | 1 | 0.5863 | 1 | 1 | 0.2585 | predicted protein |
| contig_29033_5 | 0.3909 | 1 | 1 | 0.5863 | 1 | 1 | 0.2585 | EF-hand domain-containing protein |
| contig_38608_1 | 0.3916 | 1 | 1 | 0.5868 | 1 | 1 | 0.2587 | cytochrome 450 |
| contig_17212_4 | 0.3923 | 1 | 1 | 0.587 | 1 | 1 | 0.2588 | uroporphyrinogen decarboxylase |
| contig_36054_1 | 0.3925 | 1 | 1 | 0.587 | 1 | 1 | 0.2588 | protein fucoxanthin chlorophyll a/c protein |
| contig_33296_6 | 0.394 | 1 | 1 | 0.5886 | 1 | 1 | 0.2595 | light-harvesting protein |
| contig_30481_5 | 0.3959 | 1 | 1 | 0.5909 | 1 | 1 | 0.2605 | ornithine cyclodeaminase |
| contig_25696_5 | 0.3965 | 1 | 1 | 0.5912 | 1 | 1 | 0.2607 | predicted protein |
| contig_8378_3 | 0.3981 | 1 | 1 | 0.5931 | 1 | 1 | 0.2615 | phosphoglyceromutase |
| contig_5607_2 | 0.3986 | 1 | 1 | 0.5932 | 1 | 1 | 0.2615 | protease SohB |

| | | | | | | | | |
|---|---|---|---|---|---|---|---|---|
| contig_28208_6 | 0.3991 | 1 | 1 | 0.5934 | 1 | 1 | 0.2616 | SPFH/Band 7/PHB domain protein |
| contig_26152_4 | 0.4002 | 1 | 1 | 0.5944 | 1 | 1 | 0.2621 | inositol 5-phosphatase |
| contig_5770_5 | 0.4014 | 1 | 1 | 0.5957 | 1 | 1 | 0.2626 | ketol-acid reductoisomerase |
| contig_30561_2 | 0.4024 | 1 | 1 | 0.5966 | 1 | 1 | 0.263 | cytochrome b6-f complex iron-sulfur subunit |
| contig_28745_4 | 0.4032 | 1 | 1 | 0.5972 | 1 | 1 | 0.2633 | glutamyl-trna reductase |
| contig_3037_1 | 0.4038 | 1 | 1 | 0.5975 | 1 | 1 | 0.2634 | ribosomal protein S4 (chloroplast) |
| contig_17741_5 | 0.4058 | 1 | 1 | 0.5998 | 1 | 1 | 0.2644 | endonuclease 4 |
| contig_35157_2 | 0.4082 | 1 | 1 | 0.6029 | 1 | 1 | 0.2658 | predicted protein |
| contig_29915_2 | 0.4097 | 1 | 1 | 0.6045 | 1 | 1 | 0.2665 | nucleolar protein nop56 |
| contig_17663_5 | 0.4154 | 1 | 1 | 0.6119 | 1 | 1 | 0.2698 | pyruvate dehydrogenase (acetyl-transferring) E1 component alpha subunit |
| contig_28913_5 | 0.4161 | 1 | 1 | 0.6119 | 1 | 1 | 0.2698 | predicted protein |
| contig_19536_5 | 0.4162 | 1 | 1 | 0.6119 | 1 | 1 | 0.2698 | methionine S-adenosyl transferase |
| contig_30397_1 | 0.4168 | 1 | 1 | 0.6119 | 1 | 1 | 0.2698 | FAD-dependent monooxygenase |
| contig_35046_3 | 0.4172 | 1 | 1 | 0.6119 | 1 | 1 | 0.2698 | CK1/CK1/CK1-D protein kinase |
| contig_27330_6 | 0.4174 | 1 | 1 | 0.6119 | 1 | 1 | 0.2698 | predicted protein |
| contig_20315_6 | 0.418 | 1 | 1 | 0.6119 | 1 | 1 | 0.2698 | NADH dehydrogenase subunit 5 (mitochondrion) |
| contig_16552_4 | 0.4181 | 1 | 1 | 0.6119 | 1 | 1 | 0.2698 | predicted protein |
| contig_30315_5 | 0.4184 | 1 | 1 | 0.6119 | 1 | 1 | 0.2698 | thioredoxin H4-1-like |
| contig_5668_1 | 0.4192 | 1 | 1 | 0.6125 | 1 | 1 | 0.27 | 40s ribosomal protein s20 |
| contig_6218_5 | 0.4205 | 1 | 1 | 0.6137 | 1 | 1 | 0.2706 | ferredoxin-nitrite reductase |
| contig_27603_4 | 0.4223 | 1 | 1 | 0.6158 | 1 | 1 | 0.2715 | predicted protein |
| contig_25252_3 | 0.4266 | 1 | 1 | 0.6203 | 1 | 1 | 0.2735 | Annexin protein |
| contig_11033_5 | 0.4266 | 1 | 1 | 0.6203 | 1 | 1 | 0.2735 | predicted protein |
| contig_23117_2 | 0.4268 | 1 | 1 | 0.6203 | 1 | 1 | 0.2735 | peroxisomal (S)-2-hydroxy-acid oxidase GLO5-like |
| contig_29793_4 | 0.427 | 1 | 1 | 0.6203 | 1 | 1 | 0.2735 | predicted protein |
| contig_27417_6 | 0.4276 | 1 | 1 | 0.6206 | 1 | 1 | 0.2736 | ferredoxin-nitrite reductase |

| | | | | | | | | |
|---|---|---|---|---|---|---|---|---|
| contig_15995_6 | 0.4286 | 1 | 1 | 0.6215 | 1 | 1 | 0.274 | cyclopropane-fatty-acyl-phospholipid synthase |
| contig_40422_5 | 0.4292 | 1 | 1 | 0.6218 | 1 | 1 | 0.2741 | light harvesting protein |
| contig_30573_3 | 0.4312 | 1 | 1 | 0.6235 | 1 | 1 | 0.2749 | predicted protein |
| contig_10416_5 | 0.4313 | 1 | 1 | 0.6235 | 1 | 1 | 0.2749 | ATP synthase F0 subunit 6 (mitochondrion) |
| contig_8422_4 | 0.4359 | 1 | 1 | 0.6297 | 1 | 1 | 0.2776 | predicted protein |
| contig_24804_5 | 0.4368 | 1 | 1 | 0.6303 | 1 | 1 | 0.2779 | ribosomal protein L13 |
| contig_31285_5 | 0.4372 | 1 | 1 | 0.6303 | 1 | 1 | 0.2779 | threonine ammonia-lyase biosynthetic |
| contig_9047_5 | 0.4398 | 1 | 1 | 0.6335 | 1 | 1 | 0.2793 | predicted protein |
| contig_28322_4 | 0.441 | 1 | 1 | 0.6346 | 1 | 1 | 0.2798 | farnesyl pyrophosphate synthase |
| contig_23416_1 | 0.4459 | 1 | 1 | 0.6411 | 1 | 1 | 0.2827 | predicted protein |
| contig_10991_6 | 0.4476 | 1 | 1 | 0.6428 | 1 | 1 | 0.2834 | ABC transporter |
| contig_33151_3 | 0.4497 | 1 | 1 | 0.6452 | 1 | 1 | 0.2845 | acetylglutamate kinase |
| contig_40197_2 | 0.4504 | 1 | 1 | 0.6452 | 1 | 1 | 0.2845 | protein fucoxanthin chlorophyll a/c protein |
| contig_13588_3 | 0.4512 | 1 | 1 | 0.6452 | 1 | 1 | 0.2845 | aarF domain-containing kinase |
| contig_38780_2 | 0.4513 | 1 | 1 | 0.6452 | 1 | 1 | 0.2845 | carbamoyl-phosphate synthase large subunit |
| contig_39705_5 | 0.4516 | 1 | 1 | 0.6452 | 1 | 1 | 0.2845 | inorganic pyrophosphatase |
| contig_25208_3 | 0.4518 | 1 | 1 | 0.6452 | 1 | 1 | 0.2845 | predicted protein |
| contig_27824_2 | 0.4527 | 1 | 1 | 0.6458 | 1 | 1 | 0.2847 | protein phosphatase 2C putative |
| contig_26362_4 | 0.453 | 1 | 1 | 0.6458 | 1 | 1 | 0.2847 | predicted protein |
| contig_25602_3 | 0.4534 | 1 | 1 | 0.6458 | 1 | 1 | 0.2847 | protein transport protein SEC61 subunit alpha |
| contig_9646_5 | 0.4551 | 1 | 1 | 0.6466 | 1 | 1 | 0.2851 | cell differentiation protein |
| contig_25806_5 | 0.4553 | 1 | 1 | 0.6466 | 1 | 1 | 0.2851 | 2-C-methyl-D-erythritol 2 4-cyclodiphosphate synthase |
| contig_8878_1 | 0.4553 | 1 | 1 | 0.6466 | 1 | 1 | 0.2851 | predicted protein |
| contig_30410_4 | 0.4566 | 1 | 1 | 0.6477 | 1 | 1 | 0.2856 | predicted protein |
| contig_38355_2 | 0.4569 | 1 | 1 | 0.6477 | 1 | 1 | 0.2856 | predicted protein |
| contig_19330_5 | 0.4584 | 1 | 1 | 0.6492 | 1 | 1 | 0.2862 | predicted protein |
| contig_40922_3 | 0.4608 | 1 | 1 | 0.652 | 1 | 1 | 0.2875 | V-type H-ATPase subunit C |

| | | | | | | | | |
|---|---|---|---|---|---|---|---|---|
| contig_7506_4 | 0.462 | 1 | 1 | 0.65 31 | 1 | 1 | 0.28 8 | apocytochrome b (mitochondrion) |
| contig_8116_6 | 0.4641 | 1 | 1 | 0.65 5 | 1 | 1 | 0.28 88 | 40S ribosomal protein S11 |
| contig_16863_ 4 | 0.4643 | 1 | 1 | 0.65 5 | 1 | 1 | 0.28 88 | D-3-phosphoglycerate dehydrogenase |
| contig_25119_ 2 | 0.4646 | 1 | 1 | 0.65 5 | 1 | 1 | 0.28 88 | 3-phosphoshikimate 1-carboxyvinyltransferase |
| contig_10410_ 5 | 0.4656 | 1 | 1 | 0.65 57 | 1 | 1 | 0.28 91 | LL-diaminopimelate aminotransferase |
| contig_14977_ 5 | 0.4667 | 1 | 1 | 0.65 62 | 1 | 1 | 0.28 93 | predicted protein |
| contig_23301_ 2 | 0.4668 | 1 | 1 | 0.65 62 | 1 | 1 | 0.28 93 | protein fucoxanthin chlorophyll a/c protein |
| contig_25833_ 1 | 0.4685 | 1 | 1 | 0.65 74 | 1 | 1 | 0.28 98 | predicted protein |
| contig_20147_ 4 | 0.4685 | 1 | 1 | 0.65 74 | 1 | 1 | 0.28 98 | fucoxanthin chlorophyl a/c protein |
| contig_9719_3 | 0.4714 | 1 | 1 | 0.66 09 | 1 | 1 | 0.29 14 | Co2+/Mg2+ efflux protein ApaG |
| contig_38903_ 2 | 0.4723 | 1 | 1 | 0.66 15 | 1 | 1 | 0.29 17 | ---NA--- |
| contig_26788_ 4 | 0.4739 | 1 | 1 | 0.66 32 | 1 | 1 | 0.29 24 | regulatory proteasome non-atpase subunit 2 |
| contig_33127_ 4 | 0.4745 | 1 | 1 | 0.66 32 | 1 | 1 | 0.29 24 | Not1-domain-containing protein |
| contig_9571_3 | 0.475 | 1 | 1 | 0.66 32 | 1 | 1 | 0.29 24 | ATP:ADP antiporter AAA family |
| contig_39819_ 3 | 0.4753 | 1 | 1 | 0.66 32 | 1 | 1 | 0.29 24 | predicted protein |
| contig_18307_ 6 | 0.4758 | 1 | 1 | 0.66 34 | 1 | 1 | 0.29 25 | predicted protein |
| contig_25106_ 6 | 0.4768 | 1 | 1 | 0.66 36 | 1 | 1 | 0.29 26 | predicted protein |
| contig_22050_ 5 | 0.4769 | 1 | 1 | 0.66 36 | 1 | 1 | 0.29 26 | 60S ribosomal protein L9 |
| contig_33542_ 2 | 0.4785 | 1 | 1 | 0.66 52 | 1 | 1 | 0.29 33 | nucleoside-diphosphate kinase |
| contig_24463_ 5 | 0.4793 | 1 | 1 | 0.66 58 | 1 | 1 | 0.29 35 | xanthine dehydrogenase molybdopterin binding subunit |
| contig_36079_ 3 | 0.4813 | 1 | 1 | 0.66 79 | 1 | 1 | 0.29 45 | photosystem II PsbU protein |
| contig_22941_ 4 | 0.4819 | 1 | 1 | 0.66 82 | 1 | 1 | 0.29 46 | mitochondrial carrier |
| contig_10800_ 3 | 0.483 | 1 | 1 | 0.66 86 | 1 | 1 | 0.29 48 | predicted protein |
| contig_36382_ 5 | 0.4831 | 1 | 1 | 0.66 86 | 1 | 1 | 0.29 48 | Ubiquitin-40S ribosomal protein S27a |
| contig_23660_ 5 | 0.4844 | 1 | 1 | 0.66 98 | 1 | 1 | 0.29 53 | predicted protein |
| contig_17128_ 2 | 0.4854 | 1 | 1 | 0.67 04 | 1 | 1 | 0.29 56 | predicted protein |

| | | | | | | | | |
|---|---|---|---|---|---|---|---|---|
| contig_29804_5 | 0.4863 | 1 | 1 | 0.6704 | 1 | 1 | 0.2956 | S-adenosylmethionine synthetase |
| contig_11577_6 | 0.4864 | 1 | 1 | 0.6704 | 1 | 1 | 0.2956 | Mitochondrial adenine nucleotide transporter ADNT1 |
| contig_24083_4 | 0.4866 | 1 | 1 | 0.6704 | 1 | 1 | 0.2956 | predicted protein |
| contig_14126_5 | 0.4914 | 1 | 1 | 0.6763 | 1 | 1 | 0.2982 | predicted protein |
| contig_13099_5 | 0.4919 | 1 | 1 | 0.6763 | 1 | 1 | 0.2982 | predicted protein |
| contig_15516_6 | 0.4922 | 1 | 1 | 0.6763 | 1 | 1 | 0.2982 | pyrimidine/purine nucleoside phosphorylase |
| contig_27823_2 | 0.4927 | 1 | 1 | 0.6763 | 1 | 1 | 0.2982 | predicted protein |
| contig_34697_6 | 0.4948 | 1 | 1 | 0.6786 | 1 | 1 | 0.2992 | asparagine synthase |
| contig_5503_5 | 0.4961 | 1 | 1 | 0.6793 | 1 | 1 | 0.2995 | predicted protein |
| contig_7732_1 | 0.4963 | 1 | 1 | 0.6793 | 1 | 1 | 0.2995 | predicted protein |
| contig_14552_5 | 0.4967 | 1 | 1 | 0.6793 | 1 | 1 | 0.2995 | phosphomannomutase |
| contig_23336_4 | 0.5003 | 1 | 1 | 0.6837 | 1 | 1 | 0.3014 | 1-hydroxy-2-methyl-2-(E)-butenyl-4-diphosphate synthase |
| contig_24978_5 | 0.5021 | 1 | 1 | 0.6856 | 1 | 1 | 0.3023 | predicted protein |
| contig_33374_5 | 0.5042 | 1 | 1 | 0.6877 | 1 | 1 | 0.3032 | predicted protein |
| contig_38336_2 | 0.5047 | 1 | 1 | 0.6879 | 1 | 1 | 0.3033 | class I SAM-dependent methyltransferase |
| contig_29992_5 | 0.507 | 1 | 1 | 0.6881 | 1 | 1 | 0.3034 | ---NA--- |
| contig_11078_3 | 0.5082 | 1 | 1 | 0.6881 | 1 | 1 | 0.3034 | peroxiredoxin |
| contig_16859_4 | 0.5082 | 1 | 1 | 0.6881 | 1 | 1 | 0.3034 | DNA-directed RNA polymerase II 135 kDa polypeptide |
| contig_25050_4 | 0.5084 | 1 | 1 | 0.6881 | 1 | 1 | 0.3034 | predicted protein |
| contig_23551_6 | 0.5085 | 1 | 1 | 0.6881 | 1 | 1 | 0.3034 | predicted protein |
| contig_40133_3 | 0.5088 | 1 | 1 | 0.6881 | 1 | 1 | 0.3034 | 30S ribosomal protein S15 |
| contig_28947_4 | 0.5089 | 1 | 1 | 0.6881 | 1 | 1 | 0.3034 | predicted protein |
| contig_27789_6 | 0.5093 | 1 | 1 | 0.6881 | 1 | 1 | 0.3034 | Mitochondrial-processing peptidase subunit alpha |
| contig_30420_3 | 0.5097 | 1 | 1 | 0.6881 | 1 | 1 | 0.3034 | predicted protein |
| contig_27061_1 | 0.5098 | 1 | 1 | 0.6881 | 1 | 1 | 0.3034 | acetyl-CoA carboxylase biotin carboxylase subunit |
| contig_28722_5 | 0.5103 | 1 | 1 | 0.6881 | 1 | 1 | 0.3034 | predicted protein |

| | | | | | | | | |
|---|---|---|---|---|---|---|---|---|
| contig_35847_5 | 0.5109 | 1 | 1 | 0.6881 | 1 | 1 | 0.3034 | serine/threonine-protein phosphatase PP1-beta catalytic subunit |
| contig_21557_3 | 0.5112 | 1 | 1 | 0.6881 | 1 | 1 | 0.3034 | 40S ribosomal protein S2 |
| contig_17375_3 | 0.5113 | 1 | 1 | 0.6881 | 1 | 1 | 0.3034 | cysteine synthase |
| contig_25885_5 | 0.5116 | 1 | 1 | 0.6881 | 1 | 1 | 0.3034 | acetyl-coenzyme a synthetase |
| contig_28238_5 | 0.5126 | 1 | 1 | 0.6887 | 1 | 1 | 0.3037 | ubiquinol-cytochrome c reductase iron-sulfur subunit |
| contig_38364_2 | 0.5134 | 1 | 1 | 0.6892 | 1 | 1 | 0.3039 | predicted protein |
| contig_25034_6 | 0.5138 | 1 | 1 | 0.6892 | 1 | 1 | 0.3039 | predicted protein |
| contig_41162_4 | 0.5158 | 1 | 1 | 0.6908 | 1 | 1 | 0.3046 | 60S ribosomal protein L9-like |
| contig_14595_4 | 0.516 | 1 | 1 | 0.6908 | 1 | 1 | 0.3046 | predicted protein |
| contig_25130_6 | 0.5172 | 1 | 1 | 0.6909 | 1 | 1 | 0.3046 | methionine-tRNA ligase |
| contig_22528_2 | 0.5173 | 1 | 1 | 0.6909 | 1 | 1 | 0.3046 | predicted protein |
| contig_6643_5 | 0.5174 | 1 | 1 | 0.6909 | 1 | 1 | 0.3046 | phosphoglycolate phosphatase |
| contig_31306_6 | 0.5217 | 1 | 1 | 0.696 | 1 | 1 | 0.3069 | predicted protein |
| contig_8346_5 | 0.5257 | 1 | 1 | 0.7008 | 1 | 1 | 0.309 | putative plastid division protein (chloroplast) |
| contig_34843_5 | 0.5268 | 1 | 1 | 0.7016 | 1 | 1 | 0.3093 | Major Facilitator Superfamily (MFS) |
| contig_17360_5 | 0.5292 | 1 | 1 | 0.7042 | 1 | 1 | 0.3105 | Annexin protein |
| contig_25458_4 | 0.5303 | 1 | 1 | 0.705 | 1 | 1 | 0.3108 | succinate dehydrogenase [ubiquinone] iron-sulfur subunit mitochondrial |
| contig_37899_2 | 0.5327 | 1 | 1 | 0.7063 | 1 | 1 | 0.3114 | ribosomal protein L2 (chloroplast) |
| contig_22602_4 | 0.5329 | 1 | 1 | 0.7063 | 1 | 1 | 0.3114 | putative all-trans-retinol 13 14-reductase |
| contig_18033_5 | 0.5331 | 1 | 1 | 0.7063 | 1 | 1 | 0.3114 | mitochondrial-processing peptidase subunit beta |
| contig_15379_5 | 0.5338 | 1 | 1 | 0.7063 | 1 | 1 | 0.3114 | nucleolar gtp-binding protein 1 |
| contig_6450_1 | 0.5339 | 1 | 1 | 0.7063 | 1 | 1 | 0.3114 | glutamine-fructose-6-phosphate transaminase |
| contig_38239_2 | 0.534 | 1 | 1 | 0.7063 | 1 | 1 | 0.3114 | predicted protein |
| contig_36104_5 | 0.5384 | 1 | 1 | 0.7115 | 1 | 1 | 0.3137 | 40S ribosomal protein S2 |
| contig_29950_5 | 0.5394 | 1 | 1 | 0.7116 | 1 | 1 | 0.3138 | Fe-S cluster assembly protein SufD |
| contig_28707_5 | 0.5399 | 1 | 1 | 0.7116 | 1 | 1 | 0.3138 | V-type H+-transporting ATPase subunit a |

| | | | | | | | | |
|---|---|---|---|---|---|---|---|---|
| contig_33495_6 | 0.5399 | 1 | 1 | 0.7116 | 1 | 1 | 0.3138 | triose or hexose phosphate/phosphate translocator |
| contig_40960_3 | 0.5416 | 1 | 1 | 0.713 | 1 | 1 | 0.3144 | predicted protein |
| contig_13819_5 | 0.5419 | 1 | 1 | 0.713 | 1 | 1 | 0.3144 | 7-dehydrocholesterol reductase |
| contig_11320_6 | 0.5424 | 1 | 1 | 0.713 | 1 | 1 | 0.3144 | predicted protein |
| contig_23071_3 | 0.5438 | 1 | 1 | 0.7143 | 1 | 1 | 0.3149 | D-3-phosphoglycerate dehydrogenase |
| contig_26135_5 | 0.5447 | 1 | 1 | 0.7148 | 1 | 1 | 0.3152 | Mitochondrial Carrier (MC) Family |
| contig_26293_4 | 0.5452 | 1 | 1 | 0.7148 | 1 | 1 | 0.3152 | predicted protein |
| contig_38299_1 | 0.5469 | 1 | 1 | 0.7163 | 1 | 1 | 0.3158 | ATP-binding cassette superfamily |
| contig_25423_6 | 0.5475 | 1 | 1 | 0.7163 | 1 | 1 | 0.3158 | 2 5-diketo-D-gluconic acid reductase A putative |
| contig_15835_5 | 0.5477 | 1 | 1 | 0.7163 | 1 | 1 | 0.3158 | ---NA--- |
| contig_5385_3 | 0.5501 | 1 | 1 | 0.7188 | 1 | 1 | 0.3169 | glutathione S-transferase Mu 3-like |
| contig_6339_4 | 0.5506 | 1 | 1 | 0.7188 | 1 | 1 | 0.3169 | predicted protein |
| contig_41396_3 | 0.5523 | 1 | 1 | 0.7205 | 1 | 1 | 0.3177 | predicted protein |
| contig_28624_5 | 0.5537 | 1 | 1 | 0.7217 | 1 | 1 | 0.3182 | eukaryotic peptide chain release factor subunit 1 |
| contig_29898_4 | 0.5557 | 1 | 1 | 0.7234 | 1 | 1 | 0.319 | mitochondrial 2-oxoglutarate/malate carrier protein |
| contig_17202_2 | 0.5563 | 1 | 1 | 0.7234 | 1 | 1 | 0.319 | predicted protein |
| contig_26693_5 | 0.5565 | 1 | 1 | 0.7234 | 1 | 1 | 0.319 | UMP-CMP kinase |
| contig_29933_6 | 0.5571 | 1 | 1 | 0.7236 | 1 | 1 | 0.319 | 30S ribosomal protein S1 |
| contig_39611_1 | 0.5582 | 1 | 1 | 0.7245 | 1 | 1 | 0.3194 | methionine-tRNA ligase |
| contig_37814_3 | 0.5588 | 1 | 1 | 0.7246 | 1 | 1 | 0.3195 | ribosomal protein S3 (chloroplast) |
| contig_37718_6 | 0.561 | 1 | 1 | 0.7269 | 1 | 1 | 0.3205 | predicted protein |
| contig_21568_5 | 0.5629 | 1 | 1 | 0.7286 | 1 | 1 | 0.3213 | predicted protein |
| contig_3376_2 | 0.5656 | 1 | 1 | 0.7314 | 1 | 1 | 0.3225 | elongator complex protein 1 |
| contig_21707_3 | 0.566 | 1 | 1 | 0.7314 | 1 | 1 | 0.3225 | 40S ribosomal protein S4 |
| contig_27843_1 | 0.5676 | 1 | 1 | 0.7328 | 1 | 1 | 0.3231 | Fe-S cluster assembly protein SufD |
| contig_25676_6 | 0.5681 | 1 | 1 | 0.7329 | 1 | 1 | 0.3231 | predicted protein |
| contig_26114_2 | 0.5696 | 1 | 1 | 0.7342 | 1 | 1 | 0.3237 | ATP phosphoribosyltransferase |

| | | | | | | | | |
|---|---|---|---|---|---|---|---|---|
| contig_8369_4 | 0.5728 | 1 | 1 | 0.7371 | 1 | 1 | 0.325 | Clp protease ATP binding subunit (chloroplast) |
| contig_29766_3 | 0.5728 | 1 | 1 | 0.7371 | 1 | 1 | 0.325 | GMP synthase (glutamine-hydrolyzing) domain-containing protein |
| contig_38581_3 | 0.5742 | 1 | 1 | 0.738 | 1 | 1 | 0.3254 | predicted protein |
| contig_12042_4 | 0.5745 | 1 | 1 | 0.738 | 1 | 1 | 0.3254 | coproporphyrinogen III oxidase |
| contig_39446_2 | 0.5755 | 1 | 1 | 0.7386 | 1 | 1 | 0.3257 | U6 snRNA-associated Sm-like protein LSm2 |
| contig_33115_6 | 0.5788 | 1 | 1 | 0.7418 | 1 | 1 | 0.3271 | predicted protein |
| contig_20020_4 | 0.5789 | 1 | 1 | 0.7418 | 1 | 1 | 0.3271 | predicted protein |
| contig_28316_5 | 0.5797 | 1 | 1 | 0.7421 | 1 | 1 | 0.3272 | hypothetical protein FRACYDRAFT_234640 |
| contig_9855_4 | 0.5806 | 1 | 1 | 0.7422 | 1 | 1 | 0.3272 | S-adenosyl-L-methionine-dependent methyltransferase |
| contig_13446_5 | 0.5807 | 1 | 1 | 0.7422 | 1 | 1 | 0.3272 | predicted protein |
| contig_37256_3 | 0.5832 | 1 | 1 | 0.7448 | 1 | 1 | 0.3284 | histone H3 |
| contig_10321_2 | 0.5851 | 1 | 1 | 0.7462 | 1 | 1 | 0.329 | beta-hydroxyacyl-acyl carrier protein dehydratase (FABZ) |
| contig_21965_5 | 0.5856 | 1 | 1 | 0.7462 | 1 | 1 | 0.329 | predicted protein |
| contig_25721_4 | 0.5858 | 1 | 1 | 0.7462 | 1 | 1 | 0.329 | putative manganese superoxide dismutase protein |
| contig_9226_3 | 0.5875 | 1 | 1 | 0.7478 | 1 | 1 | 0.3297 | 2-isopropylmalate synthase |
| contig_27172_4 | 0.5907 | 1 | 1 | 0.7512 | 1 | 1 | 0.3312 | peptidyl-prolyl cis-trans isomerase |
| contig_34549_1 | 0.5918 | 1 | 1 | 0.752 | 1 | 1 | 0.3316 | pirin family protein |
| contig_30338_1 | 0.5939 | 1 | 1 | 0.7534 | 1 | 1 | 0.3322 | 60S ribosomal protein L5 |
| contig_11343_2 | 0.5939 | 1 | 1 | 0.7534 | 1 | 1 | 0.3322 | probable aspartyl aminopeptidase |
| contig_20295_6 | 0.5949 | 1 | 1 | 0.754 | 1 | 1 | 0.3324 | cytochrome c oxidase subunit II (mitochondrion) |
| contig_19585_5 | 0.5957 | 1 | 1 | 0.7544 | 1 | 1 | 0.3326 | ---NA--- |
| contig_11299_6 | 0.5966 | 1 | 1 | 0.755 | 1 | 1 | 0.3329 | predicted protein |
| contig_41244_3 | 0.5981 | 1 | 1 | 0.7555 | 1 | 1 | 0.3331 | class II fructose-bisphosphate aldolase |
| contig_23045_4 | 0.5984 | 1 | 1 | 0.7555 | 1 | 1 | 0.3331 | amino acid/polyamine transporter |
| contig_25962_4 | 0.5985 | 1 | 1 | 0.7555 | 1 | 1 | 0.3331 | predicted protein |
| contig_14363_5 | 0.5998 | 1 | 1 | 0.7565 | 1 | 1 | 0.3335 | predicted protein |

| | | | | | | | | |
|---|---|---|---|---|---|---|---|---|
| contig_25076_6 | 0.6008 | 1 | 1 | 0.7571 | 1 | 1 | 0.3338 | phosphoglucomutase |
| contig_34264_6 | 0.6027 | 1 | 1 | 0.7571 | 1 | 1 | 0.3338 | probable small nuclear ribonucleoprotein E |
| contig_7315_2 | 0.6029 | 1 | 1 | 0.7571 | 1 | 1 | 0.3338 | predicted protein |
| contig_28483_6 | 0.6034 | 1 | 1 | 0.7571 | 1 | 1 | 0.3338 | predicted protein |
| contig_41157_2 | 0.6037 | 1 | 1 | 0.7571 | 1 | 1 | 0.3338 | predicted protein |
| contig_25682_4 | 0.6038 | 1 | 1 | 0.7571 | 1 | 1 | 0.3338 | Immunoglobulin G-binding protein A |
| contig_5961_5 | 0.6038 | 1 | 1 | 0.7571 | 1 | 1 | 0.3338 | cytochrome oxidase subunit III (mitochondrion) |
| contig_30128_6 | 0.6066 | 1 | 1 | 0.76 | 1 | 1 | 0.3351 | predicted protein |
| contig_3491_1 | 0.6075 | 1 | 1 | 0.7602 | 1 | 1 | 0.3352 | predicted protein |
| contig_39705_1 | 0.6078 | 1 | 1 | 0.7602 | 1 | 1 | 0.3352 | inorganic pyrophosphatase |
| contig_7288_5 | 0.61 | 1 | 1 | 0.761 | 1 | 1 | 0.3355 | protein fucoxanthin chlorophyll a/c protein |
| contig_9265_3 | 0.6114 | 1 | 1 | 0.761 | 1 | 1 | 0.3355 | mitochondrial carrier family |
| contig_25649_3 | 0.6119 | 1 | 1 | 0.761 | 1 | 1 | 0.3355 | tRNA pseudouridine(55) synthase |
| contig_9256_6 | 0.6119 | 1 | 1 | 0.761 | 1 | 1 | 0.3355 | predicted protein |
| contig_6155_6 | 0.612 | 1 | 1 | 0.761 | 1 | 1 | 0.3355 | light harvesting protein |
| contig_22469_6 | 0.612 | 1 | 1 | 0.761 | 1 | 1 | 0.3355 | predicted protein |
| contig_17918_5 | 0.6122 | 1 | 1 | 0.761 | 1 | 1 | 0.3355 | glutamate-1-semialdehyde 2 1-aminomutase |
| contig_28468_2 | 0.6124 | 1 | 1 | 0.761 | 1 | 1 | 0.3355 | ribosome recycling factor |
| contig_28061_5 | 0.6162 | 1 | 1 | 0.765 | 1 | 1 | 0.3373 | apospory-associated protein C |
| contig_10341_1 | 0.6171 | 1 | 1 | 0.7656 | 1 | 1 | 0.3375 | predicted protein |
| contig_6146_1 | 0.618 | 1 | 1 | 0.766 | 1 | 1 | 0.3377 | aliphatic amidase |
| contig_25515_1 | 0.6184 | 1 | 1 | 0.766 | 1 | 1 | 0.3377 | cysteine desulfurase |
| contig_14735_6 | 0.6197 | 1 | 1 | 0.7669 | 1 | 1 | 0.3381 | proteasome subunit beta type-5-like |
| contig_34263_4 | 0.6224 | 1 | 1 | 0.7696 | 1 | 1 | 0.3393 | nucleolar protein 56 |
| contig_40265_2 | 0.6284 | 1 | 1 | 0.7765 | 1 | 1 | 0.3423 | predicted protein |
| contig_22654_5 | 0.6295 | 1 | 1 | 0.7772 | 1 | 1 | 0.3427 | predicted protein |
| contig_32529_3 | 0.6317 | 1 | 1 | 0.7787 | 1 | 1 | 0.3433 | GDT1-like protein 2 chloroplastic |

| | | | | | | | | |
|---|---|---|---|---|---|---|---|---|
| contig_6242_5 | 0.6318 | 1 | 1 | 0.77 87 | 1 | 1 | 0.34 33 | predicted protein |
| contig_24715_ 2 | 0.6332 | 1 | 1 | 0.77 95 | 1 | 1 | 0.34 37 | predicted protein |
| contig_22967_ 5 | 0.6335 | 1 | 1 | 0.77 95 | 1 | 1 | 0.34 37 | predicted protein |
| contig_9827_1 | 0.6346 | 1 | 1 | 0.78 03 | 1 | 1 | 0.34 4 | predicted protein |
| contig_16774_ 4 | 0.6372 | 1 | 1 | 0.78 29 | 1 | 1 | 0.34 52 | probable microsomal very long chain fatty acid elongase |
| contig_9982_6 | 0.6382 | 1 | 1 | 0.78 34 | 1 | 1 | 0.34 54 | light harvesting protein |
| contig_10303_ 5 | 0.6407 | 1 | 1 | 0.78 59 | 1 | 1 | 0.34 65 | 40S ribosomal protein S3-3 |
| contig_33443_ 2 | 0.6428 | 1 | 1 | 0.78 78 | 1 | 1 | 0.34 73 | predicted protein |
| contig_33729_ 4 | 0.6475 | 1 | 1 | 0.79 17 | 1 | 1 | 0.34 9 | secreted protein |
| contig_17183_ 2 | 0.6479 | 1 | 1 | 0.79 17 | 1 | 1 | 0.34 9 | 40S ribosomal protein S17 |
| contig_37814_ 2 | 0.6484 | 1 | 1 | 0.79 17 | 1 | 1 | 0.34 9 | ribosomal protein S3 (chloroplast) |
| contig_20218_ 5 | 0.6485 | 1 | 1 | 0.79 17 | 1 | 1 | 0.34 9 | 40S ribosomal protein S26 |
| contig_24913_ 6 | 0.6485 | 1 | 1 | 0.79 17 | 1 | 1 | 0.34 9 | glutamate synthase subunit alpha |
| contig_39862_ 1 | 0.6499 | 1 | 1 | 0.79 27 | 1 | 1 | 0.34 95 | pyruvate kinase |
| contig_36080_ 5 | 0.6518 | 1 | 1 | 0.79 43 | 1 | 1 | 0.35 02 | predicted protein |
| contig_12505_ 5 | 0.6527 | 1 | 1 | 0.79 48 | 1 | 1 | 0.35 04 | threonine ammonia-lyase |
| contig_18343_ 1 | 0.6546 | 1 | 1 | 0.79 65 | 1 | 1 | 0.35 12 | glutamine synthetase |
| contig_17076_ 5 | 0.6553 | 1 | 1 | 0.79 68 | 1 | 1 | 0.35 13 | predicted protein |
| contig_39533_ 1 | 0.6559 | 1 | 1 | 0.79 69 | 1 | 1 | 0.35 13 | sodium/proton antiporter 1 |
| contig_34277_ 5 | 0.6566 | 1 | 1 | 0.79 69 | 1 | 1 | 0.35 14 | 2-isopropylmalate synthase |
| contig_37776_ 1 | 0.657 | 1 | 1 | 0.79 69 | 1 | 1 | 0.35 14 | DNA polymerase |
| contig_41409_ 3 | 0.6584 | 1 | 1 | 0.79 76 | 1 | 1 | 0.35 17 | ribosomal protein L13 |
| contig_21581_ 3 | 0.6586 | 1 | 1 | 0.79 76 | 1 | 1 | 0.35 17 | predicted protein |
| contig_40284_ 6 | 0.6604 | 1 | 1 | 0.79 91 | 1 | 1 | 0.35 23 | large subunit ribosomal protein L13e |
| contig_5934_3 | 0.6617 | 1 | 1 | 0.8 | 1 | 1 | 0.35 27 | bifunctional aconitate hydratase 2/2-methylisocitrate dehydratase |
| contig_9076_4 | 0.6622 | 1 | 1 | 0.80 01 | 1 | 1 | 0.35 27 | 60S ribosomal protein L24 putative |
| contig_28185_ 5 | 0.6629 | 1 | 1 | 0.80 01 | 1 | 1 | 0.35 27 | UMP-CMP kinase |

| | | | | | | | | |
|---|---|---|---|---|---|---|---|---|
| contig_5654_2 | 0.6634 | 1 | 1 | 0.80 01 | 1 | 1 | 0.35 27 | hypothetical protein THAOC_17223 partial |
| contig_15800_ 4 | 0.6638 | 1 | 1 | 0.80 01 | 1 | 1 | 0.35 27 | asparagine synthase |
| contig_28307_ 4 | 0.6656 | 1 | 1 | 0.80 15 | 1 | 1 | 0.35 34 | atp-dependent rna helicase uap56 |
| contig_22476_ 4 | 0.6662 | 1 | 1 | 0.80 16 | 1 | 1 | 0.35 34 | predicted protein |
| contig_15545_ 5 | 0.6683 | 1 | 1 | 0.80 35 | 1 | 1 | 0.35 43 | Small nuclear ribonucleoprotein G |
| contig_27804_ 5 | 0.6724 | 1 | 1 | 0.80 75 | 1 | 1 | 0.35 6 | Coatomer subunit alpha-1 |
| contig_29269_ 4 | 0.673 | 1 | 1 | 0.80 75 | 1 | 1 | 0.35 6 | predicted protein |
| contig_23827_ 5 | 0.6732 | 1 | 1 | 0.80 75 | 1 | 1 | 0.35 6 | predicted protein |
| contig_30717_ 5 | 0.6738 | 1 | 1 | 0.80 76 | 1 | 1 | 0.35 61 | predicted protein |
| contig_37765_ 6 | 0.6768 | 1 | 1 | 0.81 01 | 1 | 1 | 0.35 72 | serine/threonine-protein phosphatase 2A regulatory subunit B' |
| contig_9486_3 | 0.6775 | 1 | 1 | 0.81 01 | 1 | 1 | 0.35 72 | thiamine biosynthesis protein G (chloroplast) |
| contig_27473_ 6 | 0.6775 | 1 | 1 | 0.81 01 | 1 | 1 | 0.35 72 | peptidylprolyl isomerase |
| contig_38574_ 3 | 0.6792 | 1 | 1 | 0.81 14 | 1 | 1 | 0.35 78 | predicted protein |
| contig_10682_ 1 | 0.6797 | 1 | 1 | 0.81 14 | 1 | 1 | 0.35 78 | predicted protein |
| contig_14990_ 5 | 0.6804 | 1 | 1 | 0.81 17 | 1 | 1 | 0.35 79 | heme oxygenase 1 chloroplastic-like |
| contig_5949_4 | 0.6816 | 1 | 1 | 0.81 25 | 1 | 1 | 0.35 82 | predicted protein |
| contig_5530_1 | 0.6867 | 1 | 1 | 0.81 63 | 1 | 1 | 0.35 99 | NAD(P)-dependent steroid dehydrogenase-like protein |
| contig_5519_5 | 0.6867 | 1 | 1 | 0.81 63 | 1 | 1 | 0.35 99 | protein tesmin/TSO1-like CXC 5 |
| contig_25557_ 4 | 0.6869 | 1 | 1 | 0.81 63 | 1 | 1 | 0.35 99 | nucleolar complex protein 2 |
| contig_36071_ 6 | 0.687 | 1 | 1 | 0.81 63 | 1 | 1 | 0.35 99 | predicted protein |
| contig_6467_5 | 0.6884 | 1 | 1 | 0.81 68 | 1 | 1 | 0.36 01 | predicted protein |
| contig_22327_ 4 | 0.6888 | 1 | 1 | 0.81 68 | 1 | 1 | 0.36 01 | gamma-aminobutyric-acid receptor subunit beta |
| contig_30076_ 1 | 0.689 | 1 | 1 | 0.81 68 | 1 | 1 | 0.36 01 | elongation factor 2 |
| contig_5781_3 | 0.6903 | 1 | 1 | 0.81 77 | 1 | 1 | 0.36 05 | acetyl-coenzyme a synthetase |
| contig_8332_1 | 0.6913 | 1 | 1 | 0.81 82 | 1 | 1 | 0.36 07 | oxoglutarate dehydrogenase (succinyl-transferring) E1 component |
| contig_36999_ 3 | 0.6922 | 1 | 1 | 0.81 87 | 1 | 1 | 0.36 1 | predicted protein |

| | | | | | | | | |
|---|---|---|---|---|---|---|---|---|
| contig_30482_3 | 0.6939 | 1 | 1 | 0.8201 | 1 | 1 | 0.3616 | predicted protein |
| contig_39259_1 | 0.6955 | 1 | 1 | 0.8213 | 1 | 1 | 0.3621 | preprotein-translocase subunit a (chloroplast) |
| contig_38019_3 | 0.6965 | 1 | 1 | 0.8219 | 1 | 1 | 0.3624 | ferredoxin--NADP+ reductase |
| contig_12625_2 | 0.698 | 1 | 1 | 0.8229 | 1 | 1 | 0.3628 | 26S proteasome non-ATPase regulatory subunit 7 |
| contig_11835_4 | 0.7005 | 1 | 1 | 0.8247 | 1 | 1 | 0.3636 | ion transporter |
| contig_22787_6 | 0.7005 | 1 | 1 | 0.8247 | 1 | 1 | 0.3636 | bifunctional aspartate kinase/homoserine dehydrogenase I |
| contig_23621_6 | 0.7017 | 1 | 1 | 0.8254 | 1 | 1 | 0.3639 | predicted protein |
| contig_6009_6 | 0.7027 | 1 | 1 | 0.826 | 1 | 1 | 0.3642 | 40S ribosomal protein S5 |
| contig_23244_4 | 0.7033 | 1 | 1 | 0.826 | 1 | 1 | 0.3642 | predicted protein |
| contig_26772_5 | 0.7041 | 1 | 1 | 0.8261 | 1 | 1 | 0.3642 | predicted protein |
| contig_36096_1 | 0.7045 | 1 | 1 | 0.8261 | 1 | 1 | 0.3642 | predicted protein |
| contig_33176_3 | 0.7055 | 1 | 1 | 0.8267 | 1 | 1 | 0.3645 | EF-hand domain-containing protein |
| contig_15734_6 | 0.7107 | 1 | 1 | 0.8318 | 1 | 1 | 0.3668 | ALA dehydratase |
| contig_26990_2 | 0.711 | 1 | 1 | 0.8318 | 1 | 1 | 0.3668 | uracil phosphoribosyltransferase |
| contig_25589_3 | 0.7128 | 1 | 1 | 0.8328 | 1 | 1 | 0.3672 | small subunit ribosomal protein S3Ae |
| contig_36385_4 | 0.7131 | 1 | 1 | 0.8328 | 1 | 1 | 0.3672 | ---NA--- |
| contig_34378_5 | 0.7139 | 1 | 1 | 0.8328 | 1 | 1 | 0.3672 | predicted protein |
| contig_14363_4 | 0.714 | 1 | 1 | 0.8328 | 1 | 1 | 0.3672 | predicted protein |
| contig_24237_6 | 0.7154 | 1 | 1 | 0.8332 | 1 | 1 | 0.3673 | predicted protein |
| contig_3444_2 | 0.7159 | 1 | 1 | 0.8332 | 1 | 1 | 0.3673 | protoporphyrin IX methyltransferase |
| contig_24802_3 | 0.716 | 1 | 1 | 0.8332 | 1 | 1 | 0.3673 | predicted protein |
| contig_33904_2 | 0.7181 | 1 | 1 | 0.835 | 1 | 1 | 0.3681 | pectate lyase |
| contig_15455_3 | 0.7195 | 1 | 1 | 0.836 | 1 | 1 | 0.3686 | 1 2-dihydroxy-3-keto-5-methylthiopentene dioxygenase |
| contig_6653_5 | 0.7216 | 1 | 1 | 0.8365 | 1 | 1 | 0.3688 | chorismate mutase |
| contig_35860_4 | 0.7216 | 1 | 1 | 0.8365 | 1 | 1 | 0.3688 | mitochondrial manganese superoxide dismutase |
| contig_34741_6 | 0.7226 | 1 | 1 | 0.8365 | 1 | 1 | 0.3688 | 60S ribosomal protein |
| contig_9387_4 | 0.7229 | 1 | 1 | 0.8365 | 1 | 1 | 0.3688 | DUF791-domain-containing protein |

| | | | | | | | | |
|---|---|---|---|---|---|---|---|---|
| contig_18489_1 | 0.7234 | 1 | 1 | 0.8365 | 1 | 1 | 0.3688 | predicted protein |
| contig_27847_4 | 0.7241 | 1 | 1 | 0.8365 | 1 | 1 | 0.3688 | unnamed protein product |
| contig_10034_3 | 0.7241 | 1 | 1 | 0.8365 | 1 | 1 | 0.3688 | 60S ribosomal protein L10 |
| contig_34791_3 | 0.7243 | 1 | 1 | 0.8365 | 1 | 1 | 0.3688 | Regulator of chromosome condensation 1/beta-lactamase-inhibitor protein II |
| contig_38406_2 | 0.7252 | 1 | 1 | 0.8368 | 1 | 1 | 0.369 | predicted protein |
| contig_34200_5 | 0.7274 | 1 | 1 | 0.8388 | 1 | 1 | 0.3698 | fructose-1 6-bisphosphatase |
| contig_6058_5 | 0.7283 | 1 | 1 | 0.8392 | 1 | 1 | 0.37 | predicted protein |
| contig_33655_6 | 0.7289 | 1 | 1 | 0.8392 | 1 | 1 | 0.37 | predicted protein |
| contig_26112_6 | 0.73 | 1 | 1 | 0.8399 | 1 | 1 | 0.3703 | adenylate kinase 5 chloroplastic isoform X1 |
| contig_9488_1 | 0.7316 | 1 | 1 | 0.8411 | 1 | 1 | 0.3708 | 26s protease regulatory subunit 8 |
| contig_27998_6 | 0.7357 | 1 | 1 | 0.8451 | 1 | 1 | 0.3726 | 2 3-bisphosphoglycerate-dependent phosphoglycerate mutase |
| contig_27144_6 | 0.7377 | 1 | 1 | 0.8469 | 1 | 1 | 0.3734 | predicted protein |
| contig_26683_6 | 0.7397 | 1 | 1 | 0.8484 | 1 | 1 | 0.3741 | predicted protein |
| contig_17493_4 | 0.7408 | 1 | 1 | 0.8491 | 1 | 1 | 0.3744 | lamin tail domain-containing protein |
| contig_27657_5 | 0.7428 | 1 | 1 | 0.8508 | 1 | 1 | 0.3751 | solute carrier family 35 member E1 |
| contig_40882_2 | 0.7439 | 1 | 1 | 0.851 | 1 | 1 | 0.3752 | predicted protein |
| contig_1045_6 | 0.7441 | 1 | 1 | 0.851 | 1 | 1 | 0.3752 | iron-sulfur cluster formation ABC transporter (chloroplast) |
| contig_30686_5 | 0.7462 | 1 | 1 | 0.8528 | 1 | 1 | 0.376 | chlorophyll synthase chloroplastic |
| contig_7021_2 | 0.7485 | 1 | 1 | 0.8547 | 1 | 1 | 0.3768 | protein phosphatase 2C putative |
| contig_27168_5 | 0.7505 | 1 | 1 | 0.8555 | 1 | 1 | 0.3772 | predicted protein |
| contig_39293_4 | 0.7509 | 1 | 1 | 0.8555 | 1 | 1 | 0.3772 | predicted protein |
| contig_29986_5 | 0.7512 | 1 | 1 | 0.8555 | 1 | 1 | 0.3772 | H - or Na -translocating F-type V-type and A-type ATPase (F-ATPase) Superfamily |
| contig_32560_1 | 0.7516 | 1 | 1 | 0.8555 | 1 | 1 | 0.3772 | predicted protein |
| contig_23752_4 | 0.7521 | 1 | 1 | 0.8555 | 1 | 1 | 0.3772 | predicted protein |
| contig_27320_3 | 0.7529 | 1 | 1 | 0.8555 | 1 | 1 | 0.3772 | transketolase |
| contig_35543_6 | 0.7532 | 1 | 1 | 0.8555 | 1 | 1 | 0.3772 | ferredoxin--NADP+ reductase |

| | | | | | | | | |
|---|---|---|---|---|---|---|---|---|
| contig_39210_6 | 0.7573 | 1 | 1 | 0.8592 | 1 | 1 | 0.3788 | predicted protein |
| contig_39585_3 | 0.7576 | 1 | 1 | 0.8592 | 1 | 1 | 0.3788 | predicted protein |
| contig_34883_3 | 0.7586 | 1 | 1 | 0.8597 | 1 | 1 | 0.3791 | Fructose-bisphosphate aldolase |
| contig_27972_6 | 0.7603 | 1 | 1 | 0.861 | 1 | 1 | 0.3796 | atp-dependent rna helicase dbp5 |
| contig_16053_5 | 0.7624 | 1 | 1 | 0.8623 | 1 | 1 | 0.3802 | predicted protein |
| contig_40102_3 | 0.7628 | 1 | 1 | 0.8623 | 1 | 1 | 0.3802 | predicted protein |
| contig_21612_4 | 0.7636 | 1 | 1 | 0.8623 | 1 | 1 | 0.3802 | 60S ribosomal protein L27 |
| contig_21238_5 | 0.7636 | 1 | 1 | 0.8623 | 1 | 1 | 0.3802 | fucoxanthin chlorophyll a/c protein 8 |
| contig_13497_3 | 0.7685 | 1 | 1 | 0.8662 | 1 | 1 | 0.3819 | predicted protein |
| contig_34082_5 | 0.7686 | 1 | 1 | 0.8662 | 1 | 1 | 0.3819 | predicted protein |
| contig_25375_6 | 0.7688 | 1 | 1 | 0.8662 | 1 | 1 | 0.3819 | predicted protein |
| contig_33877_5 | 0.7699 | 1 | 1 | 0.8667 | 1 | 1 | 0.3821 | ---NA--- |
| contig_39943_5 | 0.7704 | 1 | 1 | 0.8667 | 1 | 1 | 0.3821 | mitochondrial alternative oxidase |
| contig_7026_3 | 0.771 | 1 | 1 | 0.8667 | 1 | 1 | 0.3821 | 26S proteasome non-ATPase regulatory subunit 2 |
| contig_13540_6 | 0.7722 | 1 | 1 | 0.8671 | 1 | 1 | 0.3823 | RNA polymerase beta subunit (chloroplast) |
| contig_7101_4 | 0.7726 | 1 | 1 | 0.8671 | 1 | 1 | 0.3823 | DNA-directed RNA polymerase II 19 kDa polypeptide |
| contig_13893_5 | 0.773 | 1 | 1 | 0.8671 | 1 | 1 | 0.3823 | NADH dehydrogenase subunit 11 (mitochondrion) |
| contig_8256_2 | 0.7749 | 1 | 1 | 0.8685 | 1 | 1 | 0.3829 | importin subunit beta putative |
| contig_25782_5 | 0.7764 | 1 | 1 | 0.8696 | 1 | 1 | 0.3834 | predicted protein |
| contig_9129_6 | 0.778 | 1 | 1 | 0.8708 | 1 | 1 | 0.3839 | predicted protein |
| contig_41308_6 | 0.779 | 1 | 1 | 0.8712 | 1 | 1 | 0.3841 | nucleoside diphosphate kinase |
| contig_34760_6 | 0.7799 | 1 | 1 | 0.8716 | 1 | 1 | 0.3843 | low molecular mass early light-inducible protein HV60 |
| contig_7125_2 | 0.7806 | 1 | 1 | 0.8717 | 1 | 1 | 0.3843 | cysteine proteinase 3-like |
| contig_8511_4 | 0.7824 | 1 | 1 | 0.8731 | 1 | 1 | 0.385 | O-acetylhomoserine aminocarboxypropyltransferase |
| contig_28443_2 | 0.7835 | 1 | 1 | 0.8735 | 1 | 1 | 0.3851 | nad-dependent epimerase/dehydratase |
| contig_28392_3 | 0.7839 | 1 | 1 | 0.8735 | 1 | 1 | 0.3851 | predicted protein |
| contig_6255_2 | 0.7873 | 1 | 1 | 0.8766 | 1 | 1 | 0.3865 | Nuclear transport factor 2 |

| | | | | | | | | |
|---|---|---|---|---|---|---|---|---|
| contig_37814_1 | 0.7891 | 1 | 1 | 0.878 | 1 | 1 | 0.3871 | ribosomal protein S3 (chloroplast) |
| contig_24756_6 | 0.7906 | 1 | 1 | 0.879 | 1 | 1 | 0.3875 | tropinone reductase |
| contig_30510_4 | 0.7946 | 1 | 1 | 0.8827 | 1 | 1 | 0.3892 | 60S ribosomal protein |
| contig_16920_6 | 0.7954 | 1 | 1 | 0.8827 | 1 | 1 | 0.3892 | predicted protein |
| contig_24876_6 | 0.7958 | 1 | 1 | 0.8827 | 1 | 1 | 0.3892 | predicted protein |
| contig_14698_3 | 0.7963 | 1 | 1 | 0.8827 | 1 | 1 | 0.3892 | AP-1 complex subunit gamma-1 |
| contig_39464_1 | 0.7975 | 1 | 1 | 0.8834 | 1 | 1 | 0.3895 | histone deacetylase 1 |
| contig_33315_3 | 0.7986 | 1 | 1 | 0.8834 | 1 | 1 | 0.3895 | gametolysin peptidase M11 |
| contig_28112_2 | 0.7987 | 1 | 1 | 0.8834 | 1 | 1 | 0.3895 | predicted protein |
| contig_39337_5 | 0.7996 | 1 | 1 | 0.8834 | 1 | 1 | 0.3895 | ---NA--- |
| contig_22675_1 | 0.8002 | 1 | 1 | 0.8834 | 1 | 1 | 0.3895 | predicted protein |
| contig_12035_1 | 0.8006 | 1 | 1 | 0.8834 | 1 | 1 | 0.3895 | predicted protein |
| contig_9857_5 | 0.801 | 1 | 1 | 0.8834 | 1 | 1 | 0.3895 | ---NA--- |
| contig_16580_4 | 0.8018 | 1 | 1 | 0.8837 | 1 | 1 | 0.3896 | casein kinase I |
| contig_13952_4 | 0.8044 | 1 | 1 | 0.886 | 1 | 1 | 0.3906 | predicted protein |
| contig_28019_5 | 0.806 | 1 | 1 | 0.887 | 1 | 1 | 0.3911 | NAD+ synthetase |
| contig_23311_6 | 0.8078 | 1 | 1 | 0.8881 | 1 | 1 | 0.3916 | predicted protein |
| contig_30562_5 | 0.8081 | 1 | 1 | 0.8881 | 1 | 1 | 0.3916 | cytochrome b6-f complex iron-sulfur subunit |
| contig_17954_4 | 0.8095 | 1 | 1 | 0.8881 | 1 | 1 | 0.3916 | predicted protein |
| contig_30385_5 | 0.8097 | 1 | 1 | 0.8881 | 1 | 1 | 0.3916 | myo-inositol-1(or 4)-monophosphatase |
| contig_15155_6 | 0.8108 | 1 | 1 | 0.8881 | 1 | 1 | 0.3916 | predicted protein |
| contig_29342_2 | 0.8113 | 1 | 1 | 0.8881 | 1 | 1 | 0.3916 | leukotriene A-4 hydrolase/aminopeptidase |
| contig_12287_5 | 0.8122 | 1 | 1 | 0.8881 | 1 | 1 | 0.3916 | predicted protein |
| contig_36065_2 | 0.8124 | 1 | 1 | 0.8881 | 1 | 1 | 0.3916 | Translation elongation factor IF5A |
| contig_29395_5 | 0.8127 | 1 | 1 | 0.8881 | 1 | 1 | 0.3916 | triosephosphate isomerase |
| contig_5361_1 | 0.8132 | 1 | 1 | 0.8881 | 1 | 1 | 0.3916 | ---NA--- |
| contig_25647_1 | 0.8134 | 1 | 1 | 0.8881 | 1 | 1 | 0.3916 | putative all-trans-retinol 13 14-reductase |

| | | | | | | | | |
|---|---|---|---|---|---|---|---|---|
| contig_6495_6 | 0.8144 | 1 | 1 | 0.8886 | 1 | 1 | 0.3918 | predicted protein |
| contig_38591_1 | 0.8166 | 1 | 1 | 0.8903 | 1 | 1 | 0.3925 | 40S ribosomal protein S18 |
| contig_28775_6 | 0.8191 | 1 | 1 | 0.8925 | 1 | 1 | 0.3935 | cullin-associated NEDD8-dissociated protein 1 |
| contig_5469_5 | 0.8198 | 1 | 1 | 0.8926 | 1 | 1 | 0.3935 | ribosomal protein S2 (chloroplast) |
| contig_17493_5 | 0.8207 | 1 | 1 | 0.8929 | 1 | 1 | 0.3937 | lamin tail domain-containing protein |
| contig_19427_5 | 0.8217 | 1 | 1 | 0.8934 | 1 | 1 | 0.3939 | 60S ribosomal protein L18a |
| contig_30406_3 | 0.8227 | 1 | 1 | 0.8938 | 1 | 1 | 0.3941 | Glycylpeptide N-tetradecanoyltransferase 2 |
| contig_17520_3 | 0.8247 | 1 | 1 | 0.8945 | 1 | 1 | 0.3944 | methyltransferase FkbM family (macronuclear) |
| contig_30519_2 | 0.8249 | 1 | 1 | 0.8945 | 1 | 1 | 0.3944 | predicted protein |
| contig_11186_4 | 0.8256 | 1 | 1 | 0.8945 | 1 | 1 | 0.3944 | predicted protein |
| contig_23306_1 | 0.8257 | 1 | 1 | 0.8945 | 1 | 1 | 0.3944 | ATP:ADP antiporter AAA family |
| contig_38073_1 | 0.8267 | 1 | 1 | 0.8946 | 1 | 1 | 0.3944 | ---NA--- |
| contig_24829_2 | 0.8269 | 1 | 1 | 0.8946 | 1 | 1 | 0.3944 | ornithine carbamoyltransferase |
| contig_9059_3 | 0.831 | 1 | 1 | 0.8968 | 1 | 1 | 0.3954 | predicted protein |
| contig_22874_2 | 0.8311 | 1 | 1 | 0.8968 | 1 | 1 | 0.3954 | inositol 5-phosphatase |
| contig_10909_2 | 0.8315 | 1 | 1 | 0.8968 | 1 | 1 | 0.3954 | Glycerol-3-phosphate dehydrogenase mitochondrial |
| contig_5869_1 | 0.8318 | 1 | 1 | 0.8968 | 1 | 1 | 0.3954 | threonine-tRNA ligase |
| contig_34431_6 | 0.8319 | 1 | 1 | 0.8968 | 1 | 1 | 0.3954 | predicted protein |
| contig_5863_5 | 0.8396 | 1 | 1 | 0.9044 | 1 | 1 | 0.3987 | predicted protein |
| contig_9759_2 | 0.8408 | 1 | 1 | 0.905 | 1 | 1 | 0.399 | clathrin heavy chain |
| contig_17080_5 | 0.8423 | 1 | 1 | 0.9061 | 1 | 1 | 0.3995 | predicted protein |
| contig_22809_5 | 0.8464 | 1 | 1 | 0.9098 | 1 | 1 | 0.4011 | predicted protein |
| contig_33750_4 | 0.847 | 1 | 1 | 0.9098 | 1 | 1 | 0.4011 | Violaxanthin de-epoxidase |
| contig_11742_4 | 0.8485 | 1 | 1 | 0.9108 | 1 | 1 | 0.4015 | predicted protein |
| contig_23607_2 | 0.8533 | 1 | 1 | 0.9134 | 1 | 1 | 0.4027 | alanyl-tRNA synthetase |
| contig_33083_6 | 0.8534 | 1 | 1 | 0.9134 | 1 | 1 | 0.4027 | Glucose-6-phosphate isomerase cytosolic |
| contig_30540_5 | 0.8536 | 1 | 1 | 0.9134 | 1 | 1 | 0.4027 | unnamed protein product |

| | | | | | | | | |
|---|---|---|---|---|---|---|---|---|
| contig_6105_6 | 0.8537 | 1 | 1 | 0.9134 | 1 | 1 | 0.4027 | photosystem II CP47 chlorophyll apoprotein (chloroplast) |
| contig_14542_6 | 0.8539 | 1 | 1 | 0.9134 | 1 | 1 | 0.4027 | predicted protein |
| contig_40291_3 | 0.8559 | 1 | 1 | 0.9143 | 1 | 1 | 0.4031 | predicted protein |
| contig_16980_6 | 0.8562 | 1 | 1 | 0.9143 | 1 | 1 | 0.4031 | translation initiation factor 3 subunit A |
| contig_6303_5 | 0.8571 | 1 | 1 | 0.9143 | 1 | 1 | 0.4031 | acyl carrier protein (chloroplast) |
| contig_20747_6 | 0.8572 | 1 | 1 | 0.9143 | 1 | 1 | 0.4031 | translation initiation factor 4E |
| contig_16462_2 | 0.8578 | 1 | 1 | 0.9143 | 1 | 1 | 0.4031 | chorismate synthase |
| contig_39369_3 | 0.8586 | 1 | 1 | 0.9144 | 1 | 1 | 0.4032 | hypothetical protein FRACYDRAFT_269748 |
| contig_2955_5 | 0.8591 | 1 | 1 | 0.9144 | 1 | 1 | 0.4032 | NADH dehydrogenase (ubiquinone) flavoprotein 2 |
| contig_19158_3 | 0.8627 | 1 | 1 | 0.9176 | 1 | 1 | 0.4046 | small ubiquitin-like modifier (SUMO) |
| contig_19141_5 | 0.8648 | 1 | 1 | 0.9181 | 1 | 1 | 0.4048 | photosystem I subunit IV (chloroplast) |
| contig_14343_6 | 0.8655 | 1 | 1 | 0.9181 | 1 | 1 | 0.4048 | riboflavin synthase alpha subunit |
| contig_6474_4 | 0.8657 | 1 | 1 | 0.9181 | 1 | 1 | 0.4048 | FAD-dependent monooxygenase |
| contig_8636_5 | 0.8657 | 1 | 1 | 0.9181 | 1 | 1 | 0.4048 | isocitrate dehydrogenase |
| contig_9214_2 | 0.8662 | 1 | 1 | 0.9181 | 1 | 1 | 0.4048 | AMP-dependent synthetase |
| contig_36986_1 | 0.8691 | 1 | 1 | 0.9206 | 1 | 1 | 0.4059 | ---NA--- |
| contig_30863_3 | 0.8755 | 1 | 1 | 0.9267 | 1 | 1 | 0.4086 | predicted protein |
| contig_9486_1 | 0.8787 | 1 | 1 | 0.9294 | 1 | 1 | 0.4098 | thiamine biosynthesis protein G (chloroplast) |
| contig_15845_2 | 0.8815 | 1 | 1 | 0.9317 | 1 | 1 | 0.4108 | predicted protein |
| contig_32725_2 | 0.8824 | 1 | 1 | 0.932 | 1 | 1 | 0.4109 | predicted protein |
| contig_8596_6 | 0.8836 | 1 | 1 | 0.9326 | 1 | 1 | 0.4112 | predicted protein |
| contig_5826_3 | 0.8842 | 1 | 1 | 0.9326 | 1 | 1 | 0.4112 | 60S ribosomal protein L2 |
| contig_29914_2 | 0.8928 | 1 | 1 | 0.9411 | 1 | 1 | 0.4149 | predicted protein |
| contig_5663_5 | 0.8943 | 1 | 1 | 0.9416 | 1 | 1 | 0.4151 | tubulin beta chain |
| contig_39949_2 | 0.8945 | 1 | 1 | 0.9416 | 1 | 1 | 0.4151 | predicted protein |
| contig_34883_6 | 0.8962 | 1 | 1 | 0.9427 | 1 | 1 | 0.4156 | Fructose-bisphosphate aldolase |
| contig_6039_6 | 0.8988 | 1 | 1 | 0.9447 | 1 | 1 | 0.4165 | large subunit ribosomal protein L27Ae |

| | | | | | | | | |
|---|---|---|---|---|---|---|---|---|
| contig_26282_6 | 0.9003 | 1 | 1 | 0.9454 | 1 | 1 | 0.4168 | dihydrodipicolinate reductase |
| contig_26918_5 | 0.9006 | 1 | 1 | 0.9454 | 1 | 1 | 0.4168 | predicted protein |
| contig_5425_1 | 0.9018 | 1 | 1 | 0.946 | 1 | 1 | 0.4171 | sulfate adenylyltransferase |
| contig_21789_5 | 0.9036 | 1 | 1 | 0.9464 | 1 | 1 | 0.4173 | Mitochondrial Carrier (MC) Family |
| contig_38474_6 | 0.9038 | 1 | 1 | 0.9464 | 1 | 1 | 0.4173 | phosphoserine transaminase |
| contig_13878_2 | 0.9041 | 1 | 1 | 0.9464 | 1 | 1 | 0.4173 | glycine--tRNA ligase |
| contig_26089_5 | 0.9057 | 1 | 1 | 0.9474 | 1 | 1 | 0.4177 | exportin-like protein |
| contig_10572_1 | 0.9081 | 1 | 1 | 0.9488 | 1 | 1 | 0.4183 | NADH dehydrogenase [ubiquinone] flavoprotein 1 mitochondrial |
| contig_7412_1 | 0.9083 | 1 | 1 | 0.9488 | 1 | 1 | 0.4183 | E3 ubiquitin-protein ligase RING2-like |
| contig_23471_1 | 0.911 | 1 | 1 | 0.9505 | 1 | 1 | 0.4191 | predicted protein |
| contig_28636_6 | 0.9111 | 1 | 1 | 0.9505 | 1 | 1 | 0.4191 | Small nuclear ribonucleoprotein G |
| contig_30869_3 | 0.9141 | 1 | 1 | 0.9526 | 1 | 1 | 0.42 | predicted protein |
| contig_33544_3 | 0.915 | 1 | 1 | 0.9526 | 1 | 1 | 0.42 | predicted protein |
| contig_3511_5 | 0.9151 | 1 | 1 | 0.9526 | 1 | 1 | 0.42 | Ycf89 (chloroplast) |
| contig_16936_1 | 0.9162 | 1 | 1 | 0.9526 | 1 | 1 | 0.42 | H/ACA ribonucleoprotein complex subunit 4 |
| contig_10866_1 | 0.9164 | 1 | 1 | 0.9526 | 1 | 1 | 0.42 | predicted protein |
| contig_25855_2 | 0.9178 | 1 | 1 | 0.9526 | 1 | 1 | 0.42 | protein transport protein Sec61 subunit gamma-like |
| contig_25457_5 | 0.9178 | 1 | 1 | 0.9526 | 1 | 1 | 0.42 | farnesyl-diphosphate farnesyltransferase |
| contig_18889_1 | 0.9186 | 1 | 1 | 0.9526 | 1 | 1 | 0.42 | ---NA--- |
| contig_24544_3 | 0.9188 | 1 | 1 | 0.9526 | 1 | 1 | 0.42 | predicted protein |
| contig_10149_5 | 0.924 | 1 | 1 | 0.9566 | 1 | 1 | 0.4218 | phosphoribosylformylglycinamidine synthase |
| contig_22796_6 | 0.9242 | 1 | 1 | 0.9566 | 1 | 1 | 0.4218 | predicted protein |
| contig_27983_4 | 0.9247 | 1 | 1 | 0.9566 | 1 | 1 | 0.4218 | alpha/beta hydrolase |
| contig_10600_6 | 0.9255 | 1 | 1 | 0.9566 | 1 | 1 | 0.4218 | predicted protein |
| contig_9755_4 | 0.9258 | 1 | 1 | 0.9566 | 1 | 1 | 0.4218 | valyl-tRNA synthetase |
| contig_16105_5 | 0.9273 | 1 | 1 | 0.9575 | 1 | 1 | 0.4222 | hypothetical protein FRACYDRAFT_264519 |
| contig_25825_6 | 0.9291 | 1 | 1 | 0.9584 | 1 | 1 | 0.4225 | predicted protein |

| | | | | | | | | |
|---|---|---|---|---|---|---|---|---|
| contig_28195_3 | 0.9294 | 1 | 1 | 0.9584 | 1 | 1 | 0.4225 | predicted protein |
| contig_28484_6 | 0.931 | 1 | 1 | 0.9594 | 1 | 1 | 0.423 | carbamoyl-phosphate synthase (glutamine-hydrolyzing) large subunit |
| contig_19155_3 | 0.9317 | 1 | 1 | 0.9595 | 1 | 1 | 0.423 | geranylgeranyl reductase |
| contig_6937_2 | 0.9332 | 1 | 1 | 0.96 | 1 | 1 | 0.4232 | predicted protein |
| contig_5926_5 | 0.9335 | 1 | 1 | 0.96 | 1 | 1 | 0.4232 | predicted protein |
| contig_39431_2 | 0.9345 | 1 | 1 | 0.9602 | 1 | 1 | 0.4234 | GDP-mannose transporter |
| contig_9849_4 | 0.935 | 1 | 1 | 0.9602 | 1 | 1 | 0.4234 | chloroplast sedoheptulose-1 7-bisphosphatase |
| contig_25573_5 | 0.9377 | 1 | 1 | 0.9624 | 1 | 1 | 0.4243 | predicted protein |
| contig_36089_2 | 0.9387 | 1 | 1 | 0.9624 | 1 | 1 | 0.4243 | archaeal ribosomal protein S17P |
| contig_10169_3 | 0.939 | 1 | 1 | 0.9624 | 1 | 1 | 0.4243 | NADH-cytochrome b5 reductase 3-like |
| contig_6079_5 | 0.9399 | 1 | 1 | 0.9626 | 1 | 1 | 0.4244 | predicted protein |
| contig_35608_4 | 0.9424 | 1 | 1 | 0.9646 | 1 | 1 | 0.4253 | chaperone dnaj-like protein |
| contig_10808_6 | 0.9442 | 1 | 1 | 0.9657 | 1 | 1 | 0.4258 | predicted protein |
| contig_11202_6 | 0.9447 | 1 | 1 | 0.9657 | 1 | 1 | 0.4258 | predicted protein |
| contig_19117_2 | 0.9464 | 1 | 1 | 0.9667 | 1 | 1 | 0.4262 | 60S ribosomal protein L21-A |
| contig_41371_1 | 0.9496 | 1 | 1 | 0.9692 | 1 | 1 | 0.4273 | 40s ribosomal protein |
| contig_17633_4 | 0.9509 | 1 | 1 | 0.9692 | 1 | 1 | 0.4273 | Adenylyl cyclase-associated protein |
| contig_37721_1 | 0.9512 | 1 | 1 | 0.9692 | 1 | 1 | 0.4273 | ---NA--- |
| contig_25820_1 | 0.9515 | 1 | 1 | 0.9692 | 1 | 1 | 0.4273 | predicted protein |
| contig_16841_6 | 0.9524 | 1 | 1 | 0.9692 | 1 | 1 | 0.4273 | predicted protein |
| contig_33453_2 | 0.9526 | 1 | 1 | 0.9692 | 1 | 1 | 0.4273 | predicted protein |
| contig_22713_3 | 0.955 | 1 | 1 | 0.971 | 1 | 1 | 0.4281 | predicted protein |
| contig_13739_4 | 0.9567 | 1 | 1 | 0.972 | 1 | 1 | 0.4286 | protein DETOXIFICATION 46 chloroplastic |
| contig_7217_3 | 0.9583 | 1 | 1 | 0.973 | 1 | 1 | 0.429 | P-loop containing nucleoside triphosphate hydrolase protein |
| contig_26865_4 | 0.96 | 1 | 1 | 0.9741 | 1 | 1 | 0.4295 | ---NA--- |
| contig_24778_4 | 0.9614 | 1 | 1 | 0.9745 | 1 | 1 | 0.4297 | predicted protein |
| contig_40323_4 | 0.9617 | 1 | 1 | 0.9745 | 1 | 1 | 0.4297 | ---NA--- |

| | | | | | | | | |
|---|---|---|---|---|---|---|---|---|
| contig_29362_6 | 0.9633 | 1 | 1 | 0.9755 | 1 | 1 | 0.4301 | Stimulated by retinoic acid gene 6 protein |
| contig_30555_4 | 0.9644 | 1 | 1 | 0.9759 | 1 | 1 | 0.4303 | fructose-1 6-bisphosphate aldolase precursor |
| contig_25157_1 | 0.9664 | 1 | 1 | 0.9773 | 1 | 1 | 0.4309 | magnesium chelatase subunit D |
| contig_24282_5 | 0.9689 | 1 | 1 | 0.9792 | 1 | 1 | 0.4317 | predicted protein |
| contig_33515_6 | 0.9708 | 1 | 1 | 0.9799 | 1 | 1 | 0.432 | predicted protein |
| contig_23787_5 | 0.9709 | 1 | 1 | 0.9799 | 1 | 1 | 0.432 | predicted protein |
| contig_33784_6 | 0.9741 | 1 | 1 | 0.9821 | 1 | 1 | 0.433 | predicted protein |
| contig_6257_1 | 0.9743 | 1 | 1 | 0.9821 | 1 | 1 | 0.433 | predicted protein |
| contig_8945_4 | 0.9751 | 1 | 1 | 0.9823 | 1 | 1 | 0.4331 | delta-9 acyl-desaturase |
| contig_28224_2 | 0.9766 | 1 | 1 | 0.9831 | 1 | 1 | 0.4334 | alpha tubulin |
| contig_30832_6 | 0.98 | 1 | 1 | 0.9856 | 1 | 1 | 0.4345 | predicted protein |
| contig_7031_3 | 0.9808 | 1 | 1 | 0.9856 | 1 | 1 | 0.4345 | transmembrane 9 superfamily member 3 |
| contig_38536_2 | 0.9815 | 1 | 1 | 0.9856 | 1 | 1 | 0.4345 | predicted protein |
| contig_27545_4 | 0.9817 | 1 | 1 | 0.9856 | 1 | 1 | 0.4345 | predicted protein |
| contig_26837_3 | 0.9833 | 1 | 1 | 0.9866 | 1 | 1 | 0.435 | AP-1 complex subunit gamma-1 |
| contig_28535_5 | 0.9843 | 1 | 1 | 0.9869 | 1 | 1 | 0.4351 | 26S proteasome non-ATPase regulatory subunit 2 |
| contig_29698_3 | 0.9875 | 1 | 1 | 0.9895 | 1 | 1 | 0.4363 | Light-harvesting complex I LH38 protein |
| contig_14932_3 | 0.9894 | 1 | 1 | 0.9907 | 1 | 1 | 0.4368 | predicted protein |
| contig_27279_6 | 0.9906 | 1 | 1 | 0.9913 | 0.9999 | 1 | 0.4371 | adenosine kinase |
| contig_28383_6 | 0.9946 | 1 | 1 | 0.9946 | 0.9946 | 1 | 0.4385 | predicted protein |

## Nt31 – Tt34

| proteinAccession | ttest_pvals | Binomial_SGoF | Conservative_SGoF | BH | SFisher | Bonferroni | q_val | Description |
|---|---|---|---|---|---|---|---|---|
| contig_22939_2 | 0 | 0 | 3E-04 | 0.0027 | 0 | 0.0027 | 0.0021 | DNA damage-binding protein 1 |
| contig_18884_5 | 0 | 1E-04 | 4E-04 | 0.0131 | 0 | 0.0261 | 0.01 | predicted protein |
| contig_9006_3 | 0 | 1E-04 | 4E-04 | 0.0189 | 0 | 0.0581 | 0.0145 | thiol-activated cytolysin |
| contig_7129_5 | 0 | 1E-04 | 5E-04 | 0.0189 | 0 | 0.0755 | 0.0145 | T9SS type A sorting domain-containing protein |

| | | | | | | | | |
|---|---|---|---|---|---|---|---|---|
| contig_24359_6 | 1E-04 | 1E-04 | 7E-04 | 0.0191 | 0 | 0.0954 | 0.0146 | phosphoenolpyruvate carboxylase |
| contig_30013_1 | 1E-04 | 2E-04 | 8E-04 | 0.0255 | 0 | 0.1586 | 0.0195 | ribonucleoside-diphosphate reductase subunit alpha |
| contig_3037_1 | 1E-04 | 2E-04 | 0.001 | 0.0255 | 0 | 0.203 | 0.0195 | ribosomal protein S4 (chloroplast) |
| contig_5654_2 | 1E-04 | 3E-04 | 0.0012 | 0.0255 | 0 | 0.2032 | 0.0195 | hypothetical protein THAOC_17223 partial |
| contig_20131_5 | 2E-04 | 3E-04 | 0.0012 | 0.0275 | 0 | 0.2546 | 0.021 | 60S ribosomal protein L21-A |
| contig_17954_4 | 2E-04 | 4E-04 | 0.0015 | 0.0275 | 0 | 0.2733 | 0.021 | predicted protein |
| contig_6836_6 | 2E-04 | 4E-04 | 0.0015 | 0.0299 | 0 | 0.3271 | 0.0229 | predicted protein |
| contig_24617_4 | 3E-04 | 5E-04 | 0.0016 | 0.0346 | 0 | 0.4377 | 0.0265 | predicted protein |
| contig_23093_4 | 3E-04 | 5E-04 | 0.0016 | 0.0346 | 0 | 0.4468 | 0.0265 | predicted protein |
| contig_35081_4 | 4E-04 | 8E-04 | 0.002 | 0.0392 | 0 | 0.5435 | 0.03 | ubiquinol oxidase 4 chloroplastic/chromoplastic |
| contig_13715_6 | 4E-04 | 8E-04 | 0.002 | 0.041 | 0 | 0.6091 | 0.0314 | dnaJ homolog subfamily B member 6 isoform X2 |
| contig_34524_6 | 4E-04 | 0.0011 | 0.0021 | 0.0425 | 0 | 0.6812 | 0.0325 | ---NA--- |
| contig_13540_6 | 5E-04 | 0.0012 | 0.0022 | 0.0425 | 0 | 0.7145 | 0.0325 | RNA polymerase beta subunit (chloroplast) |
| contig_24447_4 | 5E-04 | 0.0012 | 0.0022 | 0.0455 | 0 | 0.8104 | 0.0349 | diaminopimelate decarboxylase |
| contig_35135_5 | 7E-04 | 0.0013 | 0.0023 | 0.0561 | 0 | 1 | 0.043 | peptidyl-prolyl cis-trans isomerase A |
| contig_39276_6 | 8E-04 | 0.0015 | 0.0025 | 0.0593 | 0 | 1 | 0.0454 | orotidine 5'-phosphate decarboxylase |
| contig_33214_4 | 8E-04 | 0.0016 | 0.0025 | 0.0593 | 0 | 1 | 0.0454 | phosphoadenosine-phosphosulphate reductase |
| contig_10303_5 | 0.001 | 0.0016 | 0.0025 | 0.0709 | 0 | 1 | 0.0543 | 40S ribosomal protein S3-3 |
| contig_7982_4 | 0.0011 | 0.0016 | 0.0029 | 0.0715 | 0 | 1 | 0.0548 | photosystem II protein V (chloroplast) |
| contig_3037_3 | 0.0012 | 0.002 | 0.0029 | 0.075 | 0 | 1 | 0.0574 | ribosomal protein S4 (chloroplast) |
| contig_24799_5 | 0.0012 | 0.002 | 0.0031 | 0.075 | 0 | 1 | 0.0574 | predicted protein |
| contig_16094_5 | 0.0013 | 0.0021 | 0.0031 | 0.075 | 0 | 1 | 0.0574 | 40S ribosomal protein S23 |
| contig_22589_6 | 0.0015 | 0.0022 | 0.0031 | 0.0788 | 0 | 1 | 0.0604 | Peptidase_M16-domain-containing protein |
| contig_33684_3 | 0.0015 | 0.0022 | 0.0033 | 0.0788 | 0 | 1 | 0.0604 | nadh dehydrogenase |
| contig_36382_5 | 0.0015 | 0.0022 | 0.0033 | 0.0788 | 0 | 1 | 0.0604 | Ubiquitin-40S ribosomal protein S27a |
| contig_15800_4 | 0.0016 | 0.0023 | 0.0035 | 0.0788 | 0 | 1 | 0.0604 | asparagine synthase |
| contig_28919_4 | 0.0016 | 0.0025 | 0.0037 | 0.0788 | 0 | 1 | 0.0604 | adenosylhomocysteinase |

| | | | | | | | | |
|---|---|---|---|---|---|---|---|---|
| contig_8636_5 | 0.0019 | 0.0025 | 0.0037 | 0.0872 | 0 | 1 | 0.0668 | isocitrate dehydrogenase |
| contig_29933_6 | 0.002 | 0.0025 | 0.0042 | 0.0872 | 0 | 1 | 0.0668 | 30S ribosomal protein S1 |
| contig_12284_5 | 0.002 | 0.0029 | 0.0044 | 0.0872 | 0 | 1 | 0.0668 | ribosomal protein S9 |
| contig_9486_3 | 0.0021 | 0.0029 | 0.0045 | 0.0872 | 0 | 1 | 0.0668 | thiamine biosynthesis protein G (chloroplast) |
| contig_6525_2 | 0.0022 | 0.003 | 0.0046 | 0.0872 | 0 | 1 | 0.0668 | probable aminopeptidase NPEPL1 |
| contig_28927_1 | 0.0022 | 0.0031 | 0.0047 | 0.0872 | 0 | 1 | 0.0668 | 60S ribosomal protein |
| contig_29760_5 | 0.0022 | 0.0031 | 0.0052 | 0.0872 | 0 | 1 | 0.0668 | predicted protein |
| contig_38660_3 | 0.0023 | 0.0032 | 0.0054 | 0.0872 | 0 | 1 | 0.0668 | elongation factor 2 |
| contig_29801_5 | 0.0023 | 0.0033 | 0.0054 | 0.0872 | 0 | 1 | 0.0668 | triosephosphate isomerase |
| contig_11453_6 | 0.0025 | 0.0036 | 0.0054 | 0.0894 | 0 | 1 | 0.0685 | major facilitator transporter |
| contig_37972_6 | 0.0025 | 0.0036 | 0.0057 | 0.0894 | 0 | 1 | 0.0685 | predicted protein |
| contig_28307_4 | 0.0025 | 0.0037 | 0.0057 | 0.0894 | 0 | 1 | 0.0685 | atp-dependent rna helicase uap56 |
| contig_28236_1 | 0.0028 | 0.0037 | 0.006 | 0.0959 | 0 | 1 | 0.0735 | tRNA splicing endonuclease putative |
| contig_27595_5 | 0.0029 | 0.0044 | 0.006 | 0.0959 | 0 | 1 | 0.0735 | transitional endoplasmic reticulum ATPase |
| contig_1081_1 | 0.0029 | 0.0045 | 0.0061 | 0.0959 | 0 | 1 | 0.0735 | hypothetical protein CBC21_09400 |
| contig_24784_5 | 0.003 | 0.0045 | 0.0061 | 0.0959 | 0 | 1 | 0.0735 | DUF423-domain-containing protein |
| contig_40308_6 | 0.0031 | 0.0046 | 0.0065 | 0.0959 | 0 | 1 | 0.0735 | 40S ribosomal protein S6 |
| contig_21030_6 | 0.0031 | 0.0047 | 0.0065 | 0.0959 | 0 | 1 | 0.0735 | 60S ribosomal protein L39 |
| contig_28911_6 | 0.0031 | 0.0052 | 0.0066 | 0.0959 | 0 | 1 | 0.0735 | eukaryotic translation initiation factor 6 |
| contig_17463_2 | 0.0032 | 0.0054 | 0.0066 | 0.0959 | 0 | 1 | 0.0735 | glucose-6-phosphate isomerase |
| contig_30397_1 | 0.0033 | 0.0054 | 0.0067 | 0.0971 | 0 | 1 | 0.0744 | FAD-dependent monooxygenase |
| contig_5809_2 | 0.0035 | 0.0054 | 0.0067 | 0.0996 | 0 | 1 | 0.0763 | ---NA--- |
| contig_8777_3 | 0.0036 | 0.0057 | 0.007 | 0.0996 | 0 | 1 | 0.0763 | elongation factor G |
| contig_34416_5 | 0.0036 | 0.0057 | 0.0072 | 0.0996 | 0 | 1 | 0.0763 | predicted protein |
| contig_39090_5 | 0.0037 | 0.0059 | 0.0073 | 0.0996 | 0 | 1 | 0.0763 | Glycerol-3-phosphate dehydrogenase |
| contig_30076_1 | 0.0037 | 0.006 | 0.0074 | 0.0996 | 0 | 1 | 0.0763 | elongation factor 2 |
| contig_9849_4 | 0.0041 | 0.006 | 0.0074 | 0.108 | 0 | 1 | 0.0827 | chloroplast sedoheptulose-1 7-bisphosphatase |

| | | | | | | | | |
|---|---|---|---|---|---|---|---|---|
| contig_6708_5 | 0.0042 | 0.0061 | 0.008 | 0.108 | 0 | 1 | 0.0827 | hemagglutinin/amebocyte aggregation factor |
| contig_9486_1 | 0.0044 | 0.0061 | 0.0081 | 0.1109 | 0 | 1 | 0.0849 | thiamine biosynthesis protein G (chloroplast) |
| contig_10138_6 | 0.0045 | 0.0063 | 0.0083 | 0.1109 | 0 | 1 | 0.0849 | adenylate kinase chloroplastic |
| contig_26510_5 | 0.0045 | 0.0066 | 0.0098 | 0.1109 | 0 | 1 | 0.0849 | LL-diaminopimelate aminotransferase |
| contig_22011_5 | 0.0046 | 0.0066 | 0.0104 | 0.1111 | 0 | 1 | 0.0851 | argininosuccinate synthase |
| contig_39862_1 | 0.0047 | 0.0067 | 0.0108 | 0.1129 | 0 | 1 | 0.0865 | pyruvate kinase |
| contig_9499_6 | 0.0052 | 0.0067 | 0.0112 | 0.1201 | 0 | 1 | 0.0919 | predicted protein |
| contig_17741_5 | 0.0052 | 0.0067 | 0.0112 | 0.1201 | 0 | 1 | 0.0919 | endonuclease 4 |
| contig_16887_1 | 0.0054 | 0.007 | 0.0119 | 0.1201 | 0 | 1 | 0.0919 | predicted protein |
| contig_34697_6 | 0.0054 | 0.0072 | 0.012 | 0.1201 | 0 | 1 | 0.0919 | asparagine synthase |
| contig_23988_3 | 0.0054 | 0.0073 | 0.0127 | 0.1201 | 0 | 1 | 0.0919 | predicted protein |
| contig_27313_6 | 0.0056 | 0.0074 | 0.0128 | 0.121 | 0 | 1 | 0.0927 | predicted protein |
| contig_27603_4 | 0.0057 | 0.008 | 0.0137 | 0.121 | 0 | 1 | 0.0927 | predicted protein |
| contig_37814_1 | 0.0057 | 0.0081 | 0.0138 | 0.121 | 0 | 1 | 0.0927 | ribosomal protein S3 (chloroplast) |
| contig_25854_4 | 0.0059 | 0.0083 | 0.0143 | 0.121 | 0 | 1 | 0.0927 | L30e-like protein |
| contig_14542_6 | 0.006 | 0.0098 | 0.0144 | 0.121 | 0 | 1 | 0.0927 | predicted protein |
| contig_28628_1 | 0.006 | 0.0102 | 0.0144 | 0.121 | 0 | 1 | 0.0927 | phosphoribosylaminoimidazolecarboxamide formyltransferase/IMP cyclohydrolase |
| contig_30106_6 | 0.0061 | 0.0108 | 0.0147 | 0.121 | 0 | 1 | 0.0927 | predicted protein |
| contig_29723_5 | 0.0061 | 0.0112 | 0.0149 | 0.121 | 0 | 1 | 0.0927 | predicted protein |
| contig_11577_6 | 0.0063 | 0.0112 | 0.015 | 0.121 | 0 | 1 | 0.0927 | Mitochondrial adenine nucleotide transporter ADNT1 |
| contig_35016_2 | 0.0065 | 0.0119 | 0.0156 | 0.121 | 0 | 1 | 0.0927 | predicted protein |
| contig_5881_5 | 0.0065 | 0.0122 | 0.0172 | 0.121 | 0 | 1 | 0.0927 | elongation factor Tu (chloroplast) |
| contig_23071_3 | 0.0066 | 0.0128 | 0.0174 | 0.121 | 0 | 1 | 0.0927 | D-3-phosphoglycerate dehydrogenase |
| contig_30668_1 | 0.0066 | 0.0128 | 0.0174 | 0.121 | 0 | 1 | 0.0927 | predicted protein |
| contig_17134_1 | 0.0067 | 0.0137 | 0.0179 | 0.121 | 0 | 1 | 0.0927 | 60S ribosomal protein L22 |
| contig_37899_1 | 0.0067 | 0.0144 | 0.0179 | 0.121 | 0 | 1 | 0.0927 | ribosomal protein L2 (chloroplast) |

| | | | | | | | | |
|---|---|---|---|---|---|---|---|---|
| contig_24766_6 | 0.0067 | 0.0144 | 0.0181 | 0.121 | 0 | 1 | 0.0927 | nitrate reductase |
| contig_28995_4 | 0.007 | 0.0144 | 0.0187 | 0.1243 | 0 | 1 | 0.0952 | predicted protein |
| contig_15864_3 | 0.0071 | 0.0147 | 0.019 | 0.125 | 0 | 1 | 0.0957 | 50S ribosomal protein L15 |
| contig_27659_6 | 0.0072 | 0.015 | 0.0192 | 0.125 | 0 | 1 | 0.0957 | predicted protein |
| contig_27069_6 | 0.0073 | 0.0153 | 0.0196 | 0.1258 | 0 | 1 | 0.0963 | predicted protein |
| contig_19330_5 | 0.0074 | 0.0153 | 0.0196 | 0.1265 | 0 | 1 | 0.0969 | predicted protein |
| contig_32615_1 | 0.0079 | 0.0171 | 0.0202 | 0.1323 | 0 | 1 | 0.1013 | predicted protein |
| contig_33630_6 | 0.008 | 0.0174 | 0.0203 | 0.1323 | 0 | 1 | 0.1013 | actin |
| contig_9464_6 | 0.0081 | 0.0174 | 0.0203 | 0.1328 | 0 | 1 | 0.1017 | cytochrome c heme attachment protein (chloroplast) |
| contig_28171_1 | 0.0082 | 0.0179 | 0.0208 | 0.1331 | 0 | 1 | 0.1019 | predicted protein |
| contig_24655_5 | 0.0083 | 0.0179 | 0.0217 | 0.1334 | 0 | 1 | 0.1021 | predicted protein |
| contig_25730_2 | 0.0089 | 0.0181 | 0.0218 | 0.1425 | 0 | 1 | 0.1091 | predicted protein |
| contig_22708_3 | 0.0096 | 0.0187 | 0.0222 | 0.1508 | 0 | 1 | 0.1155 | Laminin G domain protein |
| contig_23416_6 | 0.0098 | 0.019 | 0.0229 | 0.1516 | 0 | 1 | 0.1161 | predicted protein |
| contig_9059_3 | 0.0098 | 0.0192 | 0.023 | 0.1516 | 0 | 1 | 0.1161 | predicted protein |
| contig_37313_1 | 0.0102 | 0.0195 | 0.023 | 0.1565 | 0 | 1 | 0.1198 | predicted protein |
| contig_28730_5 | 0.0104 | 0.0196 | 0.0236 | 0.1575 | 0 | 1 | 0.1206 | predicted protein |
| contig_9980_5 | 0.0108 | 0.02 | 0.0242 | 0.1615 | 0 | 1 | 0.1237 | protein fucoxanthin chlorophyll a/c protein |
| contig_37879_1 | 0.0109 | 0.0203 | 0.0242 | 0.1621 | 0 | 1 | 0.1242 | UPF0187 protein |
| contig_30486_2 | 0.0112 | 0.0203 | 0.0247 | 0.1634 | 0 | 1 | 0.1251 | signal peptidase I |
| contig_27073_1 | 0.0112 | 0.0208 | 0.0255 | 0.1634 | 0 | 1 | 0.1251 | Phospholipid hydroperoxide glutathione peroxidase |
| contig_6039_6 | 0.0118 | 0.0217 | 0.0255 | 0.1698 | 0 | 1 | 0.1301 | large subunit ribosomal protein L27Ae |
| contig_39493_5 | 0.0119 | 0.0218 | 0.0261 | 0.1701 | 0 | 1 | 0.1303 | 60s ribosomal protein |
| contig_29625_1 | 0.012 | 0.0218 | 0.0267 | 0.1701 | 0 | 1 | 0.1303 | rRNA 2'-O-methyltransferase fibrillarin |
| contig_38064_1 | 0.0122 | 0.0229 | 0.0273 | 0.1705 | 0 | 1 | 0.1306 | phosphoenolpyruvate carboxylase |
| contig_28439_5 | 0.0127 | 0.023 | 0.0275 | 0.1753 | 0 | 1 | 0.1342 | 3-isopropylmalate dehydrogenase |
| contig_24941_5 | 0.0128 | 0.023 | 0.0275 | 0.1753 | 0 | 1 | 0.1342 | triosephosphate isomerase |

| | | | | | | | | |
|---|---|---|---|---|---|---|---|---|
| contig_25119_2 | 0.0128 | 0.0236 | 0.028 | 0.1753 | 0 | 1 | 0.1342 | 3-phosphoshikimate 1-carboxyvinyltransferase |
| contig_21794_2 | 0.0133 | 0.0242 | 0.0286 | 0.1799 | 0 | 1 | 0.1378 | fucoxanthin chlorophyll a/c protein deviant |
| contig_19964_6 | 0.0137 | 0.0242 | 0.0289 | 0.1831 | 0 | 1 | 0.1402 | predicted protein |
| contig_27832_2 | 0.0138 | 0.0247 | 0.0299 | 0.184 | 0 | 1 | 0.1409 | ---NA--- |
| contig_41228_1 | 0.0142 | 0.0247 | 0.0299 | 0.184 | 0 | 1 | 0.1409 | light-harvesting protein |
| contig_5982_4 | 0.0143 | 0.0255 | 0.0301 | 0.184 | 0 | 1 | 0.1409 | abnormal spindle-like microcephaly-associated protein homolog |
| contig_29386_5 | 0.0144 | 0.0255 | 0.0306 | 0.184 | 0 | 1 | 0.1409 | predicted protein |
| contig_38925_2 | 0.0144 | 0.0261 | 0.0311 | 0.184 | 0 | 1 | 0.1409 | ---NA--- |
| contig_13757_6 | 0.0144 | 0.0267 | 0.0317 | 0.184 | 0 | 1 | 0.1409 | predicted protein |
| contig_36104_5 | 0.0147 | 0.0273 | 0.0317 | 0.1853 | 0 | 1 | 0.1419 | 40S ribosomal protein S2 |
| contig_9376_2 | 0.0149 | 0.0275 | 0.0318 | 0.1862 | 0 | 1 | 0.1426 | predicted protein |
| contig_36968_1 | 0.015 | 0.0277 | 0.0321 | 0.1862 | 0 | 1 | 0.1426 | 60S ribosomal protein L38 |
| contig_25214_5 | 0.0153 | 0.0288 | 0.0325 | 0.1889 | 0 | 1 | 0.1446 | predicted protein |
| contig_10388_3 | 0.0156 | 0.0289 | 0.0327 | 0.1903 | 0 | 1 | 0.1457 | light-harvesting protein |
| contig_27675_6 | 0.017 | 0.0301 | 0.0329 | 0.2037 | 0 | 1 | 0.156 | inosine-5'-monophosphate dehydrogenase |
| contig_20846_4 | 0.017 | 0.0306 | 0.0334 | 0.2037 | 0 | 1 | 0.156 | fructose-bisphosphate aldolase |
| contig_20218_5 | 0.0171 | 0.0308 | 0.0335 | 0.2037 | 0 | 1 | 0.156 | 40S ribosomal protein S26 |
| contig_23099_5 | 0.0172 | 0.0316 | 0.0345 | 0.2037 | 0 | 1 | 0.156 | NAD(P)-dependent alcohol dehydrogenase |
| contig_13893_5 | 0.0174 | 0.0317 | 0.035 | 0.2037 | 0 | 1 | 0.156 | NADH dehydrogenase subunit 11 (mitochondrion) |
| contig_7406_5 | 0.0174 | 0.032 | 0.0372 | 0.2037 | 0 | 1 | 0.156 | predicted protein |
| contig_20747_6 | 0.0178 | 0.032 | 0.0375 | 0.2047 | 0 | 1 | 0.1568 | translation initiation factor 4E |
| contig_4611_6 | 0.0179 | 0.0323 | 0.0377 | 0.2047 | 0 | 1 | 0.1568 | long chain acyl-coa synthetase |
| contig_35315_5 | 0.0179 | 0.0327 | 0.0384 | 0.2047 | 0 | 1 | 0.1568 | ---NA--- |
| contig_39943_5 | 0.0181 | 0.0327 | 0.0391 | 0.2049 | 0 | 1 | 0.1569 | mitochondrial alternative oxidase |
| contig_27172_4 | 0.0185 | 0.0327 | 0.0391 | 0.2075 | 0 | 1 | 0.1589 | peptidyl-prolyl cis-trans isomerase |
| contig_40448_3 | 0.0187 | 0.0329 | 0.0415 | 0.2083 | 0 | 1 | 0.1595 | RNA-binding transcriptional accessory protein |
| contig_22474_6 | 0.019 | 0.0334 | 0.0422 | 0.2095 | 0 | 1 | 0.1604 | succinyl-CoA ligase beta-chain mitochondrial precursor |

| | | | | | | | | |
|---|---|---|---|---|---|---|---|---|
| contig_9049_5 | 0.019 | 0.0335 | 0.0438 | 0.2095 | 0 | 1 | 0.1604 | enolase |
| contig_29429_4 | 0.0192 | 0.035 | 0.0459 | 0.2097 | 0 | 1 | 0.1606 | predicted protein |
| contig_26303_6 | 0.0195 | 0.0372 | 0.0469 | 0.2099 | 0 | 1 | 0.1607 | DEAD-domain-containing protein |
| contig_28019_5 | 0.0196 | 0.0375 | 0.0471 | 0.2099 | 0 | 1 | 0.1607 | NAD+ synthetase |
| contig_29346_4 | 0.0196 | 0.0377 | 0.0471 | 0.2099 | 0 | 1 | 0.1607 | homogentisate solanesyltransferase |
| contig_28140_1 | 0.02 | 0.0384 | 0.0477 | 0.2113 | 0 | 1 | 0.1618 | citrate (Si)-synthase eukaryotic |
| contig_17096_5 | 0.0202 | 0.0391 | 0.0486 | 0.2113 | 0 | 1 | 0.1618 | predicted protein |
| contig_18343_1 | 0.0203 | 0.0392 | 0.0495 | 0.2113 | 0 | 1 | 0.1618 | glutamine synthetase |
| contig_24237_6 | 0.0203 | 0.0415 | 0.0496 | 0.2113 | 0 | 1 | 0.1618 | predicted protein |
| contig_23045_4 | 0.0207 | 0.0422 | 0.0497 | 0.2132 | 0 | 1 | 0.1632 | amino acid/polyamine transporter |
| contig_14441_4 | 0.0208 | 0.0459 | 0.0503 | 0.2132 | 0 | 1 | 0.1632 | predicted protein |
| contig_27387_6 | 0.0217 | 0.0471 | 0.0505 | 0.2183 | 0 | 1 | 0.1672 | GTP-binding protein YPTC1 variant |
| contig_19117_2 | 0.0217 | 0.0471 | 0.0505 | 0.2183 | 0 | 1 | 0.1672 | 60S ribosomal protein L21-A |
| contig_35026_4 | 0.0218 | 0.0475 | 0.0515 | 0.2183 | 0 | 1 | 0.1672 | glyceraldehyde 3-phosphate dehydrogenase |
| contig_38768_2 | 0.0218 | 0.0481 | 0.0516 | 0.2183 | 0 | 1 | 0.1672 | ---NA--- |
| contig_30564_1 | 0.0222 | 0.0491 | 0.0517 | 0.2207 | 0 | 1 | 0.169 | predicted protein |
| contig_16024_6 | 0.0229 | 0.0495 | 0.052 | 0.223 | 0 | 1 | 0.1707 | Chaperonin GroL |
| contig_5781_3 | 0.0229 | 0.0497 | 0.0521 | 0.223 | 0 | 1 | 0.1707 | acetyl-coenzyme a synthetase |
| contig_40308_5 | 0.023 | 0.0497 | 0.0522 | 0.223 | 0 | 1 | 0.1707 | 40S ribosomal protein S6 |
| contig_34173_4 | 0.023 | 0.0503 | 0.0526 | 0.223 | 0 | 1 | 0.1707 | succinate dehydrogenase flavoprotein subunit |
| contig_10169_3 | 0.0235 | 0.0505 | 0.0527 | 0.2257 | 0 | 1 | 0.1728 | NADH-cytochrome b5 reductase 3-like |
| contig_30881_4 | 0.0236 | 0.0506 | 0.053 | 0.2257 | 0 | 1 | 0.1728 | pyruvate carboxylase |
| contig_24459_5 | 0.0239 | 0.0515 | 0.0535 | 0.2274 | 0 | 1 | 0.1741 | fact complex subunit ssrp1 |
| contig_9076_4 | 0.0242 | 0.0516 | 0.0542 | 0.2275 | 0 | 1 | 0.1742 | 60S ribosomal protein L24 putative |
| contig_35161_4 | 0.0242 | 0.0517 | 0.0543 | 0.2275 | 0 | 1 | 0.1742 | RNA polymerase beta' subunit (chloroplast) |
| contig_6179_6 | 0.0246 | 0.052 | 0.0546 | 0.2288 | 0 | 1 | 0.1752 | Long-chain-fatty-acid--CoA ligase putative |
| contig_17187_5 | 0.0247 | 0.0521 | 0.0546 | 0.2288 | 0 | 1 | 0.1752 | small ubiquitin-like modifier |

| | | | | | | | | |
|---|---|---|---|---|---|---|---|---|
| contig_6151_5 | 0.025 | 0.0522 | 0.0548 | 0.2301 | 0 | 1 | 0.1762 | CD209 antigen-like protein 2 isoform X1 |
| contig_9342_6 | 0.0255 | 0.0526 | 0.0549 | 0.2319 | 0 | 1 | 0.1776 | predicted protein |
| contig_29073_1 | 0.0255 | 0.0527 | 0.0552 | 0.2319 | 0 | 1 | 0.1776 | 26S proteasome non-ATPase regulatory subunit 6 |
| contig_30298_4 | 0.0258 | 0.053 | 0.0574 | 0.2338 | 0 | 1 | 0.179 | predicted protein |
| contig_21617_6 | 0.0261 | 0.0535 | 0.0583 | 0.2345 | 0 | 1 | 0.1795 | predicted protein |
| contig_37899_2 | 0.0264 | 0.0542 | 0.0587 | 0.2358 | 0 | 1 | 0.1806 | ribosomal protein L2 (chloroplast) |
| contig_28913_5 | 0.0267 | 0.0545 | 0.0603 | 0.2374 | 0 | 1 | 0.1818 | predicted protein |
| contig_34979_6 | 0.0271 | 0.0546 | 0.0606 | 0.2385 | 0 | 1 | 0.1827 | ubiquitin-like protein |
| contig_27112_5 | 0.0273 | 0.0546 | 0.0608 | 0.2385 | 0 | 1 | 0.1827 | calreticulin |
| contig_11343_2 | 0.0275 | 0.0548 | 0.0618 | 0.2385 | 0 | 1 | 0.1827 | probable aspartyl aminopeptidase |
| contig_28970_6 | 0.0275 | 0.0549 | 0.0619 | 0.2385 | 0 | 1 | 0.1827 | peptide methionine sulfoxide reductase |
| contig_38458_3 | 0.0277 | 0.0552 | 0.0656 | 0.2393 | 0 | 1 | 0.1832 | Proteasome subunit alpha/beta |
| contig_5469_5 | 0.028 | 0.0574 | 0.0657 | 0.2401 | 0 | 1 | 0.1839 | ribosomal protein S2 (chloroplast) |
| contig_34549_1 | 0.0285 | 0.0585 | 0.0681 | 0.2429 | 0 | 1 | 0.186 | pirin family protein |
| contig_23548_3 | 0.0286 | 0.0587 | 0.0685 | 0.2429 | 0 | 1 | 0.186 | ATP synthase subunit beta mitochondrial |
| contig_30343_1 | 0.0288 | 0.0604 | 0.0695 | 0.2429 | 0 | 1 | 0.186 | fucoxanthin chl a/c light-harvesting protein |
| contig_9627_5 | 0.0289 | 0.0606 | 0.0697 | 0.2429 | 0 | 1 | 0.186 | 26S proteasome non-ATPase regulatory subunit 14 |
| contig_9330_5 | 0.0298 | 0.0611 | 0.0708 | 0.2484 | 0 | 1 | 0.1902 | short-chain alcohol dehydrogenase with NAD or NADP as acceptor |
| contig_22713_2 | 0.0299 | 0.0618 | 0.0725 | 0.2484 | 0 | 1 | 0.1902 | predicted protein |
| contig_17799_3 | 0.0301 | 0.0619 | 0.0731 | 0.2484 | 0 | 1 | 0.1902 | light-harvesting protein |
| contig_32560_1 | 0.0305 | 0.0657 | 0.0731 | 0.2504 | 0 | 1 | 0.1917 | predicted protein |
| contig_10341_1 | 0.0306 | 0.0681 | 0.0834 | 0.2504 | 0 | 1 | 0.1917 | predicted protein |
| contig_25429_4 | 0.0308 | 0.0685 | 0.0897 | 0.2505 | 0 | 1 | 0.1918 | V-type proton ATPase proteolipid subunit |
| contig_33628_1 | 0.0311 | 0.0695 | 0.0897 | 0.2514 | 0 | 1 | 0.1925 | glycine-rich domain-containing protein 2-like |
| contig_33602_5 | 0.0316 | 0.0697 | 0.0904 | 0.2514 | 0 | 1 | 0.1925 | FAD-binding domain-containing protein |
| contig_39484_1 | 0.0317 | 0.0708 | 0.0934 | 0.2514 | 0 | 1 | 0.1925 | probable aspartyl aminopeptidase |
| contig_23620_5 | 0.0317 | 0.0726 | 0.0936 | 0.2514 | 0 | 1 | 0.1925 | nitrate transporter |

| | | | | | | | | |
|---|---|---|---|---|---|---|---|---|
| contig_35362_1 | 0.0318 | 0.0726 | 0.0936 | 0.2514 | 0 | 1 | 0.1925 | nitrate transporter |
| contig_28483_6 | 0.032 | 0.0838 | 0.0969 | 0.2514 | 0 | 1 | 0.1925 | predicted protein |
| contig_19373_4 | 0.0321 | 0.0838 | 0.0969 | 0.2514 | 0 | 1 | 0.1925 | fucoxanthin chlorophyll a/c-binding protein precursor |
| contig_10672_3 | 0.0323 | 0.0897 | 0.0981 | 0.2514 | 0 | 1 | 0.1925 | ATP synthase subunit |
| contig_15455_3 | 0.0325 | 0.0897 | 0.0981 | 0.2514 | 0 | 1 | 0.1925 | 1 2-dihydroxy-3-keto-5-methylthiopentene dioxygenase |
| contig_1045_6 | 0.0327 | 0.0936 | 0.0983 | 0.2514 | 0 | 1 | 0.1925 | iron-sulfur cluster formation ABC transporter (chloroplast) |
| contig_28430_6 | 0.0327 | 0.0969 | 0.1002 | 0.2514 | 0 | 1 | 0.1925 | predicted protein |
| contig_23369_5 | 0.0329 | 0.0969 | 0.1007 | 0.2514 | 0 | 1 | 0.1925 | predicted protein |
| contig_33444_6 | 0.0333 | 0.0975 | 0.1055 | 0.252 | 0 | 1 | 0.1929 | predicted protein |
| contig_10415_5 | 0.0334 | 0.0981 | 0.1057 | 0.252 | 0 | 1 | 0.1929 | DNA-binding protein 42 kDa |
| contig_33542_2 | 0.0335 | 0.0982 | 0.114 | 0.252 | 0 | 1 | 0.1929 | nucleoside-diphosphate kinase |
| contig_37779_2 | 0.0344 | 0.0983 | 0.1145 | 0.2571 | 0 | 1 | 0.1969 | protein fucoxanthin chlorophyll a/c protein |
| contig_28392_1 | 0.0345 | 0.1002 | 0.1153 | 0.2571 | 0 | 1 | 0.1969 | predicted protein |
| contig_23843_1 | 0.0349 | 0.1007 | 0.1157 | 0.2585 | 0 | 1 | 0.198 | methyltransferase domain-containing protein |
| contig_38780_2 | 0.035 | 0.1055 | 0.118 | 0.2585 | 1E-04 | 1 | 0.198 | carbamoyl-phosphate synthase large subunit |
| contig_11658_2 | 0.0358 | 0.114 | 0.118 | 0.2632 | 1E-04 | 1 | 0.2016 | pyrimidine/purine nucleoside phosphorylase |
| contig_20200_5 | 0.0367 | 0.1141 | 0.1182 | 0.2687 | 1E-04 | 1 | 0.2058 | 60s ribosomal protein l18a |
| contig_14595_3 | 0.037 | 0.1153 | 0.1191 | 0.2691 | 1E-04 | 1 | 0.2061 | predicted protein |
| contig_6178_4 | 0.0372 | 0.1157 | 0.1201 | 0.2691 | 1E-04 | 1 | 0.2061 | rubisco expression protein (chloroplast) |
| contig_11292_5 | 0.0373 | 0.1167 | 0.1205 | 0.2691 | 2E-04 | 1 | 0.2061 | heat shock protein belonging to the HSP70 family |
| contig_25208_3 | 0.0375 | 0.118 | 0.1205 | 0.2693 | 2E-04 | 1 | 0.2062 | predicted protein |
| contig_37776_1 | 0.0377 | 0.1182 | 0.1207 | 0.2693 | 2E-04 | 1 | 0.2063 | DNA polymerase |
| contig_28781_6 | 0.0382 | 0.1191 | 0.1217 | 0.2713 | 3E-04 | 1 | 0.2078 | heat shock protein 90 |
| contig_13084_2 | 0.0384 | 0.1203 | 0.1254 | 0.2721 | 4E-04 | 1 | 0.2083 | 60S ribosomal protein L14 putative |
| contig_39819_3 | 0.0387 | 0.1205 | 0.1257 | 0.2724 | 5E-04 | 1 | 0.2086 | predicted protein |
| contig_32573_1 | 0.0391 | 0.1207 | 0.1372 | 0.2724 | 6E-04 | 1 | 0.2086 | ATP synthase CF1 alpha subunit (chloroplast) |
| contig_28103_5 | 0.0391 | 0.1211 | 0.1404 | 0.2724 | 7E-04 | 1 | 0.2086 | magnesium chelatase subunit H |

| | | | | | | | | |
|---|---|---|---|---|---|---|---|---|
| contig_9641_3 | 0.0392 | 0.1226 | 0.145 | 0.2724 | 9E-04 | 1 | 0.2086 | predicted protein |
| contig_27839_5 | 0.0412 | 0.1255 | 0.1451 | 0.2833 | 0.001 | 1 | 0.2169 | predicted protein |
| contig_36895_3 | 0.0412 | 0.1371 | 0.147 | 0.2833 | 0.0013 | 1 | 0.2169 | phenylalanine-tRNA ligase beta subunit (chloroplast) |
| contig_17285_6 | 0.0414 | 0.1372 | 0.1476 | 0.2833 | 0.0015 | 1 | 0.2169 | 40S ribosomal protein S27-4 putative |
| contig_33723_4 | 0.0415 | 0.145 | 0.1491 | 0.2833 | 0.0018 | 1 | 0.2169 | predicted protein |
| contig_22850_1 | 0.0421 | 0.1451 | 0.1492 | 0.2857 | 0.0022 | 1 | 0.2188 | nitrate transporter |
| contig_15866_5 | 0.0422 | 0.147 | 0.1498 | 0.2857 | 0.0026 | 1 | 0.2188 | hypothetical protein THAOC_02924 |
| contig_37452_3 | 0.0428 | 0.1476 | 0.1506 | 0.2883 | 0.0031 | 1 | 0.2208 | ADP-ribosylation factor |
| contig_11304_3 | 0.0438 | 0.1492 | 0.1507 | 0.2926 | 0.0036 | 1 | 0.224 | predicted protein |
| contig_24282_5 | 0.0438 | 0.1498 | 0.1509 | 0.2926 | 0.0043 | 1 | 0.224 | predicted protein |
| contig_10198_5 | 0.0451 | 0.1498 | 0.1537 | 0.2992 | 0.005 | 1 | 0.2291 | predicted protein |
| contig_38999_3 | 0.0452 | 0.1507 | 0.1545 | 0.2992 | 0.0058 | 1 | 0.2291 | predicted protein |
| contig_17918_5 | 0.0459 | 0.1507 | 0.1545 | 0.3011 | 0.0068 | 1 | 0.2306 | glutamate-1-semialdehyde 2 1-aminomutase |
| contig_16936_1 | 0.0459 | 0.1509 | 0.1551 | 0.3011 | 0.0079 | 1 | 0.2306 | H/ACA ribonucleoprotein complex subunit 4 |
| contig_15496_6 | 0.0465 | 0.1509 | 0.1551 | 0.3028 | 0.0092 | 1 | 0.2319 | catalase |
| contig_22686_4 | 0.0469 | 0.1548 | 0.1589 | 0.3028 | 0.0106 | 1 | 0.2319 | predicted protein |
| contig_22629_4 | 0.0469 | 0.1548 | 0.1589 | 0.3028 | 0.0122 | 1 | 0.2319 | acetyl-coa carboxylase |
| contig_34082_5 | 0.0471 | 0.1569 | 0.1653 | 0.3028 | 0.0141 | 1 | 0.2319 | predicted protein |
| contig_22636_2 | 0.0471 | 0.1569 | 0.1653 | 0.3028 | 0.0161 | 1 | 0.2319 | 60s acidic ribosomal protein p0 |
| contig_26583_1 | 0.0475 | 0.1589 | 0.166 | 0.3038 | 0.0185 | 1 | 0.2326 | predicted protein |
| contig_5881_4 | 0.0477 | 0.1589 | 0.166 | 0.3038 | 0.021 | 1 | 0.2326 | elongation factor Tu (chloroplast) |
| contig_39157_3 | 0.0481 | 0.1655 | 0.1685 | 0.305 | 0.0239 | 1 | 0.2336 | predicted protein |
| contig_21789_5 | 0.0486 | 0.1655 | 0.1685 | 0.306 | 0.0271 | 1 | 0.2343 | Mitochondrial Carrier (MC) Family |
| contig_24751_3 | 0.0491 | 0.1682 | 0.1691 | 0.306 | 0.0307 | 1 | 0.2343 | predicted protein |
| contig_19427_5 | 0.0493 | 0.1682 | 0.1691 | 0.306 | 0.0346 | 1 | 0.2343 | 60S ribosomal protein L18a |
| contig_27862_5 | 0.0495 | 0.1687 | 0.1709 | 0.306 | 0.0389 | 1 | 0.2343 | predicted protein |
| contig_6528_1 | 0.0496 | 0.1687 | 0.1709 | 0.306 | 0.0436 | 1 | 0.2343 | 40S ribosomal protein S19-3 |

| | | | | | | | | |
|---|---|---|---|---|---|---|---|---|
| contig_5747_6 | 0.0497 | 0.1703 | 0.1711 | 0.306 | 0.0488 | 1 | 0.2343 | photosystem I assembly protein Ycf4 (chloroplast) |
| contig_39464_1 | 0.0502 | 0.1709 | 0.1765 | 0.306 | 0.0545 | 1 | 0.2343 | histone deacetylase 1 |
| contig_14932_3 | 0.0503 | 0.171 | 0.1766 | 0.306 | 0.0606 | 1 | 0.2343 | predicted protein |
| contig_39064_3 | 0.0505 | 0.171 | 0.1802 | 0.306 | 0.0673 | 1 | 0.2343 | Glycerophosphoryl diester phosphodiesterase |
| contig_5881_6 | 0.0505 | 0.1765 | 0.1808 | 0.306 | 0.0746 | 1 | 0.2343 | elongation factor Tu (chloroplast) |
| contig_11299_6 | 0.0506 | 0.1766 | 0.1815 | 0.306 | 0.0824 | 1 | 0.2343 | predicted protein |
| contig_24360_5 | 0.0512 | 0.1802 | 0.1816 | 0.306 | 0.0909 | 1 | 0.2343 | splicing factor 3B subunit 3-like |
| contig_5607_4 | 0.0515 | 0.1808 | 0.1913 | 0.306 | 0.0999 | 1 | 0.2343 | protease SohB |
| contig_27843_1 | 0.0516 | 0.1815 | 0.2144 | 0.306 | 0.1096 | 1 | 0.2343 | Fe-S cluster assembly protein SufD |
| contig_29278_3 | 0.0517 | 0.1816 | 0.215 | 0.306 | 0.1199 | 1 | 0.2343 | predicted protein |
| contig_38793_1 | 0.0519 | 0.1943 | 0.2188 | 0.306 | 0.131 | 1 | 0.2343 | predicted protein |
| contig_23910_2 | 0.052 | 0.215 | 0.2232 | 0.306 | 0.1427 | 1 | 0.2343 | ARM repeat-containing protein |
| contig_9471_3 | 0.0521 | 0.2188 | 0.2237 | 0.306 | 0.155 | 1 | 0.2343 | photosystem II CP43 chlorophyll apoprotein (chloroplast) |
| contig_11877_6 | 0.0522 | 0.2234 | 0.2972 | 0.306 | 0.1681 | 1 | 0.2343 | 30S ribosomal protein S5 |
| contig_10866_1 | 0.0525 | 0.2237 | 0.2973 | 0.306 | 0.1819 | 1 | 0.2343 | predicted protein |
| contig_30410_4 | 0.0526 | 1 | 1 | 0.306 | 0.1964 | 1 | 0.2343 | predicted protein |
| contig_17520_3 | 0.0527 | 1 | 1 | 0.306 | 0.2115 | 1 | 0.2343 | methyltransferase FkbM family (macronuclear) |
| contig_9468_6 | 0.053 | 1 | 1 | 0.306 | 0.2274 | 1 | 0.2343 | anaphase-promoting complex subunit 1 |
| contig_22808_5 | 0.0532 | 1 | 1 | 0.306 | 0.2438 | 1 | 0.2343 | predicted protein |
| contig_15155_6 | 0.0535 | 1 | 1 | 0.306 | 0.2609 | 1 | 0.2343 | predicted protein |
| contig_15547_4 | 0.054 | 1 | 1 | 0.306 | 0.2785 | 1 | 0.2343 | hypothetical protein FRACYDRAFT_234640 |
| contig_34750_5 | 0.0542 | 1 | 1 | 0.306 | 0.2967 | 1 | 0.2343 | thioredoxin-disulfide reductase |
| contig_29365_1 | 0.0543 | 1 | 1 | 0.306 | 0.3153 | 1 | 0.2343 | predicted protein |
| contig_18476_4 | 0.0545 | 1 | 1 | 0.306 | 0.3345 | 1 | 0.2343 | transmembrane 9 superfamily member 4 |
| contig_9047_5 | 0.0546 | 1 | 1 | 0.306 | 0.3541 | 1 | 0.2343 | predicted protein |
| contig_27341_1 | 0.0546 | 1 | 1 | 0.306 | 0.3741 | 1 | 0.2343 | protein patched homolog 2 |
| contig_26568_4 | 0.0548 | 1 | 1 | 0.306 | 0.3945 | 1 | 0.2343 | atpase family aaa domain-containing protein 3 |

| | | | | | | | | |
|---|---|---|---|---|---|---|---|---|
| contig_24581_4 | 0.0549 | 1 | 1 | 0.306 | 0.4152 | 1 | 0.2343 | predicted protein |
| contig_30461_3 | 0.0551 | 1 | 1 | 0.306 | 0.4361 | 1 | 0.2343 | 6 7-dimethyl-8-ribityllumazine synthase |
| contig_30318_6 | 0.0552 | 1 | 1 | 0.306 | 0.4572 | 1 | 0.2343 | pyruvate dehydrogenase complex dihydrolipoamide acetyltransferase |
| contig_39705_1 | 0.0567 | 1 | 1 | 0.3125 | 0.4785 | 1 | 0.2393 | inorganic pyrophosphatase |
| contig_38571_1 | 0.0568 | 1 | 1 | 0.3125 | 0.4995 | 1 | 0.2393 | photosystem I subunit III (chloroplast) |
| contig_23912_3 | 0.0573 | 1 | 1 | 0.3133 | 0.5206 | 1 | 0.2399 | predicted protein |
| contig_40017_2 | 0.0574 | 1 | 1 | 0.3133 | 0.5416 | 1 | 0.2399 | predicted protein |
| contig_38023_2 | 0.0581 | 1 | 1 | 0.3159 | 0.5625 | 1 | 0.2419 | predicted protein |
| contig_28094_2 | 0.0583 | 1 | 1 | 0.3159 | 0.583 | 1 | 0.2419 | agmatinase |
| contig_9651_3 | 0.0585 | 1 | 1 | 0.3159 | 0.6034 | 1 | 0.2419 | ribosomal protein RPL17 |
| contig_8365_3 | 0.0587 | 1 | 1 | 0.3161 | 0.6235 | 1 | 0.2421 | pyruvate dehydrogenase |
| contig_35278_1 | 0.059 | 1 | 1 | 0.3168 | 0.6432 | 1 | 0.2426 | light harvesting protein |
| contig_16833_4 | 0.0602 | 1 | 1 | 0.3207 | 0.6626 | 1 | 0.2456 | 60S ribosomal protein |
| contig_32502_6 | 0.0603 | 1 | 1 | 0.3207 | 0.6814 | 1 | 0.2456 | predicted protein |
| contig_38361_6 | 0.0604 | 1 | 1 | 0.3207 | 0.6997 | 1 | 0.2456 | 60S ribosomal protein L7a |
| contig_33184_2 | 0.0606 | 1 | 1 | 0.3208 | 0.7175 | 1 | 0.2456 | predicted protein |
| contig_16671_2 | 0.0608 | 1 | 1 | 0.3208 | 0.7349 | 1 | 0.2456 | ribosomal protein L21 (chloroplast) |
| contig_8116_6 | 0.0611 | 1 | 1 | 0.3211 | 0.7516 | 1 | 0.2459 | 40S ribosomal protein S11 |
| contig_29950_5 | 0.0617 | 1 | 1 | 0.3218 | 0.7678 | 1 | 0.2464 | Fe-S cluster assembly protein SufD |
| contig_6296_5 | 0.0618 | 1 | 1 | 0.3218 | 0.7833 | 1 | 0.2464 | predicted protein |
| contig_37814_3 | 0.0619 | 1 | 1 | 0.3218 | 0.7982 | 1 | 0.2464 | ribosomal protein S3 (chloroplast) |
| contig_5526_6 | 0.0633 | 1 | 1 | 0.328 | 0.8125 | 1 | 0.2512 | pol protein |
| contig_24410_6 | 0.0637 | 1 | 1 | 0.3281 | 0.826 | 1 | 0.2512 | sodium/potassium/calcium exchanger 2-like |
| contig_28207_5 | 0.0637 | 1 | 1 | 0.3281 | 0.8389 | 1 | 0.2512 | mannose-P-dolichol utilization defect 1 protein |
| contig_22513_4 | 0.0649 | 1 | 1 | 0.3329 | 0.8511 | 1 | 0.2549 | predicted protein |
| contig_2955_5 | 0.0653 | 1 | 1 | 0.3338 | 0.8626 | 1 | 0.2556 | NADH dehydrogenase (ubiquinone) flavoprotein 2 |
| contig_29535_5 | 0.0656 | 1 | 1 | 0.3338 | 0.8735 | 1 | 0.2556 | predicted protein |

| | | | | | | | | |
|---|---|---|---|---|---|---|---|---|
| contig_30367_5 | 0.0657 | 1 | 1 | 0.3338 | 0.8837 | 1 | 0.2556 | protein fucoxanthin chlorophyll a/c protein |
| contig_19340_6 | 0.0669 | 1 | 1 | 0.3386 | 0.8933 | 1 | 0.2593 | acetyl-coa carboxylase |
| contig_25871_6 | 0.0672 | 1 | 1 | 0.3393 | 0.9023 | 1 | 0.2598 | RNA-binding domain-containing protein |
| contig_6700_4 | 0.0677 | 1 | 1 | 0.3401 | 0.9107 | 1 | 0.2605 | agmatinase |
| contig_32601_3 | 0.068 | 1 | 1 | 0.3401 | 0.9186 | 1 | 0.2605 | predicted protein |
| contig_29419_6 | 0.0681 | 1 | 1 | 0.3401 | 0.9259 | 1 | 0.2605 | leucyl-tRNA synthetase |
| contig_22730_1 | 0.0685 | 1 | 1 | 0.3402 | 0.9327 | 1 | 0.2605 | predicted protein |
| contig_25490_5 | 0.0685 | 1 | 1 | 0.3402 | 0.9389 | 1 | 0.2605 | ribonucleoside-diphosphate reductase large subunit |
| contig_37814_2 | 0.0692 | 1 | 1 | 0.3425 | 0.9448 | 1 | 0.2623 | ribosomal protein S3 (chloroplast) |
| contig_22140_2 | 0.0695 | 1 | 1 | 0.3425 | 0.9501 | 1 | 0.2623 | predicted protein |
| contig_38581_3 | 0.0697 | 1 | 1 | 0.3425 | 0.9551 | 1 | 0.2623 | predicted protein |
| contig_16604_4 | 0.0702 | 1 | 1 | 0.3442 | 0.9596 | 1 | 0.2636 | myo-inositol 2-dehydrogenase putative |
| contig_28093_5 | 0.0707 | 1 | 1 | 0.3449 | 0.9638 | 1 | 0.2641 | protein dispatched homolog 1-like |
| contig_40230_1 | 0.0708 | 1 | 1 | 0.3449 | 0.9675 | 1 | 0.2641 | predicted protein |
| contig_27467_4 | 0.072 | 1 | 1 | 0.349 | 0.971 | 1 | 0.2673 | histidinol-phosphate aminotransferase |
| contig_24254_6 | 0.0724 | 1 | 1 | 0.349 | 0.9741 | 1 | 0.2673 | predicted protein |
| contig_28046_4 | 0.0725 | 1 | 1 | 0.349 | 0.9769 | 1 | 0.2673 | light harvesting protein |
| contig_5519_5 | 0.0726 | 1 | 1 | 0.349 | 0.9795 | 1 | 0.2673 | protein tesmin/TSO1-like CXC 5 |
| contig_24908_5 | 0.0731 | 1 | 1 | 0.3493 | 0.9818 | 1 | 0.2675 | transitional endoplasmic reticulum ATPase TER94 |
| contig_19000_1 | 0.0731 | 1 | 1 | 0.3493 | 0.9839 | 1 | 0.2675 | predicted protein |
| contig_5826_3 | 0.0752 | 1 | 1 | 0.3584 | 0.9858 | 1 | 0.2745 | 60S ribosomal protein L2 |
| contig_16274_4 | 0.0759 | 1 | 1 | 0.3597 | 0.9875 | 1 | 0.2755 | predicted protein |
| contig_15835_5 | 0.076 | 1 | 1 | 0.3597 | 0.989 | 1 | 0.2755 | ---NA--- |
| contig_22654_5 | 0.0769 | 1 | 1 | 0.3627 | 0.9903 | 1 | 0.2777 | predicted protein |
| contig_27901_2 | 0.0786 | 1 | 1 | 0.3698 | 0.9915 | 1 | 0.2832 | methionyl-trna formyltransferase |
| contig_21839_5 | 0.0794 | 1 | 1 | 0.3722 | 0.9925 | 1 | 0.285 | 1-deoxy-d-xylulose-5-phosphate reductoisomerase |
| contig_30052_6 | 0.0796 | 1 | 1 | 0.3723 | 0.9934 | 1 | 0.2851 | predicted protein |

| | | | | | | | | |
|---|---|---|---|---|---|---|---|---|
| contig_27872_3 | 0.0809 | 1 | 1 | 0.3758 | 0.9943 | 1 | 0.2878 | predicted protein |
| contig_24223_6 | 0.081 | 1 | 1 | 0.3758 | 0.995 | 1 | 0.2878 | predicted protein |
| contig_10561_5 | 0.0815 | 1 | 1 | 0.3758 | 0.9956 | 1 | 0.2878 | 6-phosphogluconate dehydrogenase |
| contig_25695_5 | 0.0817 | 1 | 1 | 0.3758 | 0.9962 | 1 | 0.2878 | 40S ribosomal protein S7 |
| contig_17829_4 | 0.0818 | 1 | 1 | 0.3758 | 0.9967 | 1 | 0.2878 | predicted protein |
| contig_32146_2 | 0.0819 | 1 | 1 | 0.3758 | 0.9971 | 1 | 0.2878 | silicon transporter |
| contig_23501_5 | 0.0825 | 1 | 1 | 0.3779 | 0.9975 | 1 | 0.2894 | bile acid:sodium symporter |
| contig_17958_6 | 0.0828 | 1 | 1 | 0.3779 | 0.9979 | 1 | 0.2894 | luminal-binding protein |
| contig_30131_2 | 0.0834 | 1 | 1 | 0.3793 | 0.9982 | 1 | 0.2905 | dihydrolipoyl dehydrogenase |
| contig_15603_6 | 0.0838 | 1 | 1 | 0.3804 | 0.9984 | 1 | 0.2913 | predicted protein |
| contig_41157_2 | 0.0853 | 1 | 1 | 0.3857 | 0.9986 | 1 | 0.2953 | predicted protein |
| contig_25682_4 | 0.087 | 1 | 1 | 0.392 | 0.9988 | 1 | 0.3002 | Immunoglobulin G-binding protein A |
| contig_28951_4 | 0.0872 | 1 | 1 | 0.392 | 0.999 | 1 | 0.3002 | predicted protein |
| contig_37607_6 | 0.0876 | 1 | 1 | 0.392 | 0.9991 | 1 | 0.3002 | 2-isopropylmalate synthase |
| contig_40284_6 | 0.0879 | 1 | 1 | 0.392 | 0.9993 | 1 | 0.3002 | large subunit ribosomal protein L13e |
| contig_28707_5 | 0.0879 | 1 | 1 | 0.392 | 0.9994 | 1 | 0.3002 | V-type H+-transporting ATPase subunit a |
| contig_16230_2 | 0.0884 | 1 | 1 | 0.3931 | 0.9995 | 1 | 0.301 | phosphoglycerate mutase |
| contig_32986_4 | 0.0895 | 1 | 1 | 0.3942 | 0.9995 | 1 | 0.3019 | protochlorophyllide reductase magnesium chelatase subunit (chloroplast) |
| contig_17375_3 | 0.0896 | 1 | 1 | 0.3942 | 0.9996 | 1 | 0.3019 | cysteine synthase |
| contig_25500_5 | 0.0897 | 1 | 1 | 0.3942 | 0.9997 | 1 | 0.3019 | predicted protein |
| contig_27592_6 | 0.0897 | 1 | 1 | 0.3942 | 0.9997 | 1 | 0.3019 | zeta-carotene desaturase |
| contig_24874_1 | 0.0903 | 1 | 1 | 0.3947 | 0.9998 | 1 | 0.3023 | predicted protein |
| contig_13426_4 | 0.0904 | 1 | 1 | 0.3947 | 0.9998 | 1 | 0.3023 | long-chain fatty acid--CoA ligase |
| contig_6069_2 | 0.0921 | 1 | 1 | 0.4013 | 0.9998 | 1 | 0.3073 | GLNA glutamine synthase |
| contig_16087_6 | 0.0927 | 1 | 1 | 0.402 | 0.9999 | 1 | 0.3078 | predicted protein |
| contig_36394_4 | 0.0933 | 1 | 1 | 0.402 | 0.9999 | 1 | 0.3078 | Fucoxanthin-chlorophyll a-c binding protein F chloroplastic |

| | | | | | | | | |
|---|---|---|---|---|---|---|---|---|
| contig_19536_5 | 0.0933 | 1 | 1 | 0.402 | 0.9999 | 1 | 0.3078 | methionine S-adenosyl transferase |
| contig_37721_1 | 0.0934 | 1 | 1 | 0.402 | 0.9999 | 1 | 0.3078 | ---NA--- |
| contig_12101_5 | 0.0936 | 1 | 1 | 0.402 | 0.9999 | 1 | 0.3078 | ---NA--- |
| contig_17991_3 | 0.094 | 1 | 1 | 0.4024 | 0.9999 | 1 | 0.3082 | light-harvesting protein |
| contig_5591_4 | 0.0947 | 1 | 1 | 0.4044 | 1 | 1 | 0.3097 | predicted protein |
| contig_25573_4 | 0.0959 | 1 | 1 | 0.4071 | 1 | 1 | 0.3118 | predicted protein |
| contig_40093_3 | 0.0962 | 1 | 1 | 0.4071 | 1 | 1 | 0.3118 | predicted protein |
| contig_22469_6 | 0.0967 | 1 | 1 | 0.4071 | 1 | 1 | 0.3118 | predicted protein |
| contig_3111_6 | 0.0969 | 1 | 1 | 0.4071 | 1 | 1 | 0.3118 | acyl-CoA dehydrogenase |
| contig_14853_4 | 0.0969 | 1 | 1 | 0.4071 | 1 | 1 | 0.3118 | peptidyl-prolyl cis-trans isomerase FKBP19 chloroplastic |
| contig_17394_3 | 0.0974 | 1 | 1 | 0.4071 | 1 | 1 | 0.3118 | proliferating cell nuclear antigen |
| contig_30315_5 | 0.0975 | 1 | 1 | 0.4071 | 1 | 1 | 0.3118 | thioredoxin H4-1-like |
| contig_24921_6 | 0.0981 | 1 | 1 | 0.4071 | 1 | 1 | 0.3118 | nitrate transporter |
| contig_5425_1 | 0.0981 | 1 | 1 | 0.4071 | 1 | 1 | 0.3118 | sulfate adenylyltransferase |
| contig_13616_6 | 0.0982 | 1 | 1 | 0.4071 | 1 | 1 | 0.3118 | hsp70-like protein |
| contig_37905_5 | 0.0983 | 1 | 1 | 0.4071 | 1 | 1 | 0.3118 | ---NA--- |
| contig_25394_4 | 0.0987 | 1 | 1 | 0.4078 | 1 | 1 | 0.3123 | prohibitin putative |
| contig_13811_4 | 0.0995 | 1 | 1 | 0.4099 | 1 | 1 | 0.3139 | fucoxanthin chlorophyll a/c protein 8 |
| contig_23722_3 | 0.0998 | 1 | 1 | 0.4103 | 1 | 1 | 0.3142 | light harvesting protein |
| contig_9372_6 | 0.1002 | 1 | 1 | 0.4105 | 1 | 1 | 0.3144 | predicted protein |
| contig_21557_3 | 0.1007 | 1 | 1 | 0.4105 | 1 | 1 | 0.3144 | 40S ribosomal protein S2 |
| contig_41162_4 | 0.1007 | 1 | 1 | 0.4105 | 1 | 1 | 0.3144 | 60S ribosomal protein L9-like |
| contig_11991_5 | 0.104 | 1 | 1 | 0.4209 | 1 | 1 | 0.3223 | ---NA--- |
| contig_15545_5 | 0.1043 | 1 | 1 | 0.4209 | 1 | 1 | 0.3223 | Small nuclear ribonucleoprotein G |
| contig_17212_4 | 0.1044 | 1 | 1 | 0.4209 | 1 | 1 | 0.3223 | uroporphyrinogen decarboxylase |
| contig_32774_1 | 0.1048 | 1 | 1 | 0.4209 | 1 | 1 | 0.3223 | 3-deoxy-7-phosphoheptulonate synthase |
| contig_7984_6 | 0.1049 | 1 | 1 | 0.4209 | 1 | 1 | 0.3223 | ribosomal protein L1 (chloroplast) |

| | | | | | | | | |
|---|---|---|---|---|---|---|---|---|
| contig_22487_6 | 0.105 | 1 | 1 | 0.4209 | 1 | 1 | 0.3223 | predicted protein |
| contig_39493_2 | 0.1052 | 1 | 1 | 0.4209 | 1 | 1 | 0.3223 | 60s ribosomal protein |
| contig_12115_3 | 0.1055 | 1 | 1 | 0.4209 | 1 | 1 | 0.3223 | predicted protein |
| contig_24825_6 | 0.1057 | 1 | 1 | 0.4209 | 1 | 1 | 0.3223 | predicted protein |
| contig_33008_5 | 0.1067 | 1 | 1 | 0.4238 | 1 | 1 | 0.3246 | protein fucoxanthin chlorophyll a/c protein |
| contig_14326_6 | 0.1085 | 1 | 1 | 0.4297 | 1 | 1 | 0.329 | peroxiredoxin |
| contig_28947_4 | 0.1088 | 1 | 1 | 0.4297 | 1 | 1 | 0.329 | predicted protein |
| contig_23336_4 | 0.1093 | 1 | 1 | 0.4306 | 1 | 1 | 0.3297 | 1-hydroxy-2-methyl-2-(E)-butenyl-4-diphosphate synthase |
| contig_22608_4 | 0.1097 | 1 | 1 | 0.4306 | 1 | 1 | 0.3297 | predicted protein |
| contig_25969_5 | 0.1098 | 1 | 1 | 0.4306 | 1 | 1 | 0.3297 | Tic20-like protein |
| contig_30840_2 | 0.1116 | 1 | 1 | 0.4358 | 1 | 1 | 0.3337 | 60S ribosomal protein L18a |
| contig_38702_2 | 0.112 | 1 | 1 | 0.4358 | 1 | 1 | 0.3337 | predicted protein |
| contig_22654_2 | 0.112 | 1 | 1 | 0.4358 | 1 | 1 | 0.3337 | predicted protein |
| contig_6339_4 | 0.1127 | 1 | 1 | 0.4368 | 1 | 1 | 0.3345 | predicted protein |
| contig_33176_3 | 0.1128 | 1 | 1 | 0.4368 | 1 | 1 | 0.3345 | EF-hand domain-containing protein |
| contig_33573_6 | 0.1138 | 1 | 1 | 0.4382 | 1 | 1 | 0.3356 | ---NA--- |
| contig_28215_4 | 0.114 | 1 | 1 | 0.4382 | 1 | 1 | 0.3356 | light harvesting complex protein |
| contig_27789_6 | 0.1141 | 1 | 1 | 0.4382 | 1 | 1 | 0.3356 | Mitochondrial-processing peptidase subunit alpha |
| contig_24961_6 | 0.1145 | 1 | 1 | 0.4386 | 1 | 1 | 0.3358 | mitochondrial carrier |
| contig_40949_4 | 0.1151 | 1 | 1 | 0.439 | 1 | 1 | 0.3362 | U3 small nucleolar RNA-associated protein 18 homolog |
| contig_39895_1 | 0.1153 | 1 | 1 | 0.439 | 1 | 1 | 0.3362 | protein fucoxanthin chlorophyll a/c protein |
| contig_29498_6 | 0.1157 | 1 | 1 | 0.439 | 1 | 1 | 0.3362 | aldehyde dehydrogenase |
| contig_28181_3 | 0.1157 | 1 | 1 | 0.439 | 1 | 1 | 0.3362 | dihydroxy-acid dehydratase |
| contig_27237_6 | 0.1161 | 1 | 1 | 0.4394 | 1 | 1 | 0.3365 | translation initiation factor 5B |
| contig_20909_4 | 0.1167 | 1 | 1 | 0.4407 | 1 | 1 | 0.3375 | 40S ribosomal protein |
| contig_28347_6 | 0.1177 | 1 | 1 | 0.4409 | 1 | 1 | 0.3376 | DEAD-box ATP-dependent RNA helicase 8 |
| contig_25885_5 | 0.118 | 1 | 1 | 0.4409 | 1 | 1 | 0.3376 | acetyl-coenzyme a synthetase |

| | | | | | | | | |
|---|---|---|---|---|---|---|---|---|
| contig_24544_3 | 0.118 | 1 | 1 | 0.4409 | 1 | 1 | 0.3376 | predicted protein |
| contig_16670_4 | 0.1181 | 1 | 1 | 0.4409 | 1 | 1 | 0.3376 | porphobilinogen deaminase |
| contig_26248_5 | 0.1182 | 1 | 1 | 0.4409 | 1 | 1 | 0.3376 | predicted protein |
| contig_39855_1 | 0.1188 | 1 | 1 | 0.4416 | 1 | 1 | 0.3381 | predicted protein |
| contig_29269_4 | 0.1191 | 1 | 1 | 0.4416 | 1 | 1 | 0.3381 | predicted protein |
| contig_10410_5 | 0.12 | 1 | 1 | 0.4416 | 1 | 1 | 0.3381 | LL-diaminopimelate aminotransferase |
| contig_32599_2 | 0.1201 | 1 | 1 | 0.4416 | 1 | 1 | 0.3381 | ATP-dependent DNA helicase PIF1-like |
| contig_10991_6 | 0.1203 | 1 | 1 | 0.4416 | 1 | 1 | 0.3381 | ABC transporter |
| contig_32695_4 | 0.1205 | 1 | 1 | 0.4416 | 1 | 1 | 0.3381 | predicted protein |
| contig_7506_5 | 0.1205 | 1 | 1 | 0.4416 | 1 | 1 | 0.3381 | apocytochrome b (mitochondrion) |
| contig_34816_5 | 0.1207 | 1 | 1 | 0.4416 | 1 | 1 | 0.3381 | 40S ribosomal protein S12 |
| contig_34938_2 | 0.1211 | 1 | 1 | 0.442 | 1 | 1 | 0.3384 | predicted protein |
| contig_39521_4 | 0.1217 | 1 | 1 | 0.4421 | 1 | 1 | 0.3386 | predicted protein |
| contig_24913_6 | 0.1217 | 1 | 1 | 0.4421 | 1 | 1 | 0.3386 | glutamate synthase subunit alpha |
| contig_27144_6 | 0.1226 | 1 | 1 | 0.4443 | 1 | 1 | 0.3402 | predicted protein |
| contig_30812_2 | 0.1244 | 1 | 1 | 0.448 | 1 | 1 | 0.3431 | citrate synthase |
| contig_39369_3 | 0.1248 | 1 | 1 | 0.448 | 1 | 1 | 0.3431 | hypothetical protein FRACYDRAFT_269748 |
| contig_24234_3 | 0.1248 | 1 | 1 | 0.448 | 1 | 1 | 0.3431 | predicted protein |
| contig_24765_5 | 0.1251 | 1 | 1 | 0.448 | 1 | 1 | 0.3431 | ABC transporter ATP-binding protein/permease |
| contig_38474_6 | 0.1254 | 1 | 1 | 0.448 | 1 | 1 | 0.3431 | phosphoserine transaminase |
| contig_30510_4 | 0.1255 | 1 | 1 | 0.448 | 1 | 1 | 0.3431 | 60S ribosomal protein |
| contig_36096_1 | 0.1257 | 1 | 1 | 0.448 | 1 | 1 | 0.3431 | predicted protein |
| contig_25874_5 | 0.1275 | 1 | 1 | 0.4534 | 1 | 1 | 0.3472 | predicted protein |
| contig_6467_5 | 0.1281 | 1 | 1 | 0.4544 | 1 | 1 | 0.348 | predicted protein |
| contig_17693_5 | 0.1285 | 1 | 1 | 0.4548 | 1 | 1 | 0.3483 | elongation factor-like |
| contig_12588_5 | 0.1289 | 1 | 1 | 0.455 | 1 | 1 | 0.3485 | predicted protein |
| contig_10600_6 | 0.1304 | 1 | 1 | 0.4595 | 1 | 1 | 0.3519 | predicted protein |

| | | | | | | | | |
|---|---|---|---|---|---|---|---|---|
| contig_17663_5 | 0.1308 | 1 | 1 | 0.4599 | 1 | 1 | 0.3522 | pyruvate dehydrogenase (acetyl-transferring) E1 component alpha subunit |
| contig_36050_1 | 0.1312 | 1 | 1 | 0.46 | 1 | 1 | 0.3522 | Light-harvesting complex I LH38 protein |
| contig_28238_5 | 0.1317 | 1 | 1 | 0.4607 | 1 | 1 | 0.3528 | ubiquinol-cytochrome c reductase iron-sulfur subunit |
| contig_19778_5 | 0.1334 | 1 | 1 | 0.4655 | 1 | 1 | 0.3565 | voltage-gated chloride channel |
| contig_28775_6 | 0.1352 | 1 | 1 | 0.4686 | 1 | 1 | 0.3588 | cullin-associated NEDD8-dissociated protein 1 |
| contig_25924_5 | 0.1353 | 1 | 1 | 0.4686 | 1 | 1 | 0.3588 | ATP-dependent Clp protease proteolytic subunit |
| contig_13819_5 | 0.1353 | 1 | 1 | 0.4686 | 1 | 1 | 0.3588 | 7-dehydrocholesterol reductase |
| contig_25157_1 | 0.1355 | 1 | 1 | 0.4686 | 1 | 1 | 0.3588 | magnesium chelatase subunit D |
| contig_16905_4 | 0.1361 | 1 | 1 | 0.4693 | 1 | 1 | 0.3594 | predicted protein |
| contig_2430_4 | 0.1368 | 1 | 1 | 0.4693 | 1 | 1 | 0.3594 | ubiquinol-cytochrome c r |
| contig_32529_3 | 0.137 | 1 | 1 | 0.4693 | 1 | 1 | 0.3594 | GDT1-like protein 2 chloroplastic |
| contig_33090_6 | 0.1371 | 1 | 1 | 0.4693 | 1 | 1 | 0.3594 | predicted protein |
| contig_5445_2 | 0.1372 | 1 | 1 | 0.4693 | 1 | 1 | 0.3594 | asparagine synthase B |
| contig_36089_2 | 0.138 | 1 | 1 | 0.4711 | 1 | 1 | 0.3608 | archaeal ribosomal protein S17P |
| contig_36190_5 | 0.1398 | 1 | 1 | 0.4748 | 1 | 1 | 0.3636 | long chain acyl-coa synthetase |
| contig_28224_2 | 0.1401 | 1 | 1 | 0.4748 | 1 | 1 | 0.3636 | alpha tubulin |
| contig_14974_2 | 0.1403 | 1 | 1 | 0.4748 | 1 | 1 | 0.3636 | dnaj subfamily b member 4 |
| contig_23061_6 | 0.1404 | 1 | 1 | 0.4748 | 1 | 1 | 0.3636 | predicted protein |
| contig_29766_3 | 0.1407 | 1 | 1 | 0.475 | 1 | 1 | 0.3637 | GMP synthase (glutamine-hydrolyzing) domain-containing protein |
| contig_25286_5 | 0.1423 | 1 | 1 | 0.479 | 1 | 1 | 0.3668 | ap-2 complex subunit mu |
| contig_40158_1 | 0.1428 | 1 | 1 | 0.479 | 1 | 1 | 0.3668 | extrinsic protein in photosystem II |
| contig_22578_5 | 0.1429 | 1 | 1 | 0.479 | 1 | 1 | 0.3668 | Urea-proton symporter DUR3 |
| contig_22533_3 | 0.1442 | 1 | 1 | 0.4813 | 1 | 1 | 0.3686 | predicted protein |
| contig_18580_6 | 0.1446 | 1 | 1 | 0.4813 | 1 | 1 | 0.3686 | predicted protein |
| contig_3085_5 | 0.145 | 1 | 1 | 0.4813 | 1 | 1 | 0.3686 | ferredoxin (chloroplast) |
| contig_29172_6 | 0.145 | 1 | 1 | 0.4813 | 1 | 1 | 0.3686 | arginyl-tRNA synthetase |

| | | | | | | | | |
|---|---|---|---|---|---|---|---|---|
| contig_38881_1 | 0.1451 | 1 | 1 | 0.4813 | 1 | 1 | 0.3686 | SulP family inorganic anion transporter |
| contig_10162_6 | 0.1463 | 1 | 1 | 0.4826 | 1 | 1 | 0.3696 | predicted protein |
| contig_36385_4 | 0.1464 | 1 | 1 | 0.4826 | 1 | 1 | 0.3696 | ---NA--- |
| contig_39259_1 | 0.1467 | 1 | 1 | 0.4826 | 1 | 1 | 0.3696 | preprotein-translocase subunit a (chloroplast) |
| contig_28903_4 | 0.147 | 1 | 1 | 0.4826 | 1 | 1 | 0.3696 | predicted protein |
| contig_21768_6 | 0.1472 | 1 | 1 | 0.4826 | 1 | 1 | 0.3696 | 40S ribosomal protein |
| contig_34578_1 | 0.1476 | 1 | 1 | 0.4826 | 1 | 1 | 0.3696 | predicted protein |
| contig_6077_5 | 0.1482 | 1 | 1 | 0.4826 | 1 | 1 | 0.3696 | 60s ribosomal protein l9 |
| contig_29454_5 | 0.1489 | 1 | 1 | 0.4826 | 1 | 1 | 0.3696 | peptidylprolyl isomerase |
| contig_30068_2 | 0.1491 | 1 | 1 | 0.4826 | 1 | 1 | 0.3696 | 40s ribosomal protein s4 |
| contig_24353_3 | 0.1492 | 1 | 1 | 0.4826 | 1 | 1 | 0.3696 | hemagglutinin/amebocyte aggregation factor |
| contig_22726_1 | 0.1497 | 1 | 1 | 0.4826 | 1 | 1 | 0.3696 | Valine-tRNA ligase |
| contig_5829_1 | 0.1498 | 1 | 1 | 0.4826 | 1 | 1 | 0.3696 | gametolysin peptidase M11 |
| contig_16707_4 | 0.1502 | 1 | 1 | 0.4826 | 1 | 1 | 0.3696 | nitrate reductase |
| contig_27384_5 | 0.1506 | 1 | 1 | 0.4826 | 1 | 1 | 0.3696 | predicted protein |
| contig_17114_6 | 0.1507 | 1 | 1 | 0.4826 | 1 | 1 | 0.3696 | predicted protein |
| contig_32617_4 | 0.1507 | 1 | 1 | 0.4826 | 1 | 1 | 0.3696 | VPLPA-CTERM protein sorting domain-containing protein |
| contig_25862_1 | 0.1509 | 1 | 1 | 0.4826 | 1 | 1 | 0.3696 | predicted protein |
| contig_8369_4 | 0.1522 | 1 | 1 | 0.4857 | 1 | 1 | 0.372 | Clp protease ATP binding subunit (chloroplast) |
| contig_19155_3 | 0.1526 | 1 | 1 | 0.4857 | 1 | 1 | 0.372 | geranylgeranyl reductase |
| contig_14080_4 | 0.1528 | 1 | 1 | 0.4857 | 1 | 1 | 0.372 | dihydroorotase |
| contig_38336_2 | 0.1534 | 1 | 1 | 0.4861 | 1 | 1 | 0.3722 | class I SAM-dependent methyltransferase |
| contig_6709_6 | 0.1537 | 1 | 1 | 0.4861 | 1 | 1 | 0.3722 | predicted protein |
| contig_25653_6 | 0.1542 | 1 | 1 | 0.4861 | 1 | 1 | 0.3722 | 40S ribosomal protein S8 |
| contig_34741_6 | 0.1545 | 1 | 1 | 0.4861 | 1 | 1 | 0.3722 | 60S ribosomal protein |
| contig_39564_2 | 0.1548 | 1 | 1 | 0.4861 | 1 | 1 | 0.3722 | predicted protein |
| contig_33607_5 | 0.1551 | 1 | 1 | 0.4861 | 1 | 1 | 0.3722 | S-adenosyl-L-methionine-dependent methyltransferase |

| | | | | | | | | |
|---|---|---|---|---|---|---|---|---|
| contig_6255_2 | 0.1551 | 1 | 1 | 0.4861 | 1 | 1 | 0.3722 | Nuclear transport factor 2 |
| contig_17402_3 | 0.1566 | 1 | 1 | 0.4887 | 1 | 1 | 0.3742 | hypothetical protein FRACYDRAFT_278091 |
| contig_9759_1 | 0.1569 | 1 | 1 | 0.4887 | 1 | 1 | 0.3742 | clathrin heavy chain |
| contig_33639_6 | 0.1569 | 1 | 1 | 0.4887 | 1 | 1 | 0.3742 | ---NA--- |
| contig_19428_2 | 0.1578 | 1 | 1 | 0.49 | 1 | 1 | 0.3752 | chloroplast beta-keto acyl reductase |
| contig_33347_6 | 0.158 | 1 | 1 | 0.49 | 1 | 1 | 0.3752 | V-type ATPase A subunit |
| contig_28354_5 | 0.1584 | 1 | 1 | 0.49 | 1 | 1 | 0.3752 | hypothetical protein FRACYDRAFT_169912 |
| contig_27196_4 | 0.1589 | 1 | 1 | 0.49 | 1 | 1 | 0.3752 | predicted protein |
| contig_24114_3 | 0.1589 | 1 | 1 | 0.49 | 1 | 1 | 0.3752 | predicted protein |
| contig_25721_4 | 0.1599 | 1 | 1 | 0.491 | 1 | 1 | 0.376 | putative manganese superoxide dismutase protein |
| contig_39581_5 | 0.1599 | 1 | 1 | 0.491 | 1 | 1 | 0.376 | cytochrome c1 |
| contig_5770_5 | 0.1614 | 1 | 1 | 0.4946 | 1 | 1 | 0.3788 | ketol-acid reductoisomerase |
| contig_26198_6 | 0.1625 | 1 | 1 | 0.4968 | 1 | 1 | 0.3804 | ribosomal protein L20 |
| contig_10952_4 | 0.1632 | 1 | 1 | 0.4974 | 1 | 1 | 0.3809 | diacylglycerol O-acyltransferase 2D |
| contig_30482_3 | 0.1637 | 1 | 1 | 0.4974 | 1 | 1 | 0.3809 | predicted protein |
| contig_10149_5 | 0.1645 | 1 | 1 | 0.4974 | 1 | 1 | 0.3809 | phosphoribosylformylglycinamidine synthase |
| contig_14784_5 | 0.1647 | 1 | 1 | 0.4974 | 1 | 1 | 0.3809 | atp-dependent RNA helicase |
| contig_24919_4 | 0.1648 | 1 | 1 | 0.4974 | 1 | 1 | 0.3809 | predicted protein |
| contig_10800_3 | 0.1649 | 1 | 1 | 0.4974 | 1 | 1 | 0.3809 | predicted protein |
| contig_14379_2 | 0.1653 | 1 | 1 | 0.4974 | 1 | 1 | 0.3809 | probable aspartyl aminopeptidase |
| contig_22967_5 | 0.1655 | 1 | 1 | 0.4974 | 1 | 1 | 0.3809 | predicted protein |
| contig_7075_5 | 0.1657 | 1 | 1 | 0.4974 | 1 | 1 | 0.3809 | predicted protein |
| contig_25696_5 | 0.166 | 1 | 1 | 0.4974 | 1 | 1 | 0.3809 | predicted protein |
| contig_789_5 | 0.167 | 1 | 1 | 0.4974 | 1 | 1 | 0.3809 | ribulose-1 5-bisphosphate carboxylase/oxygenase large subunit (chloroplast) |
| contig_9347_5 | 0.1673 | 1 | 1 | 0.4974 | 1 | 1 | 0.3809 | putative glycerol-3-phosphate transporter 1 |
| contig_22874_2 | 0.168 | 1 | 1 | 0.4974 | 1 | 1 | 0.3809 | inositol 5-phosphatase |

| | | | | | | | | |
|---|---|---|---|---|---|---|---|---|
| contig_26865_4 | 0.1682 | 1 | 1 | 0.4974 | 1 | 1 | 0.3809 | ---NA--- |
| contig_20020_4 | 0.1683 | 1 | 1 | 0.4974 | 1 | 1 | 0.3809 | predicted protein |
| contig_39519_2 | 0.1685 | 1 | 1 | 0.4974 | 1 | 1 | 0.3809 | predicted protein |
| contig_8511_4 | 0.1687 | 1 | 1 | 0.4974 | 1 | 1 | 0.3809 | O-acetylhomoserine aminocarboxypropyltransferase |
| contig_40323_4 | 0.169 | 1 | 1 | 0.4974 | 1 | 1 | 0.3809 | ---NA--- |
| contig_24980_4 | 0.1691 | 1 | 1 | 0.4974 | 1 | 1 | 0.3809 | Bystin |
| contig_29713_5 | 0.1701 | 1 | 1 | 0.4974 | 1 | 1 | 0.3809 | probable prefoldin subunit 2 |
| contig_40261_6 | 0.1703 | 1 | 1 | 0.4974 | 1 | 1 | 0.3809 | predicted protein |
| contig_32573_3 | 0.1708 | 1 | 1 | 0.4974 | 1 | 1 | 0.3809 | ATP synthase CF1 alpha subunit (chloroplast) |
| contig_33257_5 | 0.1709 | 1 | 1 | 0.4974 | 1 | 1 | 0.3809 | SulP family inorganic anion transporter |
| contig_33544_3 | 0.171 | 1 | 1 | 0.4974 | 1 | 1 | 0.3809 | predicted protein |
| contig_28237_2 | 0.171 | 1 | 1 | 0.4974 | 1 | 1 | 0.3809 | plant UBX domain-containing protein 10-like |
| contig_22734_3 | 0.1711 | 1 | 1 | 0.4974 | 1 | 1 | 0.3809 | predicted protein |
| contig_7021_2 | 0.172 | 1 | 1 | 0.4989 | 1 | 1 | 0.3821 | protein phosphatase 2C putative |
| contig_19765_6 | 0.1737 | 1 | 1 | 0.5031 | 1 | 1 | 0.3853 | predicted protein |
| contig_16381_6 | 0.1751 | 1 | 1 | 0.5038 | 1 | 1 | 0.3858 | predicted protein |
| contig_7979_4 | 0.1753 | 1 | 1 | 0.5038 | 1 | 1 | 0.3858 | protein fucoxanthin chlorophyll a/c protein |
| contig_41409_3 | 0.1759 | 1 | 1 | 0.5038 | 1 | 1 | 0.3858 | ribosomal protein L13 |
| contig_26227_3 | 0.1763 | 1 | 1 | 0.5038 | 1 | 1 | 0.3858 | 3-phosphoshikimate 1-carboxyvinyltransferase |
| contig_10429_5 | 0.1763 | 1 | 1 | 0.5038 | 1 | 1 | 0.3858 | hypothetical protein T265_11290 partial |
| contig_24914_6 | 0.1764 | 1 | 1 | 0.5038 | 1 | 1 | 0.3858 | bifunctional dTDP-4-dehydrorhamnose 3 5-epimerase/dTDP-4-dehydrorhamnose reductase-like |
| contig_13922_3 | 0.1765 | 1 | 1 | 0.5038 | 1 | 1 | 0.3858 | annexin |
| contig_24813_3 | 0.1766 | 1 | 1 | 0.5038 | 1 | 1 | 0.3858 | predicted protein |
| contig_38368_6 | 0.1782 | 1 | 1 | 0.5075 | 1 | 1 | 0.3886 | glutathione S-transferase |
| contig_34140_1 | 0.179 | 1 | 1 | 0.5076 | 1 | 1 | 0.3887 | predicted protein |
| contig_23563_5 | 0.1796 | 1 | 1 | 0.5076 | 1 | 1 | 0.3887 | FK506-binding protein 2 isoform X2 |

| | | | | | | | | |
|---|---|---|---|---|---|---|---|---|
| contig_6818_2 | 0.1796 | 1 | 1 | 0.5076 | 1 | 1 | 0.3887 | predicted protein |
| contig_35394_6 | 0.1799 | 1 | 1 | 0.5076 | 1 | 1 | 0.3887 | cytochrome c |
| contig_13739_4 | 0.1802 | 1 | 1 | 0.5076 | 1 | 1 | 0.3887 | protein DETOXIFICATION 46 chloroplastic |
| contig_35287_3 | 0.1806 | 1 | 1 | 0.5076 | 1 | 1 | 0.3887 | photosystem II PsbU protein |
| contig_38390_3 | 0.1808 | 1 | 1 | 0.5076 | 1 | 1 | 0.3887 | predicted protein |
| contig_30765_6 | 0.1815 | 1 | 1 | 0.5076 | 1 | 1 | 0.3887 | dynactin subunit 1 |
| contig_40859_3 | 0.1815 | 1 | 1 | 0.5076 | 1 | 1 | 0.3887 | predicted protein |
| contig_39585_3 | 0.1816 | 1 | 1 | 0.5076 | 1 | 1 | 0.3887 | predicted protein |
| contig_26554_3 | 0.1834 | 1 | 1 | 0.5115 | 1 | 1 | 0.3917 | predicted protein |
| contig_21931_4 | 0.1836 | 1 | 1 | 0.5115 | 1 | 1 | 0.3917 | chaperonin GroEL (chloroplast) |
| contig_27994_5 | 0.1841 | 1 | 1 | 0.5119 | 1 | 1 | 0.392 | nitrate transporter |
| contig_22875_4 | 0.1848 | 1 | 1 | 0.5127 | 1 | 1 | 0.3926 | calcium-translocating P-type ATPase PMCA-type |
| contig_26596_5 | 0.1851 | 1 | 1 | 0.5129 | 1 | 1 | 0.3927 | leucyl aminopeptidase |
| contig_32503_1 | 0.1869 | 1 | 1 | 0.5169 | 1 | 1 | 0.3958 | predicted protein |
| contig_9535_1 | 0.188 | 1 | 1 | 0.5185 | 1 | 1 | 0.397 | transitional endoplasmic reticulum ATPase |
| contig_7536_5 | 0.1882 | 1 | 1 | 0.5185 | 1 | 1 | 0.397 | predicted protein |
| contig_35860_4 | 0.1889 | 1 | 1 | 0.5188 | 1 | 1 | 0.3972 | mitochondrial manganese superoxide dismutase |
| contig_25806_5 | 0.189 | 1 | 1 | 0.5188 | 1 | 1 | 0.3972 | 2-C-methyl-D-erythritol 2 4-cyclodiphosphate synthase |
| contig_11078_3 | 0.1896 | 1 | 1 | 0.5194 | 1 | 1 | 0.3978 | peroxiredoxin |
| contig_28209_6 | 0.1903 | 1 | 1 | 0.5197 | 1 | 1 | 0.3979 | dihydrolipoamide acetyltransferase |
| contig_14977_5 | 0.1905 | 1 | 1 | 0.5197 | 1 | 1 | 0.3979 | predicted protein |
| contig_25589_3 | 0.1907 | 1 | 1 | 0.5197 | 1 | 1 | 0.3979 | small subunit ribosomal protein S3Ae |
| contig_5563_5 | 0.1913 | 1 | 1 | 0.5204 | 1 | 1 | 0.3985 | predicted protein |
| contig_7641_5 | 0.1929 | 1 | 1 | 0.5229 | 1 | 1 | 0.4004 | precursor cytochrome c6 |
| contig_12026_5 | 0.193 | 1 | 1 | 0.5229 | 1 | 1 | 0.4004 | predicted protein |
| contig_28172_4 | 0.1932 | 1 | 1 | 0.5229 | 1 | 1 | 0.4004 | 60S ribosomal protein L5 |
| contig_37802_5 | 0.194 | 1 | 1 | 0.524 | 1 | 1 | 0.4013 | predicted protein |

| | | | | | | | | |
|---|---|---|---|---|---|---|---|---|
| contig_32280_2 | 0.1943 | 1 | 1 | 0.524 | 1 | 1 | 0.4013 | plastid sedoheptulose-1 7-bisphosphatase |
| contig_7030_6 | 0.1948 | 1 | 1 | 0.5244 | 1 | 1 | 0.4016 | ---NA--- |
| contig_26772_5 | 0.1974 | 1 | 1 | 0.5306 | 1 | 1 | 0.4063 | predicted protein |
| contig_28624_5 | 0.1978 | 1 | 1 | 0.5306 | 1 | 1 | 0.4063 | eukaryotic peptide chain release factor subunit 1 |
| contig_5548_1 | 0.1995 | 1 | 1 | 0.5341 | 1 | 1 | 0.409 | geranylgeranyl pyrophosphate synthase |
| contig_25820_1 | 0.2007 | 1 | 1 | 0.536 | 1 | 1 | 0.4105 | predicted protein |
| contig_12419_1 | 0.2009 | 1 | 1 | 0.536 | 1 | 1 | 0.4105 | T-complex protein 1 subunit delta |
| contig_27149_6 | 0.2012 | 1 | 1 | 0.536 | 1 | 1 | 0.4105 | homogentisate phytyltransferase |
| contig_7288_5 | 0.2016 | 1 | 1 | 0.5361 | 1 | 1 | 0.4105 | protein fucoxanthin chlorophyll a/c protein |
| contig_13609_2 | 0.2027 | 1 | 1 | 0.5382 | 1 | 1 | 0.4121 | U5 small nuclear ribonucleoprotein 200 kDa helicase |
| contig_16774_4 | 0.2039 | 1 | 1 | 0.54 | 1 | 1 | 0.4135 | probable microsomal very long chain fatty acid elongase |
| contig_27950_6 | 0.2041 | 1 | 1 | 0.54 | 1 | 1 | 0.4135 | aspartyl-tRNA synthetase |
| contig_28761_2 | 0.2048 | 1 | 1 | 0.5408 | 1 | 1 | 0.4142 | protein dispatched homolog 3-like |
| contig_28417_5 | 0.2068 | 1 | 1 | 0.5448 | 1 | 1 | 0.4172 | predicted protein |
| contig_30091_6 | 0.207 | 1 | 1 | 0.5448 | 1 | 1 | 0.4172 | predicted protein |
| contig_7847_6 | 0.2082 | 1 | 1 | 0.5462 | 1 | 1 | 0.4183 | predicted protein |
| contig_15399_2 | 0.2085 | 1 | 1 | 0.5462 | 1 | 1 | 0.4183 | predicted protein |
| contig_23995_6 | 0.2086 | 1 | 1 | 0.5462 | 1 | 1 | 0.4183 | predicted protein |
| contig_22594_4 | 0.2094 | 1 | 1 | 0.5473 | 1 | 1 | 0.4191 | magnesium chelatase subunit H |
| contig_16462_2 | 0.2105 | 1 | 1 | 0.5486 | 1 | 1 | 0.4201 | chorismate synthase |
| contig_24756_6 | 0.211 | 1 | 1 | 0.5486 | 1 | 1 | 0.4201 | tropinone reductase |
| contig_15153_4 | 0.2118 | 1 | 1 | 0.5486 | 1 | 1 | 0.4201 | Ras-like GTP-binding protein YPT1 |
| contig_38597_3 | 0.2119 | 1 | 1 | 0.5486 | 1 | 1 | 0.4201 | ---NA--- |
| contig_27033_6 | 0.2119 | 1 | 1 | 0.5486 | 1 | 1 | 0.4201 | glucokinase |
| contig_9136_5 | 0.2121 | 1 | 1 | 0.5486 | 1 | 1 | 0.4201 | predicted protein |
| contig_16863_4 | 0.2129 | 1 | 1 | 0.5486 | 1 | 1 | 0.4201 | D-3-phosphoglycerate dehydrogenase |
| contig_30516_6 | 0.2131 | 1 | 1 | 0.5486 | 1 | 1 | 0.4201 | aspartate--ammonia ligase |

| | | | | | | | | |
|---|---|---|---|---|---|---|---|---|
| contig_11948_3 | 0.2135 | 1 | 1 | 0.5486 | 1 | 1 | 0.4201 | aldo/keto reductase |
| contig_25130_6 | 0.2135 | 1 | 1 | 0.5486 | 1 | 1 | 0.4201 | methionine-tRNA ligase |
| contig_23545_2 | 0.2143 | 1 | 1 | 0.5487 | 1 | 1 | 0.4202 | predicted protein |
| contig_10829_2 | 0.2144 | 1 | 1 | 0.5487 | 1 | 1 | 0.4202 | plastid fructose 1 6-bisphosphatase isoform 1 |
| contig_5908_5 | 0.2146 | 1 | 1 | 0.5487 | 1 | 1 | 0.4202 | predicted protein |
| contig_33755_6 | 0.215 | 1 | 1 | 0.5487 | 1 | 1 | 0.4202 | predicted protein |
| contig_39499_3 | 0.2158 | 1 | 1 | 0.55 | 1 | 1 | 0.4212 | adenylosuccinate synthetase |
| contig_28450_6 | 0.2174 | 1 | 1 | 0.5521 | 1 | 1 | 0.4228 | transketolase |
| contig_6450_1 | 0.2182 | 1 | 1 | 0.5521 | 1 | 1 | 0.4228 | glutamine-fructose-6-phosphate transaminase |
| contig_12639_1 | 0.2184 | 1 | 1 | 0.5521 | 1 | 1 | 0.4228 | ADP-ribosylation factor-related protein 1 |
| contig_20820_5 | 0.2184 | 1 | 1 | 0.5521 | 1 | 1 | 0.4228 | photosystem II protein D1 (chloroplast) |
| contig_40524_3 | 0.2186 | 1 | 1 | 0.5521 | 1 | 1 | 0.4228 | light harvesting protein |
| contig_36067_5 | 0.2188 | 1 | 1 | 0.5521 | 1 | 1 | 0.4228 | silicon transporter |
| contig_28732_1 | 0.2202 | 1 | 1 | 0.5543 | 1 | 1 | 0.4245 | light harvesting protein |
| contig_25123_5 | 0.2204 | 1 | 1 | 0.5543 | 1 | 1 | 0.4245 | ubiquitin-activating enzyme E1 protein 3 |
| contig_5607_2 | 0.221 | 1 | 1 | 0.5547 | 1 | 1 | 0.4248 | protease SohB |
| contig_22753_2 | 0.2221 | 1 | 1 | 0.556 | 1 | 1 | 0.4258 | nicotinate phosphoribosyltransferase 2-like |
| contig_7982_5 | 0.2225 | 1 | 1 | 0.556 | 1 | 1 | 0.4258 | photosystem II protein V (chloroplast) |
| contig_29915_2 | 0.2227 | 1 | 1 | 0.556 | 1 | 1 | 0.4258 | nucleolar protein nop56 |
| contig_30444_6 | 0.2232 | 1 | 1 | 0.556 | 1 | 1 | 0.4258 | NADH dehydrogenase (ubiquinone) 1 beta subcomplex subunit 9 |
| contig_16700_6 | 0.2234 | 1 | 1 | 0.556 | 1 | 1 | 0.4258 | serine decarboxylase |
| contig_34264_6 | 0.2237 | 1 | 1 | 0.556 | 1 | 1 | 0.4258 | probable small nuclear ribonucleoprotein E |
| contig_28392_3 | 0.2243 | 1 | 1 | 0.5568 | 1 | 1 | 0.4264 | predicted protein |
| contig_17554_4 | 0.2248 | 1 | 1 | 0.5571 | 1 | 1 | 0.4266 | predicted protein |
| contig_26646_2 | 0.2276 | 1 | 1 | 0.5632 | 1 | 1 | 0.4312 | short-chain dehydrogenase/reductase acting with NAD or NADP as acceptor |
| contig_23311_6 | 0.2294 | 1 | 1 | 0.5659 | 1 | 1 | 0.4334 | predicted protein |

| | | | | | | | | |
|---|---|---|---|---|---|---|---|---|
| contig_32710_4 | 0.2295 | 1 | 1 | 0.5659 | 1 | 1 | 0.4334 | class I SAM-dependent methyltransferase |
| contig_23323_6 | 0.2302 | 1 | 1 | 0.5669 | 1 | 1 | 0.4341 | predicted protein |
| contig_3511_5 | 0.2316 | 1 | 1 | 0.5688 | 1 | 1 | 0.4355 | Ycf89 (chloroplast) |
| contig_26348_5 | 0.2321 | 1 | 1 | 0.5688 | 1 | 1 | 0.4355 | heat shock protein 70 |
| contig_36986_1 | 0.2323 | 1 | 1 | 0.5688 | 1 | 1 | 0.4355 | ---NA--- |
| contig_6576_3 | 0.2325 | 1 | 1 | 0.5688 | 1 | 1 | 0.4355 | protein COFACTOR ASSEMBLY OF COMPLEX C SUBUNIT B CCB1 chloroplastic |
| contig_21620_6 | 0.2334 | 1 | 1 | 0.5701 | 1 | 1 | 0.4365 | predicted protein |
| contig_19673_5 | 0.2355 | 1 | 1 | 0.5727 | 1 | 1 | 0.4385 | predicted protein |
| contig_27062_5 | 0.2359 | 1 | 1 | 0.5727 | 1 | 1 | 0.4385 | heat shock protein 70 |
| contig_15734_6 | 0.2363 | 1 | 1 | 0.5727 | 1 | 1 | 0.4385 | ALA dehydratase |
| contig_41330_2 | 0.2365 | 1 | 1 | 0.5727 | 1 | 1 | 0.4385 | 60S ribosomal protein L23a |
| contig_23471_1 | 0.2365 | 1 | 1 | 0.5727 | 1 | 1 | 0.4385 | predicted protein |
| contig_12843_3 | 0.2367 | 1 | 1 | 0.5727 | 1 | 1 | 0.4385 | proteasome subunit beta type-3 |
| contig_30585_6 | 0.2371 | 1 | 1 | 0.5727 | 1 | 1 | 0.4385 | ribosomal protein S2 |
| contig_40102_2 | 0.2379 | 1 | 1 | 0.5737 | 1 | 1 | 0.4393 | predicted protein |
| contig_40133_3 | 0.2395 | 1 | 1 | 0.5767 | 1 | 1 | 0.4416 | 30S ribosomal protein S15 |
| contig_9827_1 | 0.2416 | 1 | 1 | 0.5808 | 1 | 1 | 0.4448 | predicted protein |
| contig_14784_2 | 0.242 | 1 | 1 | 0.5808 | 1 | 1 | 0.4448 | atp-dependent RNA helicase |
| contig_20096_5 | 0.2426 | 1 | 1 | 0.5808 | 1 | 1 | 0.4448 | predicted protein |
| contig_6711_4 | 0.2427 | 1 | 1 | 0.5808 | 1 | 1 | 0.4448 | predicted protein |
| contig_31457_1 | 0.2434 | 1 | 1 | 0.5814 | 1 | 1 | 0.4452 | predicted protein |
| contig_9649_2 | 0.2443 | 1 | 1 | 0.5827 | 1 | 1 | 0.4462 | predicted protein |
| contig_26152_4 | 0.245 | 1 | 1 | 0.5828 | 1 | 1 | 0.4463 | inositol 5-phosphatase |
| contig_13881_2 | 0.2451 | 1 | 1 | 0.5828 | 1 | 1 | 0.4463 | predicted protein |
| contig_5813_3 | 0.2466 | 1 | 1 | 0.5854 | 1 | 1 | 0.4483 | transketolase |
| contig_7686_2 | 0.2469 | 1 | 1 | 0.5854 | 1 | 1 | 0.4483 | glutathione-disulfide reductase |

| | | | | | | | | |
|---|---|---|---|---|---|---|---|---|
| contig_34797_4 | 0.2482 | 1 | 1 | 0.5875 | 1 | 1 | 0.4499 | GTP-binding protein SAR1 |
| contig_22327_4 | 0.2495 | 1 | 1 | 0.5896 | 1 | 1 | 0.4515 | gamma-aminobutyric-acid receptor subunit beta |
| contig_30620_5 | 0.2506 | 1 | 1 | 0.5913 | 1 | 1 | 0.4528 | Transmembrane protein 87B |
| contig_3295_5 | 0.251 | 1 | 1 | 0.5913 | 1 | 1 | 0.4528 | V-type ATPase F subunit |
| contig_14735_6 | 0.2515 | 1 | 1 | 0.5917 | 1 | 1 | 0.4531 | proteasome subunit beta type-5-like |
| contig_25957_2 | 0.2529 | 1 | 1 | 0.594 | 1 | 1 | 0.4549 | predicted protein |
| contig_6119_2 | 0.2548 | 1 | 1 | 0.5969 | 1 | 1 | 0.4571 | putative tRNA (cytidine(32)/guanosine(34)-2'-O)-methyltransferase |
| contig_30700_3 | 0.2549 | 1 | 1 | 0.5969 | 1 | 1 | 0.4571 | predicted protein |
| contig_6653_5 | 0.2566 | 1 | 1 | 0.5999 | 1 | 1 | 0.4593 | chorismate mutase |
| contig_16552_4 | 0.2571 | 1 | 1 | 0.6002 | 1 | 1 | 0.4596 | predicted protein |
| contig_29362_6 | 0.2578 | 1 | 1 | 0.6009 | 1 | 1 | 0.4602 | Stimulated by retinoic acid gene 6 protein |
| contig_34506_1 | 0.2599 | 1 | 1 | 0.605 | 1 | 1 | 0.4633 | cell division protein FtsH |
| contig_38523_3 | 0.2613 | 1 | 1 | 0.6066 | 1 | 1 | 0.4645 | h+-translocating pyrophosphatase family |
| contig_11742_4 | 0.2614 | 1 | 1 | 0.6066 | 1 | 1 | 0.4645 | predicted protein |
| contig_26293_4 | 0.2619 | 1 | 1 | 0.6067 | 1 | 1 | 0.4646 | predicted protein |
| contig_12711_6 | 0.2638 | 1 | 1 | 0.6093 | 1 | 1 | 0.4666 | predicted protein |
| contig_3444_2 | 0.2642 | 1 | 1 | 0.6093 | 1 | 1 | 0.4666 | protoporphyrin IX methyltransferase |
| contig_24988_5 | 0.2645 | 1 | 1 | 0.6093 | 1 | 1 | 0.4666 | Urea-proton symporter DUR3 |
| contig_23318_6 | 0.2649 | 1 | 1 | 0.6093 | 1 | 1 | 0.4666 | putative LOV domain-containing protein |
| contig_18211_4 | 0.2651 | 1 | 1 | 0.6093 | 1 | 1 | 0.4666 | light harvesting protein |
| contig_20441_2 | 0.2663 | 1 | 1 | 0.6093 | 1 | 1 | 0.4666 | putative glyoxalase I |
| contig_33643_5 | 0.2665 | 1 | 1 | 0.6093 | 1 | 1 | 0.4666 | predicted protein |
| contig_18892_6 | 0.2676 | 1 | 1 | 0.6093 | 1 | 1 | 0.4666 | predicted protein |
| contig_15223_4 | 0.2677 | 1 | 1 | 0.6093 | 1 | 1 | 0.4666 | predicted protein |
| contig_28144_3 | 0.2682 | 1 | 1 | 0.6093 | 1 | 1 | 0.4666 | ---NA--- |
| contig_25927_6 | 0.2682 | 1 | 1 | 0.6093 | 1 | 1 | 0.4666 | light harvesting protein |

| | | | | | | | | |
|---|---|---|---|---|---|---|---|---|
| contig_28722_5 | 0.2683 | 1 | 1 | 0.6093 | 1 | 1 | 0.4666 | predicted protein |
| contig_13566_4 | 0.2683 | 1 | 1 | 0.6093 | 1 | 1 | 0.4666 | 60S ribosomal protein L35a-4 |
| contig_20792_5 | 0.2686 | 1 | 1 | 0.6093 | 1 | 1 | 0.4666 | ---NA--- |
| contig_9105_2 | 0.2701 | 1 | 1 | 0.6118 | 1 | 1 | 0.4685 | SufE-like protein 1 chloroplastic/mitochondrial |
| contig_16777_4 | 0.2718 | 1 | 1 | 0.6142 | 1 | 1 | 0.4703 | H - or Na -translocating F-type V-type and A-type ATPase (F-ATPase) Superfamily |
| contig_28396_1 | 0.2719 | 1 | 1 | 0.6142 | 1 | 1 | 0.4703 | chaperone DnaK |
| contig_24278_4 | 0.2727 | 1 | 1 | 0.6149 | 1 | 1 | 0.4709 | predicted protein |
| contig_33528_6 | 0.2739 | 1 | 1 | 0.6169 | 1 | 1 | 0.4724 | SPX domain-containing membrane protein At4g22990-like |
| contig_25458_4 | 0.2744 | 1 | 1 | 0.6169 | 1 | 1 | 0.4724 | succinate dehydrogenase [ubiquinone] iron-sulfur subunit mitochondrial |
| contig_30555_4 | 0.2754 | 1 | 1 | 0.6182 | 1 | 1 | 0.4734 | fructose-1 6-bisphosphate aldolase precursor |
| contig_8803_5 | 0.277 | 1 | 1 | 0.6199 | 1 | 1 | 0.4747 | predicted protein |
| contig_39949_2 | 0.2775 | 1 | 1 | 0.6199 | 1 | 1 | 0.4747 | predicted protein |
| contig_25619_1 | 0.2777 | 1 | 1 | 0.6199 | 1 | 1 | 0.4747 | ornithine--oxo-acid transaminase |
| contig_30107_6 | 0.2779 | 1 | 1 | 0.6199 | 1 | 1 | 0.4747 | ATP-dependent protease HslVU ATPase subunit |
| contig_37770_2 | 0.2788 | 1 | 1 | 0.6199 | 1 | 1 | 0.4747 | photosystem I P700 apoprotein A1 (chloroplast) |
| contig_29925_5 | 0.2792 | 1 | 1 | 0.6199 | 1 | 1 | 0.4747 | glutamine synthetase |
| contig_5869_1 | 0.2796 | 1 | 1 | 0.6199 | 1 | 1 | 0.4747 | threonine-tRNA ligase |
| contig_33877_5 | 0.2798 | 1 | 1 | 0.6199 | 1 | 1 | 0.4747 | ---NA--- |
| contig_23352_4 | 0.2806 | 1 | 1 | 0.6199 | 1 | 1 | 0.4747 | malate dehydrogenase |
| contig_22713_3 | 0.2811 | 1 | 1 | 0.6199 | 1 | 1 | 0.4747 | predicted protein |
| contig_13636_6 | 0.2815 | 1 | 1 | 0.6199 | 1 | 1 | 0.4747 | acyl-CoA dehydrogenase-like protein |
| contig_6495_6 | 0.2819 | 1 | 1 | 0.6199 | 1 | 1 | 0.4747 | predicted protein |
| contig_38301_1 | 0.282 | 1 | 1 | 0.6199 | 1 | 1 | 0.4747 | predicted protein |
| contig_30867_1 | 0.2822 | 1 | 1 | 0.6199 | 1 | 1 | 0.4747 | Light-harvesting complex I LH38 protein |
| contig_37899_3 | 0.2822 | 1 | 1 | 0.6199 | 1 | 1 | 0.4747 | ribosomal protein L2 (chloroplast) |
| contig_26990_2 | 0.2828 | 1 | 1 | 0.6203 | 1 | 1 | 0.475 | uracil phosphoribosyltransferase |

| | | | | | | | | |
|---|---|---|---|---|---|---|---|---|
| contig_24220_6 | 0.284 | 1 | 1 | 0.6221 | 1 | 1 | 0.4764 | ATP-binding cassette superfamily |
| contig_10572_1 | 0.2853 | 1 | 1 | 0.6241 | 1 | 1 | 0.4779 | NADH dehydrogenase [ubiquinone] flavoprotein 1 mitochondrial |
| contig_23827_5 | 0.2866 | 1 | 1 | 0.626 | 1 | 1 | 0.4794 | predicted protein |
| contig_27562_4 | 0.2887 | 1 | 1 | 0.6269 | 1 | 1 | 0.4801 | acyl-CoA dehydrogenase |
| contig_39034_5 | 0.2892 | 1 | 1 | 0.6269 | 1 | 1 | 0.4801 | predicted protein |
| contig_25962_4 | 0.2897 | 1 | 1 | 0.6269 | 1 | 1 | 0.4801 | predicted protein |
| contig_34911_5 | 0.2898 | 1 | 1 | 0.6269 | 1 | 1 | 0.4801 | predicted protein |
| contig_34431_6 | 0.2911 | 1 | 1 | 0.6269 | 1 | 1 | 0.4801 | predicted protein |
| contig_6933_4 | 0.2916 | 1 | 1 | 0.6269 | 1 | 1 | 0.4801 | methylmalonate-semialdehyde dehydrogenase (acylating) |
| contig_29981_4 | 0.2921 | 1 | 1 | 0.6269 | 1 | 1 | 0.4801 | 20S proteasome subunit alpha 4 |
| contig_39450_1 | 0.2922 | 1 | 1 | 0.6269 | 1 | 1 | 0.4801 | CTL-like protein DDB_G0288717 |
| contig_15445_5 | 0.2923 | 1 | 1 | 0.6269 | 1 | 1 | 0.4801 | zeaxanthin epoxidase |
| contig_11033_5 | 0.2928 | 1 | 1 | 0.6269 | 1 | 1 | 0.4801 | predicted protein |
| contig_30717_5 | 0.2936 | 1 | 1 | 0.6269 | 1 | 1 | 0.4801 | predicted protein |
| contig_9763_5 | 0.2937 | 1 | 1 | 0.6269 | 1 | 1 | 0.4801 | predicted protein |
| contig_28195_3 | 0.2938 | 1 | 1 | 0.6269 | 1 | 1 | 0.4801 | predicted protein |
| contig_15143_1 | 0.2942 | 1 | 1 | 0.6269 | 1 | 1 | 0.4801 | CK1/CK1/CK1-D protein kinase variant |
| contig_14305_2 | 0.2951 | 1 | 1 | 0.6269 | 1 | 1 | 0.4801 | predicted protein |
| contig_12625_2 | 0.2953 | 1 | 1 | 0.6269 | 1 | 1 | 0.4801 | 26S proteasome non-ATPase regulatory subunit 7 |
| contig_25782_5 | 0.2955 | 1 | 1 | 0.6269 | 1 | 1 | 0.4801 | predicted protein |
| contig_24778_4 | 0.2957 | 1 | 1 | 0.6269 | 1 | 1 | 0.4801 | predicted protein |
| contig_22256_3 | 0.2959 | 1 | 1 | 0.6269 | 1 | 1 | 0.4801 | 30S ribosomal protein S15 |
| contig_5503_5 | 0.296 | 1 | 1 | 0.6269 | 1 | 1 | 0.4801 | predicted protein |
| contig_12446_4 | 0.2963 | 1 | 1 | 0.6269 | 1 | 1 | 0.4801 | UPF0187 protein |
| contig_27544_6 | 0.2965 | 1 | 1 | 0.6269 | 1 | 1 | 0.4801 | Long-chain-fatty-acid--CoA ligase putative |
| contig_37059_1 | 0.2971 | 1 | 1 | 0.6269 | 1 | 1 | 0.4801 | Ura6p |
| contig_23765_6 | 0.2972 | 1 | 1 | 0.6269 | 1 | 1 | 0.4801 | predicted protein |

| | | | | | | | | |
|---|---|---|---|---|---|---|---|---|
| contig_29531_5 | 0.2973 | 1 | 1 | 0.6269 | 1 | 1 | 0.4801 | oxidoreductase putative |
| contig_16247_6 | 0.2991 | 1 | 1 | 0.6299 | 1 | 1 | 0.4824 | imidazoleglycerol-phosphate dehydratase |
| contig_19241_5 | 0.3012 | 1 | 1 | 0.6334 | 1 | 1 | 0.485 | ---NA--- |
| contig_24083_4 | 0.3019 | 1 | 1 | 0.6337 | 1 | 1 | 0.4853 | predicted protein |
| contig_9858_4 | 0.3023 | 1 | 1 | 0.6337 | 1 | 1 | 0.4853 | argininosuccinate synthase |
| contig_27972_6 | 0.3025 | 1 | 1 | 0.6337 | 1 | 1 | 0.4853 | atp-dependent rna helicase dbp5 |
| contig_22602_4 | 0.3035 | 1 | 1 | 0.6349 | 1 | 1 | 0.4862 | putative all-trans-retinol 13 14-reductase |
| contig_16700_1 | 0.3054 | 1 | 1 | 0.6379 | 1 | 1 | 0.4885 | serine decarboxylase |
| contig_26180_3 | 0.306 | 1 | 1 | 0.6383 | 1 | 1 | 0.4888 | predicted protein |
| contig_23311_3 | 0.308 | 1 | 1 | 0.6412 | 1 | 1 | 0.491 | predicted protein |
| contig_39062_4 | 0.3083 | 1 | 1 | 0.6412 | 1 | 1 | 0.491 | 2-isopropylmalate synthase |
| contig_13656_4 | 0.3086 | 1 | 1 | 0.6412 | 1 | 1 | 0.491 | predicted protein |
| contig_23649_2 | 0.3095 | 1 | 1 | 0.6418 | 1 | 1 | 0.4914 | heat shock protein 90-2 |
| contig_5781_6 | 0.3102 | 1 | 1 | 0.6418 | 1 | 1 | 0.4914 | acetyl-coenzyme a synthetase |
| contig_30020_5 | 0.3102 | 1 | 1 | 0.6418 | 1 | 1 | 0.4914 | chaperonin GroEL |
| contig_6155_6 | 0.311 | 1 | 1 | 0.6418 | 1 | 1 | 0.4914 | light harvesting protein |
| contig_17360_5 | 0.311 | 1 | 1 | 0.6418 | 1 | 1 | 0.4914 | Annexin protein |
| contig_10732_2 | 0.3134 | 1 | 1 | 0.6457 | 1 | 1 | 0.4945 | transaldolase |
| contig_29765_5 | 0.3154 | 1 | 1 | 0.6478 | 1 | 1 | 0.4961 | predicted protein |
| contig_12144_4 | 0.3156 | 1 | 1 | 0.6478 | 1 | 1 | 0.4961 | FAD-binding domain-containing protein |
| contig_16980_6 | 0.3162 | 1 | 1 | 0.6478 | 1 | 1 | 0.4961 | translation initiation factor 3 subunit A |
| contig_32725_2 | 0.3164 | 1 | 1 | 0.6478 | 1 | 1 | 0.4961 | predicted protein |
| contig_8346_5 | 0.3165 | 1 | 1 | 0.6478 | 1 | 1 | 0.4961 | putative plastid division protein (chloroplast) |
| contig_10966_3 | 0.317 | 1 | 1 | 0.648 | 1 | 1 | 0.4962 | predicted protein |
| contig_7506_4 | 0.3175 | 1 | 1 | 0.6482 | 1 | 1 | 0.4963 | apocytochrome b (mitochondrion) |
| contig_38598_5 | 0.3181 | 1 | 1 | 0.6485 | 1 | 1 | 0.4966 | Adenylyl cyclase-associated protein |
| contig_39293_4 | 0.3187 | 1 | 1 | 0.6489 | 1 | 1 | 0.4969 | predicted protein |

| | | | | | | | | |
|---|---|---|---|---|---|---|---|---|
| contig_27013_6 | 0.3193 | 1 | 1 | 0.6493 | 1 | 1 | 0.4972 | sulfite reductase ferredoxin dependent |
| contig_35278_2 | 0.3209 | 1 | 1 | 0.6515 | 1 | 1 | 0.4989 | light harvesting protein |
| contig_30065_5 | 0.3226 | 1 | 1 | 0.6541 | 1 | 1 | 0.5009 | importin subunit alpha-1-like isoform X2 |
| contig_8012_5 | 0.3242 | 1 | 1 | 0.6565 | 1 | 1 | 0.5027 | predicted protein |
| contig_25971_2 | 0.3261 | 1 | 1 | 0.6595 | 1 | 1 | 0.505 | purine nucleoside phosphorylase I inosine and guanosine-specific |
| contig_34763_6 | 0.3271 | 1 | 1 | 0.6608 | 1 | 1 | 0.506 | myosin light chain kinase |
| contig_16919_4 | 0.3284 | 1 | 1 | 0.6625 | 1 | 1 | 0.5073 | predicted protein |
| contig_8868_5 | 0.3294 | 1 | 1 | 0.663 | 1 | 1 | 0.5077 | Enoyl-CoA hydratase |
| contig_25343_2 | 0.3295 | 1 | 1 | 0.663 | 1 | 1 | 0.5077 | predicted protein |
| contig_27320_3 | 0.3305 | 1 | 1 | 0.6637 | 1 | 1 | 0.5083 | transketolase |
| contig_10491_6 | 0.3308 | 1 | 1 | 0.6637 | 1 | 1 | 0.5083 | glutamine synthetase catalytic domain protein protein |
| contig_24202_5 | 0.3313 | 1 | 1 | 0.6639 | 1 | 1 | 0.5084 | 2-isopropylmalate synthase |
| contig_29877_2 | 0.3326 | 1 | 1 | 0.6651 | 1 | 1 | 0.5093 | predicted protein |
| contig_29793_4 | 0.3328 | 1 | 1 | 0.6651 | 1 | 1 | 0.5093 | predicted protein |
| contig_35584_4 | 0.3338 | 1 | 1 | 0.6662 | 1 | 1 | 0.5102 | nitrate transporter |
| contig_14990_5 | 0.3366 | 1 | 1 | 0.6711 | 1 | 1 | 0.5139 | heme oxygenase 1 chloroplastic-like |
| contig_25542_6 | 0.3383 | 1 | 1 | 0.6735 | 1 | 1 | 0.5158 | ---NA--- |
| contig_34233_5 | 0.3396 | 1 | 1 | 0.6746 | 1 | 1 | 0.5166 | predicted protein |
| contig_33583_4 | 0.3397 | 1 | 1 | 0.6746 | 1 | 1 | 0.5166 | hypersensitive-induced response protein 1-like |
| contig_29992_5 | 0.3409 | 1 | 1 | 0.6756 | 1 | 1 | 0.5174 | ---NA--- |
| contig_32573_2 | 0.3411 | 1 | 1 | 0.6756 | 1 | 1 | 0.5174 | ATP synthase CF1 alpha subunit (chloroplast) |
| contig_41187_5 | 0.342 | 1 | 1 | 0.6765 | 1 | 1 | 0.5181 | proteasome subunit beta type 4 |
| contig_38239_5 | 0.3432 | 1 | 1 | 0.6772 | 1 | 1 | 0.5186 | predicted protein |
| contig_25252_3 | 0.3433 | 1 | 1 | 0.6772 | 1 | 1 | 0.5186 | Annexin protein |
| contig_32749_4 | 0.3459 | 1 | 1 | 0.6815 | 1 | 1 | 0.5218 | leu phe-trna protein transferase |
| contig_39759_2 | 0.3473 | 1 | 1 | 0.6816 | 1 | 1 | 0.522 | antibiotic biosynthesis monooxygenase |
| contig_28322_4 | 0.348 | 1 | 1 | 0.6816 | 1 | 1 | 0.522 | farnesyl pyrophosphate synthase |

| | | | | | | | | |
|---|---|---|---|---|---|---|---|---|
| contig_33495_6 | 0.3481 | 1 | 1 | 0.6816 | 1 | 1 | 0.522 | triose or hexose phosphate/phosphate translocator |
| contig_7935_3 | 0.3484 | 1 | 1 | 0.6816 | 1 | 1 | 0.522 | predicted protein |
| contig_25050_4 | 0.3486 | 1 | 1 | 0.6816 | 1 | 1 | 0.522 | predicted protein |
| contig_25717_4 | 0.3487 | 1 | 1 | 0.6816 | 1 | 1 | 0.522 | ATP-binding Cassette (ABC) Superfamily |
| contig_22528_2 | 0.3498 | 1 | 1 | 0.6816 | 1 | 1 | 0.522 | predicted protein |
| contig_25014_3 | 0.3498 | 1 | 1 | 0.6816 | 1 | 1 | 0.522 | predicted protein |
| contig_34219_5 | 0.3507 | 1 | 1 | 0.6816 | 1 | 1 | 0.522 | Two component regulator three Y domain protein |
| contig_23787_5 | 0.3509 | 1 | 1 | 0.6816 | 1 | 1 | 0.522 | predicted protein |
| contig_11702_1 | 0.3518 | 1 | 1 | 0.6816 | 1 | 1 | 0.522 | glutamine synthetase catalytic domain protein protein |
| contig_24939_3 | 0.3523 | 1 | 1 | 0.6816 | 1 | 1 | 0.522 | beta-ketoacyl-[acyl-carrier-protein] synthase II |
| contig_22940_5 | 0.3526 | 1 | 1 | 0.6816 | 1 | 1 | 0.522 | PH domain and leucine-rich repeat-containing protein phosphatase |
| contig_31795_6 | 0.353 | 1 | 1 | 0.6816 | 1 | 1 | 0.522 | glyceraldehyde-3-phosphate dehydrogenase |
| contig_30834_3 | 0.3537 | 1 | 1 | 0.6816 | 1 | 1 | 0.522 | phosphoglycerate kinase |
| contig_24815_3 | 0.3537 | 1 | 1 | 0.6816 | 1 | 1 | 0.522 | MFS transporter |
| contig_10657_4 | 0.3546 | 1 | 1 | 0.6816 | 1 | 1 | 0.522 | atp-dependent RNA helicase |
| contig_11114_6 | 0.3546 | 1 | 1 | 0.6816 | 1 | 1 | 0.522 | predicted protein |
| contig_10583_5 | 0.3549 | 1 | 1 | 0.6816 | 1 | 1 | 0.522 | photosystem II stability/assembly factor HCF136 |
| contig_40835_6 | 0.3549 | 1 | 1 | 0.6816 | 1 | 1 | 0.522 | 60S ribosomal protein L35a-4 |
| contig_7031_3 | 0.3562 | 1 | 1 | 0.6816 | 1 | 1 | 0.522 | transmembrane 9 superfamily member 3 |
| contig_23548_5 | 0.3568 | 1 | 1 | 0.6816 | 1 | 1 | 0.522 | ATP synthase subunit beta mitochondrial |
| contig_22539_3 | 0.357 | 1 | 1 | 0.6816 | 1 | 1 | 0.522 | sulfate adenylyltransferase |
| contig_29387_4 | 0.3573 | 1 | 1 | 0.6816 | 1 | 1 | 0.522 | predicted protein |
| contig_8557_5 | 0.3575 | 1 | 1 | 0.6816 | 1 | 1 | 0.522 | cytochrome b6 (chloroplast) |
| contig_36079_3 | 0.3575 | 1 | 1 | 0.6816 | 1 | 1 | 0.522 | photosystem II PsbU protein |
| contig_29395_5 | 0.358 | 1 | 1 | 0.6818 | 1 | 1 | 0.5221 | triosephosphate isomerase |
| contig_9038_6 | 0.3606 | 1 | 1 | 0.6858 | 1 | 1 | 0.5252 | methionine-tRNA ligase beta subunit |
| contig_23416_1 | 0.3611 | 1 | 1 | 0.6859 | 1 | 1 | 0.5253 | predicted protein |

| | | | | | | | | |
|---|---|---|---|---|---|---|---|---|
| contig_26787_4 | 0.3618 | 1 | 1 | 0.6863 | 1 | 1 | 0.5255 | DUF4336 domain-containing protein |
| contig_7371_4 | 0.3624 | 1 | 1 | 0.6867 | 1 | 1 | 0.5258 | Lipocalin family protein |
| contig_19585_5 | 0.3638 | 1 | 1 | 0.6884 | 1 | 1 | 0.5271 | ---NA--- |
| contig_25515_1 | 0.3643 | 1 | 1 | 0.6884 | 1 | 1 | 0.5272 | cysteine desulfurase |
| contig_38355_2 | 0.3664 | 1 | 1 | 0.6909 | 1 | 1 | 0.5291 | predicted protein |
| contig_28795_5 | 0.3665 | 1 | 1 | 0.6909 | 1 | 1 | 0.5291 | protein transport protein SEC23 |
| contig_28745_4 | 0.3691 | 1 | 1 | 0.695 | 1 | 1 | 0.5322 | glutamyl-trna reductase |
| contig_23664_6 | 0.3696 | 1 | 1 | 0.695 | 1 | 1 | 0.5322 | enoyl-acp reductase |
| contig_37567_5 | 0.3713 | 1 | 1 | 0.6974 | 1 | 1 | 0.534 | ATP synthase gamma subunit |
| contig_35608_4 | 0.3721 | 1 | 1 | 0.6974 | 1 | 1 | 0.534 | chaperone dnaj-like protein |
| contig_5809_4 | 0.3724 | 1 | 1 | 0.6974 | 1 | 1 | 0.534 | ---NA--- |
| contig_13388_5 | 0.3728 | 1 | 1 | 0.6974 | 1 | 1 | 0.534 | ornithine cyclodeaminase |
| contig_10238_6 | 0.3733 | 1 | 1 | 0.6974 | 1 | 1 | 0.534 | ras-like gtp-binding protein ypt1 |
| contig_6178_5 | 0.3741 | 1 | 1 | 0.6974 | 1 | 1 | 0.534 | rubisco expression protein (chloroplast) |
| contig_30185_1 | 0.3744 | 1 | 1 | 0.6974 | 1 | 1 | 0.534 | signal peptidase I |
| contig_38910_1 | 0.3745 | 1 | 1 | 0.6974 | 1 | 1 | 0.534 | predicted protein |
| contig_8232_1 | 0.3759 | 1 | 1 | 0.6984 | 1 | 1 | 0.5348 | alanine--glyoxylate aminotransferase family protein |
| contig_27824_2 | 0.3764 | 1 | 1 | 0.6984 | 1 | 1 | 0.5348 | protein phosphatase 2C putative |
| contig_17202_2 | 0.3779 | 1 | 1 | 0.6984 | 1 | 1 | 0.5348 | predicted protein |
| contig_40753_3 | 0.3788 | 1 | 1 | 0.6984 | 1 | 1 | 0.5348 | V-type proton ATPase subunit E |
| contig_17744_4 | 0.3793 | 1 | 1 | 0.6984 | 1 | 1 | 0.5348 | integral membrane protein |
| contig_40248_2 | 0.3794 | 1 | 1 | 0.6984 | 1 | 1 | 0.5348 | predicted protein |
| contig_5318_4 | 0.3801 | 1 | 1 | 0.6984 | 1 | 1 | 0.5348 | 40S ribosomal protein S15a |
| contig_30562_5 | 0.3804 | 1 | 1 | 0.6984 | 1 | 1 | 0.5348 | cytochrome b6-f complex iron-sulfur subunit |
| contig_17447_3 | 0.3804 | 1 | 1 | 0.6984 | 1 | 1 | 0.5348 | tetrapyrrole-binding protein |
| contig_6884_5 | 0.3805 | 1 | 1 | 0.6984 | 1 | 1 | 0.5348 | putative ATP-dependent RNA helicase DHX30 isoform X3 |
| contig_14395_5 | 0.3808 | 1 | 1 | 0.6984 | 1 | 1 | 0.5348 | manganese superoxide dismutase |

| | | | | | | | | |
|---|---|---|---|---|---|---|---|---|
| contig_40922_ 3 | 0.3809 | 1 | 1 | 0.69 84 | 1 | 1 | 0.53 48 | V-type H-ATPase subunit C |
| contig_27330_ 6 | 0.3813 | 1 | 1 | 0.69 84 | 1 | 1 | 0.53 48 | predicted protein |
| contig_30838_ 5 | 0.3814 | 1 | 1 | 0.69 84 | 1 | 1 | 0.53 48 | histone H4 |
| contig_21965_ 5 | 0.382 | 1 | 1 | 0.69 87 | 1 | 1 | 0.53 5 | predicted protein |
| contig_27983_ 4 | 0.3845 | 1 | 1 | 0.70 24 | 1 | 1 | 0.53 79 | alpha/beta hydrolase |
| contig_30832_ 6 | 0.3857 | 1 | 1 | 0.70 29 | 1 | 1 | 0.53 82 | predicted protein |
| contig_40270_ 3 | 0.3862 | 1 | 1 | 0.70 29 | 1 | 1 | 0.53 82 | predicted protein |
| contig_10321_ 2 | 0.3866 | 1 | 1 | 0.70 29 | 1 | 1 | 0.53 82 | beta-hydroxyacyl-acyl carrier protein dehydratase (FABZ) |
| contig_26693_ 5 | 0.3866 | 1 | 1 | 0.70 29 | 1 | 1 | 0.53 82 | UMP-CMP kinase |
| contig_16920_ 6 | 0.3877 | 1 | 1 | 0.70 39 | 1 | 1 | 0.53 91 | predicted protein |
| contig_29898_ 4 | 0.3894 | 1 | 1 | 0.70 56 | 1 | 1 | 0.54 03 | mitochondrial 2-oxoglutarate/malate carrier protein |
| contig_28484_ 6 | 0.3911 | 1 | 1 | 0.70 56 | 1 | 1 | 0.54 03 | carbamoyl-phosphate synthase (glutamine-hydrolyzing) large subunit |
| contig_27823_ 2 | 0.3923 | 1 | 1 | 0.70 56 | 1 | 1 | 0.54 03 | predicted protein |
| contig_6911_6 | 0.3924 | 1 | 1 | 0.70 56 | 1 | 1 | 0.54 03 | pyrophosphate-dependent phosphofructose kinase |
| contig_29621_ 3 | 0.3924 | 1 | 1 | 0.70 56 | 1 | 1 | 0.54 03 | predicted protein |
| contig_11202_ 6 | 0.3924 | 1 | 1 | 0.70 56 | 1 | 1 | 0.54 03 | predicted protein |
| contig_20045_ 5 | 0.3925 | 1 | 1 | 0.70 56 | 1 | 1 | 0.54 03 | AP-1 complex subunit sigma-1 |
| contig_27039_ 5 | 0.3926 | 1 | 1 | 0.70 56 | 1 | 1 | 0.54 03 | hypothetical protein THAOC_36140 |
| contig_33729_ 4 | 0.3927 | 1 | 1 | 0.70 56 | 1 | 1 | 0.54 03 | secreted protein |
| contig_15354_ 3 | 0.3953 | 1 | 1 | 0.70 94 | 1 | 1 | 0.54 32 | predicted protein |
| contig_24876_ 6 | 0.3961 | 1 | 1 | 0.70 98 | 1 | 1 | 0.54 35 | predicted protein |
| contig_22476_ 4 | 0.3964 | 1 | 1 | 0.70 98 | 1 | 1 | 0.54 35 | predicted protein |
| contig_28714_ 6 | 0.3969 | 1 | 1 | 0.70 98 | 1 | 1 | 0.54 35 | predicted protein |
| contig_19141_ 5 | 0.3984 | 1 | 1 | 0.71 1 | 1 | 1 | 0.54 44 | photosystem I subunit IV (chloroplast) |
| contig_6572_6 | 0.3985 | 1 | 1 | 0.71 1 | 1 | 1 | 0.54 44 | catalase |
| contig_10170_ 1 | 0.4013 | 1 | 1 | 0.71 52 | 1 | 1 | 0.54 76 | predicted protein |
| contig_23961_ 3 | 0.4024 | 1 | 1 | 0.71 61 | 1 | 1 | 0.54 84 | ribosomal protein S15 |

| | | | | | | | | |
|---|---|---|---|---|---|---|---|---|
| contig_5545_2 | 0.4028 | 1 | 1 | 0.7161 | 1 | 1 | 0.5484 | NarK/NasA family nitrate transporter |
| contig_6155_1 | 0.4035 | 1 | 1 | 0.7166 | 1 | 1 | 0.5487 | light harvesting protein |
| contig_28789_2 | 0.4051 | 1 | 1 | 0.7181 | 1 | 1 | 0.5499 | predicted protein |
| contig_18485_3 | 0.4055 | 1 | 1 | 0.7181 | 1 | 1 | 0.5499 | Mitochondrial-processing peptidase subunit beta |
| contig_17027_5 | 0.4058 | 1 | 1 | 0.7181 | 1 | 1 | 0.5499 | predicted protein |
| contig_10817_6 | 0.4083 | 1 | 1 | 0.7215 | 1 | 1 | 0.5525 | silicon efflux transporter LSI2 |
| contig_23751_6 | 0.4086 | 1 | 1 | 0.7215 | 1 | 1 | 0.5525 | high mobility group protein B2 |
| contig_38903_2 | 0.4106 | 1 | 1 | 0.7238 | 1 | 1 | 0.5542 | ---NA--- |
| contig_15379_5 | 0.4116 | 1 | 1 | 0.7238 | 1 | 1 | 0.5542 | nucleolar gtp-binding protein 1 |
| contig_9719_3 | 0.4116 | 1 | 1 | 0.7238 | 1 | 1 | 0.5542 | Co2+/Mg2+ efflux protein ApaG |
| contig_18489_1 | 0.4118 | 1 | 1 | 0.7238 | 1 | 1 | 0.5542 | predicted protein |
| contig_40197_2 | 0.4123 | 1 | 1 | 0.7238 | 1 | 1 | 0.5542 | protein fucoxanthin chlorophyll a/c protein |
| contig_6242_5 | 0.4139 | 1 | 1 | 0.7257 | 1 | 1 | 0.5557 | predicted protein |
| contig_5949_4 | 0.4148 | 1 | 1 | 0.7264 | 1 | 1 | 0.5563 | predicted protein |
| contig_13878_2 | 0.4159 | 1 | 1 | 0.7267 | 1 | 1 | 0.5565 | glycine--tRNA ligase |
| contig_22744_5 | 0.4161 | 1 | 1 | 0.7267 | 1 | 1 | 0.5565 | predicted protein |
| contig_22418_5 | 0.4164 | 1 | 1 | 0.7267 | 1 | 1 | 0.5565 | protein fucoxanthin chlorophyll a/c protein |
| contig_28112_2 | 0.4174 | 1 | 1 | 0.7267 | 1 | 1 | 0.5565 | predicted protein |
| contig_31306_6 | 0.4188 | 1 | 1 | 0.7267 | 1 | 1 | 0.5565 | predicted protein |
| contig_33904_2 | 0.419 | 1 | 1 | 0.7267 | 1 | 1 | 0.5565 | pectate lyase |
| contig_17153_5 | 0.4197 | 1 | 1 | 0.7267 | 1 | 1 | 0.5565 | predicted protein |
| contig_34331_5 | 0.4199 | 1 | 1 | 0.7267 | 1 | 1 | 0.5565 | Light-harvesting complex I LH38 protein |
| contig_29871_5 | 0.42 | 1 | 1 | 0.7267 | 1 | 1 | 0.5565 | lipoprotein type 6 |
| contig_24829_2 | 0.4202 | 1 | 1 | 0.7267 | 1 | 1 | 0.5565 | ornithine carbamoyltransferase |
| contig_30325_5 | 0.4202 | 1 | 1 | 0.7267 | 1 | 1 | 0.5565 | predicted protein |
| contig_38550_1 | 0.4249 | 1 | 1 | 0.7292 | 1 | 1 | 0.5584 | polyadenylate-binding protein 1-B |
| contig_27279_6 | 0.4252 | 1 | 1 | 0.7292 | 1 | 1 | 0.5584 | adenosine kinase |

| | | | | | | | | |
|---|---|---|---|---|---|---|---|---|
| contig_5617_6 | 0.4255 | 1 | 1 | 0.72 92 | 1 | 1 | 0.55 84 | sarcoplasmic calcium-binding protein |
| contig_24221_5 | 0.4258 | 1 | 1 | 0.72 92 | 1 | 1 | 0.55 84 | ATP-binding cassette superfamily |
| contig_29943_6 | 0.4279 | 1 | 1 | 0.72 92 | 1 | 1 | 0.55 84 | Got1-domain-containing protein |
| contig_26174_3 | 0.428 | 1 | 1 | 0.72 92 | 1 | 1 | 0.55 84 | predicted protein |
| contig_7261_5 | 0.4282 | 1 | 1 | 0.72 92 | 1 | 1 | 0.55 84 | ubiquitin-conjugating enzyme E2 2 |
| contig_34843_5 | 0.4287 | 1 | 1 | 0.72 92 | 1 | 1 | 0.55 84 | Major Facilitator Superfamily (MFS) |
| contig_28302_4 | 0.4288 | 1 | 1 | 0.72 92 | 1 | 1 | 0.55 84 | predicted protein |
| contig_26157_4 | 0.4288 | 1 | 1 | 0.72 92 | 1 | 1 | 0.55 84 | predicted protein |
| contig_30540_5 | 0.4289 | 1 | 1 | 0.72 92 | 1 | 1 | 0.55 84 | unnamed protein product |
| contig_9284_2 | 0.4289 | 1 | 1 | 0.72 92 | 1 | 1 | 0.55 84 | 60S ribosomal protein L12 |
| contig_40960_3 | 0.4291 | 1 | 1 | 0.72 92 | 1 | 1 | 0.55 84 | predicted protein |
| contig_14595_4 | 0.4297 | 1 | 1 | 0.72 92 | 1 | 1 | 0.55 84 | predicted protein |
| contig_13885_3 | 0.4298 | 1 | 1 | 0.72 92 | 1 | 1 | 0.55 84 | predicted protein |
| contig_38515_1 | 0.4298 | 1 | 1 | 0.72 92 | 1 | 1 | 0.55 84 | predicted protein |
| contig_29967_5 | 0.4302 | 1 | 1 | 0.72 92 | 1 | 1 | 0.55 84 | protein dispatched homolog 1 isoform X2 |
| contig_27614_3 | 0.4306 | 1 | 1 | 0.72 92 | 1 | 1 | 0.55 84 | Regulator of nonsense transcripts 1-like protein |
| contig_5862_2 | 0.4306 | 1 | 1 | 0.72 92 | 1 | 1 | 0.55 84 | arylsulfatase |
| contig_28340_4 | 0.4318 | 1 | 1 | 0.73 03 | 1 | 1 | 0.55 92 | Ras-like protein Rab-11B |
| contig_9617_4 | 0.4331 | 1 | 1 | 0.73 05 | 1 | 1 | 0.55 94 | bifunctional methylenetetrahydrofolate dehydrogenase/methenyltetrahydrofolate cyclohydrolase FolD |
| contig_24606_4 | 0.4332 | 1 | 1 | 0.73 05 | 1 | 1 | 0.55 94 | beta-ketoacyl-ACP synthase I |
| contig_33515_6 | 0.4333 | 1 | 1 | 0.73 05 | 1 | 1 | 0.55 94 | predicted protein |
| contig_26792_2 | 0.4354 | 1 | 1 | 0.73 31 | 1 | 1 | 0.56 14 | predicted protein |
| contig_12155_4 | 0.4363 | 1 | 1 | 0.73 31 | 1 | 1 | 0.56 14 | predicted protein |
| contig_30723_5 | 0.4364 | 1 | 1 | 0.73 31 | 1 | 1 | 0.56 14 | 20S proteasome subunit beta 1 |
| contig_39202_4 | 0.4369 | 1 | 1 | 0.73 31 | 1 | 1 | 0.56 14 | Bystin |
| contig_32739_1 | 0.4373 | 1 | 1 | 0.73 31 | 1 | 1 | 0.56 14 | fibronectin type III domain-containing protein |

| | | | | | | | | |
|---|---|---|---|---|---|---|---|---|
| contig_36049_<br>1 | 0.4398 | 1 | 1 | 0.73<br>55 | 1 | 1 | 0.56<br>32 | predicted protein |
| contig_29914_<br>2 | 0.4401 | 1 | 1 | 0.73<br>55 | 1 | 1 | 0.56<br>32 | predicted protein |
| contig_33347_<br>4 | 0.4402 | 1 | 1 | 0.73<br>55 | 1 | 1 | 0.56<br>32 | V-type ATPase A subunit |
| contig_17493_<br>4 | 0.4421 | 1 | 1 | 0.73<br>72 | 1 | 1 | 0.56<br>45 | lamin tail domain-containing protein |
| contig_6257_1 | 0.4431 | 1 | 1 | 0.73<br>72 | 1 | 1 | 0.56<br>45 | predicted protein |
| contig_27301_<br>5 | 0.4432 | 1 | 1 | 0.73<br>72 | 1 | 1 | 0.56<br>45 | enolase |
| contig_28721_<br>5 | 0.4435 | 1 | 1 | 0.73<br>72 | 1 | 1 | 0.56<br>45 | elongation factor Tu |
| contig_38019_<br>3 | 0.4438 | 1 | 1 | 0.73<br>72 | 1 | 1 | 0.56<br>45 | ferredoxin--NADP+ reductase |
| contig_10116_<br>3 | 0.4441 | 1 | 1 | 0.73<br>72 | 1 | 1 | 0.56<br>45 | peptidyl-prolyl cis-trans isomerase CYP19-4 |
| contig_31150_<br>6 | 0.4451 | 1 | 1 | 0.73<br>79 | 1 | 1 | 0.56<br>51 | protein fucoxanthin chlorophyll a/c protein |
| contig_30519_<br>2 | 0.4459 | 1 | 1 | 0.73<br>79 | 1 | 1 | 0.56<br>51 | predicted protein |
| contig_27168_<br>5 | 0.4471 | 1 | 1 | 0.73<br>79 | 1 | 1 | 0.56<br>51 | predicted protein |
| contig_5475_2 | 0.4475 | 1 | 1 | 0.73<br>79 | 1 | 1 | 0.56<br>51 | coatomer subunit epsilon |
| contig_34472_<br>2 | 0.4477 | 1 | 1 | 0.73<br>79 | 1 | 1 | 0.56<br>51 | histone h4 |
| contig_8751_4 | 0.4477 | 1 | 1 | 0.73<br>79 | 1 | 1 | 0.56<br>51 | predicted protein |
| contig_35155_<br>5 | 0.4479 | 1 | 1 | 0.73<br>79 | 1 | 1 | 0.56<br>51 | fucoxanthin chlorophyll a/c protein 8 |
| contig_16696_<br>4 | 0.4488 | 1 | 1 | 0.73<br>79 | 1 | 1 | 0.56<br>51 | pyruvate kinase |
| contig_5529_5 | 0.4495 | 1 | 1 | 0.73<br>79 | 1 | 1 | 0.56<br>51 | photosystem II protein D1 (chloroplast) |
| contig_6058_5 | 0.4495 | 1 | 1 | 0.73<br>79 | 1 | 1 | 0.56<br>51 | predicted protein |
| contig_35648_<br>5 | 0.4498 | 1 | 1 | 0.73<br>79 | 1 | 1 | 0.56<br>51 | ubiquitin / ribosomal protein CEP52 fusion protein putative |
| contig_22590_<br>5 | 0.4504 | 1 | 1 | 0.73<br>79 | 1 | 1 | 0.56<br>51 | glutathione reductase |
| contig_12035_<br>1 | 0.4508 | 1 | 1 | 0.73<br>79 | 1 | 1 | 0.56<br>51 | predicted protein |
| contig_11813_<br>1 | 0.4521 | 1 | 1 | 0.73<br>94 | 1 | 1 | 0.56<br>62 | predicted protein |
| contig_17128_<br>2 | 0.4527 | 1 | 1 | 0.73<br>95 | 1 | 1 | 0.56<br>63 | predicted protein |
| contig_27171_<br>6 | 0.4547 | 1 | 1 | 0.74<br>19 | 1 | 1 | 0.56<br>81 | ribosome production factor 2 homolog |
| contig_10857_<br>4 | 0.4559 | 1 | 1 | 0.74<br>19 | 1 | 1 | 0.56<br>81 | predicted protein |
| contig_10223_<br>6 | 0.456 | 1 | 1 | 0.74<br>19 | 1 | 1 | 0.56<br>81 | predicted protein |

| | | | | | | | | |
|---|---|---|---|---|---|---|---|---|
| contig_15137_6 | 0.4561 | 1 | 1 | 0.7419 | 1 | 1 | 0.5681 | aspartate aminotransferase |
| contig_38239_2 | 0.4569 | 1 | 1 | 0.7424 | 1 | 1 | 0.5685 | predicted protein |
| contig_33419_4 | 0.4576 | 1 | 1 | 0.7427 | 1 | 1 | 0.5687 | light harvesting protein |
| contig_21525_6 | 0.4585 | 1 | 1 | 0.7434 | 1 | 1 | 0.5693 | probable small nuclear ribonucleoprotein Sm D2 |
| contig_32611_6 | 0.4615 | 1 | 1 | 0.7466 | 1 | 1 | 0.5717 | predicted protein |
| contig_25890_5 | 0.4619 | 1 | 1 | 0.7466 | 1 | 1 | 0.5717 | acetolactate synthase catalytic subunit |
| contig_27634_6 | 0.4621 | 1 | 1 | 0.7466 | 1 | 1 | 0.5717 | eukaryotic translation initiation factor 3 subunit E |
| contig_6246_4 | 0.4624 | 1 | 1 | 0.7466 | 1 | 1 | 0.5717 | predicted protein |
| contig_9488_1 | 0.4634 | 1 | 1 | 0.7474 | 1 | 1 | 0.5723 | 26s protease regulatory subunit 8 |
| contig_24818_1 | 0.4641 | 1 | 1 | 0.7477 | 1 | 1 | 0.5726 | UBC-like protein |
| contig_7217_3 | 0.4654 | 1 | 1 | 0.749 | 1 | 1 | 0.5736 | P-loop containing nucleoside triphosphate hydrolase protein |
| contig_21707_3 | 0.4659 | 1 | 1 | 0.749 | 1 | 1 | 0.5736 | 40S ribosomal protein S4 |
| contig_8596_6 | 0.4665 | 1 | 1 | 0.749 | 1 | 1 | 0.5736 | predicted protein |
| contig_22809_5 | 0.4674 | 1 | 1 | 0.749 | 1 | 1 | 0.5736 | predicted protein |
| contig_38406_6 | 0.4676 | 1 | 1 | 0.749 | 1 | 1 | 0.5736 | predicted protein |
| contig_24978_5 | 0.4682 | 1 | 1 | 0.749 | 1 | 1 | 0.5736 | predicted protein |
| contig_28787_6 | 0.4683 | 1 | 1 | 0.749 | 1 | 1 | 0.5736 | tryptophan synthase |
| contig_5402_3 | 0.4691 | 1 | 1 | 0.7495 | 1 | 1 | 0.5739 | NAD(P)-dependent oxidoreductase |
| contig_24783_5 | 0.47 | 1 | 1 | 0.7501 | 1 | 1 | 0.5744 | mitochondrial protein translocase MPT family |
| contig_38543_1 | 0.4717 | 1 | 1 | 0.7511 | 1 | 1 | 0.5751 | predicted protein |
| contig_28186_6 | 0.4725 | 1 | 1 | 0.7511 | 1 | 1 | 0.5751 | predicted protein |
| contig_30481_5 | 0.4735 | 1 | 1 | 0.7511 | 1 | 1 | 0.5751 | ornithine cyclodeaminase |
| contig_23660_5 | 0.4736 | 1 | 1 | 0.7511 | 1 | 1 | 0.5751 | predicted protein |
| contig_26114_2 | 0.4738 | 1 | 1 | 0.7511 | 1 | 1 | 0.5751 | ATP phosphoribosyltransferase |
| contig_7732_1 | 0.4739 | 1 | 1 | 0.7511 | 1 | 1 | 0.5751 | predicted protein |
| contig_14363_5 | 0.4741 | 1 | 1 | 0.7511 | 1 | 1 | 0.5751 | predicted protein |
| contig_27030_2 | 0.4745 | 1 | 1 | 0.7511 | 1 | 1 | 0.5751 | geranylgeranyl reductase |

| | | | | | | | | |
|---|---|---|---|---|---|---|---|---|
| contig_30747_4 | 0.4756 | 1 | 1 | 0.7514 | 1 | 1 | 0.5754 | bile acid:Na+ symporter BASS family |
| contig_35157_2 | 0.4757 | 1 | 1 | 0.7514 | 1 | 1 | 0.5754 | predicted protein |
| contig_5001_1 | 0.478 | 1 | 1 | 0.7536 | 1 | 1 | 0.5771 | membrane guanylyl cyclase |
| contig_30385_5 | 0.4781 | 1 | 1 | 0.7536 | 1 | 1 | 0.5771 | myo-inositol-1(or 4)-monophosphatase |
| contig_32718_6 | 0.4788 | 1 | 1 | 0.754 | 1 | 1 | 0.5774 | aspartate-semialdehyde dehydrogenase |
| contig_9753_3 | 0.4813 | 1 | 1 | 0.7565 | 1 | 1 | 0.5793 | predicted protein |
| contig_33218_4 | 0.4815 | 1 | 1 | 0.7565 | 1 | 1 | 0.5793 | predicted protein |
| contig_29776_5 | 0.4819 | 1 | 1 | 0.7565 | 1 | 1 | 0.5793 | fructose-bisphosphate aldolase |
| contig_9464_1 | 0.4829 | 1 | 1 | 0.7573 | 1 | 1 | 0.5799 | cytochrome c heme attachment protein (chloroplast) |
| contig_5961_5 | 0.4835 | 1 | 1 | 0.7575 | 1 | 1 | 0.5801 | cytochrome oxidase subunit III (mitochondrion) |
| contig_20174_5 | 0.4845 | 1 | 1 | 0.7581 | 1 | 1 | 0.5805 | glutathione S-transferase |
| contig_8378_3 | 0.4856 | 1 | 1 | 0.7581 | 1 | 1 | 0.5805 | phosphoglyceromutase |
| contig_24888_1 | 0.486 | 1 | 1 | 0.7581 | 1 | 1 | 0.5805 | chromodomain-helicase-DNA-binding protein 7 |
| contig_21612_4 | 0.4877 | 1 | 1 | 0.7581 | 1 | 1 | 0.5805 | 60S ribosomal protein L27 |
| contig_30735_2 | 0.4877 | 1 | 1 | 0.7581 | 1 | 1 | 0.5805 | myo-inositol 2-dehydrogenase putative |
| contig_5987_6 | 0.4878 | 1 | 1 | 0.7581 | 1 | 1 | 0.5805 | predicted protein |
| contig_17529_2 | 0.4878 | 1 | 1 | 0.7581 | 1 | 1 | 0.5805 | ribose-phosphate pyrophosphokinase |
| contig_25825_6 | 0.488 | 1 | 1 | 0.7581 | 1 | 1 | 0.5805 | predicted protein |
| contig_40765_3 | 0.4884 | 1 | 1 | 0.7581 | 1 | 1 | 0.5805 | triosephosphate isomerase/glyceraldehyde-3-phosphate dehydrogenase precursor |
| contig_25375_6 | 0.4894 | 1 | 1 | 0.7587 | 1 | 1 | 0.581 | predicted protein |
| contig_6766_1 | 0.4898 | 1 | 1 | 0.7587 | 1 | 1 | 0.581 | N-carbamoylputrescine amidase |
| contig_30582_1 | 0.4907 | 1 | 1 | 0.7591 | 1 | 1 | 0.5813 | silicon transporter |
| contig_8461_2 | 0.491 | 1 | 1 | 0.7591 | 1 | 1 | 0.5813 | SDR family oxidoreductase |
| contig_12505_5 | 0.4917 | 1 | 1 | 0.7594 | 1 | 1 | 0.5815 | threonine ammonia-lyase |
| contig_18571_6 | 0.4925 | 1 | 1 | 0.7598 | 1 | 1 | 0.5818 | phosphoribulokinase |
| contig_6146_1 | 0.4933 | 1 | 1 | 0.7598 | 1 | 1 | 0.5818 | aliphatic amidase |

| | | | | | | | | |
|---|---|---|---|---|---|---|---|---|
| contig_30338_1 | 0.4935 | 1 | 1 | 0.7598 | 1 | 1 | 0.5818 | 60S ribosomal protein L5 |
| contig_24923_5 | 0.4943 | 1 | 1 | 0.7603 | 1 | 1 | 0.5822 | voltage activated chloride channel CLC7 type |
| contig_35046_3 | 0.4948 | 1 | 1 | 0.7604 | 1 | 1 | 0.5823 | CK1/CK1/CK1-D protein kinase |
| contig_14496_5 | 0.4977 | 1 | 1 | 0.7632 | 1 | 1 | 0.5844 | Diaminopimelate epimerase |
| contig_11653_4 | 0.4977 | 1 | 1 | 0.7632 | 1 | 1 | 0.5844 | COP9 signalosome complex subunit 3 |
| contig_28061_5 | 0.4982 | 1 | 1 | 0.7632 | 1 | 1 | 0.5844 | apospory-associated protein C |
| contig_33369_2 | 0.5001 | 1 | 1 | 0.7653 | 1 | 1 | 0.5861 | F0F1 ATP synthase subunit gamma |
| contig_26661_4 | 0.502 | 1 | 1 | 0.7676 | 1 | 1 | 0.5878 | photosystem II oxygen evolving complex protein PsbP |
| contig_6009_6 | 0.5034 | 1 | 1 | 0.7689 | 1 | 1 | 0.5888 | 40S ribosomal protein S5 |
| contig_10909_2 | 0.5045 | 1 | 1 | 0.7695 | 1 | 1 | 0.5892 | Glycerol-3-phosphate dehydrogenase mitochondrial |
| contig_23510_1 | 0.5048 | 1 | 1 | 0.7695 | 1 | 1 | 0.5892 | predicted protein |
| contig_11344_5 | 0.507 | 1 | 1 | 0.7717 | 1 | 1 | 0.5909 | Iron-sulfur cluster co-chaperone protein HscB mitochondrial |
| contig_5427_6 | 0.5072 | 1 | 1 | 0.7717 | 1 | 1 | 0.5909 | tyrosyl-tRNA synthetase |
| contig_24322_5 | 0.5087 | 1 | 1 | 0.7732 | 1 | 1 | 0.5921 | ARM repeat superfamily protein |
| contig_27487_4 | 0.5123 | 1 | 1 | 0.7779 | 1 | 1 | 0.5957 | ---NA--- |
| contig_23942_6 | 0.5134 | 1 | 1 | 0.7787 | 1 | 1 | 0.5963 | sister-chromatid cohesion protein 3 |
| contig_22651_1 | 0.5163 | 1 | 1 | 0.7824 | 1 | 1 | 0.5991 | glutamate synthase subunit alpha |
| contig_28383_6 | 0.5184 | 1 | 1 | 0.7848 | 1 | 1 | 0.6009 | predicted protein |
| contig_7315_2 | 0.5214 | 1 | 1 | 0.7885 | 1 | 1 | 0.6038 | predicted protein |
| contig_11002_5 | 0.5224 | 1 | 1 | 0.7885 | 1 | 1 | 0.6038 | peroxiredoxin |
| contig_28073_5 | 0.5225 | 1 | 1 | 0.7885 | 1 | 1 | 0.6038 | NAD-dependent epimerase/dehydratase family protein |
| contig_37256_3 | 0.5233 | 1 | 1 | 0.7885 | 1 | 1 | 0.6038 | histone H3 |
| contig_25649_3 | 0.5237 | 1 | 1 | 0.7885 | 1 | 1 | 0.6038 | tRNA pseudouridine(55) synthase |
| contig_18202_5 | 0.524 | 1 | 1 | 0.7885 | 1 | 1 | 0.6038 | protein fucoxanthin chlorophyll a/c protein |
| contig_29804_5 | 0.5245 | 1 | 1 | 0.7886 | 1 | 1 | 0.6039 | S-adenosylmethionine synthetase |
| contig_14719_6 | 0.5268 | 1 | 1 | 0.7912 | 1 | 1 | 0.6058 | thioredoxin-like |

| | | | | | | | | |
|---|---|---|---|---|---|---|---|---|
| contig_34883_6 | 0.5283 | 1 | 1 | 0.79 22 | 1 | 1 | 0.60 67 | Fructose-bisphosphate aldolase |
| contig_24590_4 | 0.5287 | 1 | 1 | 0.79 22 | 1 | 1 | 0.60 67 | acyl-CoA dehydrogenase NM domain-like protein |
| contig_28743_4 | 0.5294 | 1 | 1 | 0.79 22 | 1 | 1 | 0.60 67 | chlorophyll a/b-binding protein |
| contig_30406_3 | 0.5296 | 1 | 1 | 0.79 22 | 1 | 1 | 0.60 67 | Glycylpeptide N-tetradecanoyltransferase 2 |
| contig_38608_1 | 0.5301 | 1 | 1 | 0.79 22 | 1 | 1 | 0.60 67 | cytochrome 450 |
| contig_30517_5 | 0.5321 | 1 | 1 | 0.79 45 | 1 | 1 | 0.60 84 | predicted protein |
| contig_36065_2 | 0.534 | 1 | 1 | 0.79 56 | 1 | 1 | 0.60 92 | Translation elongation factor IF5A |
| contig_15516_6 | 0.5348 | 1 | 1 | 0.79 56 | 1 | 1 | 0.60 92 | pyrimidine/purine nucleoside phosphorylase |
| contig_34760_6 | 0.5349 | 1 | 1 | 0.79 56 | 1 | 1 | 0.60 92 | low molecular mass early light-inducible protein HV60 |
| contig_8557_6 | 0.535 | 1 | 1 | 0.79 56 | 1 | 1 | 0.60 92 | cytochrome b6 (chloroplast) |
| contig_6105_6 | 0.5357 | 1 | 1 | 0.79 56 | 1 | 1 | 0.60 92 | photosystem II CP47 chlorophyll apoprotein (chloroplast) |
| contig_24050_3 | 0.5359 | 1 | 1 | 0.79 56 | 1 | 1 | 0.60 92 | enolase-phosphatase E1 |
| contig_24297_3 | 0.5372 | 1 | 1 | 0.79 63 | 1 | 1 | 0.60 98 | elongation factor Tu |
| contig_39738_3 | 0.5377 | 1 | 1 | 0.79 63 | 1 | 1 | 0.60 98 | Asparaginyl-tRNA synthetase |
| contig_25987_5 | 0.538 | 1 | 1 | 0.79 63 | 1 | 1 | 0.60 98 | polyadenylate-binding protein 1-B |
| contig_16388_6 | 0.5388 | 1 | 1 | 0.79 67 | 1 | 1 | 0.61 01 | predicted protein |
| contig_28948_1 | 0.5397 | 1 | 1 | 0.79 73 | 1 | 1 | 0.61 06 | geranylgeranyl reductase |
| contig_23256_4 | 0.5405 | 1 | 1 | 0.79 78 | 1 | 1 | 0.61 09 | fumarate hydratase mitochondrial |
| contig_29986_5 | 0.5419 | 1 | 1 | 0.79 78 | 1 | 1 | 0.61 09 | H - or Na -translocating F-type V-type and A-type ATPase (F-ATPase) Superfamily |
| contig_35414_3 | 0.542 | 1 | 1 | 0.79 78 | 1 | 1 | 0.61 09 | ATP synthase subunit alpha mitochondrial |
| contig_33151_3 | 0.5427 | 1 | 1 | 0.79 78 | 1 | 1 | 0.61 09 | acetylglutamate kinase |
| contig_41371_1 | 0.5436 | 1 | 1 | 0.79 78 | 1 | 1 | 0.61 09 | 40s ribosomal protein |
| contig_41222_3 | 0.5441 | 1 | 1 | 0.79 78 | 1 | 1 | 0.61 09 | predicted protein |
| contig_27523_3 | 0.5445 | 1 | 1 | 0.79 78 | 1 | 1 | 0.61 09 | plastid protoporphyrinogen oxidase |
| contig_12042_4 | 0.5446 | 1 | 1 | 0.79 78 | 1 | 1 | 0.61 09 | coproporphyrinogen III oxidase |
| contig_40265_2 | 0.5453 | 1 | 1 | 0.79 78 | 1 | 1 | 0.61 09 | predicted protein |

| | | | | | | | | |
|---|---|---|---|---|---|---|---|---|
| contig_7234_4 | 0.5453 | 1 | 1 | 0.7978 | 1 | 1 | 0.6109 | ras-related gtp-binding protein |
| contig_16696_5 | 0.5464 | 1 | 1 | 0.798 | 1 | 1 | 0.6111 | pyruvate kinase |
| contig_23306_1 | 0.5464 | 1 | 1 | 0.798 | 1 | 1 | 0.6111 | ATP:ADP antiporter AAA family |
| contig_30830_5 | 0.5476 | 1 | 1 | 0.7988 | 1 | 1 | 0.6117 | PGR5-like protein 1A chloroplastic |
| contig_26815_5 | 0.548 | 1 | 1 | 0.7988 | 1 | 1 | 0.6117 | AGC/PKA protein kinase |
| contig_3491_1 | 0.5486 | 1 | 1 | 0.7989 | 1 | 1 | 0.6118 | predicted protein |
| contig_24983_1 | 0.5493 | 1 | 1 | 0.7991 | 1 | 1 | 0.6119 | GTP-binding nuclear protein Ran |
| contig_8945_4 | 0.5498 | 1 | 1 | 0.7991 | 1 | 1 | 0.6119 | delta-9 acyl-desaturase |
| contig_27608_2 | 0.5519 | 1 | 1 | 0.8007 | 1 | 1 | 0.6131 | lysine-tRNA ligase |
| contig_21568_5 | 0.5519 | 1 | 1 | 0.8007 | 1 | 1 | 0.6131 | predicted protein |
| contig_37939_4 | 0.5564 | 1 | 1 | 0.8055 | 1 | 1 | 0.6168 | predicted protein |
| contig_26948_5 | 0.5568 | 1 | 1 | 0.8055 | 1 | 1 | 0.6168 | predicted protein |
| contig_14363_4 | 0.5573 | 1 | 1 | 0.8055 | 1 | 1 | 0.6168 | predicted protein |
| contig_25763_5 | 0.5581 | 1 | 1 | 0.8055 | 1 | 1 | 0.6168 | Dihydroflavonol 4-reductase |
| contig_22768_6 | 0.5581 | 1 | 1 | 0.8055 | 1 | 1 | 0.6168 | putative LOV domain-containing protein |
| contig_7122_5 | 0.5588 | 1 | 1 | 0.8055 | 1 | 1 | 0.6168 | ---NA--- |
| contig_27187_2 | 0.5594 | 1 | 1 | 0.8055 | 1 | 1 | 0.6168 | CAAX prenyl protease 1 |
| contig_32692_5 | 0.5595 | 1 | 1 | 0.8055 | 1 | 1 | 0.6168 | predicted protein |
| contig_28185_5 | 0.561 | 1 | 1 | 0.8065 | 1 | 1 | 0.6176 | UMP-CMP kinase |
| contig_9857_5 | 0.5614 | 1 | 1 | 0.8065 | 1 | 1 | 0.6176 | ---NA--- |
| contig_35366_6 | 0.5626 | 1 | 1 | 0.8065 | 1 | 1 | 0.6176 | chaperonin GroEL (chloroplast) |
| contig_32515_5 | 0.5631 | 1 | 1 | 0.8065 | 1 | 1 | 0.6176 | phosphoenolpyruvate synthase |
| contig_33655_6 | 0.5638 | 1 | 1 | 0.8065 | 1 | 1 | 0.6176 | predicted protein |
| contig_28068_1 | 0.5647 | 1 | 1 | 0.8065 | 1 | 1 | 0.6176 | UDP-glucose 6-dehydrogenase |
| contig_30141_2 | 0.5648 | 1 | 1 | 0.8065 | 1 | 1 | 0.6176 | Lipoamide acyltransferase component of branched-chain alpha-keto acid dehydrogenase complex mitochondrial |
| contig_15161_3 | 0.5651 | 1 | 1 | 0.8065 | 1 | 1 | 0.6176 | serine/threonine-protein phosphatase PP1-beta catalytic subunit |

| | | | | | | | | |
|---|---|---|---|---|---|---|---|---|
| contig_34246_6 | 0.566 | 1 | 1 | 0.8065 | 1 | 1 | 0.6176 | homoserine kinase |
| contig_9256_6 | 0.5665 | 1 | 1 | 0.8065 | 1 | 1 | 0.6176 | predicted protein |
| contig_30640_5 | 0.567 | 1 | 1 | 0.8065 | 1 | 1 | 0.6176 | predicted protein |
| contig_18936_6 | 0.5674 | 1 | 1 | 0.8065 | 1 | 1 | 0.6176 | 40S ribosomal protein S3a |
| contig_25363_4 | 0.5675 | 1 | 1 | 0.8065 | 1 | 1 | 0.6176 | ABC-transporter like protein |
| contig_33443_2 | 0.568 | 1 | 1 | 0.8065 | 1 | 1 | 0.6176 | predicted protein |
| contig_28453_2 | 0.5681 | 1 | 1 | 0.8065 | 1 | 1 | 0.6176 | methionine-dependent methyltransferase |
| contig_8988_3 | 0.5694 | 1 | 1 | 0.8077 | 1 | 1 | 0.6185 | DUF1295 domain-containing protein |
| contig_10451_3 | 0.5707 | 1 | 1 | 0.808 | 1 | 1 | 0.6188 | predicted protein |
| contig_24490_6 | 0.5708 | 1 | 1 | 0.808 | 1 | 1 | 0.6188 | predicted protein |
| contig_26683_6 | 0.5722 | 1 | 1 | 0.8093 | 1 | 1 | 0.6197 | predicted protein |
| contig_21753_2 | 0.5748 | 1 | 1 | 0.8123 | 1 | 1 | 0.622 | glyceraldehyde-3-phosphate dehydrogenase |
| contig_7125_2 | 0.5769 | 1 | 1 | 0.8132 | 1 | 1 | 0.6227 | cysteine proteinase 3-like |
| contig_37765_6 | 0.577 | 1 | 1 | 0.8132 | 1 | 1 | 0.6227 | serine/threonine-protein phosphatase 2A regulatory subunit B' |
| contig_13099_5 | 0.5771 | 1 | 1 | 0.8132 | 1 | 1 | 0.6227 | predicted protein |
| contig_6012_4 | 0.58 | 1 | 1 | 0.8141 | 1 | 1 | 0.6234 | V-type proton ATPase subunit B isoform |
| contig_18966_2 | 0.5802 | 1 | 1 | 0.8141 | 1 | 1 | 0.6234 | predicted protein |
| contig_28796_6 | 0.5807 | 1 | 1 | 0.8141 | 1 | 1 | 0.6234 | predicted protein |
| contig_37718_5 | 0.5808 | 1 | 1 | 0.8141 | 1 | 1 | 0.6234 | predicted protein |
| contig_39337_5 | 0.5811 | 1 | 1 | 0.8141 | 1 | 1 | 0.6234 | ---NA--- |
| contig_9759_2 | 0.5818 | 1 | 1 | 0.8141 | 1 | 1 | 0.6234 | clathrin heavy chain |
| contig_27953_1 | 0.582 | 1 | 1 | 0.8141 | 1 | 1 | 0.6234 | ribosomal protein L23 putative |
| contig_27705_5 | 0.5823 | 1 | 1 | 0.8141 | 1 | 1 | 0.6234 | predicted protein |
| contig_9532_4 | 0.5825 | 1 | 1 | 0.8141 | 1 | 1 | 0.6234 | NADH dehydrogenase [ubiquinone] iron-sulfur protein 8-A mitochondrial |
| contig_28468_2 | 0.5844 | 1 | 1 | 0.816 | 1 | 1 | 0.6249 | ribosome recycling factor |
| contig_27634_3 | 0.5854 | 1 | 1 | 0.8167 | 1 | 1 | 0.6254 | eukaryotic translation initiation factor 3 subunit E |

| | | | | | | | | |
|---|---|---|---|---|---|---|---|---|
| contig_9605_2 | 0.5888 | 1 | 1 | 0.82 05 | 1 | 1 | 0.62 83 | arginine biosynthesis ArgJ mitochondrial |
| contig_40882_ 2 | 0.5892 | 1 | 1 | 0.82 05 | 1 | 1 | 0.62 83 | predicted protein |
| contig_19474_ 5 | 0.5909 | 1 | 1 | 0.82 21 | 1 | 1 | 0.62 95 | aspartate aminotransferase |
| contig_17853_ 5 | 0.5951 | 1 | 1 | 0.82 72 | 1 | 1 | 0.63 34 | predicted protein |
| contig_28316_ 5 | 0.5969 | 1 | 1 | 0.82 9 | 1 | 1 | 0.63 48 | hypothetical protein FRACYDRAFT_234640 |
| contig_28929_ 5 | 0.5977 | 1 | 1 | 0.82 94 | 1 | 1 | 0.63 51 | 1-deoxy-D-xylulose 5-phosphate reductoisomerase |
| contig_25907_ 5 | 0.5998 | 1 | 1 | 0.83 08 | 1 | 1 | 0.63 62 | predicted protein |
| contig_7643_4 | 0.5998 | 1 | 1 | 0.83 08 | 1 | 1 | 0.63 62 | ATP synthase CF1 beta subunit (chloroplast) |
| contig_24960_ 3 | 0.6008 | 1 | 1 | 0.83 09 | 1 | 1 | 0.63 63 | predicted protein |
| contig_22705_ 1 | 0.6013 | 1 | 1 | 0.83 09 | 1 | 1 | 0.63 63 | glyceraldehyde-3-phosphate dehydrogenase precursor |
| contig_29806_ 5 | 0.6016 | 1 | 1 | 0.83 09 | 1 | 1 | 0.63 63 | predicted protein |
| contig_40101_ 4 | 0.6022 | 1 | 1 | 0.83 1 | 1 | 1 | 0.63 63 | 40S ribosomal protein S13 |
| contig_16580_ 4 | 0.6033 | 1 | 1 | 0.83 17 | 1 | 1 | 0.63 69 | casein kinase I |
| contig_17354_ 1 | 0.6039 | 1 | 1 | 0.83 19 | 1 | 1 | 0.63 7 | T9SS type A sorting domain-containing protein |
| contig_30573_ 3 | 0.605 | 1 | 1 | 0.83 2 | 1 | 1 | 0.63 71 | predicted protein |
| contig_23607_ 2 | 0.6051 | 1 | 1 | 0.83 2 | 1 | 1 | 0.63 71 | alanyl-tRNA synthetase |
| contig_29698_ 3 | 0.6085 | 1 | 1 | 0.83 54 | 1 | 1 | 0.63 97 | Light-harvesting complex I LH38 protein |
| contig_25351_ 2 | 0.6091 | 1 | 1 | 0.83 54 | 1 | 1 | 0.63 97 | predicted protein |
| contig_35523_ 6 | 0.6092 | 1 | 1 | 0.83 54 | 1 | 1 | 0.63 97 | 60S ribosomal protein L10a |
| contig_14621_ 1 | 0.6109 | 1 | 1 | 0.83 7 | 1 | 1 | 0.64 09 | predicted protein |
| contig_31285_ 5 | 0.6123 | 1 | 1 | 0.83 81 | 1 | 1 | 0.64 18 | threonine ammonia-lyase biosynthetic |
| contig_14126_ 5 | 0.6129 | 1 | 1 | 0.83 82 | 1 | 1 | 0.64 19 | predicted protein |
| contig_34452_ 4 | 0.6142 | 1 | 1 | 0.83 86 | 1 | 1 | 0.64 21 | glyceraldehyde-3-phosphate dehydrogenase precursor |
| contig_27420_ 4 | 0.6143 | 1 | 1 | 0.83 86 | 1 | 1 | 0.64 21 | polyhydroxybutyrate depolymerase |
| contig_15891_ 1 | 0.6158 | 1 | 1 | 0.83 97 | 1 | 1 | 0.64 3 | predicted protein |
| contig_30686_ 5 | 0.6168 | 1 | 1 | 0.83 97 | 1 | 1 | 0.64 3 | chlorophyll synthase chloroplastic |
| contig_29442_ 5 | 0.6178 | 1 | 1 | 0.83 97 | 1 | 1 | 0.64 3 | eukaryotic translation initiation factor 2 subunit alpha |

| | | | | | | | | |
|---|---|---|---|---|---|---|---|---|
| contig_28304_6 | 0.6183 | 1 | 1 | 0.8397 | 1 | 1 | 0.643 | predicted protein |
| contig_9471_1 | 0.6185 | 1 | 1 | 0.8397 | 1 | 1 | 0.643 | photosystem II CP43 chlorophyll apoprotein (chloroplast) |
| contig_36054_1 | 0.6188 | 1 | 1 | 0.8397 | 1 | 1 | 0.643 | protein fucoxanthin chlorophyll a/c protein |
| contig_22969_5 | 0.62 | 1 | 1 | 0.8397 | 1 | 1 | 0.643 | molecular chaperone DnaJ |
| contig_8256_2 | 0.6203 | 1 | 1 | 0.8397 | 1 | 1 | 0.643 | importin subunit beta putative |
| contig_24539_2 | 0.6209 | 1 | 1 | 0.8397 | 1 | 1 | 0.643 | phosphomannomutase/phosphoglucomutase |
| contig_26089_5 | 0.621 | 1 | 1 | 0.8397 | 1 | 1 | 0.643 | exportin-like protein |
| contig_34883_3 | 0.6211 | 1 | 1 | 0.8397 | 1 | 1 | 0.643 | Fructose-bisphosphate aldolase |
| contig_28636_6 | 0.6237 | 1 | 1 | 0.8424 | 1 | 1 | 0.6451 | Small nuclear ribonucleoprotein G |
| contig_9050_3 | 0.6252 | 1 | 1 | 0.8432 | 1 | 1 | 0.6457 | triose phosphate/phosphate translocator |
| contig_33090_4 | 0.6254 | 1 | 1 | 0.8432 | 1 | 1 | 0.6457 | predicted protein |
| contig_26806_4 | 0.6274 | 1 | 1 | 0.845 | 1 | 1 | 0.6471 | rab11 protein |
| contig_9755_4 | 0.6278 | 1 | 1 | 0.845 | 1 | 1 | 0.6471 | valyl-tRNA synthetase |
| contig_21631_6 | 0.6299 | 1 | 1 | 0.847 | 1 | 1 | 0.6486 | predicted protein |
| contig_28501_6 | 0.6341 | 1 | 1 | 0.8483 | 1 | 1 | 0.6496 | predicted protein |
| contig_9513_6 | 0.6342 | 1 | 1 | 0.8483 | 1 | 1 | 0.6496 | predicted protein |
| contig_15954_4 | 0.6342 | 1 | 1 | 0.8483 | 1 | 1 | 0.6496 | elongation factor-like |
| contig_23244_4 | 0.6344 | 1 | 1 | 0.8483 | 1 | 1 | 0.6496 | predicted protein |
| contig_27905_6 | 0.6345 | 1 | 1 | 0.8483 | 1 | 1 | 0.6496 | predicted protein |
| contig_22476_2 | 0.635 | 1 | 1 | 0.8483 | 1 | 1 | 0.6496 | predicted protein |
| contig_25833_1 | 0.6351 | 1 | 1 | 0.8483 | 1 | 1 | 0.6496 | predicted protein |
| contig_13306_5 | 0.6353 | 1 | 1 | 0.8483 | 1 | 1 | 0.6496 | predicted protein |
| contig_17562_5 | 0.6372 | 1 | 1 | 0.8501 | 1 | 1 | 0.651 | patched domain-containing protein 3-like |
| contig_18889_1 | 0.639 | 1 | 1 | 0.8513 | 1 | 1 | 0.6519 | ---NA--- |
| contig_22796_6 | 0.6392 | 1 | 1 | 0.8513 | 1 | 1 | 0.6519 | predicted protein |
| contig_25835_2 | 0.6412 | 1 | 1 | 0.8528 | 1 | 1 | 0.6531 | histidinol dehydrogenase |
| contig_20295_6 | 0.6414 | 1 | 1 | 0.8528 | 1 | 1 | 0.6531 | cytochrome c oxidase subunit II (mitochondrion) |

| | | | | | | | | |
|---|---|---|---|---|---|---|---|---|
| contig_33127_4 | 0.6436 | 1 | 1 | 0.855 | 1 | 1 | 0.6547 | Not1-domain-containing protein |
| contig_6511_6 | 0.6446 | 1 | 1 | 0.8556 | 1 | 1 | 0.6552 | glutamyl-tRNA synthetase |
| contig_21644_6 | 0.6459 | 1 | 1 | 0.8565 | 1 | 1 | 0.6559 | putative manganese superoxide dismutase protein |
| contig_27826_5 | 0.6468 | 1 | 1 | 0.8569 | 1 | 1 | 0.6562 | predicted protein |
| contig_8482_6 | 0.6489 | 1 | 1 | 0.8591 | 1 | 1 | 0.6578 | UV excision repair protein RAD23 |
| contig_17493_5 | 0.6495 | 1 | 1 | 0.8591 | 1 | 1 | 0.6578 | lamin tail domain-containing protein |
| contig_30021_2 | 0.6505 | 1 | 1 | 0.8591 | 1 | 1 | 0.6579 | predicted protein |
| contig_10661_6 | 0.6506 | 1 | 1 | 0.8591 | 1 | 1 | 0.6579 | glutaredoxin 2 |
| contig_9129_6 | 0.6528 | 1 | 1 | 0.8608 | 1 | 1 | 0.6592 | predicted protein |
| contig_11835_4 | 0.653 | 1 | 1 | 0.8608 | 1 | 1 | 0.6592 | ion transporter |
| contig_15845_2 | 0.6536 | 1 | 1 | 0.8608 | 1 | 1 | 0.6592 | predicted protein |
| contig_23551_6 | 0.6561 | 1 | 1 | 0.8633 | 1 | 1 | 0.6611 | predicted protein |
| contig_24967_5 | 0.6584 | 1 | 1 | 0.8656 | 1 | 1 | 0.6629 | PGR5-like protein 1A chloroplastic |
| contig_33594_1 | 0.6599 | 1 | 1 | 0.8666 | 1 | 1 | 0.6636 | CULlin protein 1 |
| contig_38142_5 | 0.6611 | 1 | 1 | 0.8666 | 1 | 1 | 0.6636 | ATP synthase F1 subunit alpha (mitochondrion) |
| contig_5863_5 | 0.6617 | 1 | 1 | 0.8666 | 1 | 1 | 0.6636 | predicted protein |
| contig_21238_5 | 0.6623 | 1 | 1 | 0.8666 | 1 | 1 | 0.6636 | fucoxanthin chlorophyll a/c protein 8 |
| contig_23946_2 | 0.6625 | 1 | 1 | 0.8666 | 1 | 1 | 0.6636 | ATP-binding cassette subfamily F member 3 |
| contig_28463_3 | 0.663 | 1 | 1 | 0.8666 | 1 | 1 | 0.6636 | aspartate kinase |
| contig_25876_4 | 0.6632 | 1 | 1 | 0.8666 | 1 | 1 | 0.6636 | predicted protein |
| contig_33296_6 | 0.6637 | 1 | 1 | 0.8666 | 1 | 1 | 0.6636 | light-harvesting protein |
| contig_27190_6 | 0.6663 | 1 | 1 | 0.8684 | 1 | 1 | 0.665 | 20S proteasome subunit alpha 2 |
| contig_28412_6 | 0.6671 | 1 | 1 | 0.8684 | 1 | 1 | 0.665 | predicted protein |
| contig_22960_6 | 0.6674 | 1 | 1 | 0.8684 | 1 | 1 | 0.665 | urease alpha subunit |
| contig_6097_6 | 0.6686 | 1 | 1 | 0.8684 | 1 | 1 | 0.665 | S-adenosylmethionine synthase 2 |
| contig_23621_6 | 0.6687 | 1 | 1 | 0.8684 | 1 | 1 | 0.665 | predicted protein |
| contig_34378_5 | 0.6699 | 1 | 1 | 0.8684 | 1 | 1 | 0.665 | predicted protein |

| | | | | | | | | |
|---|---|---|---|---|---|---|---|---|
| contig_25680_5 | 0.67 | 1 | 1 | 0.8684 | 1 | 1 | 0.665 | glucose-6-phosphate isomerase |
| contig_30420_3 | 0.6706 | 1 | 1 | 0.8684 | 1 | 1 | 0.665 | predicted protein |
| contig_25106_6 | 0.6706 | 1 | 1 | 0.8684 | 1 | 1 | 0.665 | predicted protein |
| contig_22684_3 | 0.6707 | 1 | 1 | 0.8684 | 1 | 1 | 0.665 | predicted protein |
| contig_38859_2 | 0.6722 | 1 | 1 | 0.8688 | 1 | 1 | 0.6653 | hypersensitive-induced response protein 1-like |
| contig_29033_5 | 0.6727 | 1 | 1 | 0.8688 | 1 | 1 | 0.6653 | EF-hand domain-containing protein |
| contig_24628_4 | 0.6729 | 1 | 1 | 0.8688 | 1 | 1 | 0.6653 | predicted protein |
| contig_30824_4 | 0.6734 | 1 | 1 | 0.8688 | 1 | 1 | 0.6653 | predicted protein |
| contig_29863_5 | 0.6756 | 1 | 1 | 0.8707 | 1 | 1 | 0.6668 | Homoserine dehydrogenase [ |
| contig_23908_1 | 0.676 | 1 | 1 | 0.8707 | 1 | 1 | 0.6668 | mitochondrial carrier protein |
| contig_7026_3 | 0.6768 | 1 | 1 | 0.8707 | 1 | 1 | 0.6668 | 26S proteasome non-ATPase regulatory subunit 2 |
| contig_40422_5 | 0.6775 | 1 | 1 | 0.8707 | 1 | 1 | 0.6668 | light harvesting protein |
| contig_29569_5 | 0.6777 | 1 | 1 | 0.8707 | 1 | 1 | 0.6668 | Sedoheptulose-1 7-bisphosphatase chloroplastic |
| contig_36999_3 | 0.6784 | 1 | 1 | 0.8709 | 1 | 1 | 0.6669 | predicted protein |
| contig_26282_6 | 0.6802 | 1 | 1 | 0.8717 | 1 | 1 | 0.6675 | dihydrodipicolinate reductase |
| contig_33470_2 | 0.6814 | 1 | 1 | 0.8717 | 1 | 1 | 0.6675 | farnesyl pyrophosphate synthase |
| contig_6613_6 | 0.6818 | 1 | 1 | 0.8717 | 1 | 1 | 0.6675 | sodium-coupled neutral amino acid transporter |
| contig_6474_4 | 0.6823 | 1 | 1 | 0.8717 | 1 | 1 | 0.6675 | FAD-dependent monooxygenase |
| contig_14698_3 | 0.6828 | 1 | 1 | 0.8717 | 1 | 1 | 0.6675 | AP-1 complex subunit gamma-1 |
| contig_16105_5 | 0.6828 | 1 | 1 | 0.8717 | 1 | 1 | 0.6675 | hypothetical protein FRACYDRAFT_264519 |
| contig_9754_5 | 0.683 | 1 | 1 | 0.8717 | 1 | 1 | 0.6675 | long-chain-fatty-acid-CoA ligase |
| contig_6079_5 | 0.686 | 1 | 1 | 0.8745 | 1 | 1 | 0.6696 | predicted protein |
| contig_25211_4 | 0.6863 | 1 | 1 | 0.8745 | 1 | 1 | 0.6696 | silicon transporter |
| contig_27545_4 | 0.6873 | 1 | 1 | 0.8745 | 1 | 1 | 0.6696 | predicted protein |
| contig_28535_5 | 0.6874 | 1 | 1 | 0.8745 | 1 | 1 | 0.6696 | 26S proteasome non-ATPase regulatory subunit 2 |
| contig_34814_4 | 0.689 | 1 | 1 | 0.8757 | 1 | 1 | 0.6706 | predicted protein |
| contig_17548_1 | 0.6923 | 1 | 1 | 0.8784 | 1 | 1 | 0.6726 | DUF1772 domain-containing protein |

| | | | | | | | | |
|---|---|---|---|---|---|---|---|---|
| contig_24180_5 | 0.6927 | 1 | 1 | 0.8784 | 1 | 1 | 0.6726 | predicted protein |
| contig_15626_6 | 0.6928 | 1 | 1 | 0.8784 | 1 | 1 | 0.6726 | predicted protein |
| contig_40291_3 | 0.6943 | 1 | 1 | 0.8795 | 1 | 1 | 0.6735 | predicted protein |
| contig_9710_5 | 0.6952 | 1 | 1 | 0.88 | 1 | 1 | 0.6738 | predicted protein |
| contig_29484_4 | 0.696 | 1 | 1 | 0.8802 | 1 | 1 | 0.674 | serine hydroxymethyltransferase |
| contig_6927_4 | 0.6991 | 1 | 1 | 0.8834 | 1 | 1 | 0.6765 | predicted protein |
| contig_41167_2 | 0.7004 | 1 | 1 | 0.8843 | 1 | 1 | 0.6772 | oxygen-evolving enhancer protein 1 precursor |
| contig_33304_4 | 0.7031 | 1 | 1 | 0.887 | 1 | 1 | 0.6793 | predicted protein |
| contig_8049_3 | 0.7043 | 1 | 1 | 0.8877 | 1 | 1 | 0.6798 | predicted protein |
| contig_9214_2 | 0.7059 | 1 | 1 | 0.8881 | 1 | 1 | 0.6801 | AMP-dependent synthetase |
| contig_6511_2 | 0.706 | 1 | 1 | 0.8881 | 1 | 1 | 0.6801 | glutamyl-tRNA synthetase |
| contig_9226_3 | 0.7065 | 1 | 1 | 0.8881 | 1 | 1 | 0.6801 | 2-isopropylmalate synthase |
| contig_5530_1 | 0.7069 | 1 | 1 | 0.8881 | 1 | 1 | 0.6801 | NAD(P)-dependent steroid dehydrogenase-like protein |
| contig_6303_5 | 0.708 | 1 | 1 | 0.8888 | 1 | 1 | 0.6806 | acyl carrier protein (chloroplast) |
| contig_8347_5 | 0.7112 | 1 | 1 | 0.8911 | 1 | 1 | 0.6824 | elongation factor Ts |
| contig_17183_2 | 0.7113 | 1 | 1 | 0.8911 | 1 | 1 | 0.6824 | 40S ribosomal protein S17 |
| contig_23301_2 | 0.7116 | 1 | 1 | 0.8911 | 1 | 1 | 0.6824 | protein fucoxanthin chlorophyll a/c protein |
| contig_26788_4 | 0.7122 | 1 | 1 | 0.8911 | 1 | 1 | 0.6824 | regulatory proteasome non-atpase subunit 2 |
| contig_39446_2 | 0.7137 | 1 | 1 | 0.8922 | 1 | 1 | 0.6832 | U6 snRNA-associated Sm-like protein LSm2 |
| contig_38574_3 | 0.7151 | 1 | 1 | 0.8933 | 1 | 1 | 0.684 | predicted protein |
| contig_10358_5 | 0.7165 | 1 | 1 | 0.8937 | 1 | 1 | 0.6843 | 60S ribosomal protein L12 |
| contig_25647_1 | 0.7166 | 1 | 1 | 0.8937 | 1 | 1 | 0.6843 | putative all-trans-retinol 13 14-reductase |
| contig_24139_1 | 0.7173 | 1 | 1 | 0.8938 | 1 | 1 | 0.6844 | chaperonin GroEL |
| contig_39533_1 | 0.7182 | 1 | 1 | 0.8942 | 1 | 1 | 0.6847 | sodium/proton antiporter 1 |
| contig_10808_6 | 0.7204 | 1 | 1 | 0.8963 | 1 | 1 | 0.6863 | predicted protein |
| contig_33453_2 | 0.7226 | 1 | 1 | 0.8983 | 1 | 1 | 0.6879 | predicted protein |
| contig_5663_5 | 0.7234 | 1 | 1 | 0.8985 | 1 | 1 | 0.688 | tubulin beta chain |

| | | | | | | | | |
|---|---|---|---|---|---|---|---|---|
| contig_38505_3 | 0.7269 | 1 | 1 | 0.9021 | 1 | 1 | 0.6908 | serine/threonine-protein phosphatase 2A regulatory subunit B' |
| contig_25953_4 | 0.7288 | 1 | 1 | 0.9036 | 1 | 1 | 0.692 | predicted protein |
| contig_14563_3 | 0.7293 | 1 | 1 | 0.9036 | 1 | 1 | 0.692 | predicted protein |
| contig_5361_1 | 0.7316 | 1 | 1 | 0.9051 | 1 | 1 | 0.6931 | ---NA--- |
| contig_34164_6 | 0.7319 | 1 | 1 | 0.9051 | 1 | 1 | 0.6931 | multispanning membrane protein |
| contig_18025_5 | 0.7322 | 1 | 1 | 0.9051 | 1 | 1 | 0.6931 | histone H2B |
| contig_8921_1 | 0.7333 | 1 | 1 | 0.9056 | 1 | 1 | 0.6935 | predicted protein |
| contig_14697_5 | 0.7346 | 1 | 1 | 0.9065 | 1 | 1 | 0.6942 | fructose 1 6-bisphosphatase |
| contig_24802_3 | 0.7356 | 1 | 1 | 0.907 | 1 | 1 | 0.6945 | predicted protein |
| contig_19038_3 | 0.7366 | 1 | 1 | 0.9075 | 1 | 1 | 0.6949 | peptidylprolyl isomerase |
| contig_16053_5 | 0.7382 | 1 | 1 | 0.9088 | 1 | 1 | 0.6959 | predicted protein |
| contig_32942_6 | 0.7402 | 1 | 1 | 0.9096 | 1 | 1 | 0.6965 | predicted protein |
| contig_6643_5 | 0.7412 | 1 | 1 | 0.9096 | 1 | 1 | 0.6965 | phosphoglycolate phosphatase |
| contig_28255_4 | 0.7424 | 1 | 1 | 0.9096 | 1 | 1 | 0.6965 | long chain acyl-coa synthetase |
| contig_26782_3 | 0.7428 | 1 | 1 | 0.9096 | 1 | 1 | 0.6965 | predicted protein |
| contig_15258_2 | 0.7432 | 1 | 1 | 0.9096 | 1 | 1 | 0.6965 | predicted protein |
| contig_28744_4 | 0.7433 | 1 | 1 | 0.9096 | 1 | 1 | 0.6965 | ascorbate peroxidase |
| contig_30693_4 | 0.7441 | 1 | 1 | 0.9096 | 1 | 1 | 0.6965 | ubiquitin-conjugating enzyme E2-16 kDa |
| contig_7380_4 | 0.7449 | 1 | 1 | 0.9096 | 1 | 1 | 0.6965 | predicted protein |
| contig_29342_2 | 0.745 | 1 | 1 | 0.9096 | 1 | 1 | 0.6965 | leukotriene A-4 hydrolase/aminopeptidase |
| contig_22932_5 | 0.7456 | 1 | 1 | 0.9096 | 1 | 1 | 0.6965 | predicted protein |
| contig_32742_5 | 0.7458 | 1 | 1 | 0.9096 | 1 | 1 | 0.6965 | predicted protein |
| contig_22496_6 | 0.746 | 1 | 1 | 0.9096 | 1 | 1 | 0.6965 | predicted protein |
| contig_9153_4 | 0.7503 | 1 | 1 | 0.9136 | 1 | 1 | 0.6996 | predicted protein |
| contig_40568_2 | 0.7505 | 1 | 1 | 0.9136 | 1 | 1 | 0.6996 | glutathione reductase |
| contig_11147_6 | 0.7525 | 1 | 1 | 0.9154 | 1 | 1 | 0.701 | predicted protein |

| | | | | | | | | |
|---|---|---|---|---|---|---|---|---|
| contig_12511_6 | 0.7534 | 1 | 1 | 0.9157 | 1 | 1 | 0.7012 | predicted protein |
| contig_5934_3 | 0.755 | 1 | 1 | 0.9158 | 1 | 1 | 0.7013 | bifunctional aconitate hydratase 2/2-methylisocitrate dehydratase |
| contig_36134_4 | 0.755 | 1 | 1 | 0.9158 | 1 | 1 | 0.7013 | fucoxanthin chlorophyll a/c protein |
| contig_35847_5 | 0.7554 | 1 | 1 | 0.9158 | 1 | 1 | 0.7013 | serine/threonine-protein phosphatase PP1-beta catalytic subunit |
| contig_19197_6 | 0.7562 | 1 | 1 | 0.9158 | 1 | 1 | 0.7013 | casein kinase I isoform X1 |
| contig_6218_5 | 0.7565 | 1 | 1 | 0.9158 | 1 | 1 | 0.7013 | ferredoxin-nitrite reductase |
| contig_40094_2 | 0.7583 | 1 | 1 | 0.9173 | 1 | 1 | 0.7024 | 40S ribosomal protein S14 |
| contig_11460_2 | 0.7592 | 1 | 1 | 0.9176 | 1 | 1 | 0.7027 | predicted protein |
| contig_33374_5 | 0.7604 | 1 | 1 | 0.9183 | 1 | 1 | 0.7032 | predicted protein |
| contig_21411_6 | 0.7621 | 1 | 1 | 0.9187 | 1 | 1 | 0.7035 | ATP-dependent RNA helicase eIF4A |
| contig_35273_1 | 0.7623 | 1 | 1 | 0.9187 | 1 | 1 | 0.7035 | Dethiobiotin synthase |
| contig_12161_5 | 0.7625 | 1 | 1 | 0.9187 | 1 | 1 | 0.7035 | predicted protein |
| contig_15393_2 | 0.7638 | 1 | 1 | 0.9196 | 1 | 1 | 0.7042 | predicted protein |
| contig_35069_1 | 0.765 | 1 | 1 | 0.9198 | 1 | 1 | 0.7043 | translation elongation factor-like protein |
| contig_26111_3 | 0.7654 | 1 | 1 | 0.9198 | 1 | 1 | 0.7043 | predicted protein |
| contig_30780_6 | 0.7663 | 1 | 1 | 0.9198 | 1 | 1 | 0.7043 | possible transporter |
| contig_25881_5 | 0.7666 | 1 | 1 | 0.9198 | 1 | 1 | 0.7043 | predicted protein |
| contig_39705_5 | 0.767 | 1 | 1 | 0.9198 | 1 | 1 | 0.7043 | inorganic pyrophosphatase |
| contig_20147_4 | 0.7701 | 1 | 1 | 0.9221 | 1 | 1 | 0.7061 | fucoxanthin chlorophyl a/c protein |
| contig_11320_6 | 0.7702 | 1 | 1 | 0.9221 | 1 | 1 | 0.7061 | predicted protein |
| contig_30869_3 | 0.7714 | 1 | 1 | 0.923 | 1 | 1 | 0.7068 | predicted protein |
| contig_19770_6 | 0.7759 | 1 | 1 | 0.9268 | 1 | 1 | 0.7097 | predicted protein |
| contig_18233_1 | 0.7761 | 1 | 1 | 0.9268 | 1 | 1 | 0.7097 | dihydrolipoamide dehydrogenase |
| contig_11625_6 | 0.7765 | 1 | 1 | 0.9268 | 1 | 1 | 0.7097 | T9SS type A sorting domain-containing protein |
| contig_30073_5 | 0.7772 | 1 | 1 | 0.9269 | 1 | 1 | 0.7098 | transaldolase |
| contig_10727_3 | 0.7782 | 1 | 1 | 0.9274 | 1 | 1 | 0.7102 | eukaryotic translation initiation factor 3 subunit G |

| | | | | | | | | |
|---|---|---|---|---|---|---|---|---|
| contig_33083_6 | 0.7794 | 1 | 1 | 0.9281 | 1 | 1 | 0.7107 | Glucose-6-phosphate isomerase cytosolic |
| contig_33488_4 | 0.7812 | 1 | 1 | 0.9292 | 1 | 1 | 0.7116 | transcription elongation factor SPT6 |
| contig_28331_5 | 0.7816 | 1 | 1 | 0.9292 | 1 | 1 | 0.7116 | transketolase |
| contig_27521_6 | 0.7846 | 1 | 1 | 0.9322 | 1 | 1 | 0.7138 | predicted protein |
| contig_16859_4 | 0.7869 | 1 | 1 | 0.9342 | 1 | 1 | 0.7154 | DNA-directed RNA polymerase II 135 kDa polypeptide |
| contig_6981_5 | 0.7885 | 1 | 1 | 0.9347 | 1 | 1 | 0.7157 | predicted protein |
| contig_25670_6 | 0.7887 | 1 | 1 | 0.9347 | 1 | 1 | 0.7157 | predicted protein |
| contig_5926_5 | 0.7892 | 1 | 1 | 0.9347 | 1 | 1 | 0.7157 | predicted protein |
| contig_8125_5 | 0.7917 | 1 | 1 | 0.9364 | 1 | 1 | 0.7171 | adenosylhomocysteinase |
| contig_29330_2 | 0.7938 | 1 | 1 | 0.9364 | 1 | 1 | 0.7171 | pilus assembly protein |
| contig_8784_1 | 0.7939 | 1 | 1 | 0.9364 | 1 | 1 | 0.7171 | 40S ribosomal protein S10 |
| contig_7643_5 | 0.794 | 1 | 1 | 0.9364 | 1 | 1 | 0.7171 | ATP synthase CF1 beta subunit (chloroplast) |
| contig_36399_5 | 0.7943 | 1 | 1 | 0.9364 | 1 | 1 | 0.7171 | triosephosphate isomerase/glyceraldehyde-3-phosphate dehydrogenase precursor |
| contig_7412_1 | 0.7943 | 1 | 1 | 0.9364 | 1 | 1 | 0.7171 | E3 ubiquitin-protein ligase RING2-like |
| contig_21581_3 | 0.7957 | 1 | 1 | 0.9372 | 1 | 1 | 0.7177 | predicted protein |
| contig_27423_4 | 0.7962 | 1 | 1 | 0.9372 | 1 | 1 | 0.7177 | predicted protein |
| contig_8371_4 | 0.8004 | 1 | 1 | 0.9379 | 1 | 1 | 0.7182 | Rab family other |
| contig_22675_1 | 0.8007 | 1 | 1 | 0.9379 | 1 | 1 | 0.7182 | predicted protein |
| contig_40734_1 | 0.8017 | 1 | 1 | 0.9379 | 1 | 1 | 0.7182 | predicted protein |
| contig_27060_3 | 0.8019 | 1 | 1 | 0.9379 | 1 | 1 | 0.7182 | calcium-binding protein |
| contig_15521_3 | 0.8037 | 1 | 1 | 0.9379 | 1 | 1 | 0.7182 | predicted protein |
| contig_34791_3 | 0.8048 | 1 | 1 | 0.9379 | 1 | 1 | 0.7182 | Regulator of chromosome condensation 1/beta-lactamase-inhibitor protein II |
| contig_30060_4 | 0.8056 | 1 | 1 | 0.9379 | 1 | 1 | 0.7182 | nitrite transporter NAR1 |
| contig_5668_1 | 0.8057 | 1 | 1 | 0.9379 | 1 | 1 | 0.7182 | 40s ribosomal protein s20 |
| contig_25275_2 | 0.8069 | 1 | 1 | 0.9379 | 1 | 1 | 0.7182 | histidine-tRNA ligase |
| contig_30644_2 | 0.8086 | 1 | 1 | 0.9379 | 1 | 1 | 0.7182 | predicted protein |

| | | | | | | | | |
|---|---|---|---|---|---|---|---|---|
| contig_33426_1 | 0.8088 | 1 | 1 | 0.9379 | 1 | 1 | 0.7182 | predicted protein |
| contig_17076_5 | 0.8098 | 1 | 1 | 0.9379 | 1 | 1 | 0.7182 | predicted protein |
| contig_18043_1 | 0.8099 | 1 | 1 | 0.9379 | 1 | 1 | 0.7182 | Light-harvesting complex I LH38 protein |
| contig_26837_3 | 0.8101 | 1 | 1 | 0.9379 | 1 | 1 | 0.7182 | AP-1 complex subunit gamma-1 |
| contig_25573_5 | 0.8103 | 1 | 1 | 0.9379 | 1 | 1 | 0.7182 | predicted protein |
| contig_30561_2 | 0.8109 | 1 | 1 | 0.9379 | 1 | 1 | 0.7182 | cytochrome b6-f complex iron-sulfur subunit |
| contig_14140_6 | 0.8119 | 1 | 1 | 0.9379 | 1 | 1 | 0.7182 | predicted protein |
| contig_12195_1 | 0.8126 | 1 | 1 | 0.9379 | 1 | 1 | 0.7182 | thioredoxin-like protein 4A isoform X1 |
| contig_28208_6 | 0.813 | 1 | 1 | 0.9379 | 1 | 1 | 0.7182 | SPFH/Band 7/PHB domain protein |
| contig_40102_3 | 0.8134 | 1 | 1 | 0.9379 | 1 | 1 | 0.7182 | predicted protein |
| contig_29868_5 | 0.8141 | 1 | 1 | 0.9379 | 1 | 1 | 0.7182 | adp-ribosylation factor 1 |
| contig_33784_6 | 0.8142 | 1 | 1 | 0.9379 | 1 | 1 | 0.7182 | predicted protein |
| contig_15405_3 | 0.8145 | 1 | 1 | 0.9379 | 1 | 1 | 0.7182 | predicted protein |
| contig_28443_2 | 0.8147 | 1 | 1 | 0.9379 | 1 | 1 | 0.7182 | nad-dependent epimerase/dehydratase |
| contig_39623_1 | 0.8151 | 1 | 1 | 0.9379 | 1 | 1 | 0.7182 | predicted protein |
| contig_27804_5 | 0.8152 | 1 | 1 | 0.9379 | 1 | 1 | 0.7182 | Coatomer subunit alpha-1 |
| contig_8226_3 | 0.8161 | 1 | 1 | 0.9379 | 1 | 1 | 0.7182 | NAD(P) transhydrogenase mitochondrial |
| contig_7582_3 | 0.8163 | 1 | 1 | 0.9379 | 1 | 1 | 0.7182 | vanadium-dependent haloperoxidase |
| contig_8332_1 | 0.817 | 1 | 1 | 0.9379 | 1 | 1 | 0.7182 | oxoglutarate dehydrogenase (succinyl-transferring) E1 component |
| contig_15868_5 | 0.8173 | 1 | 1 | 0.9379 | 1 | 1 | 0.7182 | fucoxanthin-chlorophyll a-c binding protein |
| contig_6069_1 | 0.8175 | 1 | 1 | 0.9379 | 1 | 1 | 0.7182 | GLNA glutamine synthase |
| contig_14556_3 | 0.8177 | 1 | 1 | 0.9379 | 1 | 1 | 0.7182 | anion-transporting ATPase |
| contig_6795_4 | 0.8181 | 1 | 1 | 0.9379 | 1 | 1 | 0.7182 | WSC domain protein putative |
| contig_36071_6 | 0.8182 | 1 | 1 | 0.9379 | 1 | 1 | 0.7182 | predicted protein |
| contig_38073_1 | 0.8187 | 1 | 1 | 0.9379 | 1 | 1 | 0.7182 | ---NA--- |
| contig_41369_3 | 0.8189 | 1 | 1 | 0.9379 | 1 | 1 | 0.7182 | predicted protein |

| | | | | | | | | |
|---|---|---|---|---|---|---|---|---|
| contig_19181_5 | 0.82 | 1 | 1 | 0.9385 | 1 | 1 | 0.7186 | ribosomal protein S4 |
| contig_10034_3 | 0.821 | 1 | 1 | 0.9385 | 1 | 1 | 0.7187 | 60S ribosomal protein L10 |
| contig_14343_6 | 0.8213 | 1 | 1 | 0.9385 | 1 | 1 | 0.7187 | riboflavin synthase alpha subunit |
| contig_9968_3 | 0.8232 | 1 | 1 | 0.9393 | 1 | 1 | 0.7193 | predicted protein |
| contig_24750_1 | 0.8232 | 1 | 1 | 0.9393 | 1 | 1 | 0.7193 | AP-1 complex subunit beta-1 |
| contig_17633_4 | 0.8257 | 1 | 1 | 0.9394 | 1 | 1 | 0.7194 | Adenylyl cyclase-associated protein |
| contig_15357_4 | 0.8263 | 1 | 1 | 0.9394 | 1 | 1 | 0.7194 | calreticulin |
| contig_26135_5 | 0.8271 | 1 | 1 | 0.9394 | 1 | 1 | 0.7194 | Mitochondrial Carrier (MC) Family |
| contig_27847_4 | 0.8272 | 1 | 1 | 0.9394 | 1 | 1 | 0.7194 | unnamed protein product |
| contig_9571_3 | 0.8272 | 1 | 1 | 0.9394 | 1 | 1 | 0.7194 | ATP:ADP antiporter AAA family |
| contig_9646_5 | 0.8279 | 1 | 1 | 0.9394 | 1 | 1 | 0.7194 | cell differentiation protein |
| contig_17246_6 | 0.8282 | 1 | 1 | 0.9394 | 1 | 1 | 0.7194 | predicted protein |
| contig_41171_6 | 0.8285 | 1 | 1 | 0.9394 | 1 | 1 | 0.7194 | ADP/ATP transporter on adenylate translocase putative |
| contig_27998_6 | 0.8289 | 1 | 1 | 0.9394 | 1 | 1 | 0.7194 | 2 3-bisphosphoglycerate-dependent phosphoglycerate mutase |
| contig_30861_4 | 0.8309 | 1 | 1 | 0.9411 | 1 | 1 | 0.7206 | fucoxanthin chlorophyll a/c protein 8 |
| contig_39210_6 | 0.832 | 1 | 1 | 0.9412 | 1 | 1 | 0.7207 | predicted protein |
| contig_23006_1 | 0.8323 | 1 | 1 | 0.9412 | 1 | 1 | 0.7207 | predicted protein |
| contig_3138_3 | 0.8337 | 1 | 1 | 0.9422 | 1 | 1 | 0.7215 | predicted protein |
| contig_21628_5 | 0.8353 | 1 | 1 | 0.9433 | 1 | 1 | 0.7223 | Tubulin alpha-5 chain |
| contig_19158_3 | 0.8368 | 1 | 1 | 0.9438 | 1 | 1 | 0.7227 | small ubiquitin-like modifier (SUMO) |
| contig_10682_1 | 0.8381 | 1 | 1 | 0.9438 | 1 | 1 | 0.7227 | predicted protein |
| contig_9050_5 | 0.8383 | 1 | 1 | 0.9438 | 1 | 1 | 0.7227 | triose phosphate/phosphate translocator |
| contig_11186_4 | 0.8387 | 1 | 1 | 0.9438 | 1 | 1 | 0.7227 | predicted protein |
| contig_30749_6 | 0.8388 | 1 | 1 | 0.9438 | 1 | 1 | 0.7227 | fucoxanthin-chlorophyll a/c light-harvesting protein |
| contig_21909_6 | 0.8395 | 1 | 1 | 0.9438 | 1 | 1 | 0.7227 | 26S proteasome beta type 7 subunit |
| contig_16732_6 | 0.8423 | 1 | 1 | 0.9463 | 1 | 1 | 0.7247 | membrane spanning protein of the nonaspanin family |
| contig_23694_6 | 0.8435 | 1 | 1 | 0.947 | 1 | 1 | 0.7251 | cell division cycle protein 123 homolog |

| | | | | | | | | |
|---|---|---|---|---|---|---|---|---|
| contig_27657_5 | 0.8449 | 1 | 1 | 0.9478 | 1 | 1 | 0.7258 | solute carrier family 35 member E1 |
| contig_37712_6 | 0.8464 | 1 | 1 | 0.9479 | 1 | 1 | 0.7258 | predicted protein |
| contig_25826_3 | 0.8483 | 1 | 1 | 0.9479 | 1 | 1 | 0.7258 | heat shock protein 70 |
| contig_22773_6 | 0.849 | 1 | 1 | 0.9479 | 1 | 1 | 0.7258 | phosphoglycerate kinase |
| contig_27280_6 | 0.8492 | 1 | 1 | 0.9479 | 1 | 1 | 0.7258 | dihydrolipoamide dehydrogenase |
| contig_27293_4 | 0.8495 | 1 | 1 | 0.9479 | 1 | 1 | 0.7258 | predicted protein |
| contig_24334_5 | 0.8496 | 1 | 1 | 0.9479 | 1 | 1 | 0.7258 | predicted protein |
| contig_21155_4 | 0.8497 | 1 | 1 | 0.9479 | 1 | 1 | 0.7258 | heat shock 70 kDa protein |
| contig_10416_5 | 0.8499 | 1 | 1 | 0.9479 | 1 | 1 | 0.7258 | ATP synthase F0 subunit 6 (mitochondrion) |
| contig_37718_6 | 0.8516 | 1 | 1 | 0.9487 | 1 | 1 | 0.7265 | predicted protein |
| contig_10461_4 | 0.8521 | 1 | 1 | 0.9487 | 1 | 1 | 0.7265 | ATP synthase subunit beta mitochondrial |
| contig_14030_3 | 0.8525 | 1 | 1 | 0.9487 | 1 | 1 | 0.7265 | S-adenosylmethionine carrier 1 chloroplastic/mitochondrial |
| contig_9265_3 | 0.8537 | 1 | 1 | 0.9493 | 1 | 1 | 0.7269 | mitochondrial carrier family |
| contig_14919_6 | 0.8549 | 1 | 1 | 0.95 | 1 | 1 | 0.7275 | predicted protein |
| contig_30128_6 | 0.8563 | 1 | 1 | 0.9508 | 1 | 1 | 0.7281 | predicted protein |
| contig_39431_2 | 0.8597 | 1 | 1 | 0.9539 | 1 | 1 | 0.7305 | GDP-mannose transporter |
| contig_41308_6 | 0.8604 | 1 | 1 | 0.954 | 1 | 1 | 0.7305 | nucleoside diphosphate kinase |
| contig_36255_5 | 0.8615 | 1 | 1 | 0.9545 | 1 | 1 | 0.7309 | hypothetical protein THAOC_30038 |
| contig_6937_2 | 0.8623 | 1 | 1 | 0.9548 | 1 | 1 | 0.7311 | predicted protein |
| contig_25423_6 | 0.8641 | 1 | 1 | 0.9554 | 1 | 1 | 0.7316 | 2 5-diketo-D-gluconic acid reductase A putative |
| contig_18307_6 | 0.8642 | 1 | 1 | 0.9554 | 1 | 1 | 0.7316 | predicted protein |
| contig_4841_4 | 0.8665 | 1 | 1 | 0.9573 | 1 | 1 | 0.733 | 40S ribosomal protein S14 |
| contig_15461_5 | 0.868 | 1 | 1 | 0.9581 | 1 | 1 | 0.7336 | predicted protein |
| contig_6054_4 | 0.8686 | 1 | 1 | 0.9581 | 1 | 1 | 0.7336 | ---NA--- |
| contig_25557_4 | 0.8706 | 1 | 1 | 0.9581 | 1 | 1 | 0.7336 | nucleolar complex protein 2 |
| contig_29735_5 | 0.8709 | 1 | 1 | 0.9581 | 1 | 1 | 0.7336 | dihydrolipoamide succinyltransferase |
| contig_9387_4 | 0.8713 | 1 | 1 | 0.9581 | 1 | 1 | 0.7336 | DUF791-domain-containing protein |

| | | | | | | | | |
|---|---|---|---|---|---|---|---|---|
| contig_25921_4 | 0.8714 | 1 | 1 | 0.9581 | 1 | 1 | 0.7336 | predicted protein |
| contig_38406_2 | 0.8716 | 1 | 1 | 0.9581 | 1 | 1 | 0.7336 | predicted protein |
| contig_33557_6 | 0.8745 | 1 | 1 | 0.9602 | 1 | 1 | 0.7353 | phosphoglycerate kinase |
| contig_30596_2 | 0.8748 | 1 | 1 | 0.9602 | 1 | 1 | 0.7353 | predicted protein |
| contig_24729_6 | 0.8763 | 1 | 1 | 0.9608 | 1 | 1 | 0.7357 | predicted protein |
| contig_22787_6 | 0.8766 | 1 | 1 | 0.9608 | 1 | 1 | 0.7357 | bifunctional aspartate kinase/homoserine dehydrogenase I |
| contig_23768_5 | 0.8814 | 1 | 1 | 0.9654 | 1 | 1 | 0.7392 | Flavocytochrome c |
| contig_6442_4 | 0.8827 | 1 | 1 | 0.9661 | 1 | 1 | 0.7398 | putative ATP-dependent RNA helicase DHX30 isoform X3 |
| contig_22941_4 | 0.8872 | 1 | 1 | 0.9696 | 1 | 1 | 0.7425 | mitochondrial carrier |
| contig_15995_6 | 0.8889 | 1 | 1 | 0.9696 | 1 | 1 | 0.7425 | cyclopropane-fatty-acyl-phospholipid synthase |
| contig_30865_4 | 0.8895 | 1 | 1 | 0.9696 | 1 | 1 | 0.7425 | 2-oxo acid dehydrogenase subunit E2 |
| contig_17882_6 | 0.8898 | 1 | 1 | 0.9696 | 1 | 1 | 0.7425 | light harvesting protein |
| contig_24496_1 | 0.89 | 1 | 1 | 0.9696 | 1 | 1 | 0.7425 | protein RCC2 homolog |
| contig_19309_1 | 0.8906 | 1 | 1 | 0.9696 | 1 | 1 | 0.7425 | 60S ribosomal protein L9-like |
| contig_25855_2 | 0.891 | 1 | 1 | 0.9696 | 1 | 1 | 0.7425 | protein transport protein Sec61 subunit gamma-like |
| contig_20315_6 | 0.891 | 1 | 1 | 0.9696 | 1 | 1 | 0.7425 | NADH dehydrogenase subunit 5 (mitochondrion) |
| contig_28753_4 | 0.8942 | 1 | 1 | 0.9723 | 1 | 1 | 0.7446 | FKBP-type peptidyl-prolyl cis-trans isomerase |
| contig_32836_4 | 0.8947 | 1 | 1 | 0.9723 | 1 | 1 | 0.7446 | predicted protein |
| contig_8422_4 | 0.8958 | 1 | 1 | 0.9728 | 1 | 1 | 0.745 | predicted protein |
| contig_16000_5 | 0.8968 | 1 | 1 | 0.9729 | 1 | 1 | 0.745 | FAD-dependent oxidoreductase |
| contig_25076_6 | 0.8975 | 1 | 1 | 0.9729 | 1 | 1 | 0.745 | phosphoglucomutase |
| contig_34263_4 | 0.8978 | 1 | 1 | 0.9729 | 1 | 1 | 0.745 | nucleolar protein 56 |
| contig_34200_5 | 0.9005 | 1 | 1 | 0.9744 | 1 | 1 | 0.7461 | fructose-1 6-bisphosphatase |
| contig_14405_3 | 0.901 | 1 | 1 | 0.9744 | 1 | 1 | 0.7461 | predicted protein |
| contig_6050_1 | 0.902 | 1 | 1 | 0.9744 | 1 | 1 | 0.7461 | heat shock protein 70 (chloroplast) |
| contig_9065_4 | 0.9023 | 1 | 1 | 0.9744 | 1 | 1 | 0.7461 | tRNA cyclic N6-threonylcarbamoyladenosine(37) synthase TcdA |

| | | | | | | | | |
|---|---|---|---|---|---|---|---|---|
| contig_21520_1 | 0.9024 | 1 | 1 | 0.9744 | 1 | 1 | 0.7461 | phosphoglycerate mutase 1 family |
| contig_27547_5 | 0.9042 | 1 | 1 | 0.9757 | 1 | 1 | 0.7472 | ribulose-phosphate 3-epimerase |
| contig_20080_5 | 0.9055 | 1 | 1 | 0.9764 | 1 | 1 | 0.7477 | AP-1 complex subunit beta-1-like |
| contig_28740_4 | 0.9074 | 1 | 1 | 0.9772 | 1 | 1 | 0.7483 | tubulin alpha chain |
| contig_22479_1 | 0.9076 | 1 | 1 | 0.9772 | 1 | 1 | 0.7483 | predicted protein |
| contig_26083_5 | 0.9082 | 1 | 1 | 0.9772 | 1 | 1 | 0.7483 | phosphatidylinositol transfer protein 3-like |
| contig_15022_4 | 0.9108 | 1 | 1 | 0.9789 | 1 | 1 | 0.7496 | ---NA--- |
| contig_38299_1 | 0.9114 | 1 | 1 | 0.9789 | 1 | 1 | 0.7496 | ATP-binding cassette superfamily |
| contig_22050_5 | 0.9121 | 1 | 1 | 0.9789 | 1 | 1 | 0.7496 | 60S ribosomal protein L9 |
| contig_38591_1 | 0.9137 | 1 | 1 | 0.9789 | 1 | 1 | 0.7496 | 40S ribosomal protein S18 |
| contig_27261_5 | 0.9146 | 1 | 1 | 0.9789 | 1 | 1 | 0.7496 | enoyl-CoA hydratase |
| contig_14552_5 | 0.9147 | 1 | 1 | 0.9789 | 1 | 1 | 0.7496 | phosphomannomutase |
| contig_33399_2 | 0.9151 | 1 | 1 | 0.9789 | 1 | 1 | 0.7496 | predicted protein |
| contig_26112_6 | 0.9158 | 1 | 1 | 0.9789 | 1 | 1 | 0.7496 | adenylate kinase 5 chloroplastic isoform X1 |
| contig_16841_6 | 0.9158 | 1 | 1 | 0.9789 | 1 | 1 | 0.7496 | predicted protein |
| contig_9855_4 | 0.9161 | 1 | 1 | 0.9789 | 1 | 1 | 0.7496 | S-adenosyl-L-methionine-dependent methyltransferase |
| contig_23752_4 | 0.9179 | 1 | 1 | 0.9796 | 1 | 1 | 0.7501 | predicted protein |
| contig_5385_3 | 0.9181 | 1 | 1 | 0.9796 | 1 | 1 | 0.7501 | glutathione S-transferase Mu 3-like |
| contig_38588_6 | 0.9203 | 1 | 1 | 0.9799 | 1 | 1 | 0.7504 | Ras-related protein Rab-8A |
| contig_13588_3 | 0.9204 | 1 | 1 | 0.9799 | 1 | 1 | 0.7504 | aarF domain-containing kinase |
| contig_11923_6 | 0.9205 | 1 | 1 | 0.9799 | 1 | 1 | 0.7504 | aspartate aminotransferase |
| contig_12148_3 | 0.9224 | 1 | 1 | 0.9799 | 1 | 1 | 0.7504 | predicted protein |
| contig_9976_6 | 0.9226 | 1 | 1 | 0.9799 | 1 | 1 | 0.7504 | poly(U)-binding-splicing factor PUF60 |
| contig_19001_2 | 0.9227 | 1 | 1 | 0.9799 | 1 | 1 | 0.7504 | predicted protein |
| contig_23500_4 | 0.9229 | 1 | 1 | 0.9799 | 1 | 1 | 0.7504 | proline-tRNA ligase |
| contig_30727_4 | 0.9238 | 1 | 1 | 0.9799 | 1 | 1 | 0.7504 | predicted protein |
| contig_30690_3 | 0.9241 | 1 | 1 | 0.9799 | 1 | 1 | 0.7504 | phosphoglycerate mutase |

| | | | | | | | | |
|---|---|---|---|---|---|---|---|---|
| contig_38536_2 | 0.9256 | 1 | 1 | 0.9802 | 1 | 1 | 0.7506 | predicted protein |
| contig_25207_1 | 0.9263 | 1 | 1 | 0.9802 | 1 | 1 | 0.7506 | predicted protein |
| contig_18863_3 | 0.9264 | 1 | 1 | 0.9802 | 1 | 1 | 0.7506 | nucleoside diphosphate kinase |
| contig_24715_2 | 0.9275 | 1 | 1 | 0.9802 | 1 | 1 | 0.7506 | predicted protein |
| contig_41159_3 | 0.9279 | 1 | 1 | 0.9802 | 1 | 1 | 0.7506 | predicted protein |
| contig_8757_2 | 0.9288 | 1 | 1 | 0.9802 | 1 | 1 | 0.7506 | ABC transporter |
| contig_23603_4 | 0.9289 | 1 | 1 | 0.9802 | 1 | 1 | 0.7506 | serine protease |
| contig_26535_2 | 0.9337 | 1 | 1 | 0.9845 | 1 | 1 | 0.7539 | tubulin alpha chain |
| contig_27236_4 | 0.9358 | 1 | 1 | 0.9861 | 1 | 1 | 0.7551 | translation initiation factor aif-2 |
| contig_33687_4 | 0.9399 | 1 | 1 | 0.9895 | 1 | 1 | 0.7577 | predicted protein |
| contig_28369_6 | 0.9404 | 1 | 1 | 0.9895 | 1 | 1 | 0.7577 | predicted protein |
| contig_33750_4 | 0.9411 | 1 | 1 | 0.9896 | 1 | 1 | 0.7578 | Violaxanthin de-epoxidase |
| contig_14481_1 | 0.9433 | 1 | 1 | 0.9902 | 1 | 1 | 0.7582 | CDP-diacylglycerol--glycerol-3-phosphate 3-phosphatidyltransferase |
| contig_28448_5 | 0.9435 | 1 | 1 | 0.9902 | 1 | 1 | 0.7582 | glycine dehydrogenase |
| contig_22324_4 | 0.9435 | 1 | 1 | 0.9902 | 1 | 1 | 0.7582 | ---NA--- |
| contig_33315_3 | 0.9448 | 1 | 1 | 0.9903 | 1 | 1 | 0.7583 | gametolysin peptidase M11 |
| contig_28323_4 | 0.9459 | 1 | 1 | 0.9903 | 1 | 1 | 0.7583 | leucyl aminopeptidase |
| contig_25894_5 | 0.9463 | 1 | 1 | 0.9903 | 1 | 1 | 0.7583 | predicted protein |
| contig_37722_1 | 0.9464 | 1 | 1 | 0.9903 | 1 | 1 | 0.7583 | hypothetical protein |
| contig_13952_4 | 0.9469 | 1 | 1 | 0.9903 | 1 | 1 | 0.7583 | predicted protein |
| contig_17409_1 | 0.949 | 1 | 1 | 0.9909 | 1 | 1 | 0.7588 | vesicle-associated membrane protein 7b |
| contig_5576_4 | 0.9491 | 1 | 1 | 0.9909 | 1 | 1 | 0.7588 | transketolase |
| contig_24205_6 | 0.9497 | 1 | 1 | 0.9909 | 1 | 1 | 0.7588 | protein fucoxanthin chlorophyll a/c protein |
| contig_26589_3 | 0.9505 | 1 | 1 | 0.9909 | 1 | 1 | 0.7588 | predicted protein |
| contig_25733_5 | 0.9509 | 1 | 1 | 0.9909 | 1 | 1 | 0.7588 | predicted protein |
| contig_7101_4 | 0.9514 | 1 | 1 | 0.9909 | 1 | 1 | 0.7588 | DNA-directed RNA polymerase II 19 kDa polypeptide |

| | | | | | | | | |
|---|---|---|---|---|---|---|---|---|
| contig_25457_5 | 0.9547 | 1 | 1 | 0.9932 | 1 | 1 | 0.7606 | farnesyl-diphosphate farnesyltransferase |
| contig_27589_4 | 0.955 | 1 | 1 | 0.9932 | 1 | 1 | 0.7606 | nucleolar protein 4 |
| contig_5989_3 | 0.9556 | 1 | 1 | 0.9932 | 1 | 1 | 0.7606 | valyl-tRNA synthetase |
| contig_25676_6 | 0.9571 | 1 | 1 | 0.9933 | 1 | 1 | 0.7606 | predicted protein |
| contig_13446_5 | 0.9582 | 1 | 1 | 0.9933 | 1 | 1 | 0.7606 | predicted protein |
| contig_10045_6 | 0.9588 | 1 | 1 | 0.9933 | 1 | 1 | 0.7606 | dihydrolipoamide dehydrogenase |
| contig_27477_6 | 0.9592 | 1 | 1 | 0.9933 | 1 | 1 | 0.7606 | gdp-mannose 4 6-dehydratase |
| contig_25552_1 | 0.9593 | 1 | 1 | 0.9933 | 1 | 1 | 0.7606 | predicted protein |
| contig_28105_5 | 0.9595 | 1 | 1 | 0.9933 | 1 | 1 | 0.7606 | regulatory proteasome non-atpase subunit 3 |
| contig_12107_3 | 0.9623 | 1 | 1 | 0.9945 | 1 | 1 | 0.7615 | argininosuccinate synthase |
| contig_41244_3 | 0.9623 | 1 | 1 | 0.9945 | 1 | 1 | 0.7615 | class II fructose-bisphosphate aldolase |
| contig_27719_4 | 0.9631 | 1 | 1 | 0.9945 | 1 | 1 | 0.7615 | predicted protein |
| contig_27259_5 | 0.9632 | 1 | 1 | 0.9945 | 1 | 1 | 0.7615 | ATP synthase F1 gamma subunit |
| contig_15395_3 | 0.9645 | 1 | 1 | 0.9946 | 1 | 1 | 0.7617 | predicted protein |
| contig_28089_3 | 0.9647 | 1 | 1 | 0.9946 | 1 | 1 | 0.7617 | threonine synthase |
| contig_27417_6 | 0.9665 | 1 | 1 | 0.9958 | 1 | 1 | 0.7626 | ferredoxin-nitrite reductase |
| contig_23921_2 | 0.969 | 1 | 1 | 0.9975 | 1 | 1 | 0.7639 | predicted protein |
| contig_34823_4 | 0.9698 | 1 | 1 | 0.9975 | 1 | 1 | 0.7639 | 60S ribosomal protein L18-2 |
| contig_28923_1 | 0.9706 | 1 | 1 | 0.9975 | 1 | 1 | 0.7639 | predicted protein |
| contig_30187_4 | 0.9713 | 1 | 1 | 0.9975 | 1 | 1 | 0.7639 | adenylosuccinate lyase |
| contig_28327_6 | 0.9726 | 1 | 1 | 0.9975 | 1 | 1 | 0.7639 | rab GDP dissociation inhibitor alpha |
| contig_26362_4 | 0.9735 | 1 | 1 | 0.9975 | 1 | 1 | 0.7639 | predicted protein |
| contig_23117_2 | 0.9739 | 1 | 1 | 0.9975 | 1 | 1 | 0.7639 | peroxisomal (S)-2-hydroxy-acid oxidase GLO5-like |
| contig_33265_1 | 0.9747 | 1 | 1 | 0.9975 | 1 | 1 | 0.7639 | phenylalanine-tRNA ligase beta subunit |
| contig_40249_6 | 0.9752 | 1 | 1 | 0.9975 | 1 | 1 | 0.7639 | translation elongation factor-like protein |
| contig_33797_1 | 0.9769 | 1 | 1 | 0.9975 | 1 | 1 | 0.7639 | glycine-tRNA ligase |
| contig_8539_5 | 0.9777 | 1 | 1 | 0.9975 | 1 | 1 | 0.7639 | SET domain-containing protein |

| | | | | | | | | |
|---|---|---|---|---|---|---|---|---|
| contig_5740_5 | 0.9782 | 1 | 1 | 0.9975 | 1 | 1 | 0.7639 | predicted protein |
| contig_27473_6 | 0.9799 | 1 | 1 | 0.9975 | 1 | 1 | 0.7639 | peptidylprolyl isomerase |
| contig_26987_1 | 0.98 | 1 | 1 | 0.9975 | 1 | 1 | 0.7639 | predicted protein |
| contig_32841_4 | 0.9807 | 1 | 1 | 0.9975 | 1 | 1 | 0.7639 | predicted protein |
| contig_36080_5 | 0.9821 | 1 | 1 | 0.9975 | 1 | 1 | 0.7639 | predicted protein |
| contig_3408_5 | 0.9827 | 1 | 1 | 0.9975 | 1 | 1 | 0.7639 | fucoxanthin-chlorophyll a-c binding protein |
| contig_25602_3 | 0.9827 | 1 | 1 | 0.9975 | 1 | 1 | 0.7639 | protein transport protein SEC61 subunit alpha |
| contig_17279_6 | 0.9846 | 1 | 1 | 0.9975 | 1 | 1 | 0.7639 | vacuolar protein sorting-associated protein 26 |
| contig_34277_5 | 0.9849 | 1 | 1 | 0.9975 | 1 | 1 | 0.7639 | 2-isopropylmalate synthase |
| contig_26918_5 | 0.9854 | 1 | 1 | 0.9975 | 1 | 1 | 0.7639 | predicted protein |
| contig_25846_1 | 0.9854 | 1 | 1 | 0.9975 | 1 | 1 | 0.7639 | glyceraldehyde-3-phosphate dehydrogenase |
| contig_34535_1 | 0.9855 | 1 | 1 | 0.9975 | 1 | 1 | 0.7639 | eukaryotic translation initiation factor 2 subunit 3 |
| contig_25113_4 | 0.9858 | 1 | 1 | 0.9975 | 1 | 1 | 0.7639 | YEATS domain-containing protein 4 |
| contig_12287_5 | 0.9867 | 1 | 1 | 0.9975 | 1 | 1 | 0.7639 | predicted protein |
| contig_34240_2 | 0.9887 | 1 | 1 | 0.9975 | 1 | 1 | 0.7639 | predicted protein |
| contig_17080_5 | 0.9888 | 1 | 1 | 0.9975 | 1 | 1 | 0.7639 | predicted protein |
| contig_38364_2 | 0.9889 | 1 | 1 | 0.9975 | 1 | 1 | 0.7639 | predicted protein |
| contig_18033_5 | 0.9901 | 1 | 1 | 0.9975 | 1 | 1 | 0.7639 | mitochondrial-processing peptidase subunit beta |
| contig_39771_2 | 0.991 | 1 | 1 | 0.9975 | 1 | 1 | 0.7639 | hypothetical protein THAOC_36769 |
| contig_35543_6 | 0.9911 | 1 | 1 | 0.9975 | 1 | 1 | 0.7639 | ferredoxin--NADP+ reductase |
| contig_39611_1 | 0.9912 | 1 | 1 | 0.9975 | 1 | 1 | 0.7639 | methionine-tRNA ligase |
| contig_16347_5 | 0.9912 | 1 | 1 | 0.9975 | 1 | 1 | 0.7639 | predicted protein |
| contig_27061_1 | 0.9923 | 1 | 1 | 0.9975 | 1 | 1 | 0.7639 | acetyl-CoA carboxylase biotin carboxylase subunit |
| contig_30863_3 | 0.9924 | 1 | 1 | 0.9975 | 1 | 1 | 0.7639 | predicted protein |
| contig_21850_6 | 0.9931 | 1 | 1 | 0.9975 | 1 | 1 | 0.7639 | 40S ribosomal protein S14 |
| contig_29420_3 | 0.9937 | 1 | 1 | 0.9975 | 1 | 1 | 0.7639 | thylakoid lumenal 15.0 kDa protein 2 chloroplastic |
| contig_13497_3 | 0.9939 | 1 | 1 | 0.9975 | 1 | 1 | 0.7639 | predicted protein |

| | | | | | | | | |
|---|---|---|---|---|---|---|---|---|
| contig_9982_6 | 0.994 | 1 | 1 | 0.9975 | 1 | 1 | 0.7639 | light harvesting protein |
| contig_24804_5 | 0.9943 | 1 | 1 | 0.9975 | 1 | 1 | 0.7639 | ribosomal protein L13 |
| contig_41396_3 | 0.9954 | 1 | 1 | 0.998 | 1 | 1 | 0.7642 | predicted protein |
| contig_33115_6 | 0.9961 | 1 | 1 | 0.998 | 1 | 1 | 0.7642 | predicted protein |
| contig_33536_4 | 0.9977 | 1 | 1 | 0.999 | 1 | 1 | 0.765 | sec1 family transporter sly1 |
| contig_8878_1 | 0.9984 | 1 | 1 | 0.999 | 1 | 1 | 0.765 | predicted protein |
| contig_25034_6 | 0.9993 | 1 | 1 | 0.9993 | 0.9993 | 1 | 0.7652 | predicted protein |

Supplementary table 2: List of all the enriched GO terms for the three comparisons (Nt25-Nt31, Nt25-Tt34, and Nt31-Tt34) with their corresponding functional category. For each term at each comparison, the populations with more than 75% of proteins upregulated are indicated with the name of the population and the color: Nt25 (dark blue) is non-tolerant at 25ºC, Nt31 (light blue) is non-tolerant at 31ºC, and Tt34 (red) is thermo-tolerant at 34ºC. None indicates that none of the populations for the corresponding comparison has more than 75% of the proteins upregulated for the GO term. C means carbon and N means nitrogen. The original data can be downloaded from the repository specified in the main text of the paper (file: Table_PAPER_GOupregulated_format.xlsx ).

| Functional category | Enriched GO terms | Nt25 vs. Nt31 | Nt25 vs. Tt34 | Nt31 vs. Tt34 |
|---|---|---|---|---|
| C metabolism / Respiration | phosphoenolpyruvate carboxylase activity | | | Nt31 |
| C metabolism | triose-phosphate isomerase activity | | | Nt31 |
| C metabolism | glucose 6-phosphate metabolic process | | Tt34 | |
| C metabolism | fructose 1,6-bisphosphate 1-phosphatase activity | Nt31 | | |
| C metabolism | fructose 1,6-bisphosphate metabolic process | Nt31 | | |
| C metabolism | fructose 6-phosphate metabolic process | Nt31 | | |
| C metabolism | fructose metabolic process | Nt31 | | |
| C metabolism | isomerase activity | Nt31 | | |
| C metabolism | one-carbon metabolic process | Nt31 | | |
| C metabolism | pentose-phosphate shunt | Nt31 | | |
| C metabolism | phosphopyruvate hydratase activity | Nt31 | | |

| | | | | |
|---|---|---|---|---|
| C metabolism | phosphopyruvate hydratase complex | Nt31 | | |
| C metabolism | S-adenosylmethionine metabolic process | Nt31 | | |
| C metabolism | sucrose biosynthetic process | Nt31 | | |
| C metabolism | transaldolase activity | Nt31 | | |
| C metabolism | gluconeogenesis | Nt31 | Tt34 | Nt31 |
| Photosynthesis | glyceraldehyde-3-phosphate dehydrogenase (NAD+) (phosphorylating) activity | Nt31 | Tt34 | |
| C metabolism | glycolytic process | Nt31 | Tt34 | Nt31 |
| C metabolism | transketolase activity | Nt31 | Tt34 | |
| C metabolism / Respiration | acetyl-CoA biosynthetic process | Nt31 | | |
| C metabolism / Respiration | CoA-ligase activity | Nt31 | | |
| Cell organization | cellular component organization | | Tt34 | |
| Cell organization | protein insertion into membrane | | Tt34 | |
| DNA-RNA | FACT complex | | none | |
| DNA-RNA | nucleosome binding | | none | |
| DNA-RNA | nucleotide biosynthetic process | | | Nt31 |

| | | | | |
|---|---|---|---|---|
| DNA-RNA | purine-nucleoside phosphorylase activity | | Tt34 | |
| DNA-RNA | pyrimidine-containing compound biosynthetic process | Nt31 | | |
| DNA-RNA | purine ribonucleoside monophosphate biosynthetic process | Nt31 | | |
| Energy | pyrophosphate hydrolysis-driven proton transmembrane transporter activity | | Tt34 | |
| Energy | proton transmembrane transport | | Tt34 | |
| Energy | proton-transporting ATP synthase complex, catalytic core F(1) | | Tt34 | |
| Energy | ATP hydrolysis activity | Nt31 | Tt34 | |
| Energy | ATP binding | Nt31 | Tt34 | Nt31 |
| Energy | proton motive force-driven ATP synthesis | Nt31 | Tt34 | |
| Energy | proton-transporting ATP synthase activity, rotational mechanism | Nt31 | Tt34 | |
| Fatty acids metabolism | fatty acid ligase activity | | none | |

| | | | |
|---|---|---|---|
| Fatty acids metabolism | fatty acid biosynthetic process | Nt31 | |
| Fatty acids metabolism | fatty acid synthase activity | Nt31 | |
| Fatty acids metabolism | acyl-CoA dehydrogenase activity | | Tt34 |
| Fatty acids metabolism / Proteins biosynthesis / Others | carboxylic acid biosynthetic process | | Tt34 |
| Fatty acids metabolism / Proteins catabolism / Others | carboxylic acid catabolic process | | Tt34 |
| Fatty acids metabolism / Proteins biosynthesis / Others | carboxylic acid binding | Nt31 | |
| N metabolism | nitrate assimilation | | Nt25 |
| N metabolism | nitrate reductase (NADPH) activity | | Nt25 |
| N metabolism | molybdenum ion binding | | Nt25 |
| N metabolism | nitrate transmembrane transport | Nt25 | Nt25 |
| N metabolism | nitrate transmembrane transporter activity | Nt25 | Nt25 |
| N metabolism | nitrite transmembrane transporter activity | Nt25 | Nt25 |
| N metabolism | nitrite transport | Nt25 | Nt25 |

| | | | | |
|---|---|---|---|---|
| N metabolism | glutamate-ammonia ligase activity | Nt31 | | |
| N metabolism | agmatinase activity | Nt31 | Tt34 | |
| N metabolism | nitrogen cycle metabolic process | none | | |
| N metabolism /Stress response | nitric oxide biosynthetic process | | Nt25 | |
| Photosynthesis | organelle outer membrane | | | none |
| Photosynthesis | chloroplast | | | Nt31 |
| Photosynthesis | thylakoid | | | none |
| Photosynthesis | chlorophyll binding | | Tt34 | |
| Photosynthesis | electron transporter, transferring electrons within the cyclic electron transport pathway of photosynthesis activity | | Tt34 | |
| Photosynthesis | photosynthesis, light harvesting in photosystem I | | Tt34 | |
| Photosynthesis | photosynthetic electron transport in photosystem II | | Tt34 | |
| Photosynthesis | photosystem I | | Tt34 | |
| Photosynthesis | photosystem II | | none | |
| Photosynthesis | photosynthetic electron transport chain | Nt31 | | |
| Photosynthesis | pigment biosynthetic process | Nt31 | | |

| | | | | |
|---|---|---|---|---|
| Photosynthesis | reductive pentose-phosphate cycle | Nt31 | | |
| Photosynthesis | magnesium ion binding | Nt31 | | |
| Photosynthesis | porphyrin-containing compound biosynthetic process | Nt31 | | |
| Photosynthesis | chloroplast thylakoid membrane | Nt31 | Tt34 | |
| Photosynthesis | thylakoid lumen | none | | |
| Photosynthesis / Respiration | electron transfer activity | Nt31 | | |
| Proteins biosynthesis | asparagine synthase (glutamine-hydrolyzing) activity | | | Nt31 |
| Proteins biosynthesis | RNA binding | | | Nt31 |
| Proteins biosynthesis | translation | | | Nt31 |
| Proteins biosynthesis | box C/D RNA 3'-end processing | Nt31 | | |
| Proteins biosynthesis | Cajal body | Nt31 | | |
| Proteins biosynthesis | cytoplasmic translation | Nt31 | | |
| Proteins biosynthesis | cytosolic large ribosomal subunit | Nt31 | | Nt31 |
| Proteins biosynthesis | cytosolic small ribosomal subunit | Nt31 | | Nt31 |
| Proteins biosynthesis | nucleus | Nt31 | | |
| Proteins biosynthesis | regulation of translational fidelity | Nt31 | | |

| | | | | |
|---|---|---|---|---|
| Proteins biosynthesis | ribosomal large subunit assembly | Nt31 | | |
| Proteins biosynthesis | ribosomal small subunit assembly | Nt31 | | |
| Proteins biosynthesis | rRNA processing | Nt31 | | |
| Proteins biosynthesis | spliceosomal complex assembly | Nt31 | | |
| Proteins biosynthesis | structural constituent of ribosome | Nt31 | | Nt31 |
| Proteins biosynthesis | GTP binding | Nt31 | | |
| Proteins biosynthesis | GTPase activity | Nt31 | | Nt31 |
| Proteins biosynthesis | histone glutamine methylation | Nt31 | | |
| Proteins biosynthesis | histone glutamine N-methyltransferase activity | Nt31 | | |
| Proteins biosynthesis | isoleucine biosynthetic process | Nt31 | | |
| Proteins biosynthesis | L-aspartate:2-oxoglutarate aminotransferase activity | Nt31 | | |
| Proteins biosynthesis | L,L-diaminopimelate aminotransferase activity | Nt31 | | |
| Proteins biosynthesis | large ribosomal subunit rRNA binding | Nt31 | | |
| Proteins biosynthesis | lysine biosynthetic process via diaminopimelate | Nt31 | | |

| | | | | |
|---|---|---|---|---|
| Proteins biosynthesis | maturation of SSU-rRNA from tricistronic rRNA transcript (SSU-rRNA, 5.8S rRNA, LSU-rRNA) | Nt31 | | |
| Proteins biosynthesis | modification-dependent protein catabolic process | Nt31 | | |
| Proteins biosynthesis | mRNA 5'-UTR binding | Nt31 | | |
| Proteins biosynthesis | poly(A) binding | Nt31 | | |
| Proteins biosynthesis | aminoacyl-tRNA ligase activity | Nt31 | Tt34 | |
| Proteins biosynthesis | translation elongation factor activity | Nt31 | Tt34 | Nt31 |
| Proteins biosynthesis | translational elongation | Nt31 | Tt34 | |
| Proteins biosynthesis | tRNA aminoacylation for protein translation | Nt31 | Tt34 | |
| Proteins catabolism | histidine decarboxylase activity | none | none | |
| Proteins biosynthesis / Transport | protein import into mitochondrial intermembrane space | Nt31 | | |
| Proteins biosynthesis / Transport | protein exit from endoplasmic reticulum | Nt31 | | |
| Proteins catabolism | aminopeptidase activity | | Tt34 | |
| Proteins catabolism | metalloexopeptidase activity | | Tt34 | |

| | | | | |
|---|---|---|---|---|
| Proteins catabolism | proteasomal ubiquitin-independent protein catabolic process | Nt31 | | |
| Proteins catabolism | proteasome core complex | Nt31 | | |
| Proteins catabolism | proteasome-mediated ubiquitin-dependent protein catabolic process | Nt31 | | |
| Proteins catabolism | amino acid catabolic process | Nt31 | | |
| Proteins catabolism | threonine biosynthetic process | Nt31 | | |
| Proteins catabolism | threonine-type endopeptidase activity | Nt31 | | |
| Proteins regulation | methyltransferase activity | Nt31 | | |
| Respiration | cellular respiration | | Tt34 | Nt31 |
| Fatty acids metabolism | fatty acid beta-oxidation using acyl-CoA dehydrogenase | | Tt34 | |
| Respiration | mitochondrion | | Tt34 | |
| Energy | mitochondrial proton-transporting ATP synthase complex, catalytic sector F(1) | Nt31 | | |
| Respiration | tricarboxylic acid cycle | Nt31 | | |
| Respiration | tricarboxylic acid cycle enzyme complex | Nt31 | | |

| | | | | |
|---|---|---|---|---|
| Respiration | tricarboxylic acid metabolic process | Nt31 | | Nt31 |
| Stress response | protein stabilization | | Tt34 | |
| Stress response | removal of superoxide radicals | Nt31 | | |
| Stress response | superoxide dismutase activity | Nt31 | | |
| Stress response | positive regulation of DNA repair | Nt31 | | |
| Stress response | cell redox homeostasis | Nt31 | Tt34 | |
| Stress response | cellular response to heat | Nt31 | Tt34 | |
| Stress response | cellular response to unfolded protein | Nt31 | Tt34 | |
| Stress response | chaperone cofactor-dependent protein refolding | Nt31 | Tt34 | |
| Stress response | heat shock protein binding | Nt31 | Tt34 | |
| Stress response | misfolded protein binding | Nt31 | Tt34 | |
| Stress response | protein folding chaperone | Nt31 | Tt34 | |
| Stress response | protein refolding | Nt31 | Tt34 | |
| Stress response | unfolded protein binding | Nt31 | Tt34 | |
| Transport | endoplasmic reticulum lumen | | Tt34 | |
| Transport | endoplasmic reticulum protein-containing complex | | Tt34 | |
| Transport | establishment of protein localization | | Tt34 | |
| Transport | intracellular protein transport | | Tt34 | |

| | | | | |
|---|---|---|---|---|
| Transport | molecular carrier activity | | Tt34 | |
| Transport | voltage-gated anion channel activity | | Tt34 | |
| Transport | endomembrane system | Nt31 | Tt34 | |
| Transport | inorganic ion transmembrane transport | none | | |
| Transport | transmembrane transport | none | | |
| Others | acid-thiol ligase activity | Nt31 | | Nt31 |
| Others | alcohol metabolic process | | | Nt31 |
| Others | oxidoreductase activity | | | Nt31 |
| Energy | NADPH regeneration | | Tt34 | |
| Photosynthesis | response to light stimulus | | Tt34 | |
| Others | cytosol | | Tt34 | |
| Stress response | cellular response to oxygen-containing compound | Nt31 | | |
| Others | flavin adenine dinucleotide binding | Nt31 | | |
| Others | inositol 2-dehydrogenase activity | Nt31 | | |
| Others | ligase activity, forming nitrogen-metal bonds, forming coordination complexes | Nt31 | | |

| | | | |
|---|---|---|---|
| Energy | oxidoreductase activity, acting on a sulfur group of donors, NAD(P) as acceptor | Nt31 | |
| Others | oxidoreductase activity, acting on the CH-CH group of donors | Nt31 | |
| Energy | oxidoreductase activity, acting on the CH-OH group of donors, NAD or NADP as acceptor | Nt31 | |
| Others | oxidoreductase complex | Nt31 | |
| Others | pyridoxal phosphate binding | Nt31 | |
| Energy | NAD binding | Nt31 | Tt34 |
| Energy | NADP binding | Nt31 | Tt34 |
| Others | plasma membrane | none | Tt34 |
